# Complex genetic variation in nearly complete human genomes

**DOI:** 10.1101/2024.09.24.614721

**Authors:** Glennis A. Logsdon, Peter Ebert, Peter A. Audano, Mark Loftus, David Porubsky, Jana Ebler, Feyza Yilmaz, Pille Hallast, Timofey Prodanov, DongAhn Yoo, Carolyn A. Paisie, William T. Harvey, Xuefang Zhao, Gianni V. Martino, Mir Henglin, Katherine M. Munson, Keon Rabbani, Chen-Shan Chin, Bida Gu, Hufsah Ashraf, Olanrewaju Austine-Orimoloye, Parithi Balachandran, Marc Jan Bonder, Haoyu Cheng, Zechen Chong, Jonathan Crabtree, Mark Gerstein, Lisbeth A. Guethlein, Patrick Hasenfeld, Glenn Hickey, Kendra Hoekzema, Sarah E. Hunt, Matthew Jensen, Yunzhe Jiang, Sergey Koren, Youngjun Kwon, Chong Li, Heng Li, Jiaqi Li, Paul J. Norman, Keisuke K. Oshima, Benedict Paten, Adam M. Phillippy, Nicholas R Pollock, Tobias Rausch, Mikko Rautiainen, Stephan Scholz, Yuwei Song, Arda Söylev, Arvis Sulovari, Likhitha Surapaneni, Vasiliki Tsapalou, Weichen Zhou, Ying Zhou, Qihui Zhu, Michael C. Zody, Ryan E. Mills, Scott E. Devine, Xinghua Shi, Mike E. Talkowski, Mark J.P. Chaisson, Alexander T. Dilthey, Miriam K. Konkel, Jan O. Korbel, Charles Lee, Christine R. Beck, Evan E. Eichler, Tobias Marschall

## Abstract

Diverse sets of complete human genomes are required to construct a pangenome reference and to understand the extent of complex structural variation. Here, we sequence 65 diverse human genomes and build 130 haplotype-resolved assemblies (130 Mbp median continuity), closing 92% of all previous assembly gaps^1,2^ and reaching telomere-to-telomere (T2T) status for 39% of the chromosomes. We highlight complete sequence continuity of complex loci, including the major histocompatibility complex (MHC), *SMN1*/*SMN2*, *NBPF8*, and *AMY1/AMY2*, and fully resolve 1,852 complex structural variants (SVs). In addition, we completely assemble and validate 1,246 human centromeres. We find up to 30-fold variation in α-satellite high-order repeat (HOR) array length and characterize the pattern of mobile element insertions into α-satellite HOR arrays. While most centromeres predict a single site of kinetochore attachment, epigenetic analysis suggests the presence of two hypomethylated regions for 7% of centromeres. Combining our data with the draft pangenome reference^1^ significantly enhances genotyping accuracy from short-read data, enabling whole-genome inference^3^ to a median quality value (QV) of 45. Using this approach, 26,115 SVs per sample are detected, substantially increasing the number of SVs now amenable to downstream disease association studies.

## Introduction

Genome assemblies based on highly accurate long reads, e.g., Pacific Biosciences (PacBio) high-fidelity (HiFi) reads, now almost completely resolve human genomes with excellent sequence quality and contiguity^4,5^. This technology has enabled the comprehensive characterization of structural variants (SVs), defined as variants 50 bp in length or longer, and revealed that more than half of all SVs are missed using short-read discovery^6^. Such genome assemblies have made a first draft human pangenome reference possible, which was constructed by building a graph representation reflecting an alignment of haplotype-resolved genome assemblies^1^.

Despite these advances, genome assemblies based on a single long-read sequencing technology are presently not able to contiguously assemble whole chromosomes, resulting in systematic gaps at difficult loci^2^. Closing these gaps requires combining the complementary strengths of PacBio HiFi reads (∼18 kbp in length and high base-level accuracy) and ultra-long (UL) Oxford Nanopore Technologies (ONT) reads (>100 kbp in length but with lower base-level accuracy). Combining both types of reads yielded the first gapless assembly of a human genome^7^ and, since then, the hybrid assembly process has been automated with the tools Verkko^8^ and hifiasm (UL)^9^, but so far assemblies of this quality have been restricted to few individual samples. Here, we target a diverse set of 65 samples predominantly from the 1000 Genomes Project (1kGP) cohort^10^ and assemble 39% of chromosomes to telomere-to-telomere (T2T) status. This set of haplotype-resolved hybrid assemblies provides a genomic resource that routinely closes more than 92% of gaps previously characterized in PacBio HiFi-only assemblies^2^. Moreover, our high-quality assemblies enable insights into many structurally variable but previously intractable regions of the human genome, including centromeres and disease-associated regions [e.g., major histocompatibility complex (MHC), *SMN1/SMN2*, and *AMY1/AMY2*].

## Results

### Production of 130 phased genome assemblies

#### Data production

We selected 65 human samples spanning five continental groups and 28 population groups for sequencing (**Fig. 1a**). These samples include 63 from the 1kGP, one from the International HapMap Project^11^, and another from the Genome in a Bottle effort^12^. The complete set is composed of 30 males and 35 females of either African (*n*=30), Admixed American *(n*=9), European (*n*=8), East Asian (*n*=10) or South Asian (*n*=8) descent and includes three parent–child trios (**Supplementary Table 1**).

**Figure 1.**
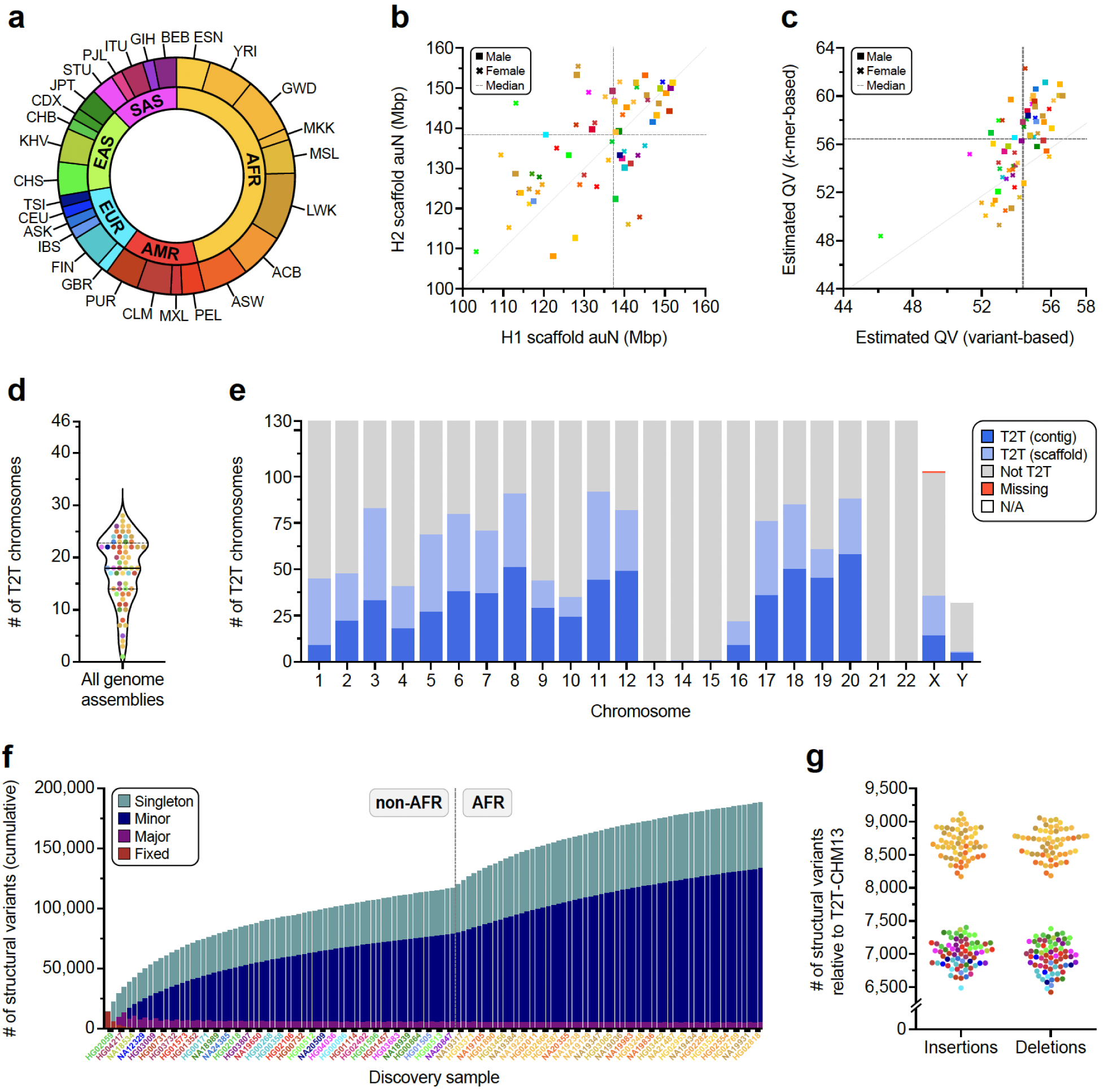
Long-read sequencing, assembly, and variant calling of 65 diverse human samples. **a)** Continental group (inner ring) and population group (outer ring) of the 65 diverse human samples analyzed in this study. **b)** Scaffold auN for haplotype 1 (H1) and haplotype 2 (H2) contigs from each genome assembly. Data points are color-coded by population and sex. Dashed lines indicate the median auN per haplotype. The dotted line indicates the unit diagonal. **c)** QV estimates for each genome assembly derived from variant calls or *k*-mer statistics **(Methods)**. **d)** The number of chromosomes assembled from telomere-to-telomere (T2T) for each genome assembly, including both single contigs and scaffolds **(Methods)**. The median (solid line) and first and third quartiles (dotted lines) are shown. **e)** The number of T2T chromosomes in a single contig (dark blue, T2T contig) or in a single scaffold (light blue, T2T scaffold). Incomplete chromosomes are labeled as “Not T2T” or “Missing” if missing entirely. Sex chromosomes not present in the respective haploid assembly are labeled as “N/A”. **f)** Cumulative nonredundant structural variants (SVs) across the diverse haplotypes in this study called with respect to the T2T-CHM13 reference genome (three trio children excluded). **g)** Number of SVs detected for each haplotype relative to the T2T-CHM13 reference genome, colored by population. Insertions and deletions are balanced when called against the T2T-CHM13 reference genome but imbalanced when called against the GRCh38 reference genome (**Extended Data Fig. 1d**).

**Extended Data Figure 1.**
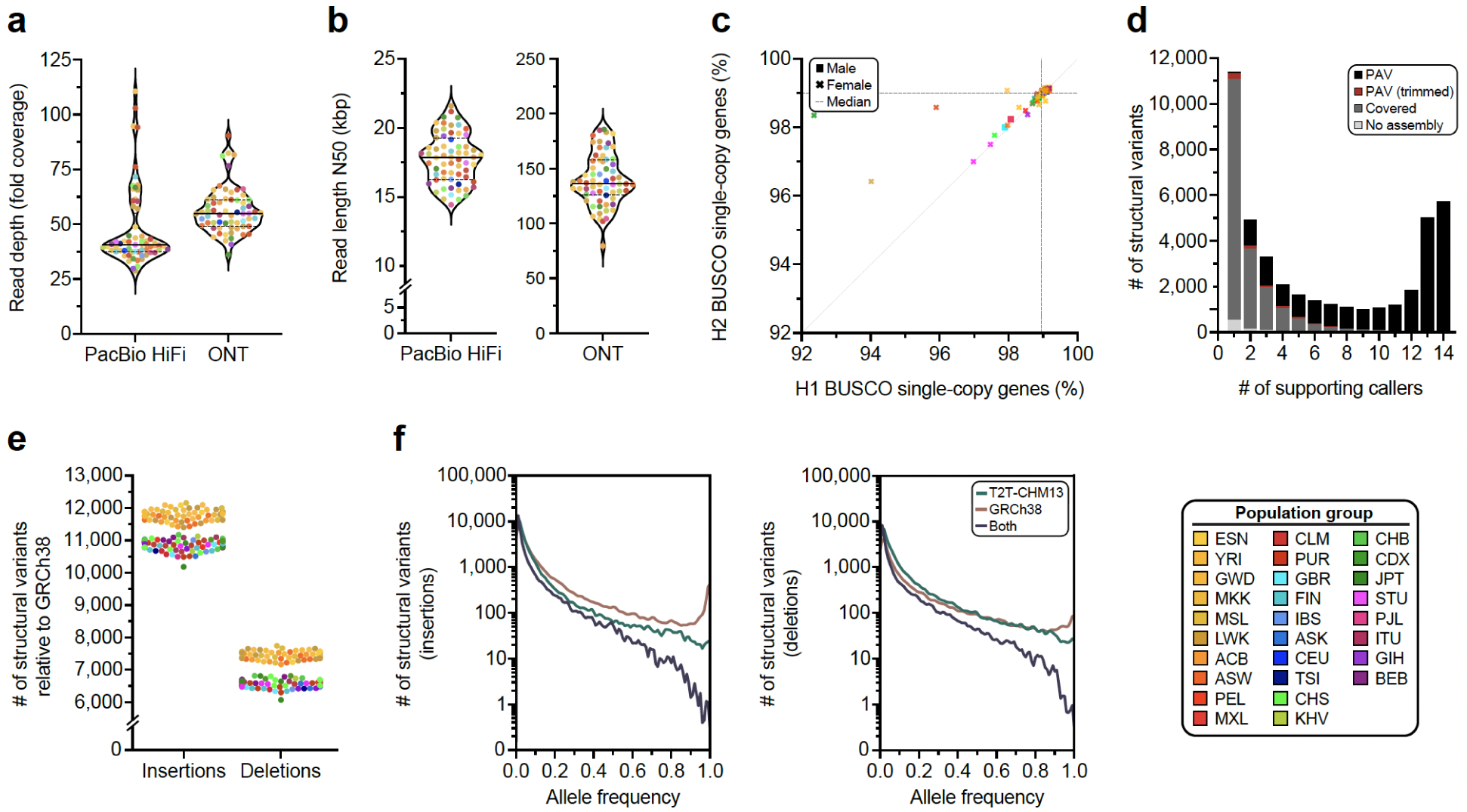
Statistics of long-read sequencing data and genome assemblies generated in this study as well as variant calls for 65 diverse human genomes. **a)** Fold coverage of the Pacific Biosciences (PacBio) high-fidelity (HiFi) and Oxford Nanopore Technologies (ONT) long-read sequencing data generated for each genome in this study. The median (solid line) and first and third quartiles (dotted lines) are shown. **b)** Read length N50 of the PacBio HiFi and ONT data generated for each genome in this study. The median (solid line) and first and third quartiles (dotted lines) are shown. **c)** Gene completeness as a percentage of BUSCO single-copy orthologs detected in each haplotype from each genome assembly **(Methods)**. **d)** The number of structural variants (SVs) detected by the Phased Assembly Variant (PAV) caller. Before applying caller-based QC, 99.75% of PAV calls are supported by at least one other call source. PAV, variant supported by PAV; PAV (trimmed), variant was removed when PAV trimmed repetitive bases mapped multiple times; Covered, region covered by an assembly, but no comparable SV found by PAV; No Assembly, SV occurs in a region where an assembly sequence was not aligned. **e)** Number of SVs called for each haplotype relative to the GRCh38 reference genome, colored by population. Insertions and deletions are imbalanced when called against the GRCh38 reference genome but balanced when called against the T2T-CHM13 reference genome (Fig. 1g). **f)** Number of SV insertions (left) and deletions (right) called against the T2T-CHM13 reference genome, GRCh38 reference genome, or both relative to their allele frequency. SVs called against both references tend to be more rare because they are less likely to appear in a reference genome. A sharp peak for high allele frequency (∼1.0) for insertions is detected relative to the GRCh38 reference genome but not the T2T-CHM13 reference genome.

We compiled a rich data resource to produce haplotype-resolved genome assemblies and capture large SVs, complex loci, and other genomic features (**Methods**). We generated ∼47-fold coverage of PacBio HiFi and ∼56-fold coverage of ONT (∼36-fold ultra-long) long reads on average per sample (**Extended Data Fig. 1a,1b; Supplementary Table 2**). In addition, we performed single-cell template strand sequencing (Strand-seq) (**Supplementary Table 2**), Bionano Genomics optical mapping (**Supplementary Table 3**), Hi-C sequencing (**Supplementary Tables 4,5**), isoform sequencing (Iso-Seq; **Supplementary Table 6**), and RNA sequencing (RNA-seq; **Supplementary Table 7**).

#### Assembly

We generated haplotype-resolved assemblies from all 65 diploid samples using Verkko^8^ as a current state-of-the-art approach for hybrid genome assembly (**Fig. 1a, Supplementary Tables 1,2**; **Methods**). The phasing signal for the assembly process was produced with Graphasing^13^, leveraging Strand-seq to globally phase assembly graphs at a quality on par with trio-based workflows^13^ (**Methods**). This approach can, therefore, be applied to samples without parental sequencing data, which enabled us to cover all 26 populations from the 1kGP by including samples that are not part of a family trio. The resulting set of 130 haploid assemblies is highly contiguous (median area under the Nx curve [auN] of 137 Mbp, **Fig. 1b**, **Supplementary Table 8**) and accurate at the base-pair level (median QV between 54 and 61, **Fig. 1c**, **Supplementary Table 9**, **Methods**). Using established methods^14^, we estimate the assemblies to be 99% complete (median) for known single-copy genes (**Extended Data Fig. 1c, Supplementary Table 10**). Combining two complementary long-read technologies also enabled us to reliably assemble and close 92% of previously reported gaps in PacBio HiFi-only assemblies^2^ (**Supplementary Figs. 1,2; Supplementary Table 11**) (**Methods**) and fully resolve biomedically important regions, such as the MHC locus for 128/130 assembled haplotypes.

We integrated a range of quality control (QC) annotations for each assembly using established tools such as Flagger^1^, NucFreq^15^, Merqury^16^, and Inspector^17^ (**Supplementary Figs. 3,4; Supplementary Tables 12,13**) to compute robust error estimates for each assembled base (**Supplementary Tables 14 -17**, **Methods**). We, thus, estimate that 99.6% of the phased sequence (median) has been assembled correctly (**Supplementary Table 18**). For the three family trios in our dataset (SH032, Y117, PR05^10^), we assessed the parental support for the respective haplotypes in the child’s assembly via assembly-to-assembly alignments and found that a median of 99.8% of all sequence assembled in contigs >100 kbp are supported by one parent assembly (**Supplementary Table 19**, **Methods**). In total, Verkko assembled 599 chromosomes as a single gapless contig from telomere to telomere (median 10 per genome) and an additional 558 as a single scaffold (median 18 per genome), i.e., in a connected sequence containing one or more N-gaps (**Figs. 1d,1e**; **Supplementary Fig. 5, Supplementary Table 20**) (**Methods**).

Despite the substantial improvements in assembly quality compared to single-technology approaches, certain regions, such as centromeres or the Yq12 region, remained challenging to assemble and evaluate. We, thus, complemented our assembly efforts by running hifiasm (UL)^9^ on the same dataset (**Supplementary Figs. 6,7; Supplementary Tables 21,22; Methods**). We observed an overall lower contiguity compared to Verkko and several cases of spurious duplications in the hifiasm assemblies (**Supplementary Table 23**) and, therefore, restricted their use to extending our analysis set for centromeres and the Yq12 region after manual curation of the relevant sequences.

#### Variant calling

From our phased assemblies, we identify 188,500 SVs, 6.3 million indels, and 23.9 million single-nucleotide variants (SNVs) against the T2T-CHM13v2.0 (T2T-CHM13) reference (**Fig. 1f**). Against GRCh38-NoALT (GRCh38), we identify 176,531 SVs, 6.2 million indels, and 23.5 million SNVs (**Supplementary Table 24**, **Data Availability**). Callset construction was performed similarly for both references and was led by PAV (v2.4.0.1)^6^ with orthogonal support from 10 other independent callers with sensitivity for SVs (eight callers) and indels and SNVs (three callers) (**Supplementary Table 25**; **Methods**). We find higher support for PAV calls across all callers (99.7%) compared to other methods (99.7% to 67.9%) (**Extended Data Fig. 1d**) (**Supplementary Table 26**). With one additional sample, we estimate our callset would increase by 842 SV insertions and deletions with a 1.86× enrichment for an African versus a non-African sample (1,117 vs. 599) (**Supplementary Methods**).

Compared with our previous resource derived from 32 phased human genome assemblies^6^, our current callset yields 1.6× more SV insertions and deletions (177,718 vs. 111,679), which increases to 3.5× for SVs greater than 10 kbp (4,541 vs. 1,299) with improvements in assembly contiguity (**Supplementary Table 27**). Mendelian inheritance error (MIE) has also decreased (**Supplementary Table 28**) with 2.7% MIE for SV insertions and deletions (55% decrease), 5.1% MIE for inversions, 4.9% MIE for indels (5.5% decrease), and 0.6% MIE for SNVs (79% decrease).

Per assembled haplotype, we identify 7,772 SV insertions and 7,745 SV deletions in the T2T-CHM13 reference (**Fig. 1g**). As expected, GRCh38 SVs are unbalanced^6,18,19^ with 11,275 SV insertions and 6,972 SV deletions per haplotype on average (**Extended Data Fig. 1e**) with excess insertions occurring in high allele-frequency variants, which can be largely explained by reference errors^20^. While GRCh38 yields more SVs per sample compared to T2T-CHM13 (**Supplementary Table 29**), the T2T-CHM13 callset across all samples contains 6% more SVs, 2% more indels, and 2% more SNVs highlighting the impact of a more complete T2T reference (**Supplementary Table 24**). We further linked variants across both references through assembly coordinates (**Supplementary Table 30**, **Methods**). We find the number of fixed variants in the callset is significantly enriched for insertions versus deletions in GRCh38 (*p*=7.88×10^-50^, Fisher’s exact test [FET]), but not T2T-CHM13 (*p*=0.40, FET), which holds when tandem repeats are excluded (GRCh38 *p*=6.47×10^-27^, T2T-CHM13 *p*=0.16, FET). Because rare alleles are less likely to become reference alleles, SVs identified in both references have lower allele frequencies on average (6.5% insertions, 7.8% deletions) compared to variants present in only T2T-CHM13 (8.3% insertions, 11.1% deletions) or GRCh38 (13.3% insertions, 14.7% deletions). As expected, a distinct peak for fixed SVs (100% AF) is apparent for GRCh38 SV insertions composed of variants in GRCh38 with no representation in T2T-CHM13 (**Extended Data Fig. 1f**). A similar but smaller peak is apparent for GRCh38 deletions **(Extended Data Fig. 1f)**, likely attributable to false duplications in GRCh38.

### An improved genomic resource

Our publicly available haplotype-resolved assemblies and the associated SV callset constitute a resource for the genomics community that is significantly improved when compared to existing genome resources. Examples of relevant improvements are detailed below.

#### Mobile Element Insertions

Mobile element insertions (MEIs) continue to diversify human genomes^21^ and constitute a significant portion of SVs (∼13% of SVs in this study). To identify MEIs in the 130 haplotype assemblies, we generated a union dataset from two independent MEI callsets each for both T2T-CHM13 (13,216 putative MEIs) and GRCh38 (13,001 putative MEIs) (**Methods**). All putative MEIs identified by a single caller were manually curated (**Methods**) (**Supplementary Figs. 8,9; Supplementary Tables 31,32**). Additionally, our comparison with an orthogonal callset showed a high average concordance of 90.6% (T2T-CHM13: 90.4%, GRCh38: 90.8%) (**Methods, Supplementary Tables 33,34**). The T2T-CHM13 callset (*Alu*: 10,689, L1: 1,718, SVA: 803, HERV-K: 5, snRNA: 1) and GRCh38 dataset (*Alu*: 10,522, L1: 1,702, SVA: 770, HERV-K: 6, snRNA: 1) had similar MEI compositions. Moreover, we find 569 full-length L1 insertions with respect to T2T-CHM13 and 551 with respect to GRCh38. Of these, an average of 96.4% possess at least one intact open reading frame (ORF) and 82.4% harbor two intact ORFs. Therefore, the vast majority of full-length L1 MEIs appear to retain the potential to retrotranspose. Compared to the MEI datasets of the previous HGSVC publication callset^6^ (n=9,453 MEIs; *Alu*: 7,738, L1: 1,775, and SVA: 540), the number of MEIs increased by more than 35% (T2T-CHM13: 39.74%; GRCh38: 37.45%). Next, we screened the PAV deletion callsets and identified 2,450 polymorphic MEIs present in T2T-CHM13 and 2,468 MEIs in GRCh38 (**Supplementary Fig. 10, Supplementary Tables 35,36**). Through a combination of high-quality haplotype-resolved assemblies and more sensitive MEI callers, we are now able to investigate MEIs in complex regions, resulting in the identification of more MEIs per sample.

#### Inversions

Identifying inversions is challenging due to the frequent location of their boundaries in long, highly identical repeat sequences. We identify 276 T2T-CHM13–based and 298 GRCh38-based inversions in the main callset. In order to perform validation of the PAV inversion calls, re-genotyping was performed using ArbiGent on Strand-seq data^22^. For T2T-CHM13–based inversions, ArbiGent was able to provide an assessment for 252 inversion calls, labeling 218 as inversions, 26 as inverted duplications, and 2 as potential misorientations (**Supplementary Table 37**, **Supplementary Methods**). To ensure accuracy of these challenging SV calls, we performed manual inspection (**Supplementary Fig. 11**)^23^ to estimate a false discovery rate (FDR) of 12.33% (**Supplementary Fig. 12**). Of the calls passing manual inspection, 97% were labeled as inversions or inverted duplications by ArbiGent (**Supplementary Tables 37,38**). To assess inversion detection sensitivity, we re-genotyped the previously reported T2T-CHM13– and GRCh38-based inversion callsets, which used a subset of 41 samples^22,24^ (**Supplementary Tables 39,40**). We observed that 179 out of 295 balanced inversions in T2T-CHM13 were identified by PAV based on requiring a 25% reciprocal overlap. There are only 76 inversions with no significant overlap with the current PAV callset, of which 42 are longer than 10 kbp, corresponding to an improved ability to detect inversions directly from assemblies compared to earlier studies^22^. Notably, we find 21 novel inversions in the PAV callset, of which 18 are detected among 24 new samples added in the current study. These include a large (1.8 Mbp) inversion at Chromosome 5q35 that overlaps with the Sotos syndrome critical region^25,26^.

#### Segmental duplication and copy number polymorphic genes

Because segmental duplications (SDs) are enriched 10-fold for copy number variation (CNV) and are the source of some of the most complex forms of genic structural polymorphism in the human genome^30,31^, we systematically assessed SD content (>1 kbp and >90% identity) for each of the 130 human genomes and then focused on the structure and content of some of the most gene-rich polymorphic regions. Overall, we find an average of 168.1 Mbp (standard deviation [s.d.], 9.2 Mbp) of SDs per human genome. As expected, there is an improved representation of interchromosomal SDs (**Supplementary Fig. 13**) when compared to the Human Pangenome Reference Consortium (HPRC) release^1^. Both the length and percent identity distributions for both intrachromosomal and interchromosomal SDs are highly reproducible with modes at 99.4% (s.d., 0.03%) and 95.0% (s.d., 0.4%) identity, respectively (**Supplementary Fig. 14**). Using T2T-CHM13 as a gauge of completeness, we estimate that 25.6 Mbp of SDs still remain unresolved per haploid genome (**Fig. 2a**). Most of these unresolved SDs (21.2 Mbp) correspond to the acrocentric short arms of Chromosomes 13, 14, 15, 21, and 22, which are known to be among the most difficult regions to assemble due to extensive ectopic recombination and the highest degree of sequence homology^7,32^. Since high-identity SDs are also a common source of misassembly^2^, we also assessed SDs for evidence of collapse and confirmed copy number using read depth (**Methods**). We find that 80-90% of SDs are accurately assembled depending on the genome (**Supplementary Fig. 15**).

**Figure 2.**
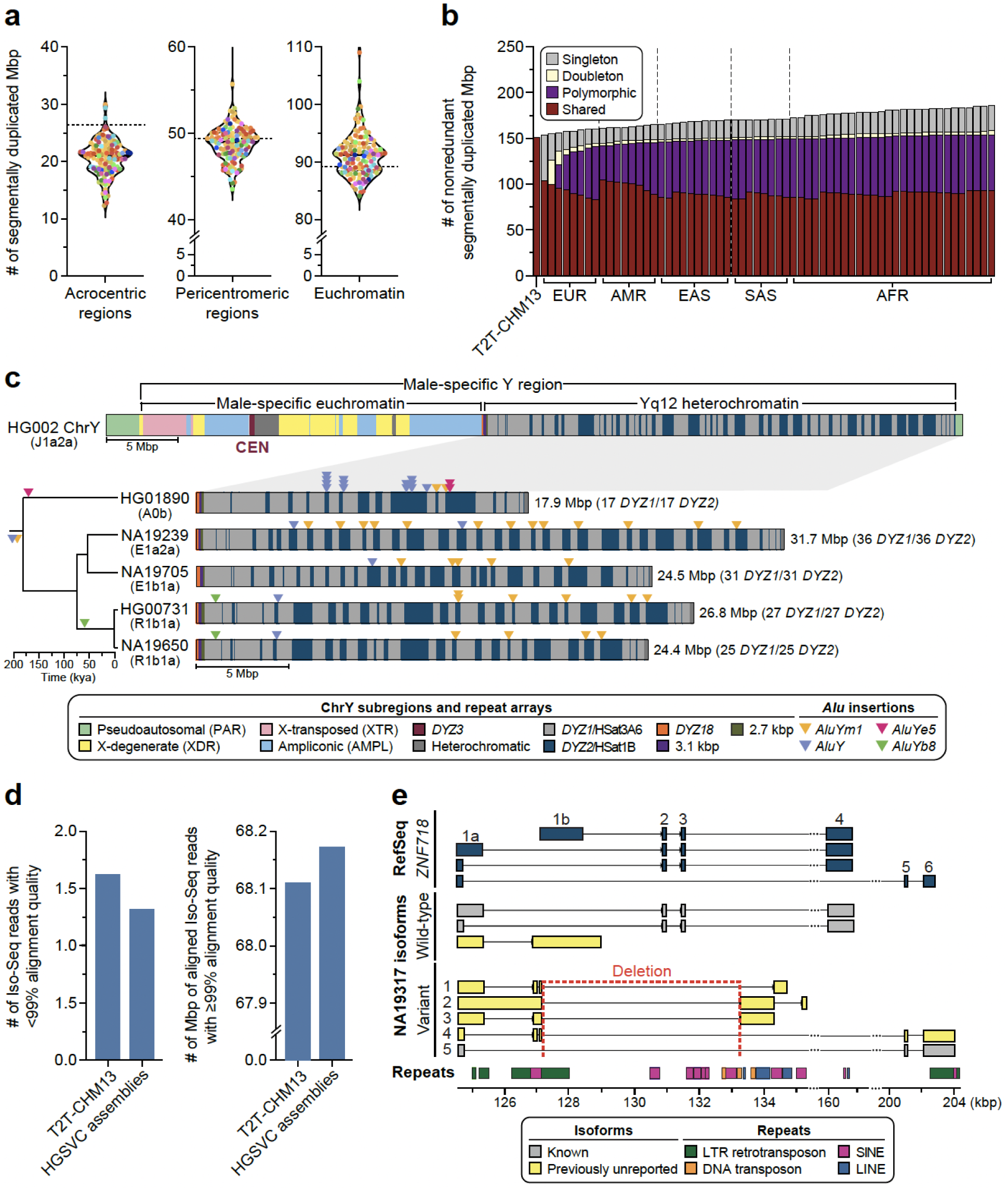
An improved genomic resource for challenging loci. **a)** Number of segmentally duplicated bases assembled in different regions of the genome for each sample in this study, excluding sex chromosomes. The dashed line indicates the number of segmentally duplicated bases in the T2T-CHM13 genome. **b)** Segmental duplication (SD) accumulation curve. Starting with T2T-CHM13, the SDs (excluding those located in acrocentric regions and chrY) of 63 samples (excluding NA19650 and NA19434) were projected onto T2T-CHM13 genome space in the continental group order of: EUR, AMR, EAS, SAS and AFR. For each bar, the SDs that are singleton, doubleton, polymorphic (>2) and shared (>90%) are indicated. **c)** Structure of a human Y chromosome on the basis of T2T-CHM13 chromosome Y reference sequence, including the centromere (CEN; top). On the bottom, repeat composition of four contiguously assembled Yq12 heterochromatic regions with their phylogenetic relationships shown on the left. The size of the region and the number of *DYZ1* and *DYZ2* repeat array blocks are shown on the right. Locations of four inserted and subsequently amplified *Alu* elements on Yq12 are shown as triangles. **d)** Comparison of total Iso-Seq reads that failed to align at ≥99% accuracy for T2T-CHM13 vs. the assemblies in this study (left), and comparison of total bases aligned to T2T-CHM13 vs. the assemblies in this study among reads that aligned to both at ≥99% accuracy (right). **e)** Expressed isoforms of *ZNF718* identified in NA19317. This individual is heterozygous for a deletion that impacts the exon-intron structure of *ZNF718* (deleting exons 2 and 3 and part of the alternate first exon 1b). Repeat classes are annotated by color at the bottom. The wild-type allele harbors a single, previously unreported isoform consisting of a canonical first exon and second exon that is typically reported as alternate first exon 1b (yellow, wild-type). The presence of the 6,142 bp long deletion on chr4:127,125-133,267 is associated with four isoforms not previously annotated in RefSeq^27^, GENCODE^28^, or CHESS^29^ (variant, yellow). All four novel isoforms begin at the canonical transcription start site, contain part of exon 1b, and lack canonical exons 2 and 3.

Focusing on SDs that passed QC (not flagged by NucFreq or Flagger), we observe that African genomes show higher SD content when compared to those of non-African origin (**Supplementary Fig. 16**), confirming a recent report^33^. Next, we performed a pangenome analysis using 10 kbp of flanking unique sequence as an anchor to project onto the T2T-CHM13 genome (**Supplementary Fig. 17**, **Methods**). As a result, we classify at least 92.8 Mbp of the SDs as shared among most humans (present in at least 90% of samples) and 61.0 Mbp as variable across the human population (**Fig. 2b**). In addition, we identify 33 Mbp of SD sequence present in a single copy or not annotated as SDs in T2T-CHM13. (**Extended Data Fig. 2**). The majority of these (23.6 Mbp, including 2.4 Mbp of chrX SDs) are novel when compared to a recent analysis of 170 human genomes^33^ and completely or partially overlap with 167 protein-coding genes (**Supplementary Fig. 18**). Predictably, these 33 Mbp represent rare SDs [19.2 Mbp (*n*=626) singleton or 3.8 Mbp (*n*=171) doubleton loci] or variable loci (9.8 Mbp/531 loci). Notably, 31 loci (0.4 Mbp) are shared among most humans but not classified as duplicated in the T2T-CHM13 human genome suggesting that this unique status in the reference represents the minor allele in the human population, a cell line artifact or, less likely, an error in the assembly. Examining genomes by continental group, the number of new SDs added per sample (diploid) is highest for Africans (3.97 Mbp/individual) when compared to non-African samples (2.88 Mbp/ individual).

Using GENCODE annotation^28^, we searched specifically for protein-coding genes that are copy number variable and >2 kbp long (**Methods**). Genomes with African ancestry harbor, on average, 468 additional paralogous genes (*n*=21,595 total genes) when compared to non-African (*n*=21,127 total genes), consistent with SD data. We identify a total of 727 multi-copy genes that have SDs spanning at least 90% of the gene body, with a large proportion corresponding to shared (*n*=335 or 46.1%) and variable (*n*=292 or 40.2%) SDs (**Supplementary Table 41**). In terms of gene copy number, we observed variation in the majority of these multi-copy genes (*n*=626 or 86.1%). Comparing the copy numbers to the HPRC assemblies^1^, we discover a similar distribution of genes (**Supplementary Fig. 19**). Among copy number polymorphic genes, we identify 16 gene families where the distribution significantly differs between the HPRC and our data (**Supplementary Fig. 19**) (adjusted p<0.05, two-sided Welch’s t-test); however, the contiguity for copy number variant genes was considerably greater in our assemblies versus HPRC; 5.88% of duplicated genes in our assemblies are within 200 kbp of a contig break or unknown base (“N”) compared to 13.95% of duplicated genes in HPRC assemblies (**Supplementary Fig. 20**).

**Extended Data Figure 2.**
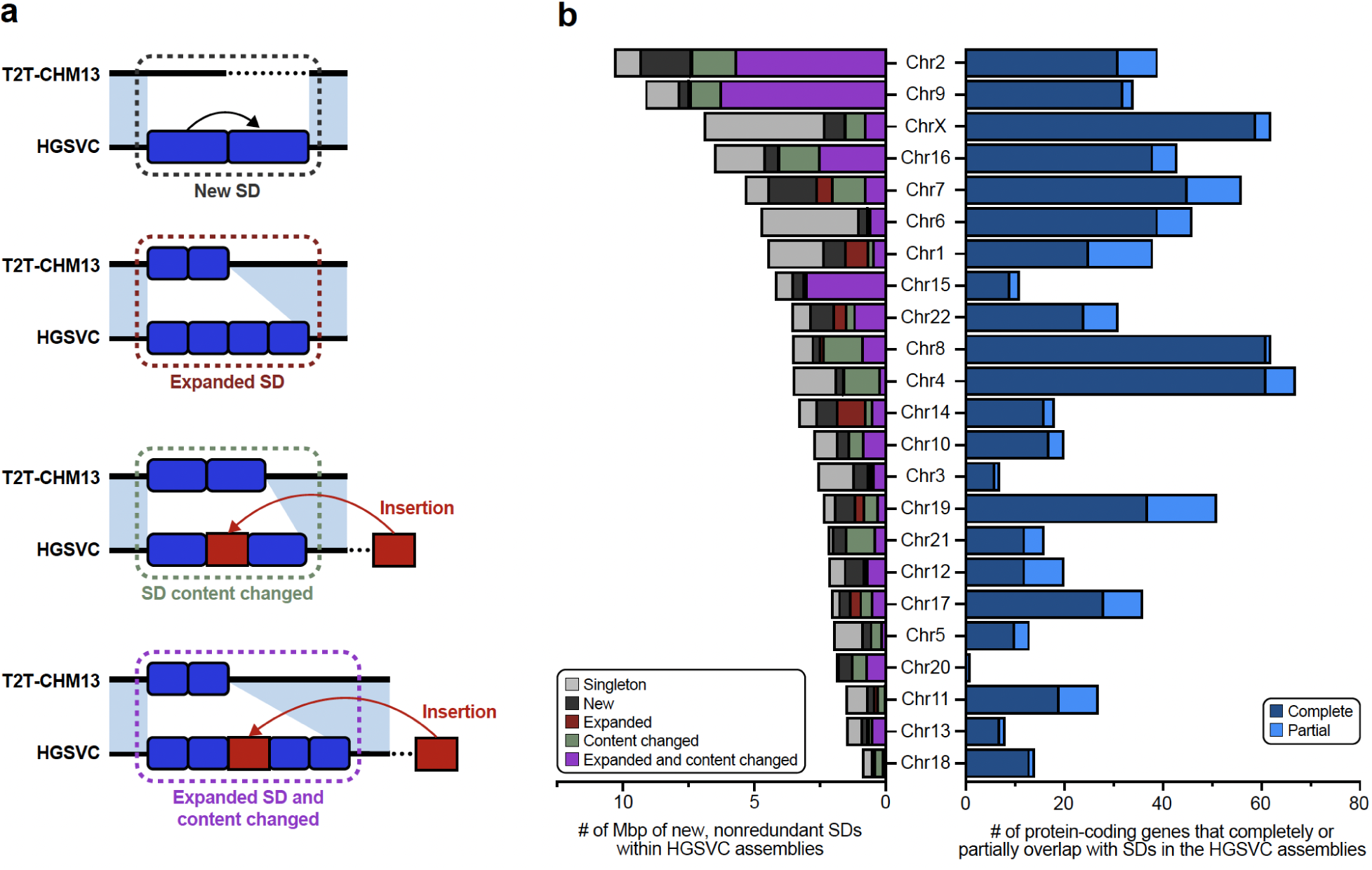
Classification and distribution of changes in SD content in the 65 genomes. **a)** Schematic depicting the four categories of non-reference SDs: 1) new (i.e., unique in the reference), 2) expanded copy number, 3) content or composition changed, and 4) expanded and content changed SDs with respect to the SDs in the reference genome, T2T-CHM13. **b)** Quantification in terms of Mbp and predicted protein-coding genes across the four categories of new SDs compared to T2T-CHM13. The left panel shows the Mbp by category, while flagging those that are singleton (i.e., duplicated in T2T-CHM13 but not in other genomes). The right panel quantifies the number of complete (100% coverage) and partial overlaps (>50% coverage) with protein-coding genes for the respective chromosomes.

#### Y chromosome variation

The Y chromosome remains among the most challenging human chromosomes to fully assemble due to its highly repetitive sequence composition^32,34^ (**Fig. 2c**). Our resource provides highly contiguous Y assemblies for 30 male samples. Seven of these (23%) assembled without breaks across the male-specific Y region (excluding the pseudoautosomal regions, six assembled as T2T scaffolds, and one has a break in the pseudoautosomal region 1), (**Supplementary Figs. 21,22**). Of these seven, four are novel fully assembled human Y chromosomes representing E1b1a, R2a and R1b1a Y lineages prevalent in populations of African, Asian, and European descent^35^. Combined with previous reports, we increase the total number of completely assembled human Y chromosomes to eight (**Supplementary Fig. 23**)^36,37^.

Our assemblies enable the investigation of the largest heterochromatic region in the human genome, Yq12, mostly composed of highly similar (but size variable) alternating arrays of *DYZ1* (*HSat3A6*, ∼3.5 kbp unit size) and *DYZ2* (*HSat1B*, ∼2.4 kbp unit size) repeats (**Fig. 2c**). The Yq12 regions across 16 samples (9 novel and 7 previously published) ranged from 17.85–37.39 Mbp (mean 27.25 Mbp, median 25.62 Mbp), with high levels of variation in the number (34 to 86 arrays; mean 60, median 58) and length of *DYZ1* (24.4 kbp to 3.59 Mbp; mean 525.7 kbp, median 455.0 kbp) and *DYZ2* (11.2 kbp to 2.20 Mbp; mean 358.0 kbp, median 273.3 kbp) repeat arrays (**Supplementary Fig. 24, Supplementary Table 42**)^36,37^. Investigating the dynamics of Yq12 remains challenging^38^; however, using the duplication and deletion patterns of four unique *Alu* insertions, we can examine this genomic region over time (**Fig. 2c, Supplementary Fig. 24**). For example, in NA19239, the presence of retrotransposon insertions (one *Alu*Y and five *Alu*Ym sequences) allows clear visualization of a tandem duplication in the region.

#### SVs disrupting genes

To examine how the SVs in our resource can better identify novel loss-of-function (LoF) variants in protein-coding genes, we intersected all 176,531 GRCh38 SVs with coding exons and find 822 unique genes overlapping with 1,308 unique SVs. The coding sequences of an average of 181 genes are disrupted by at least one SV breakpoint per genome. Most gene disruptions result from polymorphic variants, with an average of 10 genes disrupted by singleton variants in each genome. Of the 822 genes affected by SVs, 18 are annotated as intolerant to LoF in humans as measured by the loss-of-function observed/expected upper-bound fraction (LOEUF^39^ <0.35). Seven of these constrained genes are altered by polymorphic SVs (in more than 1 of 65 samples). Common SV mutations altering constrained genes suggests that the variants do not result in LoF of the gene. Indeed, we found tandem repeat unit variants in coding sequences of four constrained genes (*MUC5B*, *ACAN*, *FNM2, ARMCX4*). Three additional constrained genes have partially duplicated exons that preserve the coding frame. We also observe deletions of one or more 59 bp VNTR units that overlap the last 8 bp of exon 37 of *MUC5B,* but the sequences of codons and splice sites are not altered by these deletions^40^. These examples illustrate that the previous quantification of mutational constraint from short-read sequencing may be miscalibrated for some genes where simple repeat and SD sequences leave blind spots to short reads that are now captured by long reads, which is an area of future consideration for mutational intolerance surveys. Among all 1,308 SVs, we find 7 MEIs (6 *Alu* elements and 1 SVA). Extending this analysis with an orthogonal approach, we identify 11 additional MEIs (8 *Alu* elements, 2 SVAs, and 1 L1) that occur in protein-coding exons (**Methods**). The 18 MEIs disrupted one gene each affecting 56 transcripts (**Supplementary Table 43**). The insertions introduced a premature stop codon in most transcripts (98.2%). More than half (n=28) of these are predicted to evade nonsense-mediated decay (**Methods**). Most MEIs (72%) represent singleton (n=11) or rare events (allele frequency (AF) 0.01-0.05; n=2), with five being common insertions (AF>0.05)^41^. Furthermore, we identify 216 MEIs (188 *Alu* elements, 15 SVAs, and 13 L1s) that integrate into the untranslated regions (UTRs) of 216 protein-coding genes impacting 811 transcripts. Most of these MEIs occur in the 3’ UTR (71%) and are mainly (73%) singleton (n=101) or rare events (n=60; median: 0.016).

#### Transcriptional effect of SVs

To determine the effects of SVs on isoforms, we generated long-read Iso-Seq data for 12 of the 65 assembly samples (EBV-transformed B-lymphocyte cell lines; **Methods**). Alignment of phased long-read isoforms with haplotype-resolved personal assemblies results in improved alignment quality (**Fig. 2d**), isoform discovery (**Extended Data Fig. 3a**), and detection of imprinted genes (**Supplementary text Imprinted Loci**). Through intersection of the Iso-Seq data with 176,531 GRCh38 SVs (**Methods**), we identify a total of 136 structurally variable protein-coding gene sequences with sample-matched Iso-Seq expression data, each involving exon-disrupting SVs (**Supplementary text Expression Impacted by SVs**, **Supplementary Table 44**). Of these 136 genes, 58% (n=79) contained a common SV (AF>0.05) (**Extended Data Fig. 3b**). One example, *ZNF718*, harbors a common (AF=0.55) 6,142 bp polymorphic deletion that removes exons 2 and 3 from the canonical transcript as well as the 3’ part of an exon annotated as an alternate first exon (**Extended Data Fig. 3b**). The deletion is associated with nine unique isoforms (2 known, 7 previously unreported), five of which are shown in **Figure 2d**. In addition to three known isoforms, we also identify four previously unreported isoforms across the 14 wild-type haplotypes (**Methods**). In contrast to other protein-coding genes with a single SV (**Extended Data Fig. 3c**), we find a greater transcript diversity among the variant haplotypes of *ZNF718* compared to wild-type haplotypes.

Broadening our approach to identify SVs affecting nearby gene expression (RNA-seq), we find 122 unique SVs proximal (<50 kbp) to 98 differentially expressed genes across the 12 samples, representing an enrichment compared with randomly permuted SVs (**Extended Data Fig. 3d**) (empirical p=0.001; **Supplementary Methods; Supplementary Fig. 25, Supplementary Table 45**). Genome-wide, SVs were depleted across protein-coding genes and regulatory regions in the genome, as expected^42^ (**Extended Data Fig. 3e-f; Supplementary Fig. 26**). By intersecting these 122 SVs with Hi-C data from these 12 samples, we find that 29 of the SVs (associated with 24 genes) are potentially associated with contact density changes in chromatin conformation regions (**Extended Data Fig. 3g, Methods, Supplementary Table 46**). Finally, we identified 3,818 SVs in high linkage disequilibrium with single-nucleotide polymorphism (SNP) loci uncovered in genome-wide association studies (GWAS)^43^ (**Supplementary Table 47, Methods**), emphasizing biological relevance to human traits and diseases (**Extended Data Fig. 3h**). Collectively, these results underscore the potential effects of SVs on isoform-level transcriptomes, gene regulation, chromatin architecture, and disease associations.

**Extended Data Figure 3.**
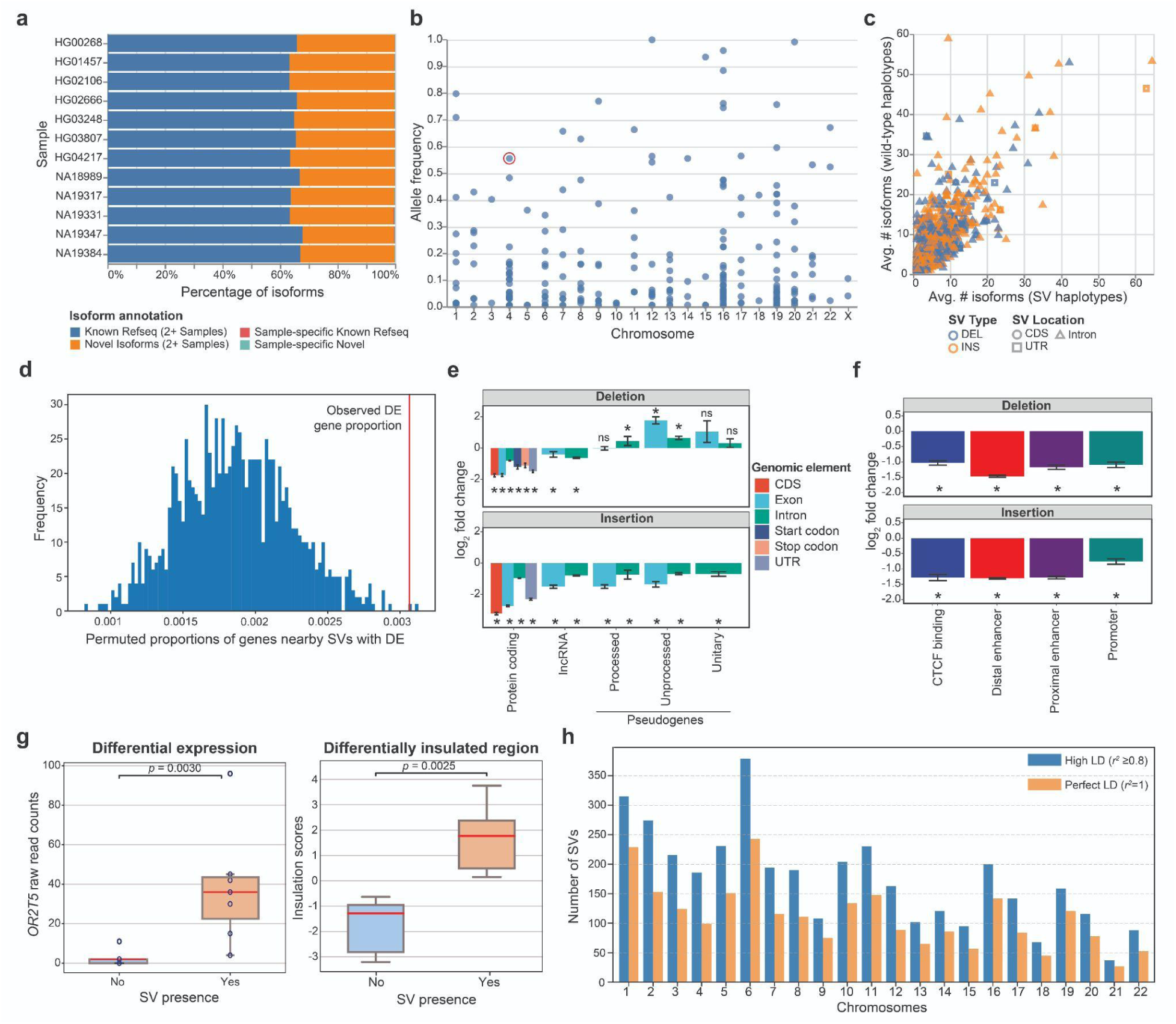
Effects of SVs on gene expression, chromosome conformation, and complex traits. **a)** The percentage of Iso-Seq isoforms identified for each sample classified as novel (present in at least two samples; orange), previously identified in RefSeq (present in at least two samples; blue), sample-specific novel (teal), or sample-specific previously identified isoforms (red). **b)** Manhattan plot of the allele frequencies for 256 SVs disrupting protein-coding exons of 136 genes with expression present in Iso-Seq. Circled in red is the 6,142 bp polymorphic deletion in *ZNF718*. **c)** Comparison of the average unique isoforms in Iso-Seq phased to wild-type and variant haplotypes for 1,471 single SV-containing protein-coding genes. The color represents the type of SV (deletion: blue, insertion: orange) and the shape indicates where the SV occurs in relation to the canonical transcript (circle: coding sequence [CDS], square: UTR, triangle: intron)**. d)** Proportion of genes located within 50 kbp of SV regions that show differential expression (DE) (RNA-seq) among individuals who carry the SVs (red line), compared with the distribution of DE gene proportions nearby simulated SV regions (1,000 permutations). **e)** Enrichments and depletions of SVs within GENCODE v45 protein-coding, long noncoding RNA (lncRNA), and pseudogene elements, subdivided into various biotypes. *empirical p<0.05 with Benjamini-Hochberg correction. ns, nonsignificant. Error bars indicate s.d. **f)** Enrichments and depletions of SVs within classes of ENCODE candidate cis-regulatory elements (cCREs). *empirical p<0.05 with Benjamini-Hochberg correction. ns, nonsignificant. Error bars indicate s.d. **g)** A differentially insulated region (DIR) in individuals with chr1-248444872-INS-63 SV, located nearby the DE gene *OR2T5*, suggests an SV-mediated novel chromatin domain could lead to increased gene expression. Box plots indicate first and third quartile, with whiskers extending to 1.5 times the interquartile range. **h)** Number of SVs per chromosome that are in high (r^2^>0.8) or perfect (r^2^=1) linkage disequilibrium (LD) with GWAS SNPs significantly associated with diseases and human traits.

### Genotyping and integrated reference panel

#### Genome-wide genotyping with PanGenie

Pangenome reference structures have enabled genome inference, a process where all variation encoded within a pangenome is genotyped in a new sample from short-read whole-genome sequencing (WGS) data. This has increased the number of SVs detectable from short-read WGS to about 18,000 per sample in previous work by the HPRC but was thus far mostly limited to common variants due to the relatively small discovery sets used to build the HPRC draft reference^1^. To expand this to variants with lower allele frequency (AF) and gaps in HPRC assemblies^2^, we constructed a pangenome graph containing all 65 samples assembled here as well as 42 HPRC samples^1^ with Minigraph-Cactus^44^ (MC) and detected variants by identifying graph bubbles relative to T2T-CHM13^44^ (**Methods**). We used PanGenie^3^ to genotype bubbles across all 3,202 samples from the 1kGP cohort based on Illumina data^45^ and used our previously developed decomposition approach to decompose the 30,490,169 bubbles into 28,343,728 SNPs, 10,421,787 indels, and 547,663 SV variant alleles^1^ (**Supplementary Fig. 27, Methods**). Leave-one-out (LOO) experiments confirmed high genotype concordance of up to ∼94% for biallelic SVs (**Supplementary Figs. 28-30**) and filtering the resulting genotypes from the 3,202 samples based on a previously developed regression model^1,6^ resulted in a set of 25,695,951 SNPs, 5,774,201 indels, and 478,587 SVs of reliably genotypable variants (**Supplementary Methods**, **Supplementary Table 48, Supplementary Figs. 31,32**). We note that this SV set is larger than our main PAV callset (188,500 SVs) because it includes the HPRC samples and at the same time retains all SV alleles at multi-allelic sites. We confirm the concordance between the two sets by observing that 392,883 SVs (82%) from the filtered PanGenie set have a matched call in the main PAV callset (**Supplementary Methods, Supplementary Fig. 33**).

We compared our genotyped set to other SV sets for the 1kGP samples, including the HPRC PanGenie genotypes we produced previously^1^ as well as the 1kGP short-read high-coverage SV callset (1kGP-HC)^45^ (**Supplementary Figs. 34,35**). On average, we found 26,115 SVs per sample, while this number was 18,462 for the HPRC genotypes and 9,596 for the 1kGP-HC SV calls. We specifically observed increases for rare variants (AF < 1; **Fig. 3a**). While the average number of rare SVs per sample was 87 for non-AFR samples in the HPRC set and 169 in the 1kGP-HC set, we can now access on average 362 rare alleles. For African samples, we detected 1,490 rare SVs per sample, while there were 382 previously for the HPRC and 477 for the 1kGP-HC set.

**Figure 3.**
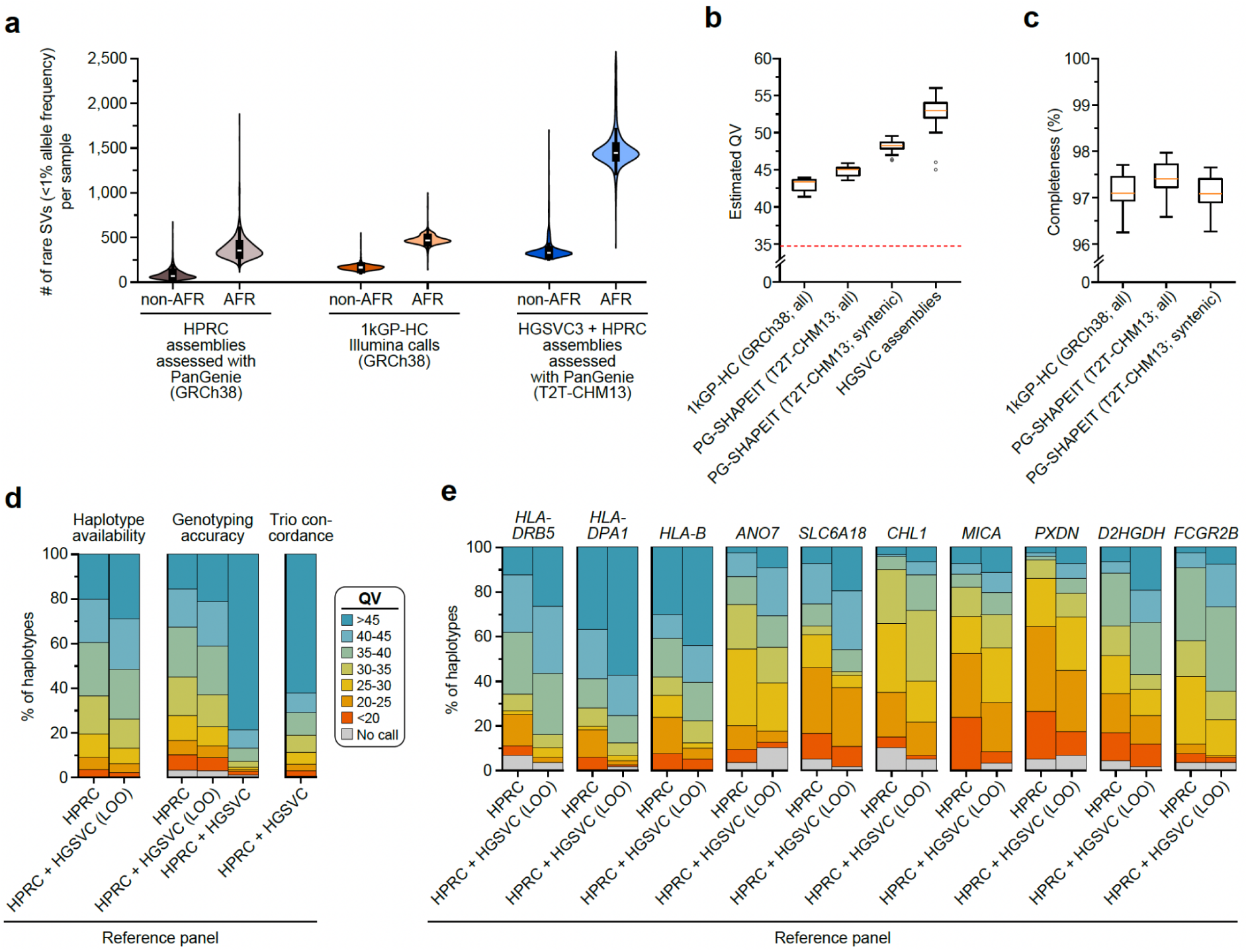
Genotyping from short-read sequencing data. **a)** Number of rare SVs, defined as those with an allele frequency of <1%, in each callset. We compared the HPRC genotyped callset (gray), the Illumina-based 1kGP-HC SV callset (orange), the combined HPRC and HGSVC genotyped callset (blue) for both non-African (non-AFR) and African (AFR) samples (*n*=3,202). The boxes inside the violins represent the first and third quartiles of the data, white dots represent the medians, and black lines mark minima and maxima of the data. **b)** Estimated QV for a subset of 60 haplotypes **(Supplementary Methods)** from the 1kGP-HC phased set (GRCh38-based), HGSVC phased genotypes (T2T-CHM13-based), and all HGSVC genome assemblies. To allow comparison between the GRCh38- and T2T-CHM13-based sets, we additionally restricted our QV analysis to “syntenic” regions of T2T-CHM13, i.e., excluding regions unique to T2T-CHM13. The red dotted line corresponds to the baseline QV that we estimated by randomizing sample labels (i.e., using PanGenie-based consensus haplotypes and reads from different samples). The median is marked in yellow and the lower and upper limits of each box represent lower and upper quartiles (Q1 and Q3). Lower and upper whiskers are defined as Q1 − 1.5(Q3–Q1) and Q3 + 1.5(Q3–Q1), and dots mark the outliers. **c)** Completeness statistics for haplotypes produced from the 1kGP-HC phased set (GRCh38-based) and the HGSVC phased genotypes (T2T-CHM13–based). To allow for comparison between the GRCh38- and T2T-CHM13-based callsets, we additionally restricted our analysis to “syntenic” regions of T2T-CHM13, i.e., excluding regions unique to T2T-CHM13. For both phased sets, completeness was computed on a subset of 30 samples. **d)** Haplotype availability, Locityper genotyping accuracy, and trio concordance across 347 polymorphic loci. Availability and accuracy are calculated for 61 HGSVC samples, while trio concordance is calculated for 602 trios. Results are grouped by the reference panel [HPRC-only, HPRC + HGSVC leave-one-out (LOO), and HPRC + HGSVC]. **e)** Locityper genotyping accuracy for 10 target loci with the highest average QV improvement.

#### Personal genome reconstruction from an integrated reference panel

Next, we asked to what extent our improved genotyping abilities allow us to reconstruct the full haplotypic sequences of genomes sequenced with short reads. In order to capture additional rare variants not represented in our discovery set and hence not genotyped, we combined our filtered PanGenie genotypes (see Section “Genome-wide genotyping with PanGenie”) with rare SNP and indel calls obtained from Illumina reads for all 3,202 1kGP samples (**Supplementary Methods**). We phased this combined set using SHAPEIT5^46^ to create a reference panel for the 1kGP set (**Supplementary Fig. 27**, Step 3). We evaluated the phasing results by comparing to our phased assemblies (SNPs, indels, SVs) as well as to a phased Illumina callset for the 1kGP samples (SNPs, indels) (**Supplementary Methods**) and observed low switch error rates, especially for samples that are part of trios (0.26%–0.18% for child samples, 0.69%–0.66% for parents, 1.21%–1.23% for unrelated samples; **Supplementary Methods, Supplementary Fig. 36,37**).

We produced consensus haplotype sequences for all 3,202 samples (6,404 haplotypes) by implanting the phased variants into T2T-CHM13 (only Chromosomes 1-22 and Chromosome X) and compared QV estimates and *k*-mer completeness values to consensus haplotypes produced from the GRCh38-based phased 1kGP-HC panel containing short-read-based SNP, indel and SV calls^45^. While the median QV of the long-read assemblies was 53, we observed a median QV of 45 for the consensus haplotypes computed from our short-read-based phased genotypes (**Fig. 3b, Supplementary Fig. 38**). To enable a fair comparison with the GRCh38-based 1kGP-HC consensus haplotypes, we additionally computed our QV estimates restricted to regions shared between T2T-CHM13 and GRCh38 (“CHM13-syntenic”). For these regions, we observed a median QV of 48, while the QV for the 1kGP-HC set was lower (median 43; **Fig. 3b**, **Supplementary Fig. 38**). Additionally, we computed the *k*-mer completeness for the consensus haplotypes (**Fig. 3c, Supplementary Fig. 38**). We observed higher *k*-mer completeness values (median 97.4%) for the consensus haplotypes produced from our phased genotypes than for the ones produced from the 1kGP-HC phased set (median 97.1%), which is consistent with the use of the more complete reference genome T2T-CHM13. When restricting to CHM13-syntenic regions, completeness values between both sets are similar. In summary, our analysis shows that the genome inference process from short reads, consisting of PanGenie genotyping followed by phasing, allows us to reconstruct the haplotype sequences to a remarkable quality that was previously only achievable from genome assemblies. The 6,404 consensus genome sequences are available as a community resource in an AGC-compressed archive^47^.

#### Targeted genotyping of complex polymorphic loci

While PanGenie performed well in this genome-wide setting, its use of *k*-mer information could make it difficult to genotype complex, repeat-rich loci with few unique *k*-mers. We therefore complemented the genome-wide genotyping by using the targeted method Locityper^48^ to genotype the 1kGP cohort across 347 polymorphic targets covering 18.2 Mbp and 494 protein-coding genes (**Methods**), including 268 challenging medically relevant genes^49^. As a reference panel, we used the haplotypes from the MC graph described above. For each short-read sample, Locityper reports a genotype for each locus by picking two of these haplotypes according to a maximum-likelihood approach.

Locityper’s performance is constrained by the haplotypes available in the reference set. Therefore, we first evaluated haplotype availability by comparing sequences of the unrelated assembled haplotypes. Across all target loci, 51.5% of our assembled haplotypes were very similar (QV ≥ 40) to some other haplotype from the full reference panel, compared to only 39.6% of haplotypes when restricting to an HPRC-only reference panel^1^ (**Fig. 3d**). Moreover, the number of almost-complete loci (average QV availability ≥ 40) increased almost twofold, from 95 loci for the HPRC-only panel to 167 loci for the full panel.

The increased haplotype availability translates into improved genotyping of polymorphic loci and we observe 80.0% haplotypes to be predicted with QV ≥ 30 using a LOO experiment compared to 74.6% haplotypes for the HPRC-only panel (**Methods**). When requiring QV ≥ 40, we still recover 42.6% of all haplotypes at this quality, compared to 33.7% for the HPRC-only panel (**Methods**). These global improvements are mirrored by improvements of individual genes, with the top 10 genes in terms of average increase in QV shown in **Figure 3e**, including *HLA-DRB5*, *HLA-DPA1*, and *HLA-B* (**Extended Data Fig. 5**). To produce a resource across these challenging loci for the community, we ran Locityper on all 3,202 samples from the 1kGP cohort. The reliability of the genotypes in the LOO experiments above is further corroborated by high Mendelian consistency observed across the 602 trios, with an average concordance QV of 42.6. Lastly, we asked what performance could potentially be achieved with Locityper for growing reference panels, where virtually all relevant haplotypes are available. To answer this, we compare the genotypes obtained for our samples when genotyping using the full reference panel to the respective assembly haplotypes. Here, Locityper predicts highly accurate haplotypes, with average QV of 45.8 and only 1.6% of the haplotypes with QV < 20, suggesting that sequence resolution of more reference haplotypes will aid future re-genotyping of challenging medically relevant genes, with applications to disease cohorts.

### Major Histocompatibility Complex

Given the complexity of the 5 Mbp MHC region (**Fig. 4a**) and its relevance to disease^50–52^, we performed a more systematic analysis of the 130 complete or near-complete MHC haplotypes. Per haplotype we annotate 27-33 HLA genes (human leukocyte antigen) and 140-146 non-HLA genes/pseudogenes along with the associated repeat content (**Supplementary Table 49**). We found that 99.2% (357/360) of the HLA alleles agree with classical typing results^53^ (**Supplementary Tables 50,51**). We identify and submit a total of 826 previously incomplete HLA allele annotations to the IPD-IMGT/HLA reference database^54^ (**Supplementary Table 52**). This includes 112 sequences from the HLA-DRB locus, important for vaccine response and autoimmune disease^55,56^. We observe a previously unknown copy number variant, a deletion of *HLA-DPA2* on one haplotype (HG03807.1), as well as low-frequency gene-level SVs, such as a deletion of *MICA* on one haplotype^57^. We identify 170 SVs not present in previously available reference haplotypes^58,59^ (**Supplementary Table 53**); the majority of these (64%) are enriched in repeats (≥70%); 30 of the remaining ones are >1 kbp.

**Figure 4.**
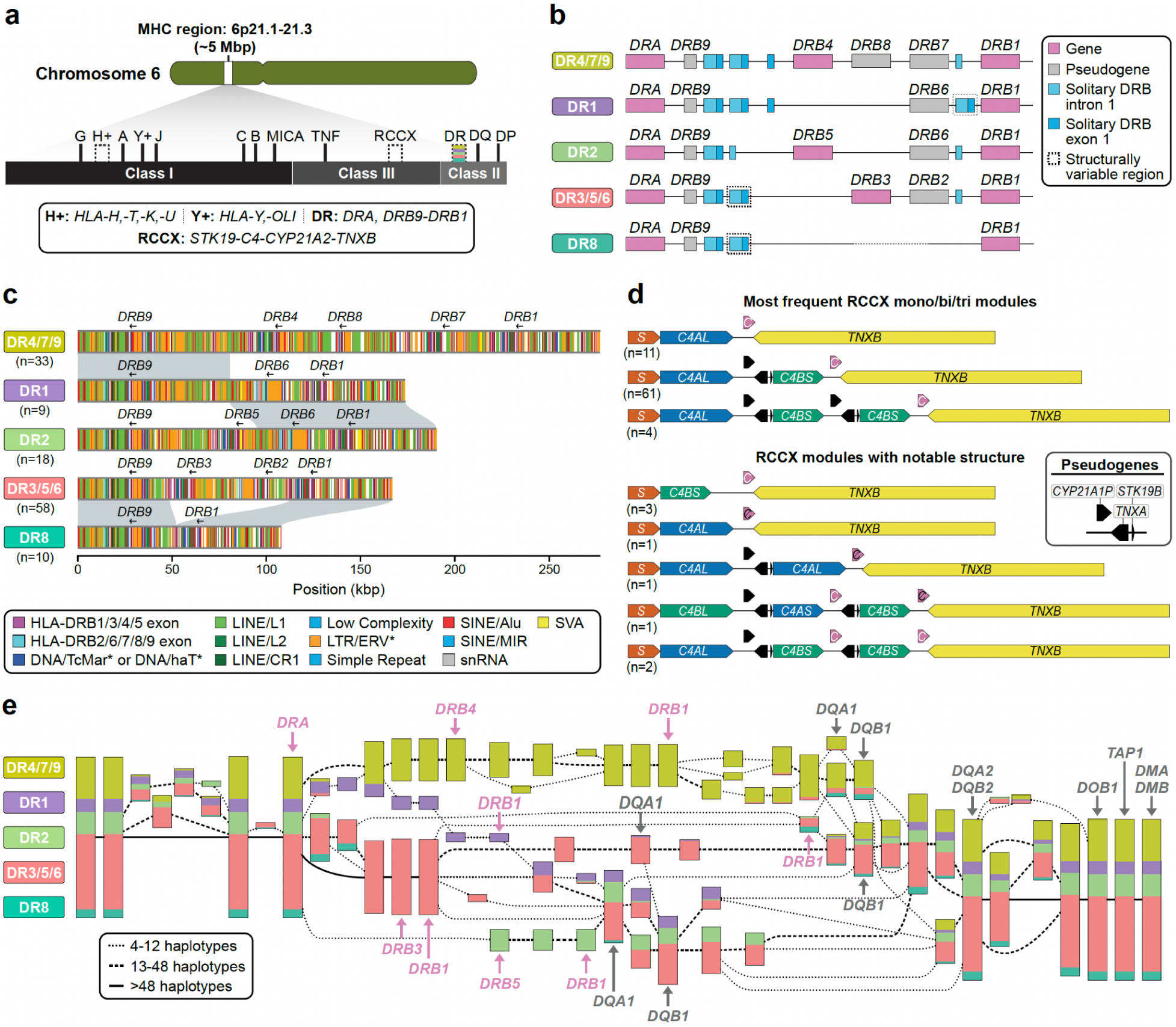
Structurally variable regions of the MHC locus. **a)** Overview of the organization of the MHC locus into class I, class II, and class III regions and the genes contained therein. Structurally variable regions are indicated by dashed lines. Colored stripes show the approximate location of the regions analyzed in panels b-d. **b)** Gene content and locations of solitary *HLA-DRB* exon 1 and intron 1 sequences in the HLA-DR region of the MHC locus by DR group, an established system for classifying haplotypes in the HLA-DR region according to their gene/pseudogene structure and their *HLA-DRB1* allele. Also shown is the number of analyzed MHC haplotypes per DR group. **c)** High-resolution repeat maps and locations of gene/pseudogene exons for different DR group haplotypes in the HLA-DR region, highlighting sequence homology between the DR1 and DR4/7/9 and DR2, and between the DR8 and DR3/5/6, haplotype groups, respectively. **d)** Visualization of common and notable RCCX haplotype structures observed in the HGSVC MHC haplotypes, showing variation in gene and pseudogene content as well as the modular structure of RCCX (S, *STK19*; black C, nonfunctional *CYP21A2*; white C, functional *CYP21A2*; *C4L*/*S*, long [(HERV-K insertion)/short(no HERV-K insertion)]. **e)** Visualization of a PGR-TK analysis of 55 MHC samples and T2T-CHM13 for 111 haplotypes in total. Colors indicate the relative proportion of distinct DR group haplotypes flowing through the visualized elements.

**Extended Data Figure 4.**
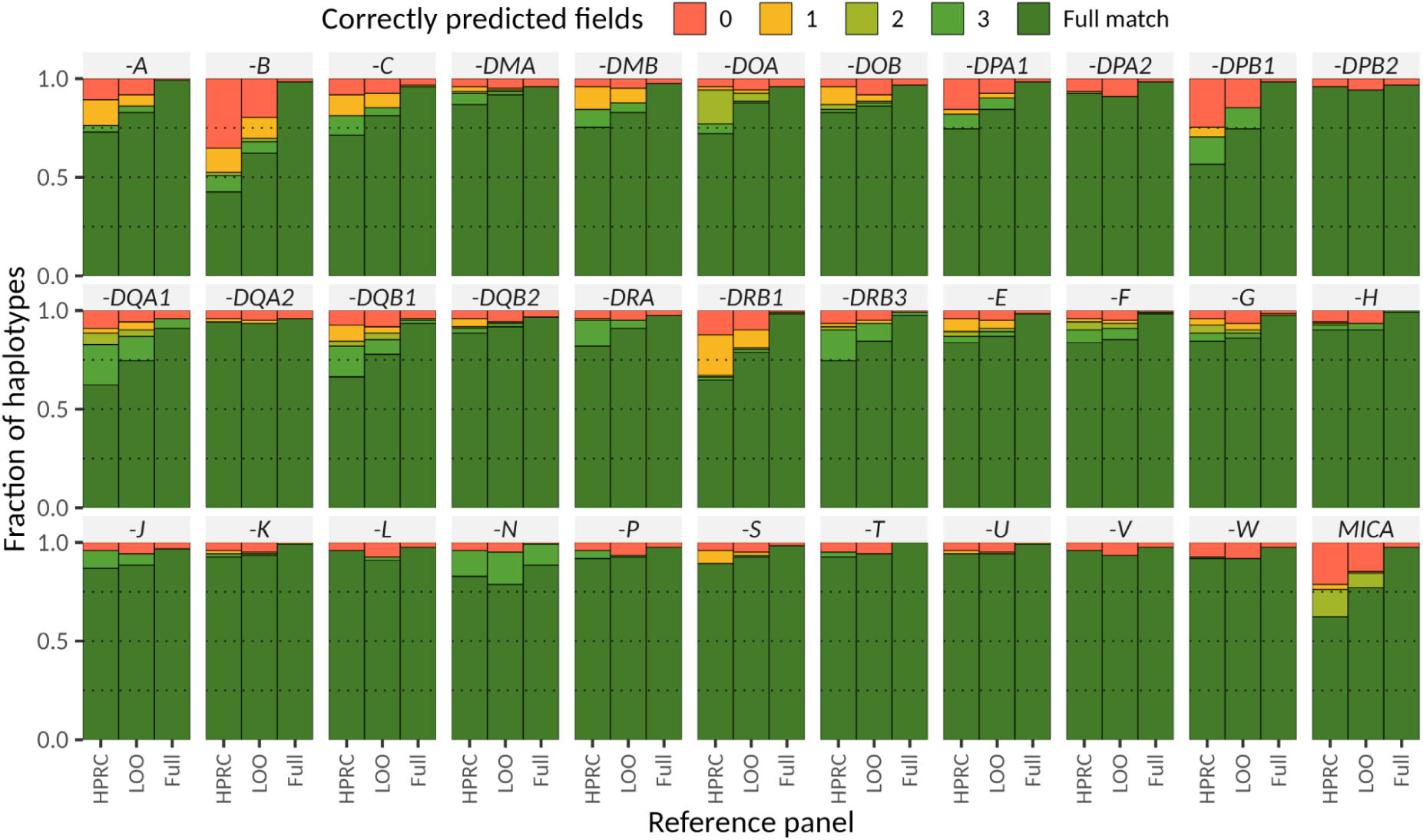
Locityper genotyping accuracy across 33 genes/pseudogenes, located at the MHC locus. Genotyping was performed for 61 Illumina short-read HGSVC datasets using three reference panels: HPRC (90 haplotypes), leave-one-out HPRC + HGSVC (LOO, 214 haplotypes), and HPRC + HGSVC (full, 216 haplotypes). Accuracy is evaluated as the number of correctly identified allele fields in the corresponding gene nomenclature.

Overall, MHC class II haplotypes are compatible with the established DR group system (**Fig. 4b**, **Supplementary Table 54**) and comprise representatives of DR5, DR8 and DR9, which were not previously analyzed in detail^58,59^. Repeat element analyses (**Supplementary Figs. 39-41**; **Methods**) suggest that an intrachromosomal deletion mediated by 150 bp of sequence homology between *HLA-DRB1* and *HLA-DRB3* on the DR3/5/6 haplotype gave rise to DR8 potentially as a result of homology-mediated double-strand breakage (**Fig. 4c**). Similarly, we find that DR1 is most likely derived from a recombination event occurring between DR2 and DR4/7/9 (**Fig. 4c**) with a LINE/L1 repeat defining the breakpoints (**Supplementary Figs. 39,42**). Finally, we also create a more comprehensive catalog of solitary HLA-DRB exon sequences^60,61^. This includes refining copy number estimates (e.g., observing a copy number of 3 instead of 1 for solitary *HLA-DRB* exon 1 sequences in the *HLA-DRB9* region of DR1) as well as identifying a previously uncharacterized solitary exon location 10 kbp 3’ of *HLA-DRB1* present on some haplotypes (**Fig. 4b, Supplementary Methods**).

Similarly, we characterize the RCCX multi-allelic cluster where tandem duplications or modules of variable copy number are each composed of four genes: serine/threonine kinase 19 (*RP1*/*STK19*), complement component 4 (*C4*), steroid 21-hydroxylase (*CYP21*), and tenascin-X (*TNX*)^62–64^. Each module typically encodes one functional variant of *C4* (*C4AS, C4AL, C4BS,* or *C4BL*) and three additional functional genes (*STK19*, *CYP21A2*, and *TNXB*) or pseudogenes (*STK19B*, *CYP21A1P*, and *TNXA*) (**Fig. 4d**, **Supplementary Fig. 43, Supplementary Table 55**); however, phasing and variant classification has been challenging due to extensive sequence homology^64,65^. Tandem duplications (aka RCCX bi-modules) are the most abundant among our assembly haplotypes (74.6% or n=97) with mono-modules and tri-modules comparable in frequency (13.1% (*n*=17) and 12.3% (*n*=16), respectively (**Supplementary Fig. 43**). The complete haplotype sequence also facilitates the detection of interlocus gene conversion events critical for RCCX evolution^66–68^. For example, we identify two haplotypes having a tri-modular RCCX with two functional *CYP21A2* copies, likely the result of interlocus gene conversion; one mono-modular and one bi-modular haplotype with no functional *CYP21A2* genes; and one tri-modular haplotype with a unique configuration where *C4B* precedes *C4A* and carries two copies of *CYP21A2*, one of which was nonfunctional (**Fig. 4d**). We suggest that the latter haplotype was generated by the introduction of one nonsense mutation and two gene conversion events, converting *CYP21A1P* into *CYP21A2* and *C4A* into a *C4B* that now unusually encodes the Rodgers (Rg) blood group epitope. We also identify seven novel *C4* amino acid variants, four residing within the beta-chain (283R->C, 530S->L, 566A->G, 614K->Q) and three within the gamma-chain (1477K->N, 1722R->C, 1724R->W) (**Supplementary Figs. 44,45**).

Finally, we tested whether the established class II DR group nomenclature could be recapitulated using unbiased, sequence-based analysis. To this end, we applied a pangenomic multiscale analysis method, PGR-TK^69^ (**Fig. 4e**), to a subset of our genomes (*n*=55) as well as T2T-CHM13^7^. We identify 63 conserved blocks in 111 haplotypes (55 diploid genomes corresponding to 110 haplotypes plus the haploid T2T-CHM13) greater than 6 kbp. Multiscale hierarchical clustering of the haplotypes perfectly reconstitutes the traditional DR group system in the region around *HLA-DRB1* (**Fig. 4e**). However, we also observe additional diversified subgroups that could serve as the basis for a more fine-grained future classification of HLA-DR haplotypes or be used in the context of GWAS, especially when coupled with the improved ability for targeted genotyping (**Extended Data Fig. 4**).

### Complex structural polymorphisms

Long-read assembled genomes significantly enhance the detection and characterization of complex structural variants (CSVs) defined here as SVs with more than two breakpoints. Because CSV breakpoints are often located in repetitive sequences, including SDs and MEIs^70–73^, we recently updated PAV^6^ to identify CSVs embedded in large complex repeats like SDs (**Methods**). Using this method, we find on average 55 complex SVs per sample (range 37 to 85) with median SV sizes ranging from 47 kbp to 222 kbp (**Supplementary Table 56**, **Data Availability**). Across all samples, we identify 1,909 complex SVs with 200 distinct complex reference signatures composed of deletions (*n*=1,498), duplications (*n*=440), inversions (*n*=196), inverted duplications (*n*=207), and higher-copy elements (*n*=142). Because many CSVs contain distal sequences, many leave alignments that mimic simple deletions (*n*=1,107; 60%) and duplications (*n*=152; 8.2%), which can mislead downstream SV calling tools. To address this, PAV links these distal and unaligned segments with the CSV to resolve its structure. As an example, we highlight two CSVs involving *NOTCH2NL* and *NBPF*, genes implicated in expansion of the human brain^6^, as well as a core duplicon associated with genomic instability^74^. While the full structures could not be resolved by previous optical mapping or sequencing experiments, we can now distinguish four distinct haplotype structures, including a reference haplotype (AF 0.15), a simple inversion around *NBPF8* (AF 0.02), a 986 kbp CSV with a distal template switch replacing *NBPF8* with *NBPF9* (AF 0.51), and an 851 kbp CSV (DEL-INV-DEL) inverting *NBPF8* and deleting *NOTCH2NLR* and *NBPF26* (AF 0.33) (**Fig. 5a**). Closer examination of the DEL-INV-DEL CSV suggests there may be two similar CSV events with sizes 986 kbp (AF 0.18) and 848 kbp (AF 0.15). By combining contiguous assemblies with improved methods, we can now detect complex structures that other methods cannot. For example, no other SV callers we ran detected the complex *NBPF8* structures or their associated deletions.

**Figure 5.**
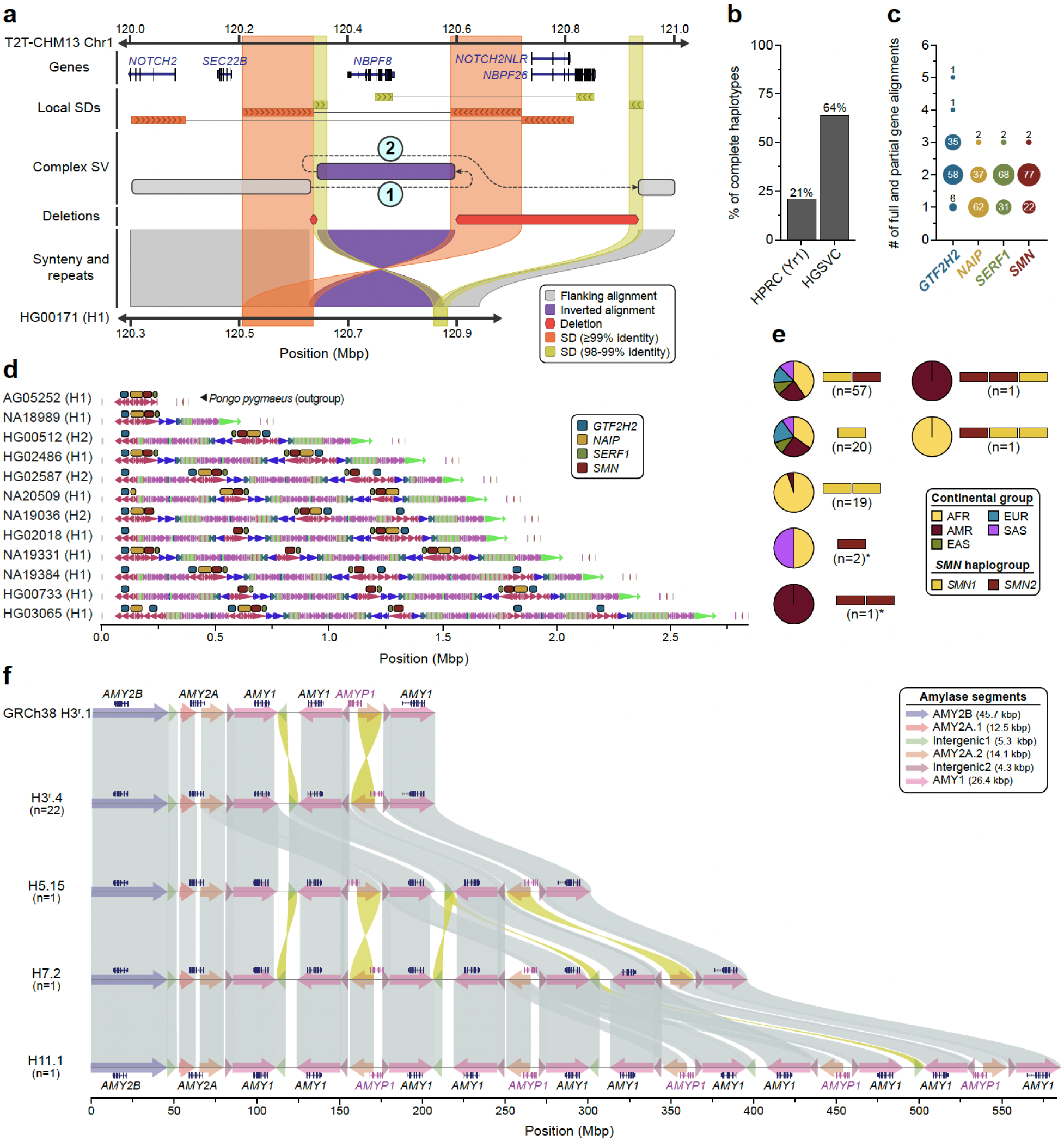
Complex SVs in human populations. **a)** An SD-mediated CSV inverts *NBPF8* and deletes two genes. Inverted SD pairs (orange and yellow bands) each mediate a template switch (dashed lines “1” and “2”). The resulting CSV inverts *NBPF8* and deletes *NOTCH2NLR* and *NBPF26*. The single recombined copy of each SD is aligned to both reference copies, obscuring the structure of the complex event by eliminating one deletion and changing the size of the inversion and the larger deletion. PAV recognizes these artifacts and refines alignments to obtain a more accurate representation of complex structures. The complex allele shown is HG00171 haplotype1-0000011. **b**) Fraction of all assemblies having complete and accurate sequence over the SMN region, stratified by study (HGSVC, HPRC-yr1). **c**) Copy number (full and partial gene alignments) of each multi-copy gene (*SMN1/2*-red, *SERF1A/B* - green, *NAIP* - gold, and *GTF2H2/C* - blue) across all human haplotypes (n=101). **d**) Visualization of DupMasker duplicons defined in 11 diverse human haplotypes spanning the SMN region. Panel depicts data from this study, the HPRC (HG02486), and one *Pongo pygmaeus* haplotype (top) used as an outgroup. **e**) Summary of *SMN1* (yellow) and *SMN2* (red) gene copies genotyped across human haplotypes (n=101). Yellow and red bars show a unique copy number of *SMN1* and *SMN2* while pie charts show proportions of continental groups carrying a given haplotype. Haplotypes that carry only the *SMN2* gene copy are highlighted by the asterisks. **f**) The amylase locus of the human genome is depicted. The H3r.4 haplotype represents the most common haplotype, H5.15 and H7.2 are haplotypes previously unresolved at the base-pair level, and H11.1 is a novel, previously undetected haplotype. Amylase gene annotations are displayed above each haplotype structure. The structure of each amylase haplotype, composed of amylase segments, is indicated by colored arrows. Sequence similarity between haplotypes ranges from 99% to 100%. The alignments highlight differences between the amylase haplotypes.

As a second example, we focus on a biomedically relevant and structurally complex region containing *SMN1* and *SMN2* gene copies. Deletion or disruption of *SMN1* is associated with spinal muscular atrophy (SMA) and its paralogue, *SMN2*, is the target of one of the most successful ASO-mediated gene therapies^75,76^. The genes are embedded in a very large SD region (∼1.5 Mbp) that has been almost impossible to fully sequence resolve despite the advances of the last two decades^1,2,6,77^ (**Supplementary Fig. 46**), including in the recent HPRC pangenome release^1,2^. We successfully assembled, validated, and characterized two thirds of all possible haplotypes (*n*=101) fully resolving the structure and copy number (full or partial gene alignments) of *SMN1/2*, *SERF1A/B*, *NAIP*, and *GTF2H2/C*. We find that about half (*n*=48) of all haplotypes carry exactly two copies of *SMN1/2*, *SERF1A/B*, and *GTF2H2/C* while *NAIP* is present mostly in a single copy. We highlight 11 human haplotypes that show increasing complexity in duplicon structure and palindromic structures (**Fig. 5b-d**). We specifically distinguish functional *SMN1* and *SMN2* copies based on our assemblies (**Supplementary Fig. 47**) and compare them with short-read based genotyping methods Parascopy^78^ and SMNCopyNumberCaller^79^. For samples where both haplotypes are fully assembled (*n*=31), predicted *SMN1/2* copy numbers match perfectly among the three methods (**Supplementary Fig. 48**). Our analysis shows that 98 haplotypes carry the ancestral *SMN1* copy but three do not and are potentially disease-risk loci that may have arisen as a result of interlocus gene conversion (**Fig. 5e, Supplementary Fig. 49**).

Finally, we analyzed the structure of the amylase locus spanning 212.5 kbp on Chromosome 1 (GRCh38; chr1:103,554,220–103,766,732) and containing the *AMY2B*, *AMY2A*, *AMY1A*, *AMY1B*, and *AMY1C* genes^80^ (**Fig. 5f**). From 130 sequence-resolved genomes, we identify 35 distinct amylase haplotypes (**Supplementary Fig. 50, Supplementary Table 57**) supported by both Verkko and optical genome mapping *de novo* assemblies, representing the largest number of base-pair resolution amylase haplotypes described to date. The length of these amylase haplotypes ranges from 111 kbp (H1^a^.1 and H1^a^.2) to 582 kbp (H11.1) (**Fig. 5f**), including those that are structurally identical to the GRCh38 (H3^r^.1) and T2T-CHM13 (H7.3) assemblies. Among the identified haplotypes, four are common: H1^a^.1 (n=14), H3^r^.1 (n=13), H3^r^.2 (n=19), and H3^r^.4 (n=22) (constituting 57% of all genomes), while 23 are singletons. We identify nine haplotypes previously supported only by optical genome mapping data and fully sequence-resolve the largest novel haplotype to date (H11.1; 11 *AMY1* [8.8 kbp] copies)^81–83^ (**Fig. 5f**).

### Centromeres

Among the most mutable regions in the human genome are the centromeres, which are large chromosomal domains that are typically composed of tandemly repeating α-satellite DNA^84^. In humans, centromeric α-satellite DNA is organized into higher-order repeats (HORs), which are assembled into arrays that can span up to several megabases on each chromosome^85–87^. The repetitive nature of the centromeres means that they are prone to homologous recombination and other mutational processes that lead to duplications, deletions, and mutations of α-satellite HORs, resulting in expansions and contractions of the centromeric α-satellite HOR array. A previous study comparing two complete sets of human centromeres revealed that approximately 22% of centromeres vary by over 1.5-fold in length, and ∼30% of them vary in their structure^88^. Here, we expand our understanding of centromere diversity by assessing the genetic and epigenetic variation of the centromeres from 65 diverse human genomes.

We first assessed the contiguity and accuracy of each centromere by aligning Verkko and hifiasm assemblies to the T2T-CHM13 reference genome and identifying contigs that traversed the centromeres from the p- to the q-arm. Then, we assessed the accuracy of the centromeric assemblies using two computational tools, Flagger^1^ and NucFreq^15^, to identify those that lacked large structural errors. Ultimately, we identified 822 centromeres from the Verkko assemblies and 777 centromeres from the hifiasm assemblies that were completely and accurately assembled. We found that approximately 28.3% of all centromeres were correctly assembled by both assemblers, while 37.7% were correctly assembled by only Verkko, and 34.1% were correctly assembled by only hifiasm. We combined these two datasets into a nonredundant set of 1,246 completely and accurately assembled centromeres (∼52 centromeres per chromosome, on average, and ∼19.5 centromeres per genome, on average; **Extended Data Fig. 5a**, **Supplementary Tables 58,59**). We used this dataset, which is 621 (twofold) more centromeres than those previously analyzed^84,88–90^, to assess the genetic and epigenetic variation across 65 diverse genomes.

**Extended Data Figure 5.**
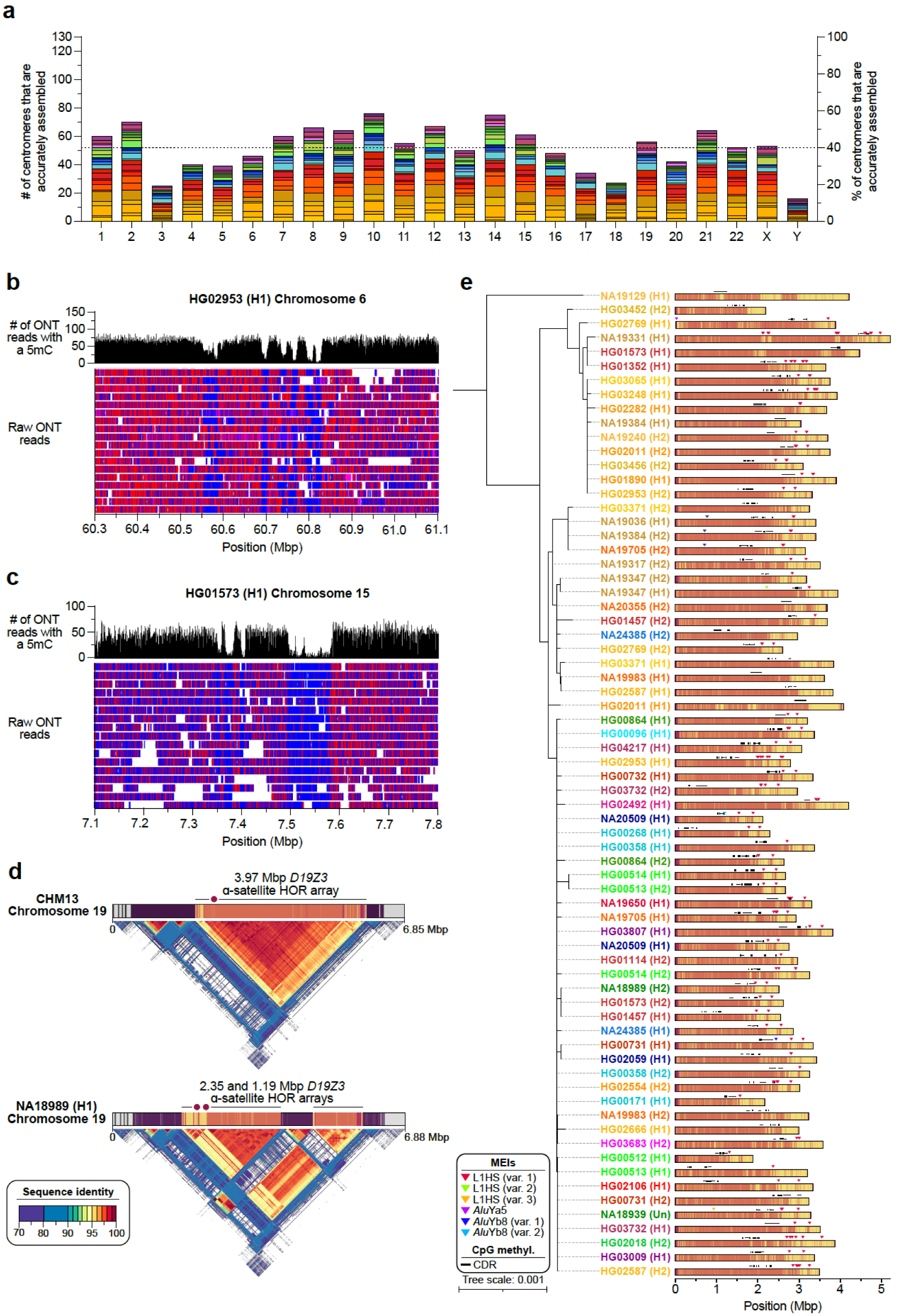
Assembly of 1,246 human centromeres across 65 diverse human genomes show genetic and epigenetic variation. **a)** Number of completely and accurately assembled centromeres across 65 diverse human genomes, colored by population group. Mean, dashed line. **b,c)** Examples of di-kinetochores, defined as two CDRs located >80 kbp apart from each other, on the **b)** HG02953 Chromosome 6 centromere and **c)** HG01573 Chromosome 15 centromere. Ultra-long ONT reads span both CDRs in each case, indicating that the CDRs occur on the same chromosome in the cell population. **d)** Differences in the ɑ-satellite HOR array organization and methylation patterns between the CHM13 and NA18989 (H1) chromosome 19 centromeres. The NA18989 (H1) chromosome 19 centromere has two CDRs, indicating the potential presence of a di-kinetochore. **e)** Mobile element insertions (MEIs) in the Chromosome 2 centromeric α-satellite HOR array. Most MEIs are consistent with duplications of the same element rather than distinct insertions, and all of them reside outside of the CDR.

We first measured the variation in the length of the centromeric α-satellite HOR array(s) on each chromosome (**Fig. 6a**, **Supplementary Table 59**). We found that, on average, centromeric α-satellite HOR arrays are 2.31 Mbp in length; however, there are clear outliers. For example, the α-satellite HOR arrays from Chromosomes 3, 4, 10, 13-16, 21, and Y are consistently smaller than average, with a mean length of 1.2, 1.1, 1.4, 1.6, 2.0, 1.5, 1.8, 1.3, and 0.7 Mbp, respectively. In contrast, the α-satellite HOR arrays on Chromosomes 1, 11, and 18 are much larger than average, with a mean length of 4.0, 3.5, and 3.9 Mbp, respectively. The greatest variation in length occurs on Chromosome 10, which has a 37.5-fold difference in length between the smallest and largest α-satellite HOR arrays; however, the greatest absolute difference in length occurs on Chromosome 18, wherein the smallest and largest α-satellite HOR arrays differ by 5.9 Mbp.

**Figure 6.**
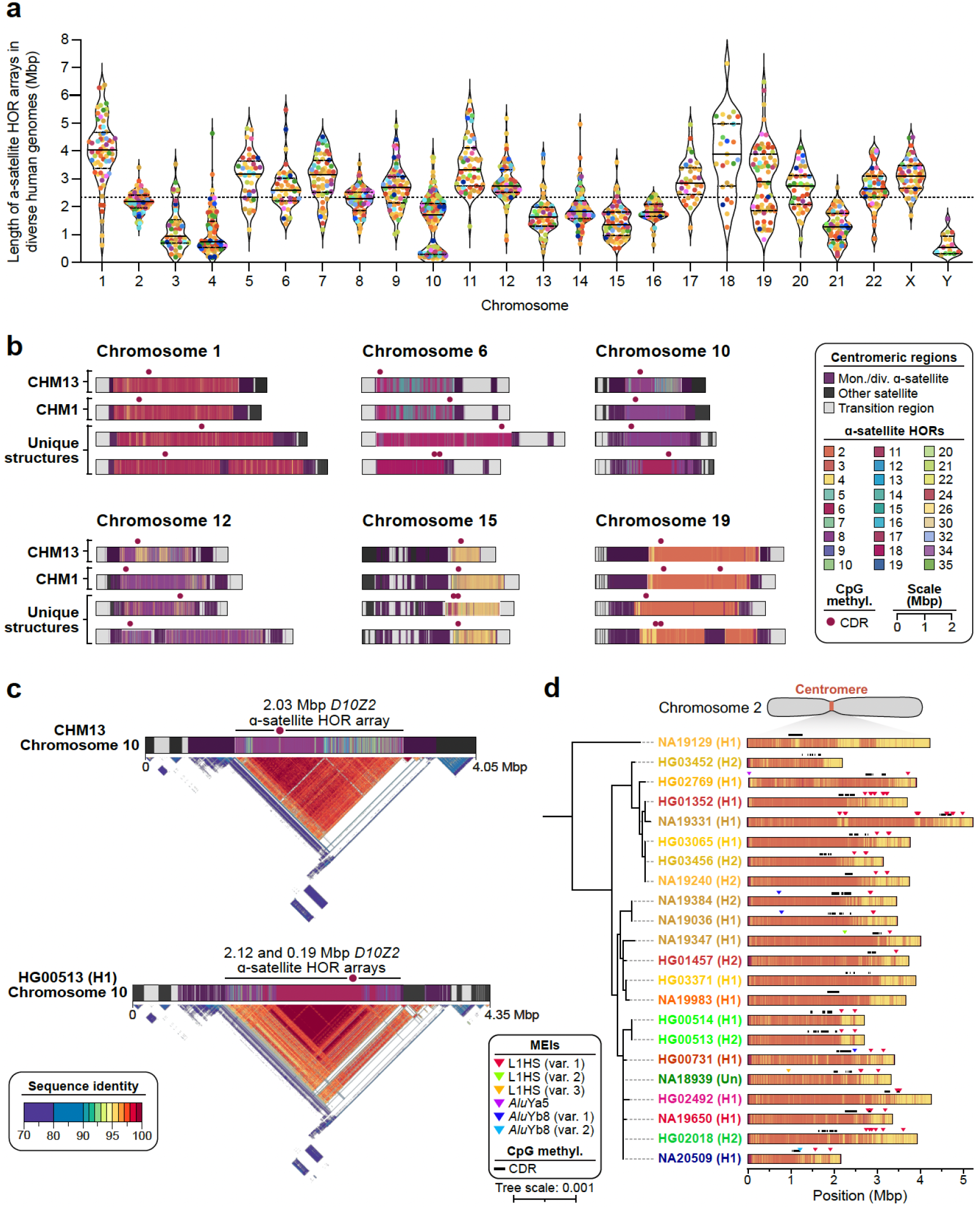
Variation in the sequence, structure, and methylation pattern among 1,246 human centromeres. **a)** Length of the ɑ-satellite higher-order repeat (HOR) array(s) for each complete and accurately assembled centromere from each genome. Each data point indicates an active ɑ-satellite HOR array and is colored by population. The median length of all α-satellite HOR arrays is shown as a dashed line. For each chromosome, the median (solid line) and first and third quartiles (dashed lines) are shown. **b)** Sequence, structure, and methylation map of centromeres from the CHM13, CHM1, and a subset of 65 diverse human genomes. The α-satellite HORs are colored by the number of α-satellite monomers within them, and the site of the putative kinetochore, known as the “centromere dip region” or “CDR”, is shown. **c)** Differences in the ɑ-satellite HOR array organization and methylation patterns between the CHM13 and HG00513 (H1) chromosome 10 centromeres. The CDRs are located on highly identical sequences in both centromeres, despite their differing locations. **d)** Mobile element insertions (MEIs) in the chromosome 2 centromeric α-satellite HOR array. Most MEIs are consistent with duplications of the same element rather than distinct insertions, and all of them reside outside of the CDR.

We compared the sequence and structure of all 1,246 centromeres and identified 4,153 new α-satellite HOR variants and novel array organizations among the active α-satellite HOR arrays (**Fig. 6b**). On Chromosome 1, for example, we found evidence of an insertion of monomeric α-satellite sequences into the *D1Z7* α-satellite HOR array, which had split the α-satellite HOR array into two distinct arrays (**Fig. 6b**). A similar bifurcation event also occurred on the Chromosome 12 and 19 centromeres, generating two α-satellite HOR arrays where there typically is only one (**Fig. 6b,c**). Additionally, we found novel α-satellite HOR array organizations for Chromosomes 6 and 10 that differ from the CHM1 and CHM13 arrays on those chromosomes^88^ (**Fig. 6b**). These array organizations, which are the most common in our dataset, are primarily composed of either 6-monomer α-satellite HORs (Chromosome 6) or 6- and 8-monomer α-satellite HORs (Chromosome 8). Finally, we found evidence of a relic α-satellite HOR variant on some centromeres, such as on Chromosome 15, where an ancestral 11-monomer α-satellite HOR variant was present in high abundance on some haplotypes but had since been substantially reduced in abundance in the CHM13 and CHM1 centromeres (**Fig. 6b**).

To determine how variation in centromeric sequence and structure affects their epigenetic landscape, we assessed the CpG methylation pattern along each centromere using native ONT data. We found that all centromeres contain at least one region of hypomethylation (termed the “centromere dip region” or CDR^84,91^) that is thought to mark the site of the kinetochore. However, in many cases, such as on Chromosomes 6, 15, and 19, there were at least two CDRs >80 kbp apart (**Fig. 6b, Extended Data Fig. 5b-d**). This suggests the presence of a “di-kinetochore”, which may form a dicentric chromosome on ∼7% of chromosomes, but additional analyses that assess the location of the centromeric histone H3 variant, CENP-A, will need to be performed to confirm these putative kinetochore sites. To determine whether the CDR associates with recently evolved and actively homogenizing α-satellite HORs, we generated sequence identity heatmaps^92^ of each centromere and found that, indeed, the CDR often resides within the most highly identical regions of the α-satellite HOR arrays (**Fig. 6c, Extended Data Fig. 5d**). Even when the α-satellite HOR array is split into two arrays, such as on Chromosome 19, the CDR associates with the array containing some of the most highly identical α-satellite HORs (**Extended Data Fig. 5d**). This suggests that the kinetochore may track with actively homogenizing α-satellite HOR sequences in response to a coevolution between centromeric DNA and proteins^93^. Additional analyses that test this centromere coevolution theory will need to be performed to confirm its validity.

In addition to variation in sequence and structure, we also noted the presence of MEIs in many of the α-satellite HOR arrays (**Methods**). We found that ∼30% of all active α-satellite HOR arrays contained at least one MEI (**Methods**). In total, we identified 89 unique polymorphic insertions with varying allele frequencies (**Supplementary Table 60**). L1 insertions were the most prevalent centromeric MEI, with 58% of all unique MEIs occurring from L1s, 41% occurring from *Alu* elements, and 1% from SVAs. One α-satellite HOR array that was highly enriched with MEIs was the *D2Z1* α-satellite HOR array on Chromosome 2 (**Fig. 6d**). This array had at least one L1HS and/or *Alu* insertion in 80% of haplotypes (**Extended Data Fig. 5e**). L1HS insertions/duplications were the most common, occurring on average three times per array, and three unique *Alu* insertions, two of which belonged to the *Alu*Yb8 subfamily and one *Alu*Ya5, were present in low allele frequency on the *D2Z1* α-satellite HOR array. Nearly all insertions, as well as their duplications, were located outside of the CDRs and typically towards the periphery. However, one *Alu*Yb8 insertion (in NA20509 haplotype 1) was located between two CDRs and appeared to ‘break’ a single CDR into two, while a pair of L1HSs were found on either side of a CDR in two haplotypes (NA19331 H1 and NA19650 H1), where they may act as boundaries to restricting the movement of the CDR and CENP-A chromatin, as suggested previously^94,95^.

## Discussion

Long-read sequencing and assembly have enabled both the full resolution of a human genome sequence^7^ and fundamentally deepened our understanding of human genetic diversity^1,6,18,19^. The development of a human pangenome reference^1,96^ should ideally be based on completely phased and assembled diverse genomes, with as few gaps and switch errors as technically possible. While there are plans to achieve this for hundreds of human genomes^97^, practically, few true T2T genomes have been generated and, thus, the pangenome is still a work-in-progress. Nevertheless, algorithms and technology have advanced significantly in the last two years, and we demonstrate that >99% of the human genome can be accurately phased and assembled by focusing on 65 diverse samples (130 human genomes). We characterize regions previously excluded or collapsed^1,2^ including centromeres, biomedically complex regions such as *SMN1/SMN2*, and thousands of more complex SV patterns. While acrocentric regions remain to be finished, we show that it is possible to fully resolve almost all variation in the genome providing new biological insights and enhancing the potential for future disease association.

As an example, we characterize one of the largest sets of complete MHC haplotypes assembled to date and show the value of this deeper pangenome representation for improved genotyping. Our detailed annotation confirms the high accuracy of our assemblies, defines precise breakpoints of rearrangement within and between haplotypes, provides a more complete characterization of genes and pseudogenes, and allows us to construct the first pangenome haplotype graph. Using these data, we apply Locityper^48^ to genotype from short-read data across 19 protein-coding genes and 14 pseudogenes from the MHC locus and compare predicted gene alleles against assembly-based gene annotation. Across all 33 loci, Locityper correctly predicts gene alleles in 81.0% cases when restricting to a limited HPRC-only reference panel (45 samples^1^). Adding our assemblies to the reference panel (n=107 samples or 214 phased genomes) increases accuracy to 86.3% (leave-one-out experiment) and 97.1% of the cases (full panel where all 214 phased genomes are leveraged). These differences highlight the significant gain in genotyping accuracy achieved by mapping short-read data to a more diverse human pangenome.

These observations for targeted genotyping are mirrored in a genome-wide setting, where this combined panel led to considerably improved genome inference. That is, given this combined resource, we are able to reconstruct a genome from short reads to a QV of up to 48, corresponding to an average base error of about 0.00158%. This process detects 26,115 SVs per sample on average from short-read sequence data and notably now recovers more rare SVs (AF<1%) than direct variant discovery from short reads. We want to emphasize, however, that these improvements are likely due to a combination of different factors, including improvements in assembly quality, the larger number of panel samples, improved versions of the Minigraph-Cactus^44^ and PanGenie^3^ softwares, and the switch to the more complete T2T-CHM13 reference genome. While it is difficult to precisely delineate the influence of all these individual factors, it appears likely that genotyping accuracy will improve even more as the HPRC, for example, increases the number of genomes to several hundreds and pushes those genomes to a T2T state^97^. This, in turn, will make disease-association studies from short reads considerably more powerful for complex variation.

The use of both ultra-long ONT and PacBio HiFi sequencing data generated from the same samples was critical to accessing previously unresolved regions of the genome. For example, we fully assembled 1,246 centromeres—42% of all possible centromeres in these samples—representing the deepest survey of completed human centromeres to date. The analysis provided both technical and biological insights. As expected, we observed considerable variation in the content and length of the α-satellite HOR array (up to 37-fold for Chromosome 10) consistent with its higher mutation rate and more rapid evolutionary turnover^2,88^. We also, however, document recent *Alu,* LINE-1, and SVA retrotransposition into the α-satellite HORs and show that these may be used to tag HOR expansions on particular human haplotypes. Using the CDR^84,91^ as a marker of kinetochore attachment, we show considerable variation in the location across human centromeres and remarkably that 7% of human chromosomes show evidence of two or more putative kinetochores (i.e., di-kinetochores). The significance of both MEIs and di-kinetochore on chromosome segregation or mis-segregation will need to be experimentally assessed and these phased genomes (and their corresponding cell lines) provide the foundation for such future work.

Finally, from a technical perspective, it should be noted that using two independent assembly algorithms, hifiasm (UL) and Verkko, nearly doubled the number of sequence-resolved centromeres. While the two methods were strongly complementary for centromeres, Verkko was clearly superior for Chromosome Y (**Supplementary Fig. 22c**). Therefore, the assembly performance differs by sequence class and, at present, there is benefit in applying both assembly algorithms in order to fully resolve the most structurally complex regions of the genome. While the performance of both Verrko and hifiasm has been shown to be very similar for large portions of the euchromatin^9^, it is likely that similar analyses of problematic/challenging regions will benefit from applying both assembly algorithms. More method development is warranted to devise an assembly tool combining the strengths of both methods.

## Methods

### Data availability

All data produced by the HGSVC and analyzed as part of this study are available under the following accessions (see **Supplementary Tables 2 -4, 6,7** for details): PacBio HiFi and ONT long reads: PRJEB58376, PRJEB75216, PRJEB77558, PRJEB75190, PRJNA698480, RJEB75739, PRJEB36100, PRJNA988114, PRJNA339722, PRJEB41778, ERP159775; Strand-seq: PRJEB39750, PRJEB12849; Bionano Genomics: PRJNA339722, PRJEB41077, PRJEB58376, PRJEB77842; Hi-C: PRJEB39684, PRJEB75193, PRJEB58376; PacBio Iso-Seq: PRJEB75191; RNA-seq: PRJEB75192, PRJEB58376. Released resources including simple and complex variant calls, genome graphs, genotyping results (genome-wide and targeted), and annotations for centromeres, MEIs, and SDs can be found in the IGSR release directory hosted publicly via HTTP and/or FTP (https://ftp.1000genomes.ebi.ac.uk/vol1/ftp/data_collections/HGSVC3/release) and on the Globus endpoint "EMBL-EBI Public Data" in directory "/1000g/ftp/data_collections/HGSVC3/working".

### Code availability

PAV is available through GitHub (github.com/EichlerLab/pav) as well as Docker and Singularity containers (see documentation on GitHub for container locations). Code for various parts of this work is available through GitHub, including those for sample selection (github.com/tobiasrausch/kmerdbg and github.com/asulovar/HGSVC3_sample_selection); Verkko genome assembly [github.com/core-unit-bioinformatics/workflow-smk-genome-hybrid-assembly (prototype branch)]; assembly evaluation [github.com/core-unit-bioinformatics/workflow-smk-assembly-evaluation (prototype branch)]; project-specific code for assembly-related evaluations, supplementary tables, and plots (github.com/core-unit-bioinformatics/project-run-hgsvc-assemblies); PanGenie genotyping and reference panel construction (github.com/eblerjana/hgsvc3); MEI, MHC, Iso-Seq, and SMN analysis (github.com/Markloftus/HGSVC3 and github.com/Markloftus/L1ME-AID); MHC annotation (github.com/DiltheyLab/MHC-annotation); MEI calling (PALMER2: github.com/WeichenZhou/PALMER and MELT-LRA: github.com/Scott-Devine/MELT-LRA); and SD analysis (SDA2: https://github.com/ChaissonLab/SegDupAnnotation2).

### Sample selection

A total of 65 samples were included in the current study. The majority (63/65) of the samples originate from the 1kGP Diversity Panel^10^, one (NA21487) from the International HapMap Project^11^, and one (NA24385, also called HG002) commonly used for benchmarking by the Genome in a Bottle (GIAB) consortium^12^ was included in all analyses with publicly available data from other efforts (**Supplementary Tables 1 -4, 6, 7**). Samples were selected to maximize genetic diversity and Y-chromosome lineages (**Supplementary Methods**).

### Data production

In addition to data generated through previous efforts^6,36^, sequencing libraries were prepared from high-molecular-weight DNA or RNA extracted from lymphoblast lines (Coriell Institute). PacBio HiFi sequencing data were generated on the Sequel II or Revio platforms using 30-hr movie times. ONT ultra-long libraries were generated using a modified fragmentase protocol and sequenced on R9.4.1 flow cells on a PromethION instrument for 96 hrs. Bionano Genomics optical mapping data using DLE-1 tagging were collected on Saphyr 2nd generation instruments. Strand-seq data were produced using BrdU incorporation and second-strand DNA removal during PCR-based library construction to generate single-nucleus barcoded libraries sequenced on an Illumina NextSeq 500 platform^98,99^. Hi-C data were collected using Proximo Hi-C kits v4.0 (Phase Genomics) and sequenced on an Illumina NovaSeq 6000. RNA-seq libraries were generated using KAPA RNA Hyperprep with RiboErase (Roche) and sequenced on an Illumina NovaSeq 6000 platform. Iso-Seq full-length cDNA libraries were created with the Iso-Seq Express protocol and sequenced on a PacBio Sequel II system. Detailed descriptions of materials and methods are available (Supplementary Methods).

### Assembly

We produced fully phased hybrid assemblies using Verkko^8^ (v1.4.1) as our primary assembler. For some of the most challenging regions (centromeres and Yq12), we additionally created hifiasm (UL)^9^ (v0.19.6) assemblies that were manually evaluated in those regions. The phasing signal for all assemblies was generated using the Graphasing pipeline^13^ (v0.3.1-alpha). All assemblies were scanned for contamination with NCBI’s Foreign Contamination Screening workflow^100^ (v0.4.0) and annotated for potential assembly errors using Flagger^1^ (v0.3.3), Merqury^16^ (v1.0), NucFreq^15^ (commit #bd080aa), and Inspector^17^ (v1.2). Assembly quality was assessed by computing QV estimates with Merqury and DeepVariant^101^ (v1.6) as described previously^6^. Gene completeness of the assemblies was evaluated using compleasm^14^ (v0.2.5) and the primate set of known single-copy genes of OrthoDB^102^ (v10). The T2T status of the assembled chromosomes and the closing status of previously reported gaps^2^ was determined relative to T2T-CHM13 reference genome^7^ by factoring in the above QC information in the evaluation of the contig-to-reference alignment produced with minimap2^103,104^ (v2.26) and mashmap^105^ (v3.1.3). The parental support for the assembled child haplotypes in the three family trios was computed by evaluating the CIGAR operations in the minimap2 contig-to-contig alignments between the parents and child.

### Variant calling

#### Genome reference

Callsets were constructed against two references, GRCh38 (GRCh38-NoALT) and T2T-CHM13 (T2T-CHM13v2.0)^7^.

#### Variant discovery and merging

For assembly-based callsets, we ran PAV^6^ (v2.4.1) with minimap2^103^ (v2.26) and LRA^106^ (v1.3.7.2) alignments, DipCall^107^ (v0.3), and SVIM-asm^108^ (v1.0.3). SVIM-asm used PAV alignments before PAV applied any alignment trimming, and DipCall produced minimap2 alignments for DipCall variants.

For PacBio HiFi callsets, we ran PBSV (https://github.com/PacificBiosciences/pbsv; v2.9.0), Sniffles^109^ (v2.0.7), Delly^110^ (v1.1.6), cuteSV^111^ (v2.0.3), DeBreak^112^ (v1.0.2), SVIM^113^ (v2.0.0), DeepVariant^101^ (v1.5.0), and Clair3^114^ (v1.0.4). The same callers and versions were run for ONT except for PBSV and DeepVariant was executed through PEPPER-Margin-DeepVariant^115^ (r0.8). The callset process was the same for both references.

SV-Pop^6^ was used to merge PAV calls from minimap2 alignments and generate per-sample support information from all other callers. Calls in T2T-CHM13 were filtered if they intersected the UCSC “CenSat” track for T2T-CHM13 (UCSC hs1) with monomeric (“mon”) records excluded or if they were in telomere repeats. GRCh38 variants intersecting modeled centromeres were removed.

### Mobile element insertions

Mobile element insertions (MEIs) were identified within the 130 sample haplotype assemblies using two separate pipelines and human references (T2T-CHM13 and GRCh38). One detection pipeline, L1ME-AID (v1.0.0-beta) (L1 Mediated Annotation and Insertion Detector) (**Code Availability**), leverages a local RepeatMasker^116^ (v4.1.6) installation with the Dfam (v3.8) database^117^ to annotate the freeze4 PAV-merged SV insertion callsets (T2T-CHM13 and GRCh38). The second pipeline called MEIs directly from the sample PacBio HiFi raw sequence reads with PALMER2 (**Code Availability**). Putative MEIs from both callers were merged using MEI coordinates, element family (*Alu*, L1, SVA, HERV-K, or snRNA), and sequence composition (**Supplementary Methods**). Next, MEIs were curated to distinguish MEIs from deletions (T2T-CHM13 or GRCh38), duplications, or potential artifacts (e.g., possible genome assembly errors; **Supplementary Methods**). All MEIs called by a single pipeline that passed QC were manually curated. Finally, both callsets were compared against an orthogonal MEI callset produced by MELT-LRA (**Supplementary Methods**, **Code Availability**). To determine intact ORFs across LINE-1 elements, we followed a previously described method^6^ to detect intact ORF1p and ORF2p from full-length (>5,900 bp) LINE-1 insertions.

Separately, MEIs within centromere HOR arrays were identified with RepeatMasker^116^ (v4.1.6), and the Dfam library^117^ (v3.8), annotation of complete and accurately assembled centromeres (**see Centromeres Methods**). The sequences of *Alu* elements, L1s, and SVAs identified by RepeatMasker within the centromere HOR array boundaries were retrieved using SAMtools^118^ (v1.15.1). Element sequences were then scrutinized with L1ME-AID (v1.0.0-beta) utilizing the same cutoffs applied to the freeze4 PAV-merged SV insertion callset to distinguish young MEIs from older mobile element fragments. Sequence of all putative MEIs that passed filtering were re-retrieved along with flanking sequence (+/-100 bp) using SAMtools^119^ (v1.15.1), and then aligned against one another using MUSCLE^120^ (v3.38.31) to distinguish unique MEIs from duplicated insertions of MEIs residing in centromere regions (**Supplementary Table 60**).

### Inversions

We performed validation of the T2T-CHM13-based and GRCh38-based PAV inversion callsets, individually, using Strand-seq-based re-genotyping of the inversion calls. Prior to genotyping, we performed Strand-seq cell selection using ASHLEYS^121^. The good quality Strand-seq cells were used as input to perform genotyping by ArbiGent^22^.

We evaluated the PAV inversion callset for one candidate carrier sample per region using manual dotplot analysis with NAHRwhals^23^. NAHRwhals was applied to detect the FDR and classify all candidate inversion regions larger than 5 kbp into distinct inversion classes.

We compared the PAV inversion callset reported with respect to T2T-CHM13 to a previously published callset^24^ based on a subset of samples reported in this study. Using the 25% reciprocal overlap criterion, we define inversions detected in both callsets as well as inversions that are new to the current study. We evaluate all novel inversion candidates manually using dotplot analysis of each putative novel inversion.

### Segmental duplication and copy number polymorphic genes

#### Identification of segmental duplications (SDs)

SD annotation was performed using SEDEF^122^ (v1.1) after masking repeats [TRF^123^ (v.4.1.0), RepeatMasker^124^ (v4.1.5), and Windowmasker^125^ (v2.2.22)]. SDs with sequence identity >90%, length >1 kbp, satellite content <70%, and free of putative erroneous regions (NucFreq or Flagger) were retained. Additionally, the highly confident SD callset was further validated by fastCN. Comparative analysis of SDs was conducted in T2T-CHM13 space. Positions of the SDs in T2T-CHM13 were mapped as follows: 1) linking SDs within 10 kbp distance, 2) identifying those SD chains that are located in alignment block of at least 100 kbp in size, and 3) projecting the chained SDs onto putative homologous SD loci containing at least one 10 kbp unique flank. Additionally, syntenic SDs were further assessed whether they share sequence content by aligning SDs with minimap2^103^ (v2.26); the following SDs were quantified: 1) SDs unobserved by T2T-CHM13, 2) having changed sequence content, and 3) expanded size (at least twofold).

#### Duplicated genes

Protein-coding transcripts from GENCODE v44 were aligned to the genome assemblies (excluding NA19650, NA19434 and NA21487) using minimap2. The mapped genes were further filtered to exclude alignments due to nested repeats, keeping minimum length of 2 kbp, percent identity of >90%, and coverage of >80%. Multi-copy genes were determined by maximum gene counts greater than one. Variable copy number genes were defined by assessing the copy number across the population (at least one of the genome assemblies with different copy number).

### Y chromosome variation

#### Construction and dating of Y phylogeny

The construction and dating of Y-chromosomal phylogeny combining the 30 males from the current study plus two males (HG01106 and HG01952 from the HPRC year 1 dataset for which contiguous Yq12 assemblies were used from Ref. 36) was done as previously described^36^. Detailed descriptions of methods are available (**Supplementary Methods**). Please note that the male sample HG03456 appears to have a XYY karyotype as reported in Ref. 45.

#### Identification of sex-chromosome contigs

Contigs containing Y-chromosomal sequences from the whole-genome assemblies were identified and extracted for the 30 male samples as previously described^36^. Y assemblies for the two HPRC samples HG01106 and HG01952 were used from Ref. ^36^.

#### Y chromosome annotation and analysis

The annotation of Y-chromosomal subregions was performed as previously described using both the GRCh38 and T2T-CHM13 Y reference sequences^36^. The centromeric α-satellite repeats for the purpose of Y subregion annotation were identified using RepeatMasker^116^ (v4.1.2-p1). The Yq12 repeat annotations were generated using HMMER v.3.3.2dev^126^ and identification of Alu insertions was performed as previously described^36^. In order to maximize the number of contiguously assembled Yq12 subregions, hifiasm assemblies of this subregion were analyzed from four samples (NA19239, HG03065, NA19347, HG00358) following manual inspection of the assembled sequences (**Supplementary Table 42**).

Dotplots to compare Y-chromosomal sequences were generated using Gepard v2.0^127^. While we also assembled the T2T (NA24385/HG002) Y as a single contig (**Supplementary Table 42**), all analyses conducted here used the existing published T2T assembly^37^.

Visualization of eight completely assembled Y chromosomes (**Supplementary Fig. 23**) was based on pairwise alignments generated using minimap2^103,104^ (v2.26) with the following options: -x asm20 -c -p 0.95 --cap-kalloc=1g -K4g -I8g -L --MD --eqx. For visualization, alignments <10 kbp in length were filtered out. Additionally, alignments were broken at SVs ≥ 50 bp in size and then binned in 50 kbp bins.

### SVs affecting genes

We annotated the potential impact of long-read SVs on genes using the coding transcripts and exons defined in GENCODE^28^ (v39), as per Ensembl VEP^128^ (v105). Long-read deletions or insertions are classified as coding overlapping events if at least one breakpoint falls within the coding exons of a gene. We consider genes that have a LOEUF score under 0.35 as intolerant to LoF variants^39^. To specifically analyze the potential impact of MEIs on genes, the merged GRCh38 MEI callset was intersected with the findings from Ensembl^129^ (release 111) VEP^128^ (see transcriptional effect of SVs below). The MEIs were categorized by insertion location (e.g., protein-coding exons, UTRs of protein-coding transcripts, and noncoding exons) and within each category the number of MEIs present, genes disrupted, and transcripts affected were quantified. The Ensembl VEP nonsense mediated decay (NMD) plugin (https://github.com/Ensembl/VEP_plugins/blob/release/112/NMD.pm) was utilized to predict which protein-coding transcripts with MEI-induced premature stop codons would escape NMD. Transcripts were further scrutinized by manually comparing the MEI location within the transcript sequence using the UCSC Genome Browser^130^. Allele frequencies were then calculated (children of trios excluded) for the exon-disrupting MEIs (**Supplementary Methods**).

### Transcriptional effect of SVs

We used the Ensembl^129^ (release 111) Variant Effect Predictor^128^ with NMD plugin to screen the PAV freeze 4 callset for SVs that disrupt gene loci in the merged GRCh38 annotation (**Supplementary Methods**). Protein-coding genes impacted by putative exon disruptions were evaluated for evidence of Iso-Seq expression across the 12 samples. Isoforms associated with these SV-containing genes were screened for the presence of unreported splice variants using SQANTI3^131^ (v5.1.2). All isoforms of these candidate genes were aligned to GRCh38p14 using pbmm2 (https://github.com/PacificBiosciences/pbmm2) (v1.5.0) and visualized with IGV^132^ to identify variant-specific patterns. We compared all isoforms phased to variant haplotypes to known transcripts represented in RefSeq^27^, CHESS^29^, and GENCODE^28^ gene annotation databases to identify novel splice products and isoforms. MUSCLE^120^ (v3.8.425) and Aliview^133^ were used to perform a multiple sequence alignment (MSA) and visualize the MSA, respectively, between wild-type and variant haplotype assemblies to identify SV breakpoints.

We next assessed SVs for enrichment near genes with altered expression in the 12 samples with Iso-Seq data. Using gene expression quantifications from short-read RNA-seq data, we performed differential expression analysis using DESeq2^134^ between individuals who carried and did not carry each SV, supplemented with outlier expression analysis for singleton SVs, and identified 122 SVs that were within 50 kbp of a GENCODE v45 annotated gene with significantly altered expression (Benjamini-Hochberg adjusted p<0.05)^28^. Permutation tests showed an enrichment of SVs (empirical p<0.001) adjacent to genes with differential expression (**Supplementary Methods**: “Transcriptional Effects of SVs”). Additionally, we assessed SV overlap with multiple GENCODE v45-derived genomic elements, such as protein-coding and pseudogene classes^28^, and ENCODE-derived candidate cis-regulatory elements^135^ (cCREs), using permutation tests to find enrichment or depletion of SVs for each annotation.

Among the 128 SV gene pairs (122 unique SVs associated with 98 genes) that exhibit significant differential gene expression changes in the 12 samples with Iso-Seq data, we first filtered out SVs with missing genotypes in 6 or more out of 12 samples. For each remaining SV, we extracted the 50 kbp upstream and downstream of the annotated transcription start site (TSS) position for each paired gene with corresponding insulation scores under 10 kbp resolution (**Supplementary Methods**). For those insulated regions intersecting with more than one SV, we applied a local multi-test correction. An FDR <0.05 from the two-sided Wilcoxon rank-sum test was considered significant. We investigated the association between variants and human phenotypes or traits by intersecting SNVs, indels, and SVs with SNPs identified in GWAS. We downloaded the GWAS summary statistics (gwas_catalog_v1.0.2-associations_e111_r2024-04-16.tsv)^43^, which contains 149,081 autosomal SNPs and we observed 1,547 SVs are at least one bp overlap with any GWAS signal. We used Plink^136^ (v1.90b6.10) to examine the linkage disequilibrium between SNVs, indels, and SVs with GWAS SNPs within 1 Mbp window size.

### Genome-wide genotyping with PanGenie

We built a pangenome graph containing 214 haplotypes with Minigraph-Cactus^44^ (v2.7.2) from the haplotype-resolved assemblies of 65 HGSVC samples and 42 samples from the HPRC^1^ and produced a CHM13-based VCF representation of the top-level bubbles of the graph that can be used as input for genotyping with PanGenie (**Supplementary Methods**). This was done by converted genotypes of male sex chromosomes to a homozygous representation, filtering out records for which at least 20% of haplotypes carry a missing allele (".") and running our previously developed decomposition approach to detect and annotate variant alleles nested inside of graph bubbles (**Supplementary Methods**). We genotyped all 30,490,169 bubbles (representing 28,343,728 SNPs, 10,421,787 indels and 547,663 SVs) across all 3,202 1kGP samples based on short reads^45^ using PanGenie^3^ (v3.1.0). We filtered the resulting genotypes based on a support vector regression approach^1,6^, resulting in 25,695,951 SNPs, 5,774,201 indels, and 478,587 SVs that are reliably genotypable (**Supplementary Methods**).

### Personal genome reconstruction from an integrated reference panel

We used our filtered genotypes across all 3,202 samples and added 70,174,243 additional rare SNPs and indels from an external short-read-based callset for the same 3,202 1kGP samples (obtained from: https://s3-us-west-2.amazonaws.com/human-pangenomics/index.html?prefix=T2T/CHM13/asse mblies/variants/1000_Genomes_Project/chm13v2.0/all_samples_3202/) (**Supplementary Methods**). We filtered out variants reported with a genotype quality below 10 and ran SHAPEIT5^46^ phase_common (v5.1.1) in order to phase this joint callset. We used the resulting reference panel in order to reconstruct personal genomes for all 3,202 samples by implanting phased variants into the CHM13 reference genome with BCFtools^118^ in order to create the 6,404 consensus haplotype sequences of all 1kGP samples (**Supplementary Methods**).

### Targeted genotyping of complex polymorphic loci

Targeted genotyping was performed using Locityper^48^ (v0.15.1) across 347 complex polymorphic target loci. Based on the input short-read WGS data, at each of the targets Locityper aims to identify two haplotypes from the reference panel that are most similar to the input data. Three reference panels were used: HPRC haplotypes (90 haplotypes); HPRC + HGSVC3 haplotypes (216 haplotypes); and leave-one-out HPRC + HGSVC3 panel (LOO; 214 haplotypes), where two assemblies corresponding to the input dataset were removed. To evaluate prediction accuracy, we constructed sequence alignments between actual and predicted haplotypes and estimated QVs as Phred-like transformed sequence divergences. Locityper accuracy is limited by haplotypes present in the reference panel; consequently, we evaluated haplotype availability QV as the highest Phred-scaled sequence divergence between actual assembled haplotypes and any haplotype from the reference panel. See **Supplementary Methods** for more details.

### Major histocompatibility complex

Immuannot-based HLA types were compared in two-field resolution to the HLA typing published earlier and obtained with PolyPheMe^53^ (http://ftp.1000genomes.ebi.ac.uk/vol1/ftp/data_collections/HLA_types/20181129_HLA_types_fu <u>ll_1000_Genomes_Project_panel.txt</u>). Of the 130 haplotypes, 58 are not in the PolyPheMe dataset and were excluded. In addition to Immuannot (MHC reference version: IPD-IMGT/HLA-V3.55.0)^54^, haplotypes were annotated using MHC-annotation v0.1 (**Code Availability**). Cases of overlapping genes were resolved after inspection by removing superfluous annotations. Reported gene counts for HLA genes and C4 annotation were based on Immuannot.

To search for structural variation in the DRB gene region, HGSVC MHC haplotypes were cut from [start of DRA] to [end of DRB1 + 20 kbp]. The coordinates were obtained using MHC-annotation v0.1. Based on their DRB1-allele as determined by Immuannot (see above), the sequences were grouped into DR groups. Within each group, every sequence was aligned with nucmer^137^ (v3.1) (-nosimplify -maxmatch) to the same sequence (arbitrarily selected as the sequence with the alphanumerically smallest ID) and plotted with a custom gnuplot script based on mummerplots output. Sequences were annotated as follows:

1. Repeat elements were masked with RepeatMasker^116^ (v4.1.2)
2. Full DRB genes and pseudogenes were searched for with minimap 2.26 (--secondary=no -c -x --asm10 -s100) by aligning the sequence from (1) against all DRB alleles from IMGT and the larger *DRB9* sequence Z80362.1; results were highlighted and masked for the next step
3. DRB exons were searched for with BLASTN^138^ (2.14.1) by aligning all DRB exons from IMGT to the sequence and filtering for highest matches; results where highlighted and masked for the next step
4. As step 3 but with introns

For each HGSVC MHC haplotype, SVs were called with PAV^6^ against eight completely resolved MHC reference haplotypes^58,59^. To determine which SVs in the HGSVC haplotypes were not present in any of the eight reference haplotypes, for each HGSVC haplotype, the “query” coordinates (i.e., the coordinates of the calls relative to the analyzed HGSVC haplotype) of the PAV calls were padded with 50 bp on each side and the intersection of SV calls (based on the padded “query” coordinates, across the eight MHC reference sequences) was computed. Only variants longer than 50 bp were included for further analysis and the smallest variant relative to any of the eight MHC references was reported. The sequences of the calls so-defined were annotated with RepeatMasker^116^ (v4.1.2).Variants were grouped by starting position on their closest MHC reference sequence and, in the case of insertions, repeat content was averaged.

We applied Immuannot (see above) to all retrieved MHC loci for the identification and annotation of protein-coding HLA-DRB genes (-*DRB1*, -*DRB3*, -*DRB4*, and -*DRB5*). Subsequently, a custom RepeatMasker^116^ (v4.1.2) library was constructed containing the exonic sequences of HLA-DRB pseudogenes (-*DRB2*:ENSG00000227442.1, -*DRB6*:ENSG00000229391.8, -*DRB7*:ENSG00000227099.1, -*DRB8*:ENSG00000233697.2, and

-*DRB9*:ENSG00000196301.3) and RCCX genes and pseudogenes (*C4*:ENSG00000244731.10, *CYP21A2*:ENSG00000231852.9, *CYP21A1P*:ENSG00000204338.9, *STK19*/*RP1*:ENSG00000204344.16, *STK19B*/*STK19P*/*RP2*:ENSG00000250535.1, *TNXB*:ENSG00000168477.21, and *TNXA*:ENSG00000248290.1). Canonical exonic sequences were sourced from the Ensembl genome browser^129^ (release 111). The exons of HLA-DRB/RCCX genes and pseudogenes within individual haplotype MHC regions were annotated using this custom library. Repetitive elements were identified using RepeatMasker (v4.1.2) with the Dfam library^117^ (v3.4). We utilized SAMtools^118^ (v1.15.1) and MUSCLE^120^ (v3.8.31) for sequence retrieval and alignment, respectively, followed by manual annotation to analyze recombination events associated with DR subregion haplotypes and within the RCCX modules (**Supplementary Methods**). Novel *C4*-coding variants were identified through comparison with Ensembl *C4A* and *C4B* protein variant tables, as well as an additional database of variants obtained from 95 human MHC haplotypes^51^.

### Complex structural polymorphisms

Complex structural variants (CSVs) were identified with a development version of PAV (methods available at 10.5281/zenodo.13800981). Briefly, the method identifies candidate variant anchors and scores variants between them. A directed acyclic graph (DAG) is constructed with alignment records as nodes and variants connecting them as edges, which is solved in *O*(N + E) time with the Bellman-ford algorithm^139^. Variants on the optimal path are accepted into the callset. CSVs intersecting centromeric repeats were eliminated. CSVs were merged into a nonredundant callset with SV-Pop by 50% reciprocal overlap and 80% sequence identity.

We evaluated complexity and copy number of SMN genes by extracting with FASTA the desired region (chr5:70300000-72100000) from assemblies reported in this study along with previously published assemblies^1,140^. Among these, we identify 101 fully assembled haplotypes. We follow by aligning exon sequences for multicopy genes (*SMN1/2, SERF1A/B, NAIP* and *GTF2H2/C*) to each assembled haplotype. To assign a specific SMN copy to each haplotype, we extracted FASTA sequence from SMN exon regions for each haplotype and concatenated them into a single sequence. We then constructed a MSA and calculated the distance among all haplotypes. We set the orangutan sequence as an outgroup and split all human haplotypes into two groups representing *SMN1* and *SMN2* gene copies where the SMN1 copy is the one closer to the outgroup.

### Centromeres

#### Centromere identification and annotation

To identify the centromeric regions within each Verkko and hifiasm (UL) genome assembly, we first aligned the whole-genome assemblies to the T2T-CHM13 (v2.0) reference genome^7^ using minimap2^103^ (v2.24) with the following parameters: -ax asm20 --secondary=no -s 25000 -K 15G --eqx --cs. We filtered the alignments to only those contigs that traversed each human centromere, from the p- to the q-arm, using BEDtools^141^ (v2.29.0) intersect. Then, we ran dna-brnn^142^ (v0.1) on each centromeric contig to identify regions containing α-satellite sequences, as indicated by a “2”. Once we identified the regions containing α-satellite sequences, we ran RepeatMasker^116^ (v4.1.0) to identify all repeat elements and their organization within the centromeric region; we also ran Hum-AS-HMMER (https://github.com/fedorrik/HumAS-HMMER_for_AnVIL) to identify α-satellite HOR sequence composition and organization. We used the resulting RepeatMasker and HumAS-AMMER stv_row.bed file to visualize the organization of the α-satellite HOR arrays with R^143^ (v1.1.383) and the ggplot2 package^144^.

#### Validation of centromeric regions

We validated the construction of each centromeric region by first aligning native PacBio HiFi and ONT data from the same genome to each relevant whole-genome assembly using pbmm2 (v1.1.0; for PacBio HiFi data; https://github.com/PacificBiosciences/pbmm2) or minimap2^103^ (v2.28; for ONT data). We, then, assessed the assemblies for uniform read depth across the centromeric regions via IGV^132^ and NucFreq^15^. Centromeres that were found to have a collapse in sequence, false duplication of sequence, and/or misjoin were flagged and removed from our analysis.

#### Estimation of α-satellite HOR array length

To estimate the length of the α-satellite HOR arrays for each human centromere, we first ran Hum-AS-HMMER (https://github.com/fedorrik/HumAS-HMMER_for_AnVIL) on the centromeric regions using the hmmer-run.sh script and the AS-HORs-hmmer3.0-170921.hmm Hidden Markov Model. Then, we used the stv_row.bed file to calculate the length of the α-satellite HOR arrays by taking the minimum and maximum coordinate of the “live” α-satellite HOR arrays, marked by an “L”, and plotting their lengths with Graphpad Prism (v9).

#### Pairwise sequence identity heatmaps

To generate pairwise sequence identity heatmaps of each centromeric region, we ran StainedGlass^145^ (v6.7.0) with the following parameters: window=5000, mm_f=30000, and mm_s=1000. We normalized the color scale across the StainedGlass plots by binning the percent sequence identities equally and recoloring the data points according to the binning.

#### CpG methylation analysis

To determine the CpG methylation status of each centromere, we aligned ONT reads >30 kbp in length from the same source genome to the relevant whole-genome assembly via minimap2^103^ (v2.28) and then assessed the CpG methylation status of the centromeric regions with Epi2me modbam2bed (https://github.com/epi2me-labs/modbam2bed; v0.10.0) and the following parameters: -e -m 5mC --cpg. We converted the resulting BED file to a bigWig using the bedGraphToBigWig tool (https://www.encodeproject.org/software/bedgraphtobigwig/) and then visualized the file in IGV. To determine the length of the hypomethylated region (termed “centromere dip region”, or CDR^84,91^ in each centromere, we used CDR-Finder (https://github.com/EichlerLab/CDR-Finder_smk). This tool first bins the assembly into 5 kbp windows, computes the median CpG methylation frequency within windows containing α-satellite [as determined by RepeatMasker^116^ (v4.1.0)], selects bins that have a lower CpG methylation frequency than the median frequency in the region, merges consecutive bins into a larger bin, filters for merged bins that are >50 kbp, and reports the location of these bins.

## Acknowledgements

Funding was provided by National Institutes of Health (NIH) grants U24HG007497 (to S.E.H., O.A., L.S., E.E.E., S.E.D., B.G., and C.L.), GM147352 (to G.A.L.), R01HG011649 (to M.J.P.C. and B.G.), K99HG012798 (to H.C.); NIH National Institute of General Medical Sciences R35GM133600 (to P.A.A., P.B., and C.R.B.), 1P20GM139769 (to M.K.K. and M.L.), 1R35GM138212 (to Z.C.); NIH National Institute of Allergy and Infectious Disease (NIAID) U01AI090905 (to A.T.D., T.P., L.A.G., and P.J.N.); NIH National Cancer Institute (NCI) R01CA261934 (to J.C. and S.E.D.), P30CA034196 (to P.A.A. and C.R.B.); National Science Foundation (NSF) CAREER 2046753 (to M.J.P.C. and K.R.), the Ministry of Culture and Science of North Rhine-Westphalia (MODS, “Profilbildung 2020” [grant no. PROFILNRW-2020–107-A]) (to A.S. and T.M.), and the German Research Foundation (DFG) grant 496874193 (to T.M.). This work was also supported, in part, by the Intramural Research Program of the National Human Genome Research Institute, NIH (to A.M.P. and S.K.), the Jürgen Manchot Foundation (to S.T.S. and A.T.D.), the Düsseldorf School of Oncology (grant SPATIAL to T.M.). E.E.E. is an investigator of the Howard Hughes Medical Institute. We thank the Centre for Information and Media Technology at Heinrich Heine University Düsseldorf for providing their computational infrastructure and support; the staff at Clemson University for their allotment of compute time on the Palmetto HPC; HPC resources at Temple University supported in part by the National Science Foundation through major research instrumentation grant number 1625061 and by the US Army Research Laboratory under contract number W911NF-16-2-0189; the staff at the Scientific Services at the Jackson Laboratory, including the Genome Technologies Service for their assistance with the work described herein; the members of the HPRC (https://humanpangenome.org) for making their data publicly available; and the people who contributed samples as part of the 1000 Genomes Project.

## Author contributions

Sample Selection: P.H., K.M.M., T.R., A.SV., C.Lee., and E.E.E.; Data Production: K.M.M., P.H., P.H.F., K.H., Q.Z., S.E.D.; Data Management: P.E., P.A.A., S.E.H., P.H., F.Y., K.M.M., Y.K., O.A., and L.S.; Assembly Production and Quality Control: P.E., W.T.H., M.H., Z.C., M.R., S.K., Y.K., H.C., A.M.P., Y.S., E.E.E., and T.M.; Variant Discovery: P.A.A, C.A.P., and C.R.B.; Mobile Elements: M.L., W.Z., P.B., R.E.M., J.C., S.E.D., C.R.B., and M.K.K.; Inversions: H.A., V.T., D.P., T.R., J.O.K., and T.M.; Segmental Duplications: D.Y., K.R., M.J.P.C., and E.E.E.; STR and VNTR Annotation: B.G. and M.J.P.C.; Chromosome Y: P.H., P.E., M.L., M.K.K., and C.Lee.; Iso-Seq Phasing: G.V.M., M.L., and M.K.K.; SV Impact on Genes: X.Z., G.V.M., M.L., M.E.T., and M.K.K.; Transcriptional Effects of SVs: G.V.M., M.L., M.J., Y.J., J.L., M.G., and M.K.K.; Hi-C and Additional Functional Analysis: C.LI., M.J.B., and X.S.; Genotyping: J.E., T.P., G.H., B.P., and T.M.; Integrated Reference Panel: J.E., T.R., M.C.Z. and T.M.; Major Histocompatibility Complex: M.L., S.T.S., C.S., Y.Z., N.R.P., P.J.N., L.A.G., P.A.A., P.E., A.S., T.P., C.R.B., H.L., T.M., M.K.K., and A.T.D.; Complex Structural Polymorphisms: P.A.A., D.P., F.Y., M.L., M.K.K., C.R.B., C.Lee., and E.E.E.; Centromeres: G.A.L., K.K.O., M.L., M.K.K., and E.E.E.; Manuscript Writing: G.A.L., P.E., P.A.A., M.L., D.P., J.E., F.Y., P.H., T.P., D.Y., X.Z., G.V.M., C.S., H.A., M.J., C.LI., X.S., M.E.T., M.J.P.C., A.T.D., M.K.K., J.O.K., C.Lee., C.R.B., E.E.E., and T.M. All authors read and approved the final manuscript. HGSVC co-chairs: J.O.K., C.Lee., E.E.E. and T.M.

## Competing Interests

E.E.E. is a scientific advisory board member of Variant Bio, Inc. C. Lee is a scientific advisory board member of Nabsys and Genome Insight. S.K. has received travel funds to speak at events hosted by Oxford Nanopore Technologies. The following authors have previously disclosed a patent application (No. EP19169090) relevant to Strand-seq: J.O.K., T.M., and D.P. The other authors declare no competing interests.

## Notes

http://ftp.1000genomes.ebi.ac.uk/vol1/ftp/data_collections/HGSVC3/release/

