## Supplementary Text and Figures for "Complex genetic variation in nearly complete human genomes"


**Note on contributing authors** **2**
**Sample selection** **2**
Methods 2
**Data production and organization** **3**
Methods 3
**Assembly production and quality control** **6**
Methods 6
Results 16
**Variant discovery and callset development** **18**
Methods 18
Results 23
**Mobile elements** **23**
Methods 23
Results 25
**Inversions** **26**
Methods 26
Results 26
**Segmental duplications** **27**
Methods 27
Results 29
**STR and VNTR annotation** **29**
Methods 30
Results 30
**Y chromosome** **31**
Methods 31
**SVs disrupting genes** **32**
**Phased transcripts, isoforms, and effects of SVs** **34**
Methods 34
Results 35
**Transcriptional effects of SVs and functional analysis** **37**
Methods 37
Results 40
**Genotyping** **42**
Methods 42
**Major Histocompatibility Complex** **51**
Methods 52
Results 53
**Complex structural polymorphisms** **58**
Methods 58

Results 60
**Centromeres** **62**
Methods 62
**Supplementary Figures** **64**
Fig. 1 64
Fig. 2 65
Fig. 3 66
Fig. 4 67
Fig. 5 68
Fig. 6 69
Fig. 7 70
Fig. 8 71
Fig. 9 72
Fig. 10 73
Fig. 11 74
Fig. 12 75
Fig. 13 76
Fig. 14 76
Fig. 15 77
Fig. 16 77
Fig. 17 78
FIg. 18 79
Fig. 19 80
Fig. 20 81
Fig. 21 82
Fig. 22 83
Fig. 23 84
Fig. 24 86
Fig. 25 88
Fig. 26 89
Fig. 27 91
Fig. 28 92
Fig. 29 93
Fig. 30 94
Fig. 31 95
Fig. 32 96
Fig. 33 97
Fig. 34 98
Fig. 35 98
Fig. 36 99
Fig. 37 100
Fig. 38 101
Fig. 39 102
Fig. 40 103

| | |
|---|---|
| Fig. 41 | 105 |
| Fig. 42 | 106 |
| Fig. 43 | 107 |
| Fig. 44 | 109 |
| Fig. 45 | 110 |
| Fig. 46 | 111 |
| Fig. 47 | 112 |
| Fig. 48 | 113 |
| Fig.49 | 113 |
| Fig. 50 | 114 |
| Fig. 51 | 116 |
| Fig. 52 | 117 |
| Fig. 53 | 118 |
| Fig. 54 | 119 |
| Fig. 55 | 120 |
| Fig. 56 | 120 |
| Fig. 57 | 121 |
| Fig. 58 | 122 |
| Fig. 59 | 123 |
| Fig. 60 | 124 |
| Fig. 61 | 125 |
| Fig. 62 | 126 |
| Fig. 63 | 127 |
| Fig. 64 | 128 |
| Fig. 65 | 129 |
| **Bibliography** | **130** |

### Note on contributing authors

Contributing authors are listed in alphabetical order by surname.

### Sample selection

Contributing authors: Evan E. Eichler, Pille Hallast, Charles Lee, Katherine M. Munson, Tobias Rausch, Arvis Sulovari

#### Methods

A total of 65 samples were included in the current study. The majority (63/65) of the samples originate from the 1000 Genomes Project (1kGP) Diversity Panel[1] and 1/65 (NA21487) from the International HapMap Project[2]. In addition, NA24385 (HG002), commonly used for benchmarking by the Genome in a Bottle (GIAB) consortium[3], was included in all analyses with publicly available data from other efforts (**Supplementary Tables 1-4,6,7**).

Samples were selected for inclusion according to four criteria: (i) Samples (n=32) for which data had previously been generated[4] were updated with additional PacBio HiFi and ultra-long Oxford Nanopore Technologies (ONT) coverage. (ii) Additional samples were chosen from the Reference Genome Improvement project (https://genome.wustl.edu/items/reference-genome-improvement/) (n=8), representing the 1kGP populations ASW, YRI, GWD, MSL, ESN, and ACB, for which deep Illumina coverage exists[5] (n=7). (iii) We calculated population-specific *k*-mers[6] for each 1kGP individual with deep-coverage Illumina data available[5] to quantify the distance of each individual relative to its respective population cluster's centroid (**Code Availability**). Briefly, we conducted a principal components analysis using all of the autosomal variants from all samples using the `SNPrelate` R package[7] (v1.26.0). The ideal number of clusters (k value) was determined using the gap statistic implemented within the `factoextra` R package (github.com/kassambara/factoextra; v1.0.7). The optimal number of clusters for EUR, AFR, EAS, SAS, and AMR samples was found to be 3, 7, 1, 4 and 6, respectively using PC1 and PC2 eigenvalues; this allowed us to compute the distance to the nearest centroid for each sample (**Code Availability**). We then chose additional samples (n=13) either closest or farthest from the centroid of their population cluster. Samples previously chosen for other long-read sequencing efforts (e.g., by the Human Pangenome Reference Consortium, HPRC) were not considered. (iv) Lastly, three samples (NA18989, HG01890, HG00358) were included to represent specific Y-chromosomal haplotypes[8].

### Data production and organization

Contributing authors: Peter A. Audano, Olanrewaju Austine-Orimoloye, Scott E. Devine, Peter Ebert, Pille Hallast, Patrick Hasenfeld, Kendra Hoekzema, Sarah E. Hunt, Youngjun Kwon, Katherine M. Munson, Likhitha Surapaneni, Feyza Yilmaz, Qihui Zhu

### Methods

Data generated as part of this project were derived from lymphoblast lines available from the Coriell Institute for Medical Research for research purposes (https://www.coriell.org/). Regular checks for mycoplasma contamination are performed at the Coriell Institute. Data file and project accession numbers are listed in **Supplementary Tables 2-4,6,7**.

#### PacBio HiFi sequence production.

##### University of Washington.

Data were generated as described previously[8]. For additional coverage, samples HG00514, HG00731, HG00732, NA19238, NA19239, NA19240, and NA18939 were also processed with the same shearing and size selection parameters but using the SMRTbell Prep Kit v3.0 (PacBio P/N 102-182-700) and SMRTbell Barcoded Adapter Plate 3.0 (PacBio P/N 102-009-200) according to manufacturer's recommendations. Additional sequencing was performed on either the PacBio Sequel II platform using chemistry v3.2 (PacBio P/N 102-333-300) with 30-hour movie and 2-hr pre-extension times, or the PacBio Revio platform with chemistry v1 (PacBio P/N 102-817-900) on collection software SMRT Link v12.0 and v13.0, using 30-hour movies and 2- or 1.417-hr pre-extension times. Revio datasets were post-processed onboard through ccs analysis and DeepConsensus v1.0 or v2.0.

##### The Jackson Laboratory

Data were generated as described previously[8]. To generate additional coverage for three samples (NA19238, NA19240 and HG03732), the high-molecular-weight (HMW) DNA was extracted using the Monarch HMW DNA Extraction Kit for cells (NEB), libraries were prepared using SMRTBell Express Template Prep Kit 3.0 (PacBio) with 5 µg of DNA was sheared using Megaruptor 3.0 (Diagenode) and sequenced on either PacBio's Sequel II (all but one) or Revio (HG03732).

##### University of Maryland Institute for Genome Sciences (UMIGS).

Data were generated essentially as described[8]. HMW genomic DNA was isolated using the Circulomics[9] DNA preparation kit following the manufacturer's protocols. Four samples were sequenced (NA19434, NA19836, HG03520, and HG02282) by shearing the HMW gDNA and isolating an average fragment size of 18 kbp. PacBio libraries were prepared from the sheared 18 kbp fragments and sequenced using PacBio's Sequel II platform with a target coverage of 30X coverage (actual coverages = 27.8X, 32.4X, 47.1X,43X, respectively for NA19434, NA19836, HG03520, and HG02282).

#### ONT-UL sequence production.

##### University of Washington.

Data were generated as described previously[8]. HMW DNA was extracted from 2 aliquots of 30 million frozen pelleted cells using the phenol–chloroform approach as described previously[10]. Libraries were prepared using the Ultra-Long DNA Sequencing Kit (SQK-ULK001, ONT) according to the manufacturer's recommendations. In brief, DNA from around 10 million cells was incubated with 6 µl of fragmentation mix at room temperature for 5 min and 75 °C for 5 min. This was followed by an addition of 5 µl of adapter (RAP-F) to the reaction mix and incubated for 30 min at room temperature. The libraries were cleaned up using Nanobind disks (Circulomics) and long fragment buffer (SQK-ULK001, ONT) and eluted in elution buffer. Libraries were sequenced on the flow cell R9.4.1 (FLO-PRO002, ONT) on a PromethION (ONT) for 96 h. A library was split into 3 loads, with each load going 24 h followed by a nuclease wash (EXP-WSH004, ONT) and subsequent reload.

##### The Jackson Laboratory.

Data were generated as described previously[8]. HMW DNA was extracted from 60 million frozen pelleted cells using the phenol–chloroform approach as previously described [11]. Libraries were prepared using the Ultra-Long DNA Sequencing Kit (SQK-ULK001, ONT) according to the manufacturer's recommendations. In brief, 50 µg of DNA was incubated with 6 µl of fragmentation mix at room temperature for 5 min and 75 °C for 5 min. This was followed by an addition of 5 µl of adapter (RAP-F) to the reaction mix and incubated for 30 min at room temperature. The libraries were cleaned up using Nanodisks (Circulomics) and eluted in an elution buffer. Libraries were sequenced on the flow cell R9.4.1 (FLO-PRO002, ONT) on a PromethION (ONT) system for 96 h. A library was generally split into 3 loads with each loaded at an interval of about 24 h or when pore activity dropped to 20%. A nuclease wash was performed using the Flow Cell Wash Kit (EXP-WSH004) between each subsequent load.

##### ONT Basecalling

For genome analysis including genome assembly, ONT reads were basecalled with Guppy 5.0.11 with configuration "dna_r9.4.1_450bps_sup_prom.cfg" distributed with the Guppy software.

For ONT-based methylation analyses, reads were basecalled a second time with Guppy 6.5.7 with configuration "dna_r9.4.1_450bps_modbases_5mc_cg_sup_prom.cfg" distributed with the Guppy software.

#### Bionano Genomics optical genome maps production.

Optical genome mapping data for 17 individuals were generated at Bionano Genomics as described previously[8] (**Supplementary Table 3**, PRJEB77842). We complemented our analysis set with data from previously published HGSVC efforts (n=13, PRJEB58376[8]; n=32, PRJEB41077[4]) and using publicly available data for HG02818 (PRJNA339722) and NA24385/HG002 (**Supplementary Table 3**). Optical genome maps of the seventeen samples were de novo assembled using the Bionano Solve v3.5 (https://bionanogenomics.com/support/software-downloads/) assembly pipeline, with default settings as described previously[4].

```
python2.7 Solve3.5.1_01142020/Pipeline/1.0/pipelineCL.py -T 64
-U -j 64 -jp 64 -N 6 -f 0.25 -i 5 -w -c 3 -y -b ${bionano_bnx}
-l ${output_dir} -t Solve3.5.1_01142020/RefAligner/1.0/ -a
```

```
Solve3.5.1_01142020/RefAligner/1.0/optArguments_haplotype_DLE1
_saphyr_human.xml -r ${reference_genome}
```

A pairwise comparison of DNA molecules (min. 250 kbp) was generated to produce the initial consensus genome maps. During an extension step, molecules were aligned to genome maps, and maps were extended based on the molecules aligning past the map ends. Overlapping genome maps were then merged. Extension and merge steps were repeated five times before a final refinement of the genome maps. Clusters of molecules aligned to genome maps with unaligned ends >30 kbp in the extension step were re-assembled to identify all alleles. To identify alternate alleles with smaller size differences from the assembled allele, clusters of molecules aligned to genome maps with internal alignment gaps of size <50 kbp were detected, and the genome maps were converted into two haplotype maps. The final genome maps were aligned to the reference genome, GRCh38.p12.

#### Strand-seq data generation and data processing.

Data for nine samples were generated as described previously[8]. We complemented our analysis set with previously published data from HGSVC efforts (n=9, PRJEB12849[12]; n=34[4], n=3[8], and n=9[13], PRJEB39750) and publicly available data for NA24385/HG002 (AWS:S3:s3-us-west-2.amazonaws.com/human-pangenomics/index.html?prefix=NHGRI_UCSC_panel/HG002/hpp_HG002_NA24385_son_v1/Strand_seq/2019-12-16-HWVTJAFXY/)

#### Hi-C data production.

In total, Hi-C data for 63 individuals were collected for analysis (**Supplementary Tables 4,5**). Specifically, Hi-C data for 20 individuals (PRJEB75193) were generated in this study using Proximo Hi-C kits v4.0 (Phase Genomics), followed by sequencing at the Jackson Laboratory for Genomic Medicine as described previously[8]. We complemented our analysis set by adding data from previous HGSVC efforts (n=30 using Proximo Hi-C kits v3.0, PRJEB39684[4]; n=10, PRJEB58376[8]) and from one external source for the three individuals NA19238, HG00513 and HG00732 (PRJNA528584[14]).

#### RNA-seq data production.

RNA-seq data were generated at The Jackson Laboratory for 21 individuals (PRJEB75192) as described previously[8], with the exception that 1 million cells were used as a starting material, and paired-end 150 bp reads were sequenced on an Illumina NovaSeq 6000 platform. We complemented our analysis set with previously published data from HGSVC efforts (n=33, PRJEB39684[4]; n=10, PRJEB58376[8]; **Supplementary Table 7**).

#### Iso-Seq data production

Iso-Seq data were generated at The Jackson Laboratory for 12 (PRJEB75191) individuals as described previously[8] (**Supplementary Table 6**).

### Assembly production and quality control

Contributing authors: Haoyu Cheng, Zechen Chong, Peter Ebert, Evan E. Eichler, William T. Harvey, Mir Henglin, Sergey Koren, Youngjun Kwon, Tobias Marschall, Adam M. Phillippy, Mikko Rautiainen, Yuwei Song

#### Methods

##### Verkko hybrid genome assembly

All samples were uniformly processed with a Snakemake v7.19.1[15] workflow implementing the hybrid genome assembly using Verkko[16] (v1.4.1; see **Code Availability**). Verkko was first executed with all PacBio HiFi and ONT data per sample to generate an unphased whole-genome assembly. The resulting assembly graph in GFA format was then forwarded into the previously published Graphasing[17] (v0.3.1-alpha) phasing pipeline that used all available Strand-seq data per sample to generate phasing paths through the assembly graph. The phase information was further processed by the Verkko companion tool Rukki[16] to generate a Verkko-compatible input file. This input file was then used in addition to the read input data for the second Verkko run resulting in fully phased assemblies for each sample. The Verkko pipeline itself executed the following tools as part of the assembly process: MBG[18] (v1.0.15), GraphAligner[19] (v1.0.17), and MashMap[20] (v3.0.6) for all samples except NA19240, HG00514, HG00732, HG00096 and NA19239. For this last set of samples, the Verkko pipeline tools were necessarily updated to MBG[18] (v1.0.16) and GraphAligner[19] (v1.0.18) to solve a bug originally discovered in the HG00096 assembly process, which is documented in the Verkko GitHub issue #189. The dataset for the sample HG00512 contained one ONT file that was later discovered to be a contaminant (library id 23-lee-007_PCA100115_3F-run15, **Supplementary Table 2**). We examined the issue with the Verkko developers (*personal communication*) and decided not to rerun the assembly because Verkko's process of aligning the ONT reads against an initial HiFi-based assembly graph is inherently robust against such contamination.

Verkko command to produce unphased assemblies:

```
Unset
verkko --hifi ALL-HIFI-READS.fastq --nano ALL-ONT-READS.fastq
--screen human -d WORK-DIR-UNPHASED
```

Verkko command to produce phased assemblies:

```
Unset
verkko --hifi ALL-HIFI-READS.fastq --nano ALL-ONT-READS.fastq
--screen human -d WORK-DIR-PHASED --assembly WORK-DIR-UNPHASED
--paths STRANDSEQ-PHASING-PATHS.gaf
```

The phased assembly output of Verkko consisted of the following assembly units:

1. haplotypes 1 and 2: the main haploid assemblies for each sample
2. unassigned: a set of unphased contigs, i.e., contigs that are not assigned to a specific haplotype.
3. disconnected: a set of usually short assembled sequences that could not be connected to any other component in the assembly graph.
4. rDNA: the set of sequences identified by Verkko as rDNA sequences (see option "--screen")
5. EBV: the set of sequences identified by Verkko as Epstein-Barr viral sequences
6. mito: the set of sequences identified by Verkko as mitochondrial sequences (see option "--screen")

If not explicitly stated otherwise, the unassigned, disconnected, rDNA, EBV and mitochondrial sequences were omitted from any downstream analysis.
We emphasize here that, due to the characteristics of the Strand-seq technology, it is not possible to *a priori* determine the parent of origin for a given assembled haplotype of a sample. Hence, the common denominations haplotype 1 (hap1/H1) and haplotype 2 (hap2/H2) are in principle interchangeable and, in particular, have not been harmonized - in terms of the parent of origin - across samples.

#### Hifiasm hybrid genome assembly

Hifiasm (UL)[21] (v0.19.6) assemblies were generated in three steps. First, a pseudo-phased assembly was generated using both the HiFi and ONT reads:

Unset
```
hifiasm -o {output.prefix} -t {threads} --ul {input.ont_fastq}
{input.hifi_fastq}
```

The resulting assembly graph in GFA format was then forwarded into the previously published Graphasing phasing pipeline[17] as for Verkko above. The resulting phasing paths and sequences were then used to generate the yak (github.com/lh3/yak) databases for phasing the assemblies. With these yak databases, the command was as follows:

Unset
```
hifiasm -o {output.prefix} -t {threads} -1 {input.yak_h1} -2
{input.yak_h2} --ul /dev/null /dev/null
```

This process uses the same error corrected reads as the initial run but applies the phasing information to the resulting assembly graph.

#### Standardized evaluation of genome assemblies

All phased assemblies were subjected to the same evaluation pipeline implemented in a Snakemake[15] (v7.19.1) workflow (**Code Availability**). The input to the evaluation pipeline consisted of FASTA files representing the different assembly units, e.g., haplotypes 1 and 2 or the unphased ("unassigned") sequences for Verkko assemblies (see "Verkko hybrid genome assembly" above), plus the same HiFi and ONT reads that were used for the assembly.

##### Identification of contaminants

We employed NCBI's Foreign Contamination Screening (FCS) tool[22] (v0.4.0) with database release 2023-01-24 to screen all assembly units for remaining sequencing adapters or common contaminants such as mycoplasma. Sequences flagged as contaminants of any type were removed from the assembly sequence files - with the exception of Verkko's "EBV" output - and the cleaned-up assemblies were then further characterized as explained in the following steps.

FCS-GX command line to screen for foreign organism contamination:

```
Unset
python3 fcs.py screen genome --fasta ASSEMBLY.fasta --out-dir OUT_DIR
--gx-db r2023-01-24/all --tax-id 9606
```

FCS-adaptor command line to screen for adaptor and vector contamination:

```
Unset
run_fcsadaptor.sh --fasta-input ASSEMBLY.fasta --output-dir OUT_DIR
--euk --container-engine singularity --image fcs-adaptor.sif
```

##### Assembly contig-to-reference alignment

All assembled contigs were aligned against two human reference assemblies: a variant of GRCh38 excluding the ALT sequences (GRCh38-NoAlt described in Ref. [4]) and the T2T-CHM13v2.0 assembly (T2T-CHM13, UCSC hs1)[23], which includes a complete Y chromosome of the individual HG002/NA24385. Assemblies of female samples were aligned against a T2T-CHM13 reference with N-masked PAR regions of the Y chromosome. All alignments were performed with minimap2[24,25] (v2.26) and filtered to exclude unmapped sequences (flag 1540) with SAMtools[26] (v1.17). The contig-to-reference alignments were used to derive a reference chromosome assignment for the assembled contigs by selecting the chromosome with the highest number of matches to the assembled contig as reported by minimap2. The sex (male/female) for each assembled haplotype of the male samples was determined in the same way.

```
Unset
minimap2 -a -x asm20 --cs --eqx -t THREADS -R READGROUP REF-ASSEMBLY.fasta
INPUT-ASSEMBLY.fasta
```

##### Read-to-assembly alignment

All HiFi and ONT reads used for the assembly were aligned against the complete assembly for the purpose of quality control. All alignments were performed with minimap2[24,25] (v2.26) and not filtered by default. The minimap2 parameter presets "map-hifi" and "map-ont" were used to align the HiFi and the ONT reads, respectively.

Unset
```
minimap2 -a -x READ-PRESET --MD --eqx --cs -t THREADS -R READGROUP
INPUT-ASSEMBLY.fasta ALL-READS.fastq
```

##### Flagging regions of putative assembly collapses with NucFreq

We used an adapted variant of the tool NucFreq[27] (fork NucFreqTwo / branch "split-two-phases" / commit bd080aa) to identify positions where the HiFi read-to-assembly alignments (see section "Read-to-assembly alignment" above, filtered here with flag 3844) showed an elevated count of the second most common base in the alignments for the respective position in the assembly. We applied the following filtering criteria to identify those positions: (1) the second most common base must be observed in at least two alignments at that position; (2) the second most common base must be observed in at least 10% of all alignments at that position. Next, regions with an increased density of such discordant alignment positions were flagged if at least five discordant positions occurred in a window of at most 500 bp. A common interpretation of NucFreq flagged regions is that they represent potential assembly collapses if they coincide with regions that exhibit an elevated read depth. We hence annotated all NucFreq regions with the median HiFi read depth normalized to a percentage value relative to the global median HiFi depth for the respective sample for comparability using custom scripts. In the integrative QC analysis (see section "Integrative analysis of QC annotations" below), all regions flagged by NucFreq were labeled "NUCFRQ".

The adapted version of NucFreq we used (termed "NucFreqTwo") is quasi-identical to the original NucFreq code but separates the computations for identifying the flagged regions from the plotting output of said regions. These minor changes were necessary because the original NucFreq code broke for more fragmented assemblies, i.e., in cases where a lot of flagged regions were plotted automatically. The following command was executed to identify the regions flagged for potential assembly collapses:

Unset
```
NucFreqTwo.py --infile HIFI-READ-TO-ASSEMBLY.bam --flag-discordant-pct 10
--flag-discordant-abs 2 --flag-min-interval 500 --flag-num-hets 5
--min-region-size 500000
# last parameter was introduced to run NucFreq[Two] genome-wide as opposed
# to the original NucFreq which is commonly executed with a list
# of regions (BED file) selected for flagging/plotting. The above filter
# skips over regions smaller than 500000 in the BAM file
```

##### Flagging regions of putative assembly errors with Flagger

We used Flagger[28] (v0.3.3), a read-alignment-based pipeline, to detect putative assembly errors in the diploid assemblies. HiFi reads were aligned to the phased assemblies, i.e., combining haplotype 1 and 2, in a haplotype-aware manner. For improved mapping in repeat regions, Meryl[29] (v1.0) and Winnowmap2[30,31] (v2.03) were used to generate repeat-aware read alignments:

```
Unset
meryl count k=15 output MERYL_DIR ASSEMBLY.fasta
meryl print greater-than distinct=0.9998 MERYL_DIR > REP_K15.txt
winnowmap -W REP_K15.txt -t THREADS -I 10G -Y -ax --MD --cs -L --eqx
ASSEMBLY.fasta HIFI_READS.fastq.gz
```

Flagger was executed using a WDL pipeline. The command line used for execution is as follows:

```
Unset
java -jar cromwell.jar run flagger_end_to_end.wdl --inputs INPUT.json
--metadata-output OUTPUT.json
```

The configuration options in the INPUT.json file were set as follows:

```
Unset
{
  "FlaggerEndToEnd.refSDBed": REF_SD.bed,
  "FlaggerEndToEnd.preprocess.variantCallingMemory": 48,
  "FlaggerEndToEnd.sampleName": SAMPLE_NAME,
  "FlaggerEndToEnd.fai": CONCAT_ASSEMBLY.fasta.gz.fai,
  "FlaggerEndToEnd.stats.threadCount": 4,
  "FlaggerEndToEnd.refBiasedRegionFactorArray": [1.25, 0.75],
  "FlaggerEndToEnd.flagger_alt_removed.isDiploid": true,
  "FlaggerEndToEnd.stats_alt_removed.threadCount": 4,
  "FlaggerEndToEnd.flagger.isDiploid": true,
  "FlaggerEndToEnd.stats_alt_removed.memSize": 4,
  "FlaggerEndToEnd.maxReadDivergence": 0.02,
  "FlaggerEndToEnd.refCntrCtBed": REF_CENTROMERE_TRANSITION.bed,
  "FlaggerEndToEnd.refSexBed": REF_SEX_CHROMS.bed,
  "FlaggerEndToEnd.variantCaller": "dv",
  "FlaggerEndToEnd.stats.memSize": 4,
  "FlaggerEndToEnd.secphaseOptions": "--hifi",
  "FlaggerEndToEnd.project.isAssemblySplit": false,
```

```
  "FlaggerEndToEnd.assemblyFastaGz": CONCAT_ASSEMBLY.fasta.gz,
  "FlaggerEndToEnd.suffix": "",
"FlaggerEndToEnd.refBiasedBlocksBedArray":[REF_HIFI_R1_BIASED.bed,REF_HIFI_R
2_BIASED.bed],
  "FlaggerEndToEnd.refCntrBed": REF_CENTROMERIC_SATELLITES.bed,
  "FlaggerEndToEnd.refBiasedRegionNameArray": ["hifi_biased_high",
"hifi_biased_low"],
  "FlaggerEndToEnd.refName": "hs1",
  "FlaggerEndToEnd.readAlignmentBam": HIFI_READS_TO_ASSEMBLY.bam,
  "FlaggerEndToEnd.hap1ToRefBam": HAP1_ASSEMBLY_TO_REF.bam,
  "FlaggerEndToEnd.hap2ToRefBam": HAP2_ASSEMBLY_TO_REF.bam
}
```

In cases where the primary alignment is deemed inaccurate due to assembly errors, Secphase (github.com/mobinasri/secphase; v0.4.3; integrated in Flagger) identifies the correct haplotype among the secondary alignments, ensuring more accurate read assignment to each haplotype.

For the integrative QC analysis (see section "Integrative analysis of QC annotations" below), all non-haploid Flagger labels were used. The Flagger labels are all prefixed with “FLG” and were shortened as follows: error (-ERR), unknown (-UNK), collapsed (-COL), duplicated (-DUP).

##### Variant-based QV estimation with DeepVariant

We used DeepVariant[32] (v1.6.0) to call short variants in the HiFi read-to-assembly alignments (see section “Read-to-assembly alignment” above, filtered here with flag 3844) that could indicate small assembly errors. The resulting callset was quality-filtered with bcftools[26] (v1.17) and all remaining variants were counted as putative assembly errors:

Unset
```
/opt/deepvariant/bin/run_deepvariant --model_type PACBIO --ref
INPUT-ASSEMBLY.fasta --reads HIFI_READS.bam --num_shards THREADS
--output_vcf DV_CALLS.vcf --noruntime_report --novcf_stats_report
--sample_name SAMPLE

--intermediate_results_dir TMP_RUN
```

Unset
```
bcftools view -f PASS -i 'QUAL>=10 && FORMAT/DP>=5' DV_CALLS.vcf
```

All variant calls remaining after filtering were counted in base pairs as putative assembly errors and a sequence QV estimate was derived as described previously[4] after adjusting the number of error-free base pairs by subtracting the total length of all gaps (stretches of Ns) in the respective sequence. In the integrative QC analysis (see section "Integrative analysis of QC annotations" below), all regions flagged by DeepVariant were labeled as “DEEPVR”.

##### *K*-mer based analysis of assemblies with Merqury

We used Meryl[29] (v1.0) and Merqury[29] (v1.0) to estimate the QV values of the assemblies. The QV values were calculated as phred-scaled scores[33], based on the error rates determined by comparing the *k*-mer databases generated from the sample sequencing datasets with the assembled sequences. Illumina sequencing data, which has high base accuracy, was used to construct the *k*-mer databases.

To generate the *k*-mer database from the sequencing dataset, the following command was executed:

```
Unset
meryl k=21 count ILLUMINA_READS.fastq.gz output READS_DB.meryl
```

To compute the QV value for the entire diploid assembly, as well as the QV values for each haplotype (hap1 and hap2), the following command was executed:

```
Unset
merqury.sh READS_DB.meryl HAP1_ASSEMBLY.fa.gz HAP2_ASSEMBLY.fa.gz
MERQURY_OUTPUT
```

##### Assessing gene completeness of the assemblies with compleasm

We used compleasm[34] (v0.2.5) to assess the gene completeness of the assembled haplotypes in terms of the presence/absence of known single-copy orthologs in the OrthoDB[35] (v10) primate database.

```
Unset
compleasm run --mode busco -L DB_DIR/ -l primates_odb10 --threads THREADS

-o OUT_DIR/ -a INPUT-ASSEMBLY.fasta
```

The results of compleasm includes the information of single-copy genes that exhibit a fragmented alignment to the respective assembly or that show additional copies. That information was incorporated into the integrative QC analysis (see section "Integrative analysis of QC annotations" below) by labeling the respective regions as “BSCFRG” (fragmented) and “BSCDUP” (duplicated), respectively.

##### Identifying complete chromosome assemblies

We labeled chromosomes as T2T by applying the following set of criteria:

1. Telomeres were assembled on both arms of the chromosome.
2. The chromosome spans the respective T2T-CHM13 reference chromosome approximately from end to end.
3. The QV estimate for the assembled sequence is above 50 (see sections “Variant-based QV estimation with DeepVariant” and “K-mer based analysis of assemblies with Merqury” above).

The required input data to check criteria 1 and 2 were computed as follows:

```
Unset
(1-telomere): seqtk telo INPUT-ASSEMBLY.fasta > ASSEMBLY.telo.tsv
```

```
Unset
(2-ref span): mashmap -f one-to-one --pi 99 --seqLength 100000 --dense

-r REF-ASSEMBLY.fasta -q INPUT-ASSEMBLY.fasta --output APPROX-ALIGN.paf
```

The adapted contig-to-reference alignment strategy using MashMap[20] (v3.1.3) resulted from the observation that base-level alignments occasionally could not traverse complex regions, e.g., on chromosome 16. This resulted in fragmented alignments of contiguously assembled chromosomes, which was avoided by resorting to the approximate, i.e., not base-level accurate alignment strategy of mashmap. Mashmap was executed with a stringent setting for reporting alignments only above a threshold of 99% sequence identity to then identify a high-confidence anchor region, i.e., the largest reported alignment. The span of the assembled chromosome relative to the reference was then computed relative to this anchor region. An assembled sequence was considered reference spanning if it covered more than 95% of the reference sequence length. This threshold was lowered to 60% for the Y chromosome due to the recently reported high variation in total length[8], and to 90% for the T2T-CHM13 chromosome 9 to account for the multi-megabase HSat3 duplication in the CHM13 cell line[23].

We note here that for the sample HG00732, the X chromosome assembly in one haplotype was deteriorated to a degree that we labeled it as missing (**Fig. 1e**). Based on the progressive loss of read depth on the X chromosome across the three HiFi sequencing batches for that sample (**Supplementary Fig. 5**), we assume that this is an artifact caused by the cell line.

##### Detection of phasing inconsistencies (Mir/Peter E.)

The Graphasing pipeline[17] assumes that all unitigs in the input assembly graph are specific to a single haplotype. Consequently, haplotype switches present in the input assembly graph will be propagated into the final assembly. We developed the following method to localize these events (referred to as “breakpoints” for brevity) and applied it to the Verkko assemblies.

Strand-seq based (phasing/switch-error) breakpoints were detected using a method similar to that used in the breakpointR R package[36]. The method leverages the haplotype-informative signal inferred from the alignment orientation of the Strand-seq reads, i.e., either in 'Crick' (C) or in 'Watson' (W) direction relative to the assembly sequence. Consequently, a change in the predominant alignment orientation, e.g., from mostly Crick at the beginning of an assembled contig to mostly Watson at its end, indicates a putative phasing error introduced by the assembler that is not supported by the Strand-seq signal.

Our method scans each unitig in windows of varying size by fixing the number of read alignments per window. For each window, the following two-sample binomial proportion test statistic $Z$ is calculated:

$$Z = \frac{p_1 + p_2}{\sqrt{p(1-p)(\frac{1}{W_1+C_1} + \frac{1}{W_2+C_2})}}$$

$$p_1 = \frac{W_1}{W_1+C_1},\ p_2 = \frac{W_2}{W_2+C_2},\ p = \frac{W_1+W_2}{W_1+C_1+W_2+C_2}$$

With $W_1$ and $C_1$ equal to the number of Watson and Crick reads in the back half of the sliding window respectively, and $W_2$ and $C_2$ equal to the number of Watson and Crick reads in the front half of the sliding window respectively. P-values were calculated from the Z statistics, after which a Bonferroni correction is applied at the unitig level to account for the multiple testing problem in this setting. Finally, breakpoints can be identified as locations with p-values lower than a given threshold, with adjacent locations merged into "peaks". The starting and ending coordinate of each peak is reported, giving a high-confidence window in which the breakpoint exists. For breakpoint calling, we used a window size of 300, and a p-value threshold of $10^{-8}$. To filter out small noise peaks, peaks were further filtered to those wider than 75 continuous locations below the p-value threshold. Breakpoints are calculated during the Graphasing pipeline if `calc_breakpoints: True` is set in the configuration file. Besides the Strand-seq reads, the input for Verkko assemblies are the homopolymer compressed assembly graph ("assembly.homopolymer-compressed.gfa") and the assembly HiFi coverage ("assembly.hifi-coverage.csv"), while for hifiasm only the unitigs graph ("*p_utg.gfa") is needed. In both cases, the paths to these files, as well as the "assembler" flag need to be specified in the sample sheet listed in the configuration file.

```
Unset
snakemake -d ../wd/ --configfiles config/config.yaml
// see text above for exec details Verkko vs hifiasm
```

##### Assembly Analysis with Inspector

Inspector[37] (v1.2) was used to evaluate the assembly errors of 65 assemblies, which were recorded as BED files. Assembly completeness was also checked using inspector reference mode aligned against the T2T-CHM13 genome.

```
Unset
inspector.py -c INPUT-ASSEMBLY.fasta -r HiFi_RAWREADS.fastq.gz --ref
REF-Genome.fasta -o OUTPUT_DIRECTORY/ -t8 --datatype hifi

inspector.py -c INPUT-ASSEMBLY.fasta -r ONT_RAWREADS.fastq.gz --ref
REF-Genome.fasta -o OUTPUT_DIRECTORY/ -t8 --datatype nanopore
```

For each sample, 28 metrics were evaluated using a reference-based approach for both Verkko and Hifiasm assemblies, employing either HiFi raw reads or ONT raw reads with the T2T-CHM13 reference (**Supplementary Tables 12,13,21,22**). Basic contig and alignment statistics were calculated, and assembly errors were classified into structural assembly errors (>50 bp) and small-scale errors (≤50 bp). The types of assembly errors considered included base substitution, collapse, expansion, inversion, and haplotype switch. Overall assembly quality (QV) was measured as the ratio of the total bases affected by errors plus the number of inversion errors to the total assembly length. All evaluated parameters are depicted in (**Supplementary Figs. 3,4,6,7**). In the integrative QC analysis (see below), the Inspector flagged regions were labeled “ISPC” plus the suffix “HF” or “ON” to indicate the input read type, i.e., PacBio HiFi or ONT, respectively.

#### Determining the status of previously reported assembly gaps

The evaluation of the status of a set of n=592 previously reported assembly gaps observed in PacBio HiFi-only assemblies[38] was implemented by combining the base-level alignments generated with minimap2 (see section “Assembly contig-to-reference alignment” above) with the approximate alignments generated with mashmap (see section “Identifying complete chromosome assemblies” above) in custom code (see **Code availability**). The output of both aligners was combined to better handle the substantial size variation of the reported assembly gaps (1 bp min, ~30 Mbp max, ~3.6 kbp median). As an example, the largest gaps are located on chromosome Y (q12 region, ~30 Mbp) and on the short arms of the acrocentric chromosomes (up to ~3.8 Mbp on chromosome 13), which are regions that notoriously break alignments due to their repetitive nature. Briefly, a gap was considered covered if any of the two aligners produced a spanning alignment. Next, we annotated the respective spanning contig with Flagger and NucFreq (see respective sections on Flagger and NucFreq above) in analogy to the curation process applied for the centromeres and the chromosome Y analysis (see main text). For this analysis, we decided not to consider any potential errors identified by Flagger or NucFreq only in the actual gap interval, which can be as small as a single base pair, but to limit the overall error rate of the entire assembled sequence for both tools to <1%.

We emphasize here one special case that resulted from our sample and data selection that included NA24385/HG002, which is the sample that was used to produce the first complete Chromosome Y as part of the T2T-CHM13v2 reference[39]. Consequently, our assembly of

NA24385 is extremely biased towards the reference we use as alignment target. We assume that this bias in conjunction with the substantial size variation reported for the chromosome Yq12 region[8] explains why only our NA24385 assembly appears to have closed the ~30 Mbp gap in Yq12 despite other complete Y chromosomes in our dataset (see main text).

### Results

#### Inspector

The Inspector analysis using both PacBio HiFi and ONT reads for the Verkko and the hifiasm assemblies resulted in QV estimates ranging from 43.7 to 56.0, with a median of 48.8 for Verkko. Similarly, Hifiasm assemblies exhibited QV values ranging from 45.8 to 57.1, with a median of 49.6. Verkko assemblies assessed using ONT raw reads had QV values ranging from 31.2 to 34.2, with a median of 23.8, whereas Hifiasm assemblies had QV values ranging from 31.3 to 36.1, with a median of 33.0 **(Supplementary Tables 12,13,21,22**).

Structural errors, defined as the sum of structural assembly errors including collapse, expansion, inversion, and haplotype switch, were also calculated. Verkko assemblies evaluated with HiFi raw reads had structural error counts ranging from 4 to 51, with a median of 51. In comparison, Hifiasm assemblies evaluated with HiFi raw reads had structural error counts ranging from 13 to 127, with a median of 28.

Genome completeness was assessed using the T2T-CHM13 as the reference. Verkko assemblies evaluated with HiFi raw reads demonstrated completeness ranging from 85.73% to 94.15%, with a median of 93.38%. Hifiasm assemblies evaluated with HiFi raw reads exhibited completeness ranging from 88.83% to 96.43%, with a median of 93.77%.

#### Integrative analysis of QC annotations

The merged QC labels (Methods, section “Genome assembly and quality control”) revealed a strong bias for certain tools to flag singleton regions, i.e., regions that were not supported by another tool. This observation was most pronounced for the Inspector analysis using ONT reads, which uniquely flagged >99% of all the flagged regions per Verkko assembly on average (**Supplementary Table 14**). We thus decided to exclude the ONT-based Inspector results, assuming that the inherently higher error-rate in ONT reads makes this data type unsuitable for discovering in particular small-scale errors in the Verkko assemblies (**Supplementary Table 15**).

Next, we computed the overlap and preferential association among all QC labels and regions that are prone to give rise to spurious alignments due to their repetitive nature (telomeres, segmental duplications >98% identity and centromeres), which may affect the performance of all QC tools that rely on read-to-assembly alignments (**Supplementary Table 16**). Limiting to the statistically significant associations (**Supplementary Table 17**; Fisher’s exact test, two-sided, from Python’s scipy.stats; p-values multiple-testing corrected with Benjamini-Yekutieli procedure[40]), we found a depletion of the Flagger-related labels (FLG*) and of NucFreq in centromeres, which is consistent with the manual curation of those regions before accepting them as correctly assembled (see main text). Other enrichments suggest that high-identity segmental duplications may either still pose a challenge for hybrid assemblers or are prone to give rise to false positives as many QC flags exhibit a pronounced positive odds ratio in that region type (BUSCO duplicated genes; Flagger false duplications; Strand-seq breakpoints). Almost all tools show an enrichment in assembled telomeres (DeepVariant, Flagger errors and unknowns, Merqury and NucFreq) except for Inspector, which may suggest that Inspector lacks sufficient sensitivity in these areas of the assemblies.

Finally, we created a merged region set for all QC-related annotations and counted only those regions as true positives, i.e., presumably genuine assembly errors, where two distinct tools flagged overlapping intervals (**Supplementary Table 18**). Regions that were not flagged by at least two tools (in the following: "clean" regions) represent 99.6% (median) of the phased sequence in the Verkko assemblies. The minimal ("size min", **Supplementary Table 18**) region size for both flagged and clean regions (median 1 bp) suggests, though, that a slightly altered merging strategy, i.e., merging based on a distance criterion instead of requiring overlap, would immediately result in a different estimate of the remaining errors in the assemblies. Moreover, we emphasize here that the above estimate refers only to the phased sequences of the assemblies and that sequences such as the rDNA (as identified by Verkko) are excluded from these considerations; the median assembly size (~5.98 Gbp, **Supplementary Table 15**) falls clearly short of a pseudo-diploid T2T-CHM13 reference assembly (~6.02 Gbp). However, restricting the view on the phased sequences is partly a technical necessity because Flagger's inference of the "haploid" label or NucFreq's approach of scanning the read-to-assembly alignments for a locally increased abundance of mismatching nucleotides are not applicable if the underlying assembled sequence has an unclear phasing status.

### Variant discovery and callset development

Contributing authors: Peter A. Audano, Christine R. Beck, Carolyn A. Paisie

#### Methods

##### Genome references

Callsets were constructed against two references, GRCh38 (GRCh38-NoALT) and T2T-CHM13 (T2T-CHM13v2.0)[23]. The GRCh38 reference was previously constructed[4] by obtaining the hg38 reference assembly from the UCSC Genome Browser[41] and removing ALT sequences leaving just the primary assembly (includes chromosome scaffolds as well as unplaced and unlocalized scaffolds). The T2T-CHM13 reference is the hs1 reference assembly obtained from the UCSC Genome Browser and was not modified.

##### Variant discovery

###### PAV

We ran PAV[4], a development version now released under v2.4.1 (**Code Availability**). All default parameters were used specifying only "reference", "aligner", and "assembly_table" in the PAV configuration JSON. Both Verkko and hifiasm assemblies were run with PAV. From

Verkko, we input haplotypes 1 and 2 and the unassigned contigs as three distinct haplotypes. From hifiasm, we input haplotypes 1 and 2. Each PAV callset was replicated using minimap2[24] (v2.26) and LRA[42] (v1.3.7.2) alignments resulting in two independent callsets for each aligner. In total, eight PAV callsets were produced (two assemblies × two aligners × two references).

Genotypes reported in the original PAV VCF files follow the order "h1|h2|un" (haplotype 1, haplotype 2, unassigned).

##### DipCall

We ran DipCall[43] (v0.3) on Verkko and hifiasm assemblies using default parameters for both GRCh38 and T2T-CHM13 references. From both Verkko and hifiasm we input haplotypes 1 and 2.

##### SVIM-asm

We ran SVIM-asm[44] (v1.0.3) on Verkko and hifiasm assemblies using svim-asm haploid using the following parameters: "--tandem_duplications_as_insertions --interspersed_duplications_as_insertions" for both GRCh38 and T2T-CHM references. SVIM-asm uses the same alignments as PAV, which was generated with PAV's CRAM target to create input files for SVIM-asm. From both Verkko and hifiasm we input haplotypes 1 and 2.

##### Read alignments and simple repeat annotations

PacBio HiFi reads were aligned to both GRCh38 and T2T-CHM13 references using pbmm2 (https://github.com/PacificBiosciences/pbmm2; v1.12.0) with "--sort --preset CCS -L 0.1 -c 0". ONT reads were aligned to both GRCh38 and T2T-CHM references using minimap2 (v2.26) with "-ax map-ont --eqx -R".

Tandem repeat annotations were generated by retrieving "Simple Repeats" tracks from UCSC for both references. Records were merged to long non-overlapping regions where any records intersecting or within 200 bp were condensed to a single record. These annotations were input into PBSV and other variant callers using tandem repeat annotations on the reference.

##### PBSV

CCS reads were aligned as described above for both GRCh38 and T2T-CHM references and variants were called using PBSV (https://github.com/PacificBiosciences/pbsv; v2.9.0). The PBSV workflow was executed separately for each sample and for each chromosome. First SV signatures were discovered with "pbsv discover --tandem-repeats <TR.bed> --region <CHROM>" where TR.bed is the simple repeat table described above. Variants were called with "pbsv call" with "--ccs -A 2 -O 2 -P 20 -m 20" for CCS. For each sample, the per-chromosome calls were concatenated, sorted, and compressed with BCFtools[45] (v1.17).

##### Sniffles

Variants were called using Sniffles2[46] (v2.0.7) using the following parameters: "--tandem-repeats TR.bed --minsvlen 5" for both GRCh38 and T2T-CHM13 references for both CCS and ONT reads. "TR.bed" is the simple repeats table described above (PBSV).

##### Delly

Variants were called using Delly[47] (v1.1.6) using default parameters for both GRCh38 and T2T-CHM13 references for both CCS and ONT reads.

##### cuteSV

Variants were called using cuteSV[48] (v2.0.3) using the suggested parameters(https://github.com/tjiangHIT/cuteSV) for both GRCh38 and T2T-CHM13 references for both CCS and ONT reads.

##### DeBreak

Variants were called using DeBreak[49] (v1.0.2) using the following parameters: "debreak --bam merged.sort.bam --outpath debreak_out/ " for both GRCh38 and T2T-CHM13 references for both CCS and ONT reads. Parameter "--rescue_dup" was added for T2T-CHM13.

##### SVIM

Variants were called using SVIM[50] (v2.0.0) using default parameters for "svim alignment" for both GRCh38 and T2T-CHM13 references for both CCS and ONT reads.

##### DeepVariant

For HiFi reads, variants were called using DeepVariant[32] (v1.5.0) using default parameters with the model "--model_type=PACBIO" for both references. For ONT reads, variants were called using PEPPER-Margin-DeepVariant[51] r0.8 using default parameters with the model "--ont_r9_guppy5_sup" for both references.

##### Clair3

Variants were called using Clair3[52] (v1.0.4) using default parameters with "--platform="hifi" --model_path=hifi_sequel2" for CCS reads and with "--platform="ont" --model_path=r941_prom_sup_g5014" for ONT reads.

#### Callset annotation, merging, and QC

##### Merging and support intersects

Variants were compared by SV-Pop[4] (v3.4.4) with parameters tuned for each variant type. These comparisons were used to construct a merged callset as well as compare the merged callset to output from other callers to identify support (i.e., determine if another caller makes the same variant call).

###### Insertions and deletions

A multi-stage merging criterion was used to merge insertions and deletions. All variants are compared by the first criteria and matches are set aside. The remaining variants are compared by the second set of criteria, and so on. Pairs of variants that pass any of the stages are merged together. The staged system allows for the merging process to try different sets of criteria designed to intersect large and small variants and account for breakpoint differences we have previously observed. For example, 50% reciprocal overlap works well for very large SVs, but small breakpoint differences for smaller SVs have a much larger impact on overlap. The fourth stage was added to better intersect SVs in tandemly repeated loci[53].

Parameters: “nr::exact:ro(0.5):szro(0.8,200):szro(0.8,unlimited,3):match(0.8)”

1. exact: Exact match by size and position
    a. Best matches first
2. ro(0.5): 50% reciprocal overlap
    a. For large SVs
3. szro(0.8,200): Within 200 bp and 80% size overlap
    a. For smaller SVs and indels, less sensitive to small breakpoint differences
4. szro(0.8,unlimited,3): Within 3× SV size and 80% size overlap
    a. For tandemly repeated variants that may span large regions

All stages require SV sequences to match by 80% identity using a circular alignment in the case of variants shifted through breakpoint repeats. Only variants of the same type are intersected. Some variant callers did not make SV sequences readily available, so we dropped this sequence match requirement when intersecting with those callsets.

To reduce merging artifacts around the SV size cutoff (50 bp), SVs and indels were intersected as a single callset, and we separated SVs and indels after merging. Support intersects were done the same way with SVs and indels together from the PAV and supporting callers.

##### Inversions

Inversions were intersected by 20% reciprocal overlap to allow for a range of variant sizes that often results from calling inversions from different methods.

##### SNVs

SNVs were intersected only if they exactly matched by both reference position and alternate base.

##### Linking variants across references

A complete callset was produced on both T2T-CHM13 and GRCh38, although it is not possible to identify which variants are in both references from these callsets alone. Using assembly coordinates for each variant call reported by PAV, we translated all variants in both references to assembly coordinates and intersected using the same parameters outlined above for intersecting variants in reference space. This was performed independently for each assembled haplotype.

For each variant in the merged callset, we examined the results of linking in all haplotypes it was identified in. If all haplotypes supporting the call pointed to the same merged sample in the other reference, we linked the merged variants. For example, if a variant in T2T-CHM13 is derived from three haplotypes, then all three haplotype variants should be assigned to the same merged variant in GRCh38 to link the merged variant in T2T-CHM13 to the variant in GRCh38.

#### Annotations for T2T-CHM13

##### Centromere and satellite annotation for T2T-CHM13

We obtained the "CenSat" track from the UCSC browser for the T2T-CHM13 reference genome on 2024-03-13 (https://hgdownload.soe.ucsc.edu/gbdb/hs1/censat/censat.bb). We extracted the type from each record using the "name" column by using the column value up to the first "_" (e.g. "bsat_16_8" becomes "bsat"). Annotation column "REF_CENSAT" was created by intersecting variants with this table and creating a sorted nonredundant list of cen/sat types each variant intersected.

For quality control, filtered regions were obtained by taking all records that were not type "mon" (Monomeric αSat). We created a nonredundant merge with BEDTools[54] (v2.31.1), intersecting the remaining (non-mon) records within 50 bp, resulting in 227 Mbp of filtered reference loci. Variants intersecting these regions by 50 bp were excluded from all callsets. We found excessive numbers of variants from these regions, specifically in large α-satellite HORs and Human Satellite III (HSat3) repeats.

##### Simple repeats

For T2T-CHM13, we obtained the "Simple Repeats" track from the UCSC genome browser for hs1 on 2024-03-13 (https://hgdownload.soe.ucsc.edu/gbdb/hs1/bbi/simpleRepeat.bb). For GRCh38, we obtained the "Simple Repeats" track from the UCSC genome browser for hg38 on 2023-11-27 (http://hgdownload.soe.ucsc.edu/goldenPath/hg38/database/simpleRepeat.txt.gz).

##### Telomere repeats

For T2T-CHM13, we extracted a set of regions from the simple repeats track (see section "Simple repeats" above) where the consensus sequence was any rotation of "TTAGGG" or its reverse complement and excluded all regions that were not within 10 kbp of either end of the chromosome (142 kbp of reference sequence).

##### Modeled centromeres

For GRCh38, we obtained the "Centromeres" track from the UCSC Genome Browser for hg38 on 2024-03-21

(http://hgdownload.soe.ucsc.edu/goldenPath/hg38/database/centromeres.txt.gz).

#### Merged callset quality filters

##### Filters applied to all variant types

Any merged variant that was identified in only unphased assemblies (i.e., no h1 or h2 assemblies) was removed from the T2T-CHM13 and GRCh38 callsets.

For T2T-CHM13, variants were dropped that were composed of 50% or more of excluded centromere and satellite sequence (see section "Annotations for T2T-CHM13" above) or that contained 50% or more of telomeric repeats located at chromosome ends (see section "Annotations for T2T-CHM13" above). For GRCh38, variants that were composed of 50% or more of modeled centromeres were dropped (see section “Annotations for T2T-CHM13” above).

##### Support by callset concordance

To determine if a variant in the merged callset is supported by another caller, we intersect the merge across all samples (unphased diploid callsets) or haplotypes (phased callsets) with the other callset. We chose a strategy that was designed to mitigate placing too much emphasis on a single sample, such as the lead sample for a variant in a merge, or that allowed random matches to arbitrarily support a variant call.

For each caller, we intersect the full merged callset with each individual haplotype (for phased callers) or sample (for unphased callers) using the same parameters as described above to perform the intersect. For each variant, we compare the pattern of samples or haplotypes in the merge with the pattern of samples or haplotypes from the caller with two metrics, Jaccard similarity (sample intersection / union) and Fisher’s exact test (two-sided) using SciPy[55] (v1.11.4) (scipy.stats.fisher_exact). We determine the variant is supported by the caller if the FET *p*-value is 0.01 or less or if the Jaccard similarity is 0.90 or greater, which mostly rescues FET *p*-values in the case that the variant is found in all samples ($p$ = 1.0 if the variant was identified and supported in 100% of the samples or haplotypes).

##### Insertions, deletions, and SNVs

All SNVs, insertions and deletions (including SVs and indels) were accepted into the final callset if they had support from at least three callers. Since SVIM-asm was run using PAV’s alignments, we found that the callsets for all but the largest variants were very similar. For orthogonality, we required that variants be supported by a caller other than PAV and SVIM-asm. Because PAV is able to detect large variants that all other assembly-based callers are unable to detect, we allowed PAV-only calls if they were also supported by PAV with hifiasm assemblies.

##### Inversions

Inversions were accepted into the final callset if they had support from PAV with hifiasm assemblies or were supported by at least three callers.

##### Callset growth estimates

Estimates for callset growth by the addition of a sample was performed by averaging the number of new variants each sample would add in a leave-one-out experiment and counting the number of new variants we would obtain from adding the left-out sample. This was simulated using the existing merge all samples with trio children removed (n = 62 samples) without re-merging. Averages reported in the paper were also computed without trio children and were obtained by running this experiment on all samples (n = 63), all AFR samples (n = 29), and all non-AFR samples (n = 33).

### Results

#### Mendelian inheritance error (MIE)

When HG00514 is excluded from MIE analysis, which has a high HiFi indel error rate compared to other samples (**Supplementary Table 28**), indel MIE falls to 2.1% (-43%), although the indel enrichment was removed during the QC process and is not apparent in per-sample variant calls (**Supplementary Table 29**). When we filter these MIE calls for candidate *de novo* variants and cell line artifacts (Methods), we find 7–40 SV insertions, no inversions, 277–1,690 indels, and 1,027–3,319 SNVs per trio sample.

### Mobile elements

Contributing authors: Parithi Balachandran, Christine R. Beck, Jonathan Crabtree, Scott E. Devine, Miriam K. Konkel, Mark Loftus, Ryan Mills, Weichen Zhou

### Methods

Mobile element insertions (MEIs) were identified within the 130 sample haplotypes assemblies using two separate pipelines and human references (T2T-CHM13 and GRCh38):

One pipeline utilized a custom calling algorithm, L1ME-AID (v1.0.0-beta) (L1 Mediated Annotation and Insertion Detector) (https://github.com/Markloftus/L1ME-AID), that leveraged RepeatMasker[56] (v4.1.6), with the Dfam (v3.8) library[57]**,** annotation of the PAV merged SV insertion callsets (T2T-CHM13 and GRCh38). PAV SV sequences were designated as MEIs by L1ME-AID if they were annotated by Repeatmasker as containing non-LTR retroposon sequence (*Alu*, L1, SVA) or snRNA, had a poly-A tail, and were semi-low divergence (*Alu*: <6%, L1/SVA: <15%) against the Dfam element consensus sequence. Additionally, elements are selected for younger/active subfamilies (e.g., AluY, L1HS, L1PA2, etc.). All candidate sequences were then further selected for minimum mobile element sequence percentage (*Alu*: 70%, L1/SVA: 20% - to allow for transduced sequence).

The second pipeline called MEIs directly from the sample HIFI raw sequence reads with PALMER2 (https://github.com/WeichenZhou/PALMER). PALMER2 represents an enhanced iteration of the PALMER algorithm, engineered to directly call non-reference TE families from aligned, assembled contigs (with PALMER --input ${INPUT_hap.bam} --workdir ${path_to_workingspace} --output ${output_prefix} --ref_ver ${reference_version} --ref_fa ${reference.fa} --type ${TE_type} --chr ${chr} --mode asm). This updated version is crafted to bypass the "masking" phase commonly required in the original workflow[58,59], thereby expediting the process by extracting insertion signals straight from the ultra-long contig sequences. By inheriting the comprehensive feature set of its predecessor, PALMER2 retains the capability of meticulously resolving the structure of TE insertions, characterizing signature retrotransposition hallmarks, and identifying customized non-reference insertion sequences. PALMER2 incorporates a merging utility to streamline the consolidation of individual sample callsets into an integrated, unified callset from diverse genomes.

MEIs identified by either caller were then integrated by comparing the MEI coordinate (chromosome and position), element family (*Alu*, L1, SVA), and sequence composition. If

PALMER2 called an MEI within 1 kbp of a PAV called MEI site, and the two sequences were identified as belonging to the same mobile element family, a pairwise global alignment of their sequences would be performed using Biopython[60] (v1.82; Bio.pairwise2 module). After alignment, the hamming distance (i.e., the proportion of disagreeing components) between the two aligned sequences would be calculated using SciPy (v1.11.1; scipy.spatial.distance.hamming)[55]. If the sequence alignment hamming distance was <0.1, the two MEI calls would be considered identical and merged.

Following merging, the combined MEI callsets underwent additional refinement/quality control by checking if each MEI was a potential deletion from the human reference genome (T2T-CHM13 or GRCh38 depending on callset), duplication, or potential artifact from errors in genome assembly. For each MEI 500 bases of flanking sequence (500 up-stream and 500 down-stream) of the insertion site within the human reference was retrieved and given to BLAT[61] (Standalone BLAT v36x2, -minScore=850) to query for homologous sites within the human reference, and within an ancestral reference (Chimpanzee (NHGRI_mPanTro3-v2.0_pri/GCA_028858775.2), genome. All resulting BLAT hits containing a gap +/-20% difference in size compared to the query MEI insertion call sequence length were further investigated. RepeatMasker[56] (v4.1.6), using the Dfam (v3.8) library[57], was utilized to annotate any elements within the homologous BLAT hits. MEI calls were filtered from the final merged dataset if any secondary sites within the human and/or chimpanzee reference genome were discovered that contain the same mobile element (based on family, orientation, and percent divergence). Finally, any MEIs still remaining and called by a single pipeline were manually curated. An F-test was performed (statistic=1.563603, p-value=0.105) to test for equal variances between sample superpopulation total MEI counts (AFR/non-AFR total MEIs from T2T-CHM13 callset). Since the F-test p-value was ≥0.01 (i.e., variances were equal), and the Shapiro-Wilk test p-values for each population were non-significant (AFR P-value=0.787, statistic=0.97859; non-AFR p-value=0.736, statistic=0.979299; i.e., the data was normally distributed), a Student's t-test was performed to test for a significant difference between African and non-African sample MEI counts (mean=1968 (AFR) and 1423 (non-AFR), statistic=49.7971, p-value=2.6181e-52, degrees of freedom=63.0). All statistical tests were two-tailed.

Callsets were then compared against an orthogonal MEI callset produced by MELT-LRA (see following section, **Supplementary Tables 33,34**).

**Mobile Element Locator Tool-Long Read Assembly (MELT-LRA).** MELT-LRA (https://github.com/Scott-Devine/MELT-LRA), uses PAV calls derived from whole genome assemblies to identify insertions ranging from 50 bp to 20 kbp that are caused by MEIs. Insertions in the 50 bp to 20 kbp range are first identified from the raw PAV calls, and these insertions are then individually compared to consensus sequences for *Alu*, L1, and SVA elements (the three currently active mobile elements in humans) using Smith Waterman alignments between the PAV calls and MEI consensus sequences. Once candidate MEIs are identified from these alignments, they are further annotated using previously developed MELT tools that have been extensively validated with short-read whole genome sequencing and PCR[62,63]. Features annotated include flanking target site duplications (TSDs), poly (A) tails, MEI subfamilies, ORFs (for full-length L1 elements), and genotypes. The MELT-LRA package also includes new visualization tools developed to facilitate this analysis. A total of 12362 MEIs were identified from PAV calls generated through GRCh38 alignments and 12434 MEIs from T2T (CHM13) alignments. This includes: 10352 and 10428 *Alu* insertions from the GRCh38 and T2T alignments, respectively; 1381 and 1367 L1 insertions from the GRCh38 and T2T alignments, respectively; and 629 and 639 SVA insertions from the GRCh38 and T2T alignments, respectively. All but 160 of these MEIs were found in at least one other MEI callset generated with the three independent MEI pipelines (see “Results” below). These 160 were removed from the final callset.

#### Results

Within the resulting merged and refined T2T-CHM13 and GRCh38 MEI callsets there were an average of 13,109 unique MEI calls (T2T-CHM13: n=13216; GRCh38: n=13001) with one or more caller support [average of 11,960 unique MEI calls with the support of both callers (T2T-CHM13: n=11979 (90.6%); GRCh38: n=11941 (91.8%))] (**Supplementary Tables 31,32**). The callsets containing only MEIs with the support of two or more callers showed a comparable amount of element family representation (T2T-CHM13: *Alu*: 9948, L1: 1416, SVA: 615; GRCh38: *Alu*: 9932, L1: 1404, SVA: 605). As expected, we identify a significantly greater number of MEIs within samples of African descent (Mean 1968 AFR, 1423 non-AFR, statistic=49.7971, p-value=2.6181e-52, degrees of freedom=63.0, Student's T-test, two-tailed, T2T-CHM13 reference) (**Supplementary Fig. 10**).

Across the 1426 centromeres analyzed 383 (30.74%) were found to contain at least one MEI in the active alpha-satellite HOR array. In total we identified 806 mobile elements. Though most of these elements (88.95%) were the same element/duplication of a single MEI event across the sample haplotypes (89 unique insertion events; 52 L1, 36 *Alu* element, and 1 SVA insertion). We then selected two unique chromosomes to analyze further (chromosomes 2 and 20). The Chromosome 2 centromere was completely and accurately assembled in 70 haplotypes. Within the HOR array of these 70 centromeres, 56 contained at least one MEI. There were three unique L1 insertions (all L1HS) as well as three unique *Alu* element insertions (two *Alu*Yb8 and one *Alu*Ya5) identified. Chromosome 20 was completely and accurately assembled in 42 haplotypes, 22 which contained at least one MEI in the HOR array. Across these 22 HOR arrays there were 3 unique insertions identified, two L1 insertions (both L1HS) and 1 *Alu* (*Alu*Y) element insertion.

### Inversions

Contributing authors: Hufsah Ashraf, Jan O. Korbel, Tobias Marschall, David Porubsky, Tobias Rausch, Vasiliki Tsapalou

#### Methods

##### Genotyping

We identify in total 276 T2T-CHM13 based and 298 GRCh38 based inversions as part of the filtered PAV callset. We performed cell selection prior to genotyping using ASHLEYS, an automated Strand-seq cell quality assessment approach[64]. Strand-seq based regenotyping was performed using ArbiGent, a Bayesian probability framework based regenotyper that determines inversion genotype likelihoods for inversions and copy number changes, using

Strand-seq reads as an input[13]. ArbiGent is available as a module in MosaiCatcher v2[65] and was executed using default parameters (`--config arbigent=True`).

##### Manual Evaluation

To manually evaluate the False Discovery Rate (FDR) of inversions in the candidate PAV callset, we used NAHRwhals (Höps et al. 2023), a tool for visualizing and detecting complex rearrangements in genome assemblies via dotplot analysis. The analysis utilized the T2T-CHM13 assembly and focused on candidate inversions larger than 5 kbp, a threshold that enables confident characterization and classification through manual dotplot inspection.

#### Results

For T2T-CHM13 based PAV callset, ArbiGent was able to provide an assessment for 252 out of 276 inversion calls, while marking 218 as inversions, 26 as inverted duplications and 2 as potential misorients (**Supplementary Fig. 12, Supplementary Table 37**). To assess the genotyping performance, we tested the genotypes for Hardy-Weinberg equilibrium. 197 inversions belonging to autosomal chromosomes with no missing sample genotype were tested and 169 (~86%) of them were found to be in Hardy-Weinberg equilibrium. For GRCh38 based PAV callset, 230 out of 298 inversions were confidently assessed by ArbiGent, marking 187 as inversions, 21 as inverted duplications, and 16 as potential misorients (**Supplementary Table 38**). 177 inversions belonging to autosomal chromosomes with no missing sample genotype were tested for Hardy-Weinberg equilibrium, 153 (~86%) of which passed. Additionally, we re-genotyped previously developed T2T-CHM13 and GRCh38 based inversion callsets that have been based on a subset of samples[13,66] (**Supplementary Tables 39,40**). ArbiGent genotypes showed a Hardy-Weinberg equilibrium statistic of 89% and 93% for balanced inversions reported in the T2T-CHM13 and GRCh38 callsets, respectively.

By manual inspection, 195 inversions were confirmed as true positives, resulting in an FDR of 12.33%. The most prevalent inversion class was homology-mediated inversions, comprising 67.3% of the identified inversions (132 instances). Simple inversions followed, accounting for approximately 14.8% (29 instances). More complex cases, -such as homology-mediated inversions accompanied by deletions-, constituted ~13% of the true positive inversions (26 instances). Inversions with deletions made up 5% (10 instances). The least abundant category was inversions paired with two deletions, which were observed in only 2 cases (1%). Finally, false positives were detected in 28 instances.

### Segmental duplications

Contributing authors: Mark J.P. Chaisson, Evan E. Eichler, Keon Rabbani, DongAhn Yoo

### Methods

#### Annotations

Segmental duplications (SDs) were identified using SEDEF[67] (v1.1) with the default parameters, on soft-masked genome assemblies, employing TRF[68] v.4.1.0 ("*trf [asm.fa] 2 7 7 80 10 50 2000 -l 30 -h -ngs*"), RepeatMasker[69] v.4.1.5 ("*RepeatMasker -s -e ncbi -xsmall -species human [asm.fa]*"), and Windowmasker[70] (v2.2.22) (*windowmasker -mk_counts -mem 16384 -smem 2048 -infmt fasta -sformat obinary -in [asm.fa] -out [asm.count] &&*

*windowmasker -infmt fasta -ustat [asm.count] -dust T -outfmt interval -in [asm.fa] -out [asm.interval]*); the softmasking was performed using BEDTools-merged regions of the overall repeatmasking results, via seqtk (v1.3) ("*seq -l 50 -M*"). SDs with sequence identity >90%, length > 1 kbp, and satellite content <70% were retained. The putative false SDs were filtered out via intersecting with erroneous regions (lower or higher read depth) from NucFreq or Flagger. Additionally, the highly confident segmental duplication calls validated by fastCN previously generated using T2T-CHM13v1.1 (excluding chrY) genome were used for the consequent comparative analyses (**https://www.biorxiv.org/content/10.1101/2024.06.04.597452v1.full**). SDs in different genomes were compared by mapping them onto T2T-CHM13. Mapping of the assemblies were done by 1) linking SDs within 10 kbp distance, 2) identifying those SD chains that are located in mappable regions; retaining overlap with at least 100 kbp long alignment block, and 3) projecting the chained SDs onto putative homologous SD loci containing at least one 10 kbp unique flank. Accumulation curve of the SDs was generated based on the number of bases of SDs projected onto T2T-CHM13 reference space; note that the data excludes NA19650, NA19434 and NA21487 samples with significantly different SD content. Additionally, homologous SDs by location were further assessed for content using pairwise alignment (minimap 2.26) of the SD bases to identify novel SDs; SDs meeting the criteria of 1) lacking homologous SDs in other T2T-CHM13 SDs by position, 2) having changed sequence content (less than 80% of the sequence conserved), and 3) exhibiting expanded size (at least 2-fold) were considered to be candidates of new SDs. Homozygous and heterozygous genotypes among SDs were determined by comparing SD calls in the haplotype assemblies.

#### Duplicated genes

Protein-coding transcripts (n=20,026) from GENCODE v44 (Liftoff to T2T-CHM13) were mapped to each assembly to identify genes on each assembly; this excludes the NA19650, NA19434 and NA21487 samples unused by the SD analysis. The mapping was performed via minimap2 ("*-cx asm20 -f 5000 -k15 -w10 -p 0.05 -N 200 -m200 -s200 -z10000 --secondary=yes --eqx*"). The mapped genes were further filtered to exclude repeat-only mappings, minimum length of 2 kbp, percent identity of >90% and coverage of >80%. Multi-copy genes were determined by finding the genes greater than count of one and variable copy number genes were defined as those genes containing different copy number in at least one of the genome assemblies.

Gene bodies identified from the transcript alignments ("*minimap2 -x slice -a*") were remapped back to each assembly ("*minimap2 -x asm20 -p 0.2 -N 100 -m 10 -E2,0 -s 10*"), filtering out alignments containing less than 95% of the coding sequences, or that had greater than 5% divergence. Because there are multiple overlapping alignments, partially mapped reads, and different isoforms, the alignments were grouped by exon overlap using Leiden community detection to avoid marking the same region in the same haplotype as multiple separate duplications (**Supplementary Fig. 51**). Final gene annotations were filtered out if they were marked as erroneous by Flagger[28]. To summarize duplications across all samples, we took the union of the isoform community generated for each haplotype, and considered the isoform with the most alignments as the 'representative' isoform. Duplications were defined as separate alignments of the same community. This is implemented in a pipeline called SegDupAnnotation2 (SDA2) (https://github.com/ChaissonLab/SegDupAnnotation2).

#### Comparison of SDA2 annotations versus HPRC annotations of gene duplications

In the draft human pangenome of 94 haplotypes[28], a combination of liftoff and an earlier version of SDA2 annotations were used to detect multicopy genes. We compared duplicated genes detected by SDA2 versus HPRC to annotate discrepancies. Of the 1,115 genes annotated as duplicated in HPRC, we find that 31 are not listed as protein coding in our database, 104 have one of the copies annotated in flagger regions, 412 have different annotations of gene names from multicopy-gene families. The remaining discrepancies are largely due to community detection in SDA2 and different thresholds of completeness of the gene model.

#### Comparison of gene duplications using SEDEF versus SDA2 annotations.

The genes in novel segmental duplications annotated by SEDEF should largely be duplicated genes. We compared the gene lists from SDA2 annotations to those in novel SDs. We considered 900 full genes in novel SDs which reflect a superset of the final list of genes because they are not filtered by fastCN[71,72]. Of these 244 or 27% were excluded by the SDA2 identity filter (<90% identity), 55 or 6% dropped out by length, 85 or 9% dropped out by low gene model alignment (<50% alignment). The remaining 189 genes were not considered by the SDA2 pipeline. Among these, 116 were single-exon genes, 132 were shorter than 5 kbp, and 77 were missing from the original set of genes used to annotate assemblies using SDA2.

### Results

Examining genomes by superpopulation, number of new segmental duplications added per sample (diploid) is highest in AFR of 3.97 Mbp/individual, compared to other populations, 2.81, 3.37, 2.76, 2.57 Mbp/individual in EUR, AMR, EAS, SAS, respectively.

In addition to the segmental duplications that are found novel by their location/synteny, we also examined the sequence content of the syntenic ones, by directly aligning the sequences (**Supplementary Fig. 18a**). Among the segmental duplications that are homologous by position, we find that 14.6 Mbp show less than 80% of sequence identity compared to T2T genome, 4.4 Mbp that are at least two-fold expanded length in the HGSVC3 genomes compared with T2T, and 29.0 Mbp with sequence changed (<80% identity) at the same time expanded (>2-fold), suggesting at least 48 Mbp of the segmental duplications that share positional homology, are not necessarily conserved (**Supplementary Fig. 18a-b**). Quantifying the number of genes that are located in the novel or more variable segmental duplications, we find on average 26 complete overlap of protein-coding genes per chromosome (598 genome-wide) and 5 partial overlap (at least 50% coverage) per chromosome (118 genome-wide).

Compared to the previously known human segmental duplications across 170 haplotypes (Jeong et al. 2024), we observed 23.6 Mbp of new SDs (including 7.3 Mbp that are found in more than one sample, and 16.3 Mbp of that are singleton). This represents complete overlap with 124 protein-coding genes, and partial overlap with 43 protein-coding genes, genome-wide (**Supplementary Fig. 18c**).

We separately used an alignment-based pipeline to annotate full-length multi-copy genes in each assembly using multi-mapped alignments of GENCODE genes (release 44). The analysis was limited to multi-exon genes at least 5 kbp to distinguish from processed pseudogenes.

### STR and VNTR annotation

Contributing authors: Bida Gu, Mark J.P. Chaisson

#### Methods

Annotation of tandem repeats

We used vamos[73] (v1.3.2) and the corresponding motif catalog (v2.1) (https://www.biorxiv.org/content/10.1101/2024.08.07.607105v1) to annotate TRs in 94 HPRC and 130 HGSVC3 haplotype assemblies. A unified diploid VCF file was generated from these annotations, encompassing 112 samples. For each TR locus, distinct alleles were represented in the ”ALTANNO” field as strings of annotated motifs. Diploid genotypes were denoted by allele indices as ordered in the ”ALTANNO” field. Homopolymers were also annotated, but given their limited capacity to reflect complex TR patterns, we excluded them from most analyses to focus on more informative TR structures.

#### Results

TR Allele Diversity

To assess the variability of TRs, we calculated the average number of alleles per annotated haplotype for TR loci with more than 20 annotated haplotypes, using two measurement methods: annotation length and motif composition (**Supplementary Fig. 52a**). As anticipated, locus variability decreased when alleles were measured solely by length compared to motif composition. Despite these differences, the majority of TRs were found to be constant across genomes, suggesting that most TRs are evolutionarily highly conserved. Interestingly, although the HGSVC3 genomes were derived from a more balanced population selection and displayed a similar TR compositional variability to the HPRC genomes, they showed reduced length variability. Our analysis identified 114,817 loci with decreased length variability in the HGSVC3 genomes, with 3,659 (3.2%) located in coding regions. Given that coding TRs account for 5.0% of the total TRs, this suggests that the reduced TR length variability observed in the HGSVC3 genomes is predominantly enriched in noncoding regions. Because the population diversity is high in both datasets, this likely reflects an indication of higher quality assembly. Principal Component Analysis (PCA) and genome wide statistical tests were performed as described previously (https://www.biorxiv.org/content/10.1101/2024.08.07.607105v1), revealing clear clustering of major populations (**Supplementary Fig. 52b**) and 9,297 population informative loci (corrected p-value $< 0.05$; $p < 10^{-8}$) among the 112 samples.

Intersection of tandem repeats in HGSVC2, HGSVC3, and HPRC genomes

Since the motif database was built using HPRC genomes, unique TRs from the HGSVC3 genomes might not be detected within the original framework. To address this gap, we compared the raw outputs from the Tandem Repeats Finder with only centromere regions masked for the HPRC, HGSVC2, and HGSVC3 genomes. As shown in **Supplementary Fig. 52c**, the majority of loci (732,035, 89.0%) were found to be shared among all three datasets, while unique TRs in each dataset largely reflected population-specific differences. Given the relatively higher quality of the HPRC and HGSVC3 assemblies, we conducted a separate comparison between these two datasets. This comparison revealed 30,030 unique TR loci in the HGSVC3 genomes (as shown in **Supplementary Fig. 53**), whereas 21,516 unique TR loci were identified in the HPRC genomes. Importantly, none of these unique loci were found in regions completely unsequenced in the other dataset, indicating comprehensive coverage of both datasets on the CHM13 reference genome. This finding demonstrates the consistency between the two datasets and underscores the robustness of our TR identification approach. As expected, the unique TR loci were primarily concentrated in ribosomal DNA (rDNA) regions, where the CHM13 reference genome is represented by a perfectly repeated model sequence. This outcome suggests that areas with less robust reference data, such as rDNA, tend to contain unique TR loci, emphasizing the need for continued refinement of reference genomes for more accurate TR analysis.

Comparing tandem repeats calls to SV calls

We annotated tandem repeat motif composition for 1,243,719 TR loci on GRCh38, and compared against the integrated SV callset. Among these, 46,314 showed a discrepancy of at least 50 bases between any allele and the reference, indicating a net structural variant. The majority of structural variant calls are accounted for by tandem repeat loci. Of the 176,438 SV loci in the combined variant callset, 113,412 overlap TRs (TR-SV), defined by either insertion variants that are inside of a tandem repeat (N=76,094), or are deletion loci contained entirely within a TR locus (N=37,318). The TR-SV are contained within 31,418 TR loci, indicating clustering of SV calls in TR sequences. Nearly all (31327/31418) TR-SV overlap multiallelic TRs, and roughly 50% (15,944/31,418) have at least 10% of samples with a different length allele.

### Y chromosome

Contributing authors: Peter Ebert, Pille Hallast, Miriam K. Konkel, Mark Loftus, Charles Lee

#### Methods

##### Construction and dating of Y phylogeny

The construction and dating of Y-chromosomal phylogeny combining the 30 males from the current study plus two males (HG01106 and HG01952 from the Human Pangenome Reference Consortium (HPRC) year 1 dataset for which contiguous Yq12 assemblies were used from Ref. [8]) was done as described in detail in Ref. [8]. Please note that the male sample HG03456 appears to have a XYY karyotype as reported in Ref. [5].

All sites were called from the Illumina high-coverage data[5] using the approx. 10.4 Mbp of Y-chromosomal sequence previously defined as accessible to short-read sequencing[74]. BCFtools[45,75] (v1.16) was used with minimum base quality 20, mapping quality 20 and ploidy 1. SNVs within 5 bp of an indel call (SnpGap) and indels were removed, followed by filtering all calls for a minimum read depth of 3 and a requirement of ≥85% of reads covering the position to support the called genotype, followed by removal of sites with ≥5% of missing calls, that is, missing in more than 2 out of 32 samples, were removed using VCFtools[76] (v.0.1.16). After filtering, a total of 10,405,284 sites remained, including 11,129 variant sites.

The Y haplogroups of each sample were predicted as previously described[77] and correspond to the International Society of Genetic Genealogy nomenclature (ISOGG, https://isogg.org, v.15.73). A coalescence-based method implemented in BEAST[78] (v.1.10.4) was used to estimate the ages of internal nodes. RAxML[79] (v.8.2.10) with the GTRGAMMA substitution model was used to construct a starting maximum-likelihood phylogenetic tree for BEAST. Markov chain Monte Carlo samples were based on 200 million iterations, logging every 1,000 iterations, with the first 10% of iterations discarded as burn-in. A constant-sized coalescent tree prior, the GTR substitution model, accounting for site heterogeneity (gamma) and a strict clock with a substitution rate of $0.76 \times 10^{-9}$ (95% CI = $0.67 \times 10^{-9}$ – $0.86 \times 10^{-9}$) single-nucleotide mutations per bp per year was used[80]. A prior with a normal distribution based on the 95% CI of the substitution rate was applied. A summary tree was produced using Tree-Annotator (v.1.10.4) and visualized using the FigTree software (v.1.4.4, **Supplementary Fig. 21**). For figures showing the Y phylogeny subsets of relevant samples were extracted and visualized.

### SVs disrupting genes

Contributing authors: Miriam K. Konkel, Mark Loftus, Gianni V. Martino, Mike E. Talkowski, Xuefang Zhao

#### Methods

##### Annotate the genes overlapped by long-read SVs

To analyze the impact of MEIs on genes, the merged GRCh38 MEI callset was intersected with the findings from Ensembl[81] (release 111) Variant Effect Predictor (VEP)[82] (see section "Transcriptional effect of SVs and functional analysis" below). The MEIs were categorized by insertion location (e.g., protein-coding exons, UTRs of protein-coding transcripts, and noncoding exons) and within each category the number of MEIs present, genes disrupted, and transcripts affected were quantified per category. Using the Ensembl VEP nonsense mediated decay (NMD) plugin [https://github.com/Ensembl/VEP_plugins/blob/release/112/NMD.pm], we predicted which protein-coding transcripts with MEI-induced premature stop codons would escape NMD. The transcripts were further scrutinized by manually comparing the MEI location within the transcript sequence using the UCSC Genome Browser[83]. To ensure that the premature stop codon met one of the four requirements for NMD escape according to the exon-junction complex model[84]. Allele frequencies in the 124 haplotypes (children of trios excluded) were then calculated for the exon-disrupting MEIs. In the event of a "." (indicating misassembly) in the genotyping information, the haplotype was excluded from the calculation.

#### Comparison of Short-read and Long-read SVs that affect genes

Contributing authors: Xuefang Zhao, Michael E. Talkowski

##### Methods:

We benchmarked long-read SVs against SVs generated from matched short-read WGS data. Two SVs are considered to be the same event if they meet the following criteria:

1. Deletions and Duplications that are under 5 kbp should have the same SV types, and share a reciprocal overlap of 10% or more.
2. Deletions and Duplications that are over 5 kbp should have the same SV types, and share a reciprocal overlap of 50% or more.
3. The breakpoints of insertions must be within 100 bp of each other.
4. Insertions can match duplications if the insertion point is within 100 bp of the first breakpoint of the duplication.

##### Results

Out of the 65 samples, 63 had matched short-read SV calls from the 1000 Genomes Project (1kGP). Of the 176,531 SV sites identified from long-read data, 98.9% (N=174,641) are present in at least one of these 63 samples. Consistent with our previous study, long-read sequencing demonstrates superior sensitivity in detecting SVs within highly repetitive regions, such as segmental duplications and simple repeats. Specifically, 82.4% of deletions and 92.1% of insertions in these regions from long-read data do not overlap with short-read SVs. In contrast, a more substantial overlap between the two technologies is observed in the remaining 90% of the genome, which is less repetitive. Here, 11.4% of deletions and 32.1% of deletions and insertions are not overlapped by short-read SVs.

In the 63 samples, an average of 10,918 deletions and 19,541 insertions were observed per individual genome. Consistent with the site-level results, there is a notable overlap between sequencing technologies for SVs in less repetitive regions. Specifically, 89.4% of deletions and 64.5% of insertions in these less repetitive sequences were detected by short-read SV calls in the matched genomes. In contrast, 19.7% of deletions and 9.0% of insertions in the more repetitive regions were detected by short-read SVs.

Coding exons from 818 unique genes were found to overlap with 1,300 unique SV sites across the 63 samples. For long-read SVs that overlap coding sequences and are located outside highly repetitive genomic regions, 84% of the deletions and 63% of the insertions are also detected by short-read SVs. In contrast, only 25.1% of these deletions and 7.3% of these insertions in repetitive regions are overlapped by short-read SVs.

By contrast, from short-read sequencing data we find 206 genes altered in these 65 samples by SV not captured in the long-read assemblies, most of which (83.2%) were disrupted by large CNVs. We leveraged VaPoR to evaluate these SVs by directly comparing raw PacBio alignments against the GRCh38 reference genome. Among the SV sites that are assessable by VaPoR, 27.8% have support from at least one PacBio alignment, indicating potential false negatives in the long-read assembly methods. Interestingly, manual review of these variants in one sample, HG00171 (11 SVs), indicates 1 (8%) was a true deletion between reference assembly gaps that missed by PAV, 2 (17%) are involved in complex SVs that were not fully resolved, 4 (33%) were in contiguously assembled loci with no sign of the deletion, 1 (8%) was present but shifted between the callsets and did not intersect, 2 (17%) were overlapping at the same locus and likely represent two distinct events that may have been lost to merging in the long-read callset, and 1 (8%) was in an uncallable locus.

### Phased transcripts, isoforms, and effects of SVs

Contributing authors: Miriam K. Konkel, Mark Loftus, Gianni V. Martino

#### Methods

##### RNA-seq pre-processing

Initial quality control of the Illumina RNA-seq reads for the 12 samples with corresponding Iso-Seq data was performed using Trimmomatic[85] (v0.39). The trimmed paired-end reads were then aligned to the indexed T2T-CHM13 (NCBI RefSeq GCF_009914755.1-RS_2023_03) using STAR[86] (v2.7.10b). Splice-junction results from all samples were compiled and provided to STAR for a realignment to optimally identify transcript diversity. Cufflinks[87] (v2.2.1) was then utilized to annotate and quantify the RNA-seq expression data

##### Phasing Iso-Seq

For preparation of the 12 PacBio Iso-Seq samples, primers and poly(A) tails were first removed using Lima (v2.1.0) [https://github.com/pacificbiosciences/barcoding/] and isoseq3 refine [https://github.com/PacificBiosciences/IsoSeq], respectively. Iso-Seq reads were then aligned to both haplotype assemblies of the matching sample using pbmm2 (v1.5.0) [https://github.com/PacificBiosciences/pbmm2]. Next, reads were phased through comparison of alignment quality (CIGAR [compact idiosyncratic gapped alignment report] approach) and sequence similarity (*k*-mer approach). Sequence matches and variations (i.e., mismatches and indels) were isometrically rewarded and penalized, respectively, utilizing the CIGAR strings produced by read alignment to each haplotype. The CIGAR-based approach subsequently phased reads to the sample haplotype assembly that produced the alignment with the greater score. The *k*-mer approach compared the 15-mers present in each read to the 15-mers in the corresponding aligned sequence from both haplotypes. The best alignment was identified as the haplotype that shared the most unique *k*-mers with the respective read. Final read-phasing designations utilized an ensemble approach where reads were phased to a sample haplotype if both phasing approaches were in agreement. Disagreement between approaches was settled through determination of which alignment produced a greater quantity of matches. In the absence of measurable differences, according to their scoring premises, an "UnableToBePhased" designation was assigned.

#### Phased Iso-Seq Annotation

Only Iso-Seq with a mapping quality >= 99% were utilized for downstream analyses. These reads were first divided into sample haplotype-specific bins based on their phasing designations (i.e., Hap1, Hap2, or Unphasable). Next, reads within each bin were clustered using isoseq3 (v3.8.2) [https://github.com/PacificBiosciences/IsoSeq] cluster, which removes singleton reads and combines similar reads to form transcripts. These transcripts were aligned to the T2T-CHM13 (NCBI RefSeq GCF_009914755.1-RS_2023_03) reference genome using pbmm2 (v1.5.0) [https://github.com/PacificBiosciences/pbmm2]. Finally, isoforms were generated by running isoseq3 collapse on the mapped transcripts. Isoforms were then annotated and filtered using SQANTI3[88] (v5.1.2) QC and rule-based filter. For each sample, the isoforms formed from the separate haplotype phasing bins were merged, and unique identifiers for each isoform were generated using sample-agnostic characteristics to facilitate cross-sample tracking of similar isoforms.

#### Short- and Long-Read Comparison

We selected all isoforms shared across at least two out of twelve samples present in the PacBio Iso-Seq and/or Illumina RNA-seq, identified all unique protein-coding and noncoding genes represented in either dataset, and compared the gene distribution between both datasets. To achieve this, we filtered the isoforms assembled from Illumina RNA-seq short reads (Cufflinks v2.2.1) [87] retaining only isoforms with non-zero abundance (FPKM). Unique isoforms from the short-read RNA-seq were then identified based upon isoform characteristics (e.g., chromosome, locus, isoform length, gene annotation, and biotype) per sample, as well as across the whole 12 sample set. For Iso-Seq, we used the filtered isoforms generated by the isoseq3 (v3.8.2) pipeline (see **Methods: Phased PacBio Iso-Seq Work-up**), which removed singleton reads. Furthermore, isoforms predicted to undergo nonsense-mediated decay by SQANTI3[88] (v5.1.2) were also excluded. Isoforms were given unique identifiers based upon sample-agnostic features (gene annotation, splice sites, isoform length, etc.) to compare the total unique genes across all 12 samples. Lastly, we compared the average number of unique isoforms per unique gene represented for each sample for both sequencing approaches.

### Results

#### Isoform Phasing

The availability of long-read Iso-Seq data and haplotype-resolved sample-specific genome assemblies allows the detailed investigation of allele-specific transcription. To ensure haplotype-resolved read alignment quality, we required agreement between a CIGAR and a *k*-mer-based approach (see Methods section "Phasing Iso-Seq" above). Congruence between both approaches was observed 98.3% of the time across the 12 samples. On average, 58.44% of the reads could be phased (read phasing range 43.98-71.98%) for a given sample (**Supplementary Fig. 54**), permitting the determination of allele-specific expression (**Supplementary Fig. 55**).

#### Imprinted Loci

Our allele-specific Iso-Seq analysis uncovered ten genes whose isoforms were transcribed from a single haplotype. As imprinted genes show a similar pattern, we reviewed the literature, which confirmed that the ten genes (*ZDBF2*, *NAP1L5*, *FAM50B*, *PEG10*, *IPW*, *MKRN3*, *SNRPN*, *PEG3*, *L3MBTL1*, *LPAR6*) are paternally imprinted in in EBV-transformed B-lymphocyte cell lines[89–91] (**Supplementary Fig. 55**). Furthermore in concordance with the literature, isoforms of the *IPW*, *MKRN3*, and *SNRPN* genes, which are located in the Prader-Willi Syndrome imprinting control region[92], displayed monoallelic expression from the same haplotype within a given sample. The detection of known imprinted genes through phasing allows distinction of paternal from maternal chromosomes. This information can be utilized to determine the origin of a genetic variant in a proband if a parental genome is unavailable. Furthermore, the full agreement of our findings with published imprinted genes provides strong evidence for both our phasing approach and haplotype assembly accuracy.

#### Comparison of Alignments to Phased Assemblies and Reference

While a reference genome serves as a useful baseline for transcriptomic data comparison, the lack of sample specificity can introduce errors into read alignments when the reference genome is discordant from a transcript. Therefore, we quantified the improvement in alignment afforded by mapping long-read Iso-Seq data (12 samples from EBV-transformed B-lymphocytes) to phased sample-specific assemblies versus the T2T-CHM13 reference genome. Out of 26,533,152 total reads, 26,352,868 reads aligned to the personal assemblies and 26,367,272 aligned to the T2T reference. Following comparison of read alignment qualities between the genomes, we observed a set of 105,317 reads that aligned poorly to the assemblies (<50% accuracy) but well to the T2T reference. Interestingly, we found that 93,408 (89.0%) of these poorly aligned reads belonged to just three samples and 99,043 (94.2%) mapped to immunoglobulin genes. These results indicate that most of these poor read alignments can be attributed to somatic rearrangements of immunoglobulin genes in EBV-transformed B-lymphocytes. One possible explanation for the poorly aligned reads being overrepresented in three samples could be that these samples are less clonal than others, leading to increased difficulty for accurate alignments. We further scrutinized the remaining poorly aligned reads (0.02% of total reads) checking for possible misassembly at the read alignment coordinates in the sample-specific assemblies. In total we identified 15 gene loci containing misassembly (4 premature contig ends, 11 incompletely assembled) and 37 loci within 5 kbp of an unassembled region.

To reduce sequencing artifacts, we identified and filtered singleton reads that failed to cluster into transcripts. After removal of singleton reads, we found an equal number of reads (22,432,063) aligned to both the T2T reference and the phased assemblies. However, the phased reads aligned to the assemblies resulted in 21,682,216 reads with at least 99% alignment quality while alignment to T2T produced 21,470,225 reads at the same threshold. This improved alignment of 211,991 reads afforded by phase-specific alignment of Iso-Seq reads to sample-specific haplotype-resolved assemblies marks a 22% reduction in the number of suboptimal alignments (<99% alignment accuracy) resulting from mapping to the T2T-CHM13 reference genome.

#### Comparison of Short- and Long-read RNA-seq

Long-read RNA sequencing, which generates reads that span the full length of transcripts, captures a wider breadth of gene isoforms compared to short-read sequencing methods[93]. However, short-read RNA-seq allows for the quantification of transcript abundances. To better gauge the differences in our expression data, we compared isoform profiles produced from Illumina RNA-seq versus PacBio Iso-Seq across twelve samples. The comparison of total unique genes expressed in at least two samples revealed that short read RNA seq identified a greater total number of genes (29,858 unique protein coding/noncoding genes) compared to long-read Iso-Seq (11,134 unique protein coding/noncoding genes; see **Supplementary Fig. 56**). The lower retrieval of unique genes by Iso-Seq is indicative of incomplete saturation of the transcriptome (mean of 2,211,096 Iso-Seq reads per sample). Additionally, RNA-seq identified on average more unique isoforms (51,413 isoforms) on a per sample basis than the Iso-Seq (33,315 isoforms). However, a greater number of unique isoforms across all twelve samples was identified by Iso-Seq (68,097 unique isoforms) than the short-read RNA-seq (63,028 unique isoforms; **Supplementary Fig. 56**). This disparity can be explained by the Iso-Seq identifying an average of 7,324 unique isoforms per sample while the RNA-seq only identified an average of 523 unique isoforms per sample. Furthermore, the overall average number of unique isoforms per unique gene identified by Iso-Seq across the 12 samples was 6.72, while for short-read RNA-seq identified an overall average of 1.99 unique isoforms per unique genes (**Supplementary Fig. 57**). Despite capturing fewer genes Iso-Seq was able to resolve in excess of three times more isoforms for those genes than the Illumina short-read approach.

### Transcriptional effects of SVs and functional analysis

Contributing authors: Marc Jan Bonder, Mark Gerstein, Matthew Jensen, Yunzhe Jiang, Miriam K. Konkel, Jiaqi Li, Chong Li, Mark Loftus, Gianni V. Martino, Xinghua Shi

### Methods

#### Identification of SV Impacts

To identify SVs with evidence for impacting the transcriptome, we used the Ensembl[81] (release 111) VEP[82] with nonsense-mediated decay (NMD) plugin [https://github.com/Ensembl/VEP_plugins/blob/release/112/NMD.pm] and screened the PAV freeze 4 callset for variants that disrupt gene loci in the merged GRCh38 annotation[94] (NCBI RefSeq GCF_000001405.40-RS_2023_03, Ensembl 111, Gencode v45). Protein-coding genes impacted by putative exon disruptions were then evaluated for evidence of Iso-Seq expression (in >1 sample) across the 12 samples. Isoforms affected by SVs and phased to variant haplotypes were designated as SVs with potential functional impact. Isoforms associated with these SV-containing genes were screened for the presence of unreported splice variants using SQANTI3[88] (v5.1.2). All isoforms of these candidate genes were aligned to GRCh38p14 using pbmm2 (v1.5.0) [https://github.com/PacificBiosciences/pbmm2] and visualized with the Integrative Genomics Viewer (IGV)[95] to identify variant-specific patterns. To determine if the isoforms were previously reported and to identify novel splice products, we compared all isoforms phased to variant haplotypes to known transcripts represented in Refseq[94], GENCODE[96], and CHESS[97] gene annotation databases. MUSCLE[98] (v3.8.425) was used to perform a multiple sequence alignment (MSA) between wild-type and variant haplotype assemblies to identify breakpoints caused by SVs. We utilized Aliview[99] for visualization of the MSA. Variant and wild-type isoforms were also compared by MSA with the assemblies. Select loci were further screened for the presence of transposable elements that may contribute to SV formation or splicing using Repeatmasker[56] (v4.1.6).

#### Enrichment of SVs in genomic regions

Genes and transcripts of the human reference genome GRCh38 assembly were derived from the GENCODE[96] annotation (v45). We further assessed four classes of ENCODE v3 cis-regulatory elements (cCREs), including promoters (<200 bp from gene TSS), proximal (between 200-2000bp from gene TSS) and distal enhancers (>2000 bp from gene TSS), and predicted elements bound by CTCF[100]. Insertions and deletions were intersected with genes and transcripts using BEDTools[54] (v2.30.0) intersect with a minimum overlap of 1 bp. We also extracted exons and introns for long noncoding RNA (lncRNA) and pseudogene transcripts, as well as coding sequences (CDSs), exons, introns, start codons, stop codons, and untranslated regions (UTRs) for protein-coding transcripts. For a given gene, we counted the insertions and deletions within different genomic elements (such as CDSs, exons, and introns) and normalized the counts by the length of the genomic element. We also calculated the proportion of these genomic elements for different transcript types that were affected by insertions or deletions.

We conducted a permutation test to assess the significance of the depletion or enrichment of SVs for these genomic elements. To achieve this, we generated a catalog of shuffled SVs using BEDTools[54] (v2.30.0) shuffle. Given the heterogeneity of chromosomes, we ensured that the shuffled SVs remained on the same chromosome (using the -chrom option) and avoided gaps in the GRCh38 assembly (using the -excl option). Similarly, we intersected the shuffled SVs with genomic elements and counted the number of hits to establish the null distribution. Fold changes were determined as the ratio of observed to permuted values for each transcript type, SV type (insertion and deletion), and genomic element (CDSs, exons, and introns, etc). Empirical p-values were calculated and adjusted for multiple comparisons, with significance reported for adjusted p-values less than 0.05.

#### Differential gene expression and outliers

Expression matrices for the 12 individuals were obtained from short-read RNA-seq data. Differential expression analysis was performed using DESeq2[101] (v1.38.3) for each SV carried by 2-10 individuals out of the 12. SVs were filtered into two categories: 1) direct overlap with an annotated exon (GENCODE v45), or 2) no overlap with an exon but located within 50 kbp of an annotated gene. For genes overlapping or near an SV, the p-values from DESeq2 were corrected using Benjamini-Hochberg adjustment, considering the total number of SV-gene pairs in each category (overlapping with exon or flanking). In these filtered gene-SV pair sets, genes with an adjusted p-value less than 0.05 were reported as differentially expressed.

Outlier expression analysis was performed for SVs carried by one or 11 individuals out of the 12. The gene expression FPKM matrix was used as input, assuming that FPKM follows a log-normal distribution to calculate p-values. The SV-gene pairs were identified and p-values were adjusted using the same methods as in the differential expression analysis. Genes with an adjusted p-value less than 0.05 were considered outliers. In each case, we annotated SVs associated with DE or outlier genes with overlapping cCREs identified using BEDTools[54] (v2.30.0) intersect.

We performed permutation tests to compare the impact of SVs on gene expression with random effects. SVs were randomly redistributed for 1,000 permutations using BEDTools (v2.30.0) shuffle while maintaining chromosome assignment, preventing overlaps, and excluding GRCh38 assembly gaps[54]. To enable a sufficient number of shuffles while maintaining efficiency in differential expression analysis, we developed a method to directly extract DE results from a pre-calculated set that includes all possible individual contrasts within our dataset. This procedure can be applied consistently to both the real SVs and the shuffled sets. We found the number of SV-proximal DE genes in the real SV set was higher than each of the permuted results, suggesting a nonrandom association between SVs and expression change (empirical $p<0.001$). In addition to using a 50 kbp window around the TSSs, we also varied the window size to 5 kbp, 25 kbp, and 100 kbp. We observed that narrower windows yielded a higher proportion of expression-altering SVs, underscoring the importance of sequence integrity near TSSs.

#### Hi-C analysis

We analyzed Hi-C data from 63 individuals to assess the chromatin structures across these genomes. Particularly, we focused on the identification of boundary regions that separate the two adjacent topologically associating domains (TADs) in these individuals. TAD boundaries are recognized for their high conservation across mammalian species and are more evolutionarily constrained than TADs themselves[102,103]. On average, we have identified 21,021 TAD boundaries per genome in these samples (**Supplementary Fig. 58, Supplementary Table 46**).

To identify these TAD boundaries, raw Hi-C sequencing reads were aligned to the GRCh38 reference genome for each individual. Specifically, raw sequencing reads were

preprocessed using *Juicer* software tools (v1.6) with the default BWA-MEM aligner[104,105]. Unaligned reads, including abnormal split reads and duplicate reads, were removed. Additionally, read pairs with a mapping quality (MAPQ) value of less than 30 were filtered out. The resulting high-quality read pairs (Hi-C contacts) were used to construct chromatin contact maps (in *.hic* format, a special format for highly compressed binary files to store contact matrics at various resolutions). These Hi-C contact maps, along with TAD boundary identifications for each sample, were generated using a pipeline following our previous work[8,106]. The files are primarily supported by *Juicer*, *JuicerTools*, and *Juicebox* command line tools for downstream analysis and visualization[107]. Data normalization was done using SCALE matrix balancing[104], which addresses the limitations of KR normalization in unexpectedly not providing coverage for specific regions or chromosomes.
To determine the optimal resolution for calling TAD boundaries, we utilized a script from *Juicer* to calculate the Hi-C map resolution[104], which was directly downloaded from Rao's study[108]. The Hi-C map resolution reflects the finest scale at which local features can be reliably detected, and the average resolutions were 9,137 (bp) for GRCh38 reference mapping in our samples. We employed an algorithm utilizing *Insulation Score* (*IS*) to detect the coordinates and corresponding boundary scores (BS) for each TAD boundary under 10 kbp resolution, using the FAN-C toolkit[109] (v0.9.26b2) with default parameters. TAD boundaries were detected at a 100 kbp window size with a minimum boundary score cut-off value of 0.2, as referenced from the 4DN domain calling protocol[110–112]. The *IS* algorithm defines a sliding window and sums contacts within this window along the Hi-C matrix diagonal. Regions with low insulation scores represent high boundary scores and are further identified as TAD boundaries, while regions with high insulation scores (low boundary scores) are typically found within domains, referred to as TAD regions (**Supplementary Fig. 58**).

##### SV effects on GWAS loci

We investigated the association between variants and human phenotypes or traits by intersecting single nucleotide variants (SNVs), insertions/deletions (INDELs), and structure variants (SVs) with SNPs identified in genome-wide association studies (GWAS). We downloaded the GWAS summary statistics[113] (*gwas_catalog_v1.0.2-associations_e111_r2024-04-16.tsv*), which contains 149,081 autosomal SNPs and we observed 133,695 SNVs present in the GWAS dataset and 10,008 INDELs and 1,547 SVs are at least one bp overlap with any GWAS signal. We used Plink[114] (v1.90b6.10) to examine the linkage disequilibrium (LD) between SNVs, INDELs, and SVs with GWAS SNPs within 1 Mbp window size.

#### Results

##### Expression Impacted by SVs

Structural Variations (SVs) have the propensity to cause drastic alterations to the genomic landscape, with the possibility of impacting transcript diversity through the disruption of coding sequence[115], altering splicing[116], or by rewiring regulatory networks[117]. Using the Ensembl[81] (release 111) VEP[82] to intersect 176,531 SVs (PAV freeze 4 callset) with merged Ensembl GRCh38p14 gene annotations, we identified 1,505 unique SVs with evidence for protein-coding exon disruption within 1,070 unique genes across the 130 haplotype assemblies. In the subset of assemblies with corresponding Iso-Seq data (12 samples; 24 haplotypes) we identified 582 unique SVs that disrupted protein-coding exons within 470 unique genes. After exclusion of 33 immunoglobulin genes, we identified 136 genes with Iso-Seq expression. Of these 136 genes, 85 were found to contain splicing variations absent from the GRCh38 Refseq Annotation[94]. Visual comparison using IGV[95] of isoforms phased to variant and wild-type haplotypes for candidate genes revealed SV-driven transcript diversification.

Following scrutinization of the candidate genes, we found *ZNF718* to best exemplify the impact of SV-driven transcriptome diversification. Associated with immune function[118], *ZNF718* was found to contain a 6,142 bp deletion on chr4:127,125-133,267 affecting 10 haplotypes (8 samples). This deletion resulted in the loss of exons 2 and 3 of the canonical transcript, as well as the majority of an alternate first exon (**Fig. 2e**). We identified a total of 16 unique isoforms in the twelve samples, nine of which were expressed across the ten variant haplotypes, and seven across the 14 wild-type haplotypes. No isoforms were found in common between the variant and wild-type haplotypes. Of the nine unique isoforms expressed by the variant haplotypes, seven were previously unreported in Refseq [94], GENCODE[96], and CHESS[97] databases, while the remaining two were known transcripts that lack the deleted exons. Some unreported isoforms displayed previously uncharacterized splice sites located up and downstream of the SV (isoforms 1, 3, 4; **Fig. 2e**), while others (isoform 2) displayed a loss of all splicing. In both cases, the result was the retention of an intronic sequence that had not been observed previously. In the absence of the deleted portion, the repetitive element landscape appeared to influence the aberrant splicing patterns. Apparent activation of novel splice sites were localized within *Alu* elements. Additionally, all previously unrecognized isoforms appeared to terminate in a transposable element. In contrast, the wild-type haplotypes harbored three known and four unreported isoforms, which all contained some portion of the deleted exons. All isoforms harbored by the wild-type haplotypes contained some portion of the deleted exons, indicating the disruptive impact of this deletion.

#### Enrichment of SVs in Genomic Elements

Permutation tests revealed significant depletion (empirical $p<0.05$, Benjamini-Hochberg correction) of both deletions and duplications across most GENCODE-defined genomic elements, including all biotypes within protein-coding genes and long noncoding RNAs, mirroring our previous findings[62]. Depletions were also observed for SVs within ENCODE-derived cCREs, including promoters and proximal and distal enhancers. However, we found significant enrichments for deletions within unprocessed pseudogenes, in addition to intronic regions of processed pseudogenes (**Extended Data Fig. 3e**), consistent with the SV-mediated mechanisms that lead to pseudogene formation[119].

#### Effects of SVs on Chromatin Conformation

Among the 128 SV-gene pairs (122 unique SVs associated with 98 genes) that exhibit significant differential gene expression changes in the 12 samples with Iso-Seq data, we aimed to identify SVs within these pairs that are also associated with differential insulated regions (DIRs). DIRs represent regions with differential insulation profiles in the presence/absence of SVs across the 12 analyzed samples. Specifically, we first filtered out SVs with missing genotypes in more than 6/12 samples. For each remaining SV, we extracted the 50 kbp upstream and downstream of the annotated TSS position for each paired gene with corresponding insulation scores under 10 kbp resolution. For those insulated regions intersecting with more than one SV, we applied a local multi-test correction. An FDR < 0.05 from the two-sided Wilcoxon rank-sum test was considered as being significant. As a result, we identified 49 SV-gene pairs (29 SVs associated with 24 genes) that were associated with 40 differential insulated regions (DIRs)[8,106] (**Supplementary table 46**).

##### Effects of SVs on GWAS loci

We identified 794,097 SNVs, 138,369 INDELs, and 3,818 SVs in high LD ($R^2 \geq 0.8$) with GWAS SNPs. Specifically, 468,217 SNVs, 77,821 INDELs, and 2,430 SVs were found to be in the highest LD ($R^2 = 1$) with GWAS SNPs (**Supplementary Table 47**, **Supplementary Fig. 59**). Among the SVs in high LD with GWAS SNPs, we observed 3 deletions (chr10-132173822-DEL-112, chr10-132191104-DEL-55, and chr14-67910288-DEL-152) that were identified as SV-differentially expressed gene pairs and linked to DIRs (**Supplementary Table 46**). Notably, chr14-67910288-DEL-152 (associated with the *RAD51B* gene) is in perfect LD with GWAS SNP rs10136790 ($R^2 = 1$), an SNP reported to be associated with the trait of balding measurement[120]. This deletion is also in high LD with rs11158717 (R2 = 0.8536), a locus associated with blonde hair, compared to black and brown hair colors[121]. In addition, chr10-132173822-DEL-112 (associated with the *DPYSL4* gene) is in high LD with GWAS SNPs rs2814162 and rs11146237 ($R^2 = 0.8235$ and 0.9743, respectively). The SNP rs2814162 is associated with vertex-wise sulcal depth for brain measurement, a robust but underexplored measure of localized brain folding that has been previously associated with multiple neurodevelopmental disorders. Sulcal depth is considered a promising neuroimaging marker that could advance people's understanding of cortical morphology[123]. SNP rs11146237, which is also in high LD with the SV chr10-132191104-DEL-55, is associated with body mass index and body weight[124]. Furthermore, we identified four INDELs (chr4-11038-INS-15, chr4-37975-DEL-1, chr4-79676-INS-2, and chr4-82371-DEL-1) that are all in high LD with one GWAS SNP, rs2187874, which is associated with the *ZNF718* gene - a candidate gene that illustrates the impact of SV-driven transcriptome diversification in our study. The SNP rs2187874 has been linked to the response to tumor necrosis factor (TNF) inhibitor in rheumatoid arthritis and joint damage measurement[125].

### Genotyping

Contributing authors: Jana Ebler, Glenn Hickey, Tobias Marschall, Benedict Paten, Timofey Prodanov, Tobias Rausch, Michael C. Zody

#### Methods

##### Building a pangenome graph with Minigraph-Cactus

We built two pangenome graphs with Minigraph-Cactus[126] (v2.7.2). First, we constructed a 132-genome graph composed of the 65 HGSVC samples (130 haplotypes) including

GRCh38 and T2T-CHM13 references. Second, we constructed a 216-genome graph including the 130 genomes along with 42 samples (84 haplotypes) from the HPRC year 1 release [28] (**Supplementary Fig. 27**, Step 1). The input and output data for these two graphs are available at https://s3-us-west-2.amazonaws.com/human-pangenomics/index.html?prefix=pangenomes/scratch/2024_02_26_minigraph_cactus_hgsvc3/ and https://s3-us-west-2.amazonaws.com/human-pangenomics/index.html?prefix=pangenomes/scratch/2024_02_23_minigraph_cactus_hgsvc3_hprc/, respectively.

The command to build the HGSVC graph was:

```
cactus-pangenome ./js1 ./hgsvc3-2024-02-23-mc-chm13.seqfile.local
--outName hgsvc3-2024-02-23-mc-chm13 --outDir
hgsvc3-2024-02-23-mc-chm13 --reference CHM13 GRCh38 --vcf
--vcfReference CHM13 GRCh38 --odgi --xg --chrom-vg clip --chrom-og
--gbz clip full --xg clip full --haplo clip --giraffe clip
--logFile hgsvc3-2024-02-23-mc-chm13.log --batchSystem slurm
--mgCores 64 --mapCores 16 --consCores 64 --indexCores 64
--maxMemory 1.5TB --batchLogsDir batch-logs --coordinationDir
/data/tmp
```

The command to build the HGSVC+HPRC graph was:

```
cactus-pangenome ./js1
./hgsvc3-hprc-2024-02-23-mc-chm13.seqfile.local --outName
hgsvc3-hprc-2024-02-23-mc-chm13 --outDir
hgsvc3-hprc-2024-02-23-mc-chm13 --reference CHM13 GRCh38 --vcf
--vcfReference CHM13 GRCh38 --odgi --xg --chrom-vg clip --chrom-og
--gbz clip full --xg clip full --haplo clip --giraffe clip
--logFile hgsvc3-hprc-2024-02-23-mc-chm13.log --batchSystem slurm
--mgCores 64 --mapCores 16 --consCores 64 --indexCores 64
--maxMemory 1.5TB --batchLogsDir batch-logs --coordinationDir
/data/tmp
```

```
cactus-graphmap-join ./js1 --outName
hgsvc3-hprc-2024-02-23-mc-chm13 --outDir
hgsvc3-hprc-2024-02-23-mc-chm13 --vg
hgsvc3-hprc-2024-02-23-mc-chm13/chrom-alignments/chr*.vg  --hal
hgsvc3-hprc-2024-02-23-mc-chm13/chrom-alignments/chr*.hal
--reference CHM13 GRCh38 --vcf --vcfReference CHM13 GRCh38 --odgi
--chrom-vg clip --chrom-og --gbz clip full --logFile
hgsvc3-hprc-2024-02-23-mc-chm13.join.log --indexCores 64
--maxMemory 1.5TB --batchLogsDir batch-logs --coordinationDir
/data/tmp --batchSystem slurm
```

Note that this second command was used to rerun the indexing step due to running out of memory on our cluster. This issue has been fixed in recent versions of Cactus.

We produced normalized versions of the VCF output of Minigraph-Cactus using vcfwave[76] for realignment, similar to as was done for the HPRC year 1 data and described here:

https://github.com/ComparativeGenomicsToolkit/cactus/blob/hprc-v1.1/doc/mc-pangenomes/hprc-v1.1-mc.md#vcf-postprocessing. The commands run for this were:

```
for i in hgsvc3-2024-02-23-mc-chm13.GRCh38
hgsvc3-hprc-2024-02-23-mc-chm13.GRCh38 hgsvc3-2024-02-23-mc-chm13
hgsvc3-hprc-2024-02-23-mc-c

hm13 ; do docker run -it --rm -v $(pwd)/:/data --user $(id
-u):$(id -g) ghcr.io/pangenome/pggb:202402032147026ffe7f bash -c
"/data/vcf-b

ubwave.sh /data/${i}.raw.vcf.gz
/data/${i}-vcfbub.a100k.wave.vcf.gz" ; done

wget -q
https://s3-us-west-2.amazonaws.com/human-pangenomics/T2T/CHM13/ass
emblies/analysis_set/chm13v2.0.fa.gz

wget -q
http://ftp.1000genomes.ebi.ac.uk/vol1/ftp/data_collections/HGSVC2/
technical/reference/20200513_hg38_NoALT/hg38.no_alt.fa.gz

for i in hgsvc3-2024-02-23-mc-chm13.GRCh38
hgsvc3-hprc-2024-02-23-mc-chm13.GRCh38; do bcftools norm
${i}-vcfbub.a100k.wave.vcf.gz -f hg3

8.no_alt.fa.gz | bcftools sort | bgzip --threads 2 >
${i}-vcfbub.a100k.wave.norm.vcf.gz ; tabix -fp vcf
${i}-vcfbub.a100k.wave.norm.vcf

.gz ; done

for i in hgsvc3-2024-02-23-mc-chm13
hgsvc3-hprc-2024-02-23-mc-chm13 ; do bcftools norm
${i}-vcfbub.a100k.wave.vcf.gz -f chm13v2.0.fa.gz

| bcftools sort | bgzip --threads 2 >
${i}-vcfbub.a100k.wave.norm.vcf.gz ; tabix -fp vcf
${i}-vcfbub.a100k.wave.norm.vcf.gz ; done
```

##### VCF generation and preprocessing

We used the VCF file: https://s3-us-west-2.amazonaws.com/human-pangenomics/pangenomes/scratch/2024_02_23_minigraph_cactus_hgsvc3_hprc/hgsvc3-hprc-2024-02-23-mc-chm13.vcf.gz (Note: VCF prior to the vcfwave-based decomposition) generated by the Minigraph-Cactus pipeline[126] (see Methods section “Buliding a pangenome graph with Minigraph-Cactus” above) from the 216-genome containing all 65 samples as well as 42 samples previously assembled by the HPRC[28] as a basis for genotyping (**Supplementary Fig. 27**, Step 2). This VCF contains a record for each top-level bubble of the graph, as well as haplotype information for all samples contained in the graph. Prior to genotyping, we preprocessed this VCF as follows. We converted the genotypes for male samples on chromosomes X and Y to a homozygous representation by duplicating the haplotype in the VCF that carries the smaller number of "." alleles. Next, we filtered out all records for which at least 20% of the haplotypes carry a missing allele ("."). This resulted in a set of 30,490,169 bubbles. These bubbles are often huge and represent not just a single variant event, but rather accumulate many individual variant alleles across the different samples in the corresponding genomic region. Therefore, we ran a bubble decomposition method that we had developed earlier[28] to find the actual variant alleles nested inside of bubbles. Briefly, the decomposition approach uses the traversals of reference and alternative sequences of the bubbles to determine variation nested inside of the bubble structures [28]. As a result, the pipeline adds annotations to the ID tag of the INFO field of each of our VCF records. For each alternative allele in the VCF, the ID tag contains IDs encoding all nested variant alleles it is composed of, separated by a colon ("bubble vcf", PanGenie input). The decomposition pipeline also outputs a second VCF with a separate, bi-allelic record for each such nested variant ID, defining its reference and alternative sequence[28]. This second VCF can be used later in order to convert genotypes computed for the bubbles to genotypes for all individual variant alleles nested in the graph based on the annotations ("decomposed vcf"). We distinguish the following variant types for the decomposed alleles: SNPs, indels (1-49 bp), SV deletions, SV insertions and other SVs (≥ 50 bp). SV deletions include all alleles for which length(REF) ≥ 50 bp and length(ALT) = 1, SV insertions include all alleles for which length(REF) = 1 and length(ALT) ≥ 50bp. "SV other" contains all other cases where length(REF) ≥ 50 bp or length(ALT)≥50 bp. In total, we obtained 39,313,178 variant alleles after decomposing all 30,490,169 bubbles. In the following, all evaluations are based on the decomposed bubbles.

##### PanGenie input data

PanGenie was run using the VCFs described in the previous section (available here: http://ftp.1000genomes.ebi.ac.uk/vol1/ftp/data_collections/HGSVC3/release/Genotyping_1kGP/20240716_PanGenie-genotypes/:

- MC_hgsvc3-hprc_chm13_filtered_bubbles.vcf (bubble vcf)
- MC_hgsvc3-hprc_chm13_filtered_decomposed.vcf (decomposed vcf)

As a reference genome, we used T2T-CHM13 (v2.0) obtained from: https://s3-us-west-2.amazonaws.com/human-pangenomics/T2T/CHM13/assemblies/analysis_set/chm13v2.0.fa.gz

Illumina reads were obtained from the locations provided below. One FASTA file per sample was generated by concatenating individual FASTA/FASTQ files into a single file.

- 3,202 1kGP samples: http://ftp.sra.ebi.ac.uk/vol1/fastq/
- NA24385: https://ftp-trace.ncbi.nlm.nih.gov/giab/ftp/data/AshkenazimTrio/HG002_NA24385_son/NIST_Illumina_2x250bps/reads/
- NA21309: https://s3-us-west-2.amazonaws.com/human-pangenomics/index.html?prefix=working/HPRC_PLUS/NA21309/raw_data/Illumina/child/
- HG01123, HG02486, HG02559: https://s3-us-west-2.amazonaws.com/human-pangenomics/index.html?prefix=submissions/30E441F3-6820-4BF6-BCF4-E64D56C8D6A4--TRUSEQ/

We provide PanGenie command lines in the sections below.

##### Leave-one-out experiments

We performed "leave-one-out" experiments to evaluate the performance of PanGenie[127] (v3.1.0) on all 107 assembly samples contained in our VCF. This was done by iteratively removing a single sample from the PanGenie input VCF ("bubble vcf") and genotyping it with PanGenie based on the remaining samples using 1000 Genomes high coverage Illumina reads[5]. PanGenie was run with additional parameter -a 108 using the following commands for each panel sample:

```
bcftools view --samples ^<sample>
MC_hgsvc3-hprc_chm13_filtered_bubbles.vcf | bcftools view --min-ac
1 > MC_hgsvc3-hprc_chm13_filtered_bubbles_<sample>.vcf

PanGenie-index -v
MC_hgsvc3-hprc_chm13_filtered_bubbles_<sample>.vcf -r chm13v2.0.fa
-o index -t 24

PanGenie -a 108 -f index -i <(gunzip -c <sample-reads>) -o
pangenie_<sample> -j 24 -t 24 -s <sample>

cat pangenie_<sample>_genotyping.vcf | python3
convert-to-biallelic.py
MC_hgsvc3-hprc_chm13_filtered_decomposed.vcf.gz >
pangenie_<sample>_genotyping_biallelic.vcf
```

The genotypes of the left-out sample were then used as a ground truth for evaluation. Because PanGenie is a re-genotyping method, it can only genotype variants contained in the input panel VCF, that is, it is not able to detect variants unique to the genotyped sample. For this reason we removed all variant alleles (after decomposition) unique to the left-out sample contained in the truth set for evaluation. In order to evaluate the genotype performance, we used the weighted genotype concordance[4,28]. We evaluated the results within the following regions:

- All regions
- Biallelic bubbles: include all bubbles in the graph with at most two branches
- Multiallelic bubbles include all bubbles in the graph with more than two branches
- SegDup regions: UCSC segmental duplications track (CHM13) [128]
- HPRC gap regions: regions with known gaps in HPRC assemblies [38]
- CMRG regions: challenging medically relevant genes [129]

Results for structural variants (≥ 50bp) are shown in **Supplementary Figs. 28-30**. Overall, we observed high genotype concordances across all regions. Performance is best within biallelic regions of the graph and drops in multiallelic regions. This is expected as multiallelic bubbles often contain a large number of different variant alleles. While genotyping performances within CMRG regions are very similar to the overall concordances, they are slightly worse within segmental duplications, and the worst performance was observed within regions where the HPRC assemblies have gaps. This is expected, since our panel contains 42 HPRC samples, and locally incomplete assemblies negatively affect the genotyping.

#### Genotyping the 1kGP cohort with PanGenie

We genotyped all 3,202 samples of the 1000 Genomes cohort based on high coverage Illumina reads[5] using PanGenie[127] (v3.1.0). We first ran PanGenie-index once using the command:

```
PanGenie-index -v MC_hgsvc3-hprc_chm13_filtered_bubbles.vcf -r
chm13v2.0.fa -o index -t 24
```

Then, we ran genotyping on each sample using the command:

```
PanGenie -a 108 -f index -i <(gunzip -c <sample-reads>) -o
pangenie_<sample> -j 24 -t 24 -s <sample>

cat pangenie_<sample>_genotyping.vcf | python3
convert-to-biallelic.py
MC_hgsvc3-hprc_chm13_filtered_decomposed.vcf.gz >
pangenie_<sample>_genotyping_biallelic.vcf
```

We filtered the resulting genotypes using an approach developed previously for the HPRC and HGSVC projects[4,28] that is based on a regression model. We trained the model based on a positive and a negative set which we defined based on the following filters:

- **ac0_fail:** a variant allele was genotyped with a allele frequency of 0.0 across all samples
- **mendel_fail**: the mendelian consistency across trios is less than 85% for a variant allele. Here, we use a strict definition of mendelian consistency which excludes all trios with only 0/0, only 0/1 and only 1/1 genotypes.
- **gq_fail**: less than 50 high quality genotypes were reported for this variant allele
- **self_fail**: genotyping accuracy of a variant allele across the panel samples is less than 90%
- **nonref_fail**: not a single non-0/0 genotype was genotyped correctly across all panel samples

To select for genotypable variants, we trained a machine-learning classifier to assign a pass or fail filter value to each genotyped variant. To train the model, all variant alleles that pass all five filters are included in the positive set. The negative set contains all variant alleles that passed the *ac0_fail* filter but failed at least three of the other filters. In a similar manner as before[4,28], we used Support Vector Regression based on features including the mendelian consistency, allele frequencies and number of alleles transmitted from parent to child samples in order to compute scores between 1 (good) and -1 (bad) for all remaining SV variant alleles. Our final, filtered variant set includes the positive set and all variant alleles with a score ≥ -0.5. We show the number of variants in the unfiltered, positive and filtered sets in **Supplementary Table 48**. Since our focus is on SVs, we applied our machine learning approach only to SVs. Our filtered set contained 89%, 88% and 81% of all deletions, insertions and other SVs that we genotyped. These percentages are much higher than the ones we previously observed when we analyzed HPRC data [28], which is mainly because of several improvements made to the PanGenie software (implemented in versions > 3.0.0, version used here: v3.1.0) in the meantime that led to smaller fractions of variants with allele frequencies of zero across all genotyped samples.

To evaluate the PanGenie genotypes, we compared the allele frequencies of SV alleles that we observed across all assembly samples in our input VCF to the allele frequencies of the same variant alleles after genotyping all 1kGP samples. This was done on all 2,590 unrelated samples. We observed that allele frequencies matched quite well, with pearson correlations of 95.2%, 94.9% and 90.8% for the unfiltered SV deletions, SV insertions and other SV alleles; and 98.1%, 95.7% and 91.9% respectively, after filtering (**Supplementary Fig. 31**). We furthermore compared heterozygosities to allele frequencies and observed a behavior close to what is expected by Hardy-Weinberg equilibrium (**Supplementary Fig. 32**).

We furthermore compared our genotypes to other SV sets, like the PanGenie genotypes we had computed previously for the HPRC data [28] as well as the Illumina-based NYGC SV callset[5]. While our genotypes are based on CHM13, the other sets are both based on GRCh38, making a one-to-one comparison more difficult. In order to account for different SV representations between the three sets, we compared them based on the number of SV sites per sample. We ran truvari collapse[130] using parameters -r 500 -p 0.95 -P 0.95 -s 50 -S 100000 to merge similar SV alleles in our filtered genotyped set, as well as in the HPRC genotypes, since no SV merging was performed for both sets. We show the results in **Supplementary Fig. 34**. To ensure that the improvements we see for the our genotypes were not just due to artifacts arising from the different underlying reference genomes of the compared callsets, we additionally counted the number of heterozygous SVs per sample (**Supplementary Fig. 35**), which revealed the same trends.

The pipeline used for the Leave-one-out experiments and the 1kGP genotyping, as well as instructions to replicate the results can be found here: https://github.com/eblerjana/hgsvc3/tree/main/experiments/genotyping/

#### Comparison of PAV and Minigraph-Cactus based calls

We compared CHM13-based SV calls produced by PAV to the ones obtained from the Minigraph-Cactus graph. Four callsets were compared: the unfiltered PAV calls (“PAV-unfiltered”), the final, filtered PAV calls (“PAV”), the Minigraph-Cactus based panel VCF used as input to PanGenie (“MC-panel”) and the filtered PanGenie genotypes (“PanGenie”). Variants were compared based on reciprocal overlap, breakpoint deviations and variant length deviations. More specifically, two SVs were considered to represent the same event if at least one of the following conditions was true:

- reciprocal overlap > 50%
- $abs(s_1 - s_2) < 200$ and $abs(e_1 - e_2) < 200$ and $0.5 < \frac{l_2}{l_1} < 1.5$

Here, ($s_1$, $e_1$) and ($s_2$, $e_2$) are the start and end coordinates of the MC/PanGenie-filtered and the PAV SVs, respectively (relative to the reference genome), and $l_1$ and $l_2$ the variant lengths. Furthermore, the variant type must be the same, unless one of the SVs has type “OTHER”. The latter ones are allowed to match with any other type.

We performed four pairwise comparisons: MC-panel vs. PAV-unfiltered, MC-panel vs. PAV, PanGenie vs. PAV-unfiltered and PanGenie vs. PAV. Results are shown in **Supplementary Fig. 33.** For overlapping SV calls, counts and variant types were determined relative to the MC / PanGenie-filtered SVs.

##### Unmatched calls

We found 49,868 SVs that are present in the PAV callset, but could not be matched in the MC-panel. Of those, 14,215 SVs were overlapping with calls previously filtered out when creating the MC-panel VCF from the Minigraph-Cactus VCFs (see section "VCF generation and preprocessing" above), while the remaining 35,653 SVs had no overlap with any of these filtered out calls. Furthermore, we found 85,315 SVs in the PanGenie set for which no match was found in the PAV calls. Since PanGenie was run on the MC-panel containing HPRC assemblies as well (in addition the the HGSVC3 samples), some of these variants might be SVs unique to HPRC samples and thus not included in the PAV set, which was produced based on the HGSVC3 samples only. Therefore, we checked the allele frequencies of these variants across HPRC and HGSVC3 samples and found that for 55,095 of 85,315 SVs the allele frequency across HGSVC3 samples was non-zero.

We further analyzed the 35,653 PAV calls that could not be matched in the MC-panel and the 55,095 SVs in the (MC-based) PanGenie filtered set that could not be matched in the PAV calls. We found that many of these calls (i.e., 24,235 of 55,095 and 15,119 of 35,653 SVs) are located within 5mb around the start and end positions of chromosomes. Therefore, we hypothesize that many of these unmatched are cases in which variant representations between the callsets are too different to be picked up by our matching criteria (see section "Comparison of PAV and Minigraph-Cactus based calls" above). Especially in repetitive contexts, coordinates of variant calls might be shifted, or larger records might be reported as multiple smaller calls and thus are missed by our matching script.

##### Generating a 1kGP reference panel with SHAPEIT

We used our final filtered PanGenie genotypes for all 3,202 1kGP samples (see section "Genotyping the 1kGP cohort with PanGenie" above) as a basis for phasing all 3,202 short-read 1kGP samples (**Supplementary Fig. 27**, Step 3). Since this genotyped set is based on variant calls obtained from 107 assembly samples, we included additional rare SNPs and indels from an external short-read based callset for the 3,202 1kGP samples obtained from https://s3-us-west-2.amazonaws.com/human-pangenomics/index.html?prefix=T2T/CHM13/assemblies/variants/1000_Genomes_Project/chm13v2.0/all_samples_3202/. This was done by selecting all variants from this short-read based set that reported genotype 0/0 for all samples overlapping with our 107 assembly samples. In this way, we added 70,174,243 additional variants to our genotyped VCF, resulting in a total of 101,980,762 variants.

In a next step prior to phasing, we set all genotypes with a quality score below 10 to missing (GQ < 10). Then, we ran SHAPEIT5[131] (v5.1.1) using the phase_common command. We provided pedigree information for all trios part of the 3,202 samples, as well as genetic maps for CHM13 obtained from https://github.com/JosephLalli/phasing_T2T/tree/master/t2t_lifted_chrom_maps. Phasing was run separately on each chromosome. For the sex chromosomes, a list of haploid samples (all males) was provided to SHAPEIT.

##### Evaluating phased genotypes

In order to evaluate the quality of our phased genotypes, we compared phasing results to other phased sets on a subset of samples. Other sets included the MC-based input panel

VCF containing the assembly haplotypes (Section "VCF generation and preprocessing"), as well as a phased Illumina-based short variant callset for the 3,202 1kGP samples obtained from https://s3-us-west-2.amazonaws.com/human-pangenomics/T2T/CHM13/assemblies/variants/1000_Genomes_Project/chm13v2.0/. We used WhatsHap's compare function[132] to compute switch error rates (**Supplementary Fig. 36**). To compute switch error rates specific to regions provided in a BED file, we treated each BED interval as a separate phased block to avoid penalizing switch errors that happened outside of the regions of interest. In this way, we computed switch error rates for the following regions:

- CHM13-syntenic: all regions shared between GRCh38 and CHM13. Those were derived from the UCSC CHM13-unique track for CHM13v2.0.
- CHM13-easy: regions outside of other difficult regions such as tandem repeats, homopolymers, difficult to map regions, segmental duplications and high/low GC content. Obtained from: https://ftp-trace.ncbi.nlm.nih.gov/ReferenceSamples/giab/release/genome-stratifications/v3.3/CHM13@all/Union/CHM13_notinalldifficultregions.bed.gz
- CHM13-lowmap-segdup: regions with low mappability and segmental duplication regions. Obtained from: https://ftp-trace.ncbi.nlm.nih.gov/ReferenceSamples/giab/release/genome-stratifications/v3.3/CHM13@all/Union/CHM13_alllowmapandsegdupregions.bed.gz
- CHM13-tandem-repeat: tandem repeat regions and homopolymers, obtained from: https://ftp-trace.ncbi.nlm.nih.gov/ReferenceSamples/giab/release/genome-stratifications/v3.3/CHM13@all/LowComplexity/CHM13_AllTandemRepeatsandHomopolymers_slop5.bed.gz

In line with our expectations, switch error rates were highest for unrelated samples, i.e., samples not part of any of the 1kGP trios, lower for parent samples and lowest for the children (**Supplementary Fig. 36**). Results for CHM13-syntenic regions were almost identical to the overall error rates (orange and blue line, **Supplementary Figure 36**). Overall, the best results were observed for the "CHM13-easy" regions.

The overall switch error rates for unrelated samples (all regions) were between 1.16 and 1.50 when comparing to the MC panel VCF, and 1.16-1.55 when comparing to the Illumina phased calls (**Supplementary Fig. 36**). For parent samples, we observed switch error rates between 0.50-0.79 when comparing to the MC panel and 0.39-1.18 when comparing to the Illumina phased calls. For children, switch error rates were below 0.42 and 0.19 for the MC panel and the Illumina phased calls, respectively. We also performed a multiway comparison of all three sets using WhatsHap's compare function [132]. Briefly, it reports the number of times the different sets agreed or disagreed on the phasing of variant sites. While the phasings of the MC panel and the Illumina calls largely agree, we observed slightly higher rates of disagreements between our and the other two sets (**Supplementary Fig. 37**), especially for the unrelated samples.

#### Personal genome reconstruction

We generated consensus haplotype sequences for all 1kGP samples based on the phasing results. For each haplotype of each sample, we implanted the variants present on that haplotype into the CHM13 reference genome (chromosomes 1-22 and chromosome X). In order to evaluate the quality of the resulting haplotype sequences, we computed QVs and completeness for a subset of 60 haplotypes (30 samples) by comparing *k*-mers of each haplotype to the *k*-mers seen in short-read sequencing data of the respective sample (*k*-mer size used: 21 bp). This was done analogously to how these statistics are computed in Merqury[29]. We furthermore restricted our QV and completeness calculations to certain regions. Given a BED file, we masked all sequences outside of the desired BED intervals with Ns and implanted variants only in the non-masked regions. The Ns were later removed and assembly statistics computed only for the remaining sequence. As a baseline estimate, we additionally computed QV and completeness statistics by flipping sample labels, i.e., by comparing the consensus haplotype for one sample to reads of another sample. In addition to generating consensus haplotypes from our phased genotypes, we also generated consensus haplotypes from the GRCh38-based high-coverage 1kGP set[5], which contains phased genotypes for SNPs, indels and SVs across all 3,202 samples, called from Illumina data and phased using SHAPEIT. The QV and completeness values we observed across all 60 evaluated haplotypes are shown in **Supplementary Fig. 38**.

The pipeline used for phasing and the generation of consensus haplotypes, as well as instructions to replicate the results can be found here: https://github.com/eblerjana/hgsvc3/tree/main/experiments/phasing/

#### Targeted genotyping using Locityper

We used Locityper v0.15.1[133] to genotype 3,202 samples from the 1kGP cohort[5]. Target loci were selected based on 268 challenging medically relevant genes[129] (5 genes not present in the CHM13 reference genome were discarded), extended by genes and pseudogenes from six polymorphic gene families: CFH, CYP2, HLA, KCNK, KIR, MUC and a hyperpolymorphic *LPA* gene. Short genes were extended to both sides to up to 10 kbp, and genes within 1 kbp of each other were combined together. Finally, reference panels were constructed based on the CHM13-based HPRC/HGSVC3 Minigraph-Cactus VCF file, while simultaneously extending locus boundaries if they overlap a pangenomic bubble (variant in the VCF file). Loci that needed to be extended by >300 kbp to one of the sides were discarded, producing a final set of 347 target loci. 61 samples with short read data and HGSVC assemblies were genotyped with two additional reference panels: leave-one-out panel, corresponding to the full panel without the actual sample assemblies; and HPRC-only panel, consisting of the reference genomes and HPRC assemblies. Assembled haplotypes from two individuals (HG00733 and HG02818) were mislabeled as HPRC assemblies, resulting in a bigger HPRC-only panel, which may have diminished the genotyping improvement achieved by switching from the HPRC to the HPRC/HGSVC3 LOO reference panel.

In order to compare actual and predicted haplotypes, we used Locityper align module, which uses largest common *k*-mer subsequence between two haplotypes, and completes the gaps using wavefront alignment algorithm[134]. We then calculated sequence divergence between haplotypes by dividing edit distance by the alignment length, and used Phred-like[33] transformation to convert sequence divergences into QV scores.

Haplotype availability analysis does not require sequencing data and only depends on the assembly sequences. Specifically, for a given sample haplotype we searched for an unrelated haplotype from a limited reference panel that would produce maximal QV score. To evaluate genotyping accuracy, we computed QV scores between predicted and actual haplotypes. All examined loci were diploid, correspondingly, we permuted predicted and actual haplotype pairs to achieve the lowest sequence divergence. Genotypes were marked as low quality and discarded if either weighted distance to other highly probable genotypes was over 30, or there were over 1000 and over 20% unexplained reads [133]. When evaluating trio concordance, we compared predicted child haplotypes against predicted parent haplotypes, and reported corresponding QVs. Similarly to genotyping, we permuted against possible haplotype configurations to achieve the lowest sequence divergence.

### Major Histocompatibility Complex

Contributing authors: Peter A. Audano, Christine R. Beck, Chen-Shan Chin, Alexander T. Dilthey, Peter Ebert, Lisbeth A. Guethlein, Miriam K. Konkel, Heng Li, Mark Loftus, Tobias Marschall, Paul J. Norman, Nicholas R. Pollock, Timofey Prodanov, Stephan Scholz, Arda Söylev, Ying Zhou

#### Methods

##### HLA-DRB and RCCX Visualization

Visualization of DR subregion haplotypes and RCCX modules was performed using a custom Python script, the pyGenomeViz library [https://github.com/moshi4/pyGenomeViz], and the DNA Features Viewer library [https://edinburgh-genome-foundry.github.io/DnaFeaturesViewer/].

##### Visual inspection of *HLA-DRB* haplotypes and solitary exon analysis

To search for structural variation in the *DRB* gene region, HGSVC MHC haplotypes were cut from [start of DRA] to [end of DRB1 + 20 kbp]. The coordinates were obtained using MHC-annotation 0.1 (https://github.com/DiltheyLab/MHC-annotation). Based on their DRB1-allele as determined by Immunoannot, the sequences were grouped into DR groups. Within each group, every sequence was aligned with nucmer 3.1 (-nosimplify -maxmatch) to the same sequence (arbitrarily selected as the sequence with the alphanumerically smallest id) and plotted with a custom gnuplot script based on mummerplots output. Sequences where annotated as follows:

1. Repeat elements were masked with RepeatMasker (v4.1.2)
2. Full DRB genes and pseudogenes were searched for with minimap (v2.26) (--secondary=no -c -x --asm10 -s100) by aligning the sequence from (1) against all DRB alleles from IMGT and the larger DRB9 sequence Z80362.1; results were highlighted and masked for the next step
3. DRB exons were searched for with BLASTN (v2.14.1) by aligning all DRB exons from IMGT to the sequence and filtering for highest matches; results where highlighted and masked for the next step
4. As step 3 but with introns

#### Comparison with 1000 Genomes HLA types

Immunoanot-based HLA types were compared in two-field resolution to the typing published earlier and obtained with PolyPheMe[135] (http://ftp.1000genomes.ebi.ac.uk/vol1/ftp/data_collections/HLA_types/20181129_HLA_types_full_1000_Genomes_Project_panel.txt). 58 out of the 130 haplotypes are not in the PolyPheMe dataset and were excluded.

#### MHC haplotype gene content analysis

In addition to Immuannot, haplotypes were annotated using MHC-annotation (v0.1). Cases of overlapping genes were resolved after inspection by removing superfluous annotations. Reported gene counts for HLA genes and C4 annotation were based on Immuannot.

#### SV analysis

Separately for each HGSVC MHC haplotype, structural variants were called with PAV[4] against 8 completely resolved MHC reference haplotypes[136,137]. To determine which SVs in the HGSVC haplotypes were not present in any of the 8 reference haplotypes, for each HGSVC haplotype, the “query” coordinates (i.e., the coordinates of the calls relative to the analyzed HGSVC haplotype) of the PAV calls were padded with 50 bp on each side and the intersection of SV calls (based on the padded “query” coordinates, across the 8 MHC reference sequences) was computed. Only variants longer than 50 bp were included for further analysis and the smallest variant relative to any of the 8 MHC references was reported. The sequences of the calls so-defined were annotated with RepeatMasker[56]. Variants were grouped by starting position on their closest MHC reference sequence and, in case of insertions, repeat content was averaged.

#### Pangenome analysis for the MHC class II region with PGR-TK

The MHC class II region (MHC delta block) exhibits very high diversity. It is unclear whether standard alignment and variant methodologies (small & structural) can be applied to this region. For instance, when attempting to align assembled contigs to the CHM13 reference, most alignments are fragmented, even with parameters that tolerate more differences. For example, using minimap2 for assembly-to-assembly alignment with up to 5% difference allowed (parameters: -x asm20 -r 10000,100000 -z 100000,100000), it generated 282 fragmented alignments out of the 111 haplotypes (HGSVC batch1 + batch2+ CHM13), and 167 of them had a soft or hard clip greater than 100 kbp on the 5-prime region (**Supplementary Fig. 60**).

We can evaluate the similarity/difference between the haplotypes using a *k*-mer-based, alignment-free method. We count all 25-mers from all haplotype sequences. As shown in the plot below, the peak at 111 indicates conserved *k*-mers across the haplotype sequences. However, there are many significant non-conserved *k*-mers, reflecting the high diversity of the MHC class II region in the human population. There are 60.9% (857,179 out of 1,407,809) 25-mers shown in less than 50% of the haplotype sequences (**Supplementary Fig. 61)**.

In our PGR-TK haplotype cluster analysis, we only select bundles longer than 6 kbp to generate the pangnome graph such that it is more comprehensible. The haplotype sequences in the bundle share a set of sparse minimizers, which can be aligned to each other through a sparse dynamic programming algorithm. At the sequence level, they are typically highly similar to each other. The plot below compares the *k*-mer Jaccard (with $k$ = 25) distribution of different haplotypes of the entire MHC class II region (~CHM13, chr6:32,168,394-32,812,380) to each other and those haplotype sequences within each bundle (**Supplementary Fig. 62)**.

### Results

#### MHC and HLA-DRB

The Major Histocompatibility Complex (MHC), a region of approximately 5 Mbp on the short arm of chromosome 6 (**Fig. 4a**), is the most gene-rich, polymorphic, and disease-associated region of the human genome[138–140]. Due to a combination of extensive structural variation in the HLA-DR and RCCX regions (**Fig. 4a**), long-range linkage disequilibrium[141], and difficulties in obtaining full sequence resolution from short reads[142], however, the majority of MHC disease association signals--for example in the class II region for multiple sclerosis[143]--have not been fine-mapped[144]. Based on the 128 complete and 2 near-complete MHC haplotypes in our assemblies, representing the largest resource of complete MHC haplotypes so far, we characterized the fine-scale structure of genetic variation in the MHC region, providing the basis for more accurate mapping of MHC association signals and addressing key open questions about the relationship between different MHC haplotypes.

Our comparison of our MHC haplotypes with available MHC reference haplotypes[136,137] show a high concordance with respect to the number of annotated[137,145] genes [27-33 Human Leukocyte Antigen (HLA) and 140-146 non-HLA genes and pseudogenes per haplotype] as well as to the overall haplotype repeat content (**Supplementary Table 49**). Our HLA haplotypes were in excellent agreement with classical HLA typing results[135], with concordant results observed for 357 out of 360 compared alleles; **Supplementary Table 50**). A total of 826 HLA allele sequences not present yet at full-length resolution in the IPD-IMGT/HLA reference database were submitted [146] (**Supplementary Table 52**), including 112 sequences from the HLA-DRB locus, expression variants of which are associated with vaccine response[147]. The linkage between the allelic states and presence/absence patterns of HLA genes were in agreement with known MHC class I and MHC class II haplotype structures around *HLA-H*, *-K*, *-T*, *-U*, *HLA-Y* and *-OLI*[148], as well as in the HLA-DR region **(Supplemental Fig. 63)**. While the HGSVC3 MHC sequences did not contain any major novel HLA gene-defined haplotype structures, we did observe a previously unknown copy number variant, deletion of *HLA-DPA2* on one haplotype (HG03807.1), as well as low-frequency gene-level structural variants, such as deletion of *MICA* on one haplotype[149], a gene associated with histocompatibility[150]. A total of 170 SVs not present in previously available reference haplotypes[136,137] were detected (**Supplementary Table 53)**; the majority of these (64%) had a repeat element content ≥70%, and 30 of the remaining ones were larger than 1 kbp. Taken together, while not indicating any major novel haplotype structures, these results further confirm the high accuracy of the HGSVC3 assemblies in a region as complex as the MHC and inform fine-mapping of HLA-related association signals. Additionally, we used Locityper[133] to genotype 61 of our samples using Illumina datasets across 19 protein coding genes and 14 pseudogenes from the MHC locus and compared predicted gene alleles against assembly-based gene annotation, constructed using Immuannot[145]. Across all 33 loci, Locityper fully predicted gene alleles in 81.0% cases using a limited HPRC-only reference panel. Adding HGSVC3 assemblies to the reference panel allowed Locityper to accurately predict gene alleles in 86.3% (leave-one-out panel) and 97.1% cases (full panel). When evaluating predicted protein product, Locityper achieved 88.3%, 90.6% and 97.8% accuracy with HPRC-only, leave-one-out and full reference panels, respectively.

We next carried out in-depth analyses at the nucleotide level of the two major structurally variable regions of the MHC, the HLA-DR and RCCX (**Figure 4a**). We classified the HLA-DR haplotypes of the HGSVC3 assemblies using the established DR group system, which groups HLA-DR haplotypes according to their HLA-DRB1 allele and gene/pseudogene content (**Figure 4b, Supplementary Table 54, Supplementary Fig. 64**). Our MHC assemblies contained representatives of all known DR groups, with counts ranging from 9 (DR1) to 25 (for each of DR5 and DR6); no haplotype configurations incompatible with the established DR group system were found. Based on visual inspection of pairwise alignments, with the exceptions of two small (~3.7 and 4.4 kbp) structural variants that contained solitary HLA-DRB exon sequences (see paragraph "Solitary HLA-DRB exon sequences […]" below), no large SVs distinguishing between DR haplotypes within the same DR group were identified. DR5, DR8 and DR9 were absent from previous analyses of fully resolved MHC haplotypes[136,137]; in addition, the relationship between the two contracted DR haplotypes DR1 and DR8, which carry only one functional HLA-DRB gene (DRB1; **Fig. 4a**), and other HLA-DR haplotypes has not been investigated at the nucleotide level.

For DR5 and DR9, pairwise sequence alignments (**Supplementary Fig. 41**) showed that these haplotype share their large-scale structure with DR3/DR7 and DR4/9, respectively, the other haplotype groups with the same DRB3- and DRB5-carrying gene configuration (**Fig. 4c**). For DR8, pairwise alignments suggested the highest degree of relatedness to either DR1 or DR3/5/6 (**Supplementary Fig. 41**). Leveraging the fully resolved nature of our MHC assemblies, we constructed high-resolution maps of repeat elements and gene/pseudogene exons within the assembled DR haplotypes (**Supplementary Fig. 39; Methods**). Shared repetitive element architecture encompassing the *HLA-DRB1* gene within DR8 to those found flanking the *HLA-DRB3* and *HLA-DRB1* genes in the DR3/5/6 haplotypes (**Fig. 4c**) indicated that, consistent with earlier analyses of DR8 based on only HLA allele sequences[151,152], a sequence contraction event between the *HLA-DRB1* and *HLA-DRB3* genes of a DR3/5/6-class haplotype may have given rise to the DR8 haplotype, which could be fine-mapped to lie within the proximity of exon 4 in HLA-DRB1*08 (**Supplementary Fig. 40**). The putative breakpoint showed a short length (<150 bp) of homology between *HLA-DRB1* and *HLA-DRB3* on DR3/5/6 spanning exon 4 and the surrounding downstream sequence (intron 3) in some DR8 haplotypes suggesting that homology-mediated double strand DNA break repair may be a more likely explanation than nonallelic homologous recombination (NAHR). For DR1, a haplotype structure that had previously been suggested to have arisen from a contraction event within an ancestral *HLA-DRB5* carrying DR2-like haplotype[153], high-resolution repeat mapping suggested that DR1 is most likely derived from a recombination event between DR2 and DR4/7/9. DR1 can be split into two segments (**Fig. 4d**), the 5′ segment of which exhibits the highest degree of repeat architecture similarity to the 5′ region of the *HLA-DRB4*-carrying haplotype groups (DR4/7/9), and the 3′ segment of which shows the highest degree of similarity to a region found in DR2. Breakpoint analysis identified homologous LINE/L1 elements at the putative breakpoints in DR1, DR/4/7/9 (L1 located between *HLA-DRB9* and *HLA-DRB4* within DR4/7/9), and DR2 (L1 located between *HLA-DRB5* and *HLA-DRB6*) (**Supplementary Fig. 42**).

Solitary HLA-DRB exon sequences have been hypothesized to represent remnants of ancestral nonhomologous recombination events, possibly mediated by the presence of LINE1 and Alu in HLA-DRB intron 1 sequences[154,155]; in previous analyses of incompletely resolved MHC haplotypes, they were found in the 3′ region of *HLA-DRB9*[154]. Based on the set of completely resolved HGSVC haplotypes, we created a comprehensive catalog of the locations of HLA-DRB intron and exon sequences across DR groups (**Fig. 4b**). For the region around *HLA-DRB9*, we refined previously available estimates of the copy number of solitary intron and exon sequences, e.g., observing a copy number of 3 instead of 1 for solitary HLA-DRB exon 1 sequences in the *HLA-DRB9* region of DR1. Solitary exon 1 (DR1 group haplotypes) and intron 1 (for all DR groups but DR8) sequences were also found approximately 10 kbp 3′ of *HLA-DRB1*, i.e., at a locus not previously associated with the presence of solitary intron or exons (**Fig. 4b**). Furthermore, the presence of solitary exon 1 sequence in the *HLA-DRB1* region of DR1 and in the DRB9 region of DR3, DR5, DR6, and DR8, was variable even within haplotypes of the same DR group (**Fig. 4b**); the underlying two structural variants (corresponding to the solitary exon-polymorphism in the DRB1 and DRB9 regions, respectively) were identified during the structural analysis of MHC haplotypes (see paragraph “We next carried out in-depth analyses [...]” above) and carried HLA-DRB intron 1 and exon 1 sequences as well as repeat elements (**Supplementary Fig. 65**).

Last, we asked whether the established DR group nomenclature could be recapitulated using unbiased and data-driven analysis. To this end, we analyzed a subset of the HGSVC3 MHC haplotypes (n=55) as well as T2T-CHM13[23] using PGR-TK, a pangenomic multiscale analysis method[156] (**Fig. 4e**). We examined conserved blocks in 111 haplotypes (110 HGSVC haplotypes + CHM13) greater than 6 kbp, identified by PGR-TK. Multiscale hierarchical clustering of the haplotypes perfectly recovered the traditional DR group system in the region around *HLA-DRB1* (**Fig. 4e**); at the same time, the identified additional diversified subgroups could serve as the basis for a more fine-grained future classification of HLA-DR haplotypes or be used in the context of genome-wide association studies.

#### RCCX

The RCCX region in the class III region of the MHC is characterized by a modular pattern of copy number variation. Each module encodes one copy of the Complement component 4 (*C4*) gene (“C”) as well as one functional or nonfunctional/degenerate copy of Serine/threonine kinase 19 (“R”), Steroid 21-hydroxylase (“C”), and Tenascin XB protein (“X”), referred to as *STK19*, *CYP21A2*, and *TNXB* (functional forms) and *STK19B*, *CYP21A1P*, and *TNXA* (pseudogene / degenerate forms), respectively.

RCCX varies in module copy number, commonly ranging from 1-3 modules per haplotype, although rare four module haplotypes have been observed previously[157,158]. Duplications and deletions of RCCX modules as discrete genetic units are commonly observed as the region is prone to NAHR[158–160]. Typically, when the RCCX is present as a mono-module, all four genes within the unit are functional, barring any mutation/recombination. However, when the RCCX contains multiple units (bi/tri-module), *C4* is the sole gene that retains its functional capacity across all discrete RCCX modules, while the additional copies of the other three genes (*STK19*, *CYP21A2*, and *TNXB*) often are present as nonfunctional pseudogenes (*STK19B*, *CYP21A1P*, and *TNXA*)

The *C4* gene is composed of 41 exons that together encode for an inactive 1744 amino acid long gylcoprotein. This inactive form contains three polypeptide chains (alpha, beta, and gamma) and becomes activated through enzymatic cleavage. The *C4* gene exists in four allelic variants, distinguished by four amino acid residues in exon 26 (positions p1120-1125, *C4* alpha chain), referred to as “A” (PCPVLD) and “B” (LSPVIH) variants of *C4* [161,162] as well as by the presence or absence of a ~6.36 kbp human endogenous retrovirus (HERV) insertion within intron 9, referred to as “short” (S) and “long” (L) versions of *C4*.

For RCCX, we carried out a comprehensive investigation of the region’s modular structure of genetic variation (**Supplementary Fig. 43**). RCCX haplotypes consist of 1 - 4 RCCX modules[157,163,164], each module typically encoding one functional variant of the *C4* gene (*C4AS*, *C4AL*, *C4BS*, *C4BL*) and three additional genes in their functional (*STK19*, CYP21A2, and *TNXB*) or pseudogenic (*STK19B*, *CYP21A1P*, and *TNXA*) forms (**Supplementary Fig. 43**); determining the phase and relative order of variants in the region from short-read sequencing or other molecular typing techniques[157,165] is challenging due to extensive sequence homologies. Consistent with prior reports[159,166], RCCX bi-modules were, at 74.6% (n=97), observed most frequently in the HGSVC3 MHC haplotypes, followed by 13.1% (n=17) for mono-modules and 12.3% (n=16) for tri-modules (**Supplemental Fig. 43**). We were able to detect and completely resolve rare RCCX haplotype configurations and characterize likely gene conversion events, which play an important role in RCCX evolution[160,167,168]. For example, we observed two sample haplotypes with tri-modular RCCX structures containing two functional *CYP21A2* copies, likely generated by gene conversion turning the *CYP21A1P* pseudogene into its functional counterpart; one mono-modular and one bi-modular haplotype with no functional *CYP21A2* genes; and one tri-modular haplotype exhibiting the unique configuration of *C4B* preceding *C4A* and carrying two copies of *CYP21A2*, one of which was nonfunctional (**Fig. 4d**). The latter haplotype was likely generated by the introduction of one nonsense mutation and two gene conversion events, turning *CYP21A1P* into *CYP21A2* and *C4A* into a *C4B* that unusually encoded the Rodgers (Rg) blood group epitope. An in-depth analysis of *C4* copy numbers (**Supplementary Fig. 44**) and allelic states (Methods) revealed seven novel amino acid variants, four residing within the beta-chain (283R->C, 530S->L, 566A->G, 614K->Q), and three within the gamma-chain (1477K->N, 1722R->C, 1724R->W) (**Supplementary Fig. 45)**. In addition to providing a more complete catalog of allelic variation, our results thus provide the basis for assessing the differential effects of allelic, copy number and structural variation in RCCX-associated autoimmune phenotypes[169].

#### C4 copy number analysis

Of a total of 259 *C4* copies, 53.3% (n=138) were identified as *C4A* and 46.7% (n=121) as *C4B*. Most individuals (56.9%) harbored four *C4* gene copies, ranging from a minimum of two *C4* copies (15.4%) per individual to a maximum of six *C4* copies (3.1%); *C4A* gene copies ranged from 1-3 copies and *C4B* from 0-4 copies per genome. Most genomes contained at least two *C4A* (72.3%) copies, and/or two copies of C4B (61.5%).

Most (66.4%, n=172) *C4* copies were seen in the long form, *C4L*, and ranged from 1-5 copies within a genome; while the short form *C4S* constituted 33.7% (n=87) of all *C4* copies and ranged from 0-3 copies within a genome. Interestingly, an absence of the short *C4* form, *C4S*, was more common in non-African samples as we identified only one African sample (HG02769) containing zero *C4S*, consistent with[165].

##### Manual curation of HG00514.hap2

*STK19* and *TNXB* were found to be present and functional within all assemblies except HG00514.hap2. This sample haplotype contained a nonsense mutation within *TNXB* caused by an indel. Analysis of the parents showed functional *TNXB* gene copies in all four assembled haplotypes. Further inspection of the sequence revealed that the indel occurred within a homopolymeric stretch, a region where long-read PacBio sequencing may have a reduce accuracy[170–173]. The combination of indel in homopolymeric sequence combined with intact genes in both parents suggests that this variant may be more likely the result of a sequencing artifact; *TNXB* was therefore treated as functional in this haplotype.

### Complex structural polymorphisms

Contributing authors: Peter A. Audano, Christine R. Beck, Evan E. Eichler, Miriam K. Konkel, Charles Lee, Mark Loftus, David Porubsky, Feyza Yilmaz

#### Methods

##### Complex Variant Discovery

Complex variants (CSVs) were identified with a development version of PAV[174] (**Methods**). Briefly, the method uses assemblies trimmed in query space, identifies candidate variant anchor pairs, and scores variants between each candidate anchor pair. A graph is constructed where nodes are scored alignment records (positive values) and edges are scored variant calls (negative values) between alignment records. The score from root to a node is the sum of all alignment record scores and variant scores along the path. Because PAV traces through increasing assembly coordinates without overlaps due to alignment trimming, the graph is a directed acyclic graph (DAG), and the optimal path through the graph can be found in $O(N + E)$ time with the Bellman-Ford algorithm[175] where N is the number of nodes and E is the number of edges. Variants on the optimal path are accepted into the callset. For each variant call, PAV computes the reference context (i.e., DEL, DUP, INV, etc) and the query context (template switches, templated insertions, and untemplated insertions), which are summarized in the manuscript.

Similar to the main callset, CSVs intersecting centromeric repeats were eliminated. Complex variants were merged into a nonredundant callset with SV-Pop by 50% reciprocal-overlap and 80% sequence identity (SV-Pop merge parameter “nr::ro(0.5):match(0.8)”).

##### SMN analysis

We start by extracting the FASTA sequence across all phased genome assemblies from this study as well as from previously published studies (HPRC and T2T-primate). For this we aligned each assembly to the reference genome T2T-CHM13 using minimap2[24] (v2.24). Then we use rustybam (v0.1.33, 10.5281/zenodo.8106233) ‘liftover’ function to subset alignments in PAF format to a desired region (chr5:70300000-72100000). Then we used query coordinates in each subsetted PAF file to extract the query FASTA sequence from each assembly. Next, in order to evaluate the copy number of each multicopy gene *(SMN1/2, SERF1A/B* and *GTF2H2/C*), we extract the sequence of each gene from

T2T-CHM13 based on the available gene annotation (https://s3-us-west-2.amazonaws.com/human-pangenomics/index.html?prefix=T2T/CHM13/assemblies/annotation/chm13v2.0_RefSeq_Liftoff_v5.1.gff3.gz). Then, we align the full gene sequence as well as exonic sequence only to each of the extracted FASTA files. Next we on each FASTA we run DupMasker[176] (v4.1.2-p1) annotation to define ancestral duplicons in each sequence for visualization purposes.

To assign a specific SMN copy to each haplotype, we extracted FASTA sequence from SMN exon regions for each haplotype (n=101) and concatenated them into a single sequence. We constructed multiple sequence alignment using the R package DECIPHER[177] (v2.28.0). We then calculated the distance among all haplotypes and constructed an UPGMA tree using R package ape[178] (v5.7-1) and phangorn[179] (v2.11.1). We set the orangutan sequence as an outgroup. Finally, we split all human haplotypes into two groups representing *SMN1* and *SMN2* gene copies.

Additionally, we constructed a custom RepeatMasker[56] (v4.1.2) library containing the canonical exon sequences of *GTF2H2* (ENSG00000145736.15), *NAIP* (ENSG00000249437.8), *SERF1A* (ENSG00000172058.16), and *SMN1* (ENSG00000172062.17). Exon sequences were sourced from the Ensembl genome browser[81] (release 111). Utilizing RepeatMasker and this custom library, we identified all low divergence (<5%) gene exon hits within our sample haplotype assemblies. Exon sequences were retrieved using SAMtools[45] (v1.15.1), then stitched together to form the full gene exonic sequence. *SMN* gene copies were manually curated as either *SMN1* or *SMN2* based on carriage of either the C or T nucleotide in exon 7.

Finally, we utilized Illumina short read data from the 1000 Genomes Project for the same individuals, and processed it with Parascopy[180] (v1.16.0) and SMNCopyNumberCaller[181] (v1.1.2) to independently obtain *SMN1/2* copy numbers. Illumina-based and assembly-based copy number predictions matched perfectly across all 31 examined individuals.

#### Amylase haplotype detection using Verkko and optical genome mapping assemblies

We characterized the amylase locus, a ~212.5 kbp region on chromosome 1 (GRCh38; chr1:103,554,220–103,766,732), in 65 samples (**Supplementary Tables 3,57**). To resolve haplotype structures at this locus, we compared Verkko and optical genome mapping (OGM) datasets, as described previously[182,183]. Fragmented OGM assemblies (HG00268, HG01596, HG02011, HG02018, HG03456, and NA19983) were reprocessed using a local de novo assembly approach. The step-by-step process of this approach involves: 1-collecting unaligned molecules and molecules which aligned to amylase locus; 2-running de novo assembly using the molecules from step 1 and aligning the assembly to GRCh38p.12:

```
python2.7 Solve3.5.1_01142020/Pipeline/1.0/pipelineCL.py -T 64 -U
-j 64 -jp 64 -N 6 -f 0.25 -i 5 -w -c 3 \

-y \

-b ${bionano_bnx} \

-l ${output_dir} \

-t Solve3.5.1_01142020/RefAligner/1.0/ \
```

```
-a
Solve3.5.1_01142020/RefAligner/1.0/optArguments_haplotype_DLE1_sap
hyr_human.xml \

-r ${reference_genome}
```

To compare the Verkko amylase locus haplotype structures to OGM, we first converted fasta files of Verkko assemblies to in silico maps using fa2cmap_multi_color.pl (Bionano Solve™ v3.5.1). The resulting in silico maps and de novo assemblies of OGMs were aligned to the GRCh38p.12 reference assembly in silico map using the refAligner tool (Bionano Solve™ v3.5.1).

```
Solve3.5.1_01142020/RefAligner/1.0/RefAligner

-ref ${referenceFile} \ -maxthreads ${threads} \

-i ${inputfile} \

-o ${outputdir}EXP_REFINEFINAL1
```

Next, OGM contigs that aligned to amylase locus were pairwise aligned to Verkko contigs using the same alignment tool, refAligner [Bionano Solve™ (v3.5.1)]. Pairwise alignment was performed between Verkko hap1 and hap2, and OGM hap1 and hap2 contigs using refAligner alignment results. This process generated two separate alignment files for each sample. The variant files and alignment confidence scores produced by these alignments were used to determine which Verkko haplotype matched with which OGM contig. Matching pairs were validated by visual inspection using the Bionano Access™ software. Amylase haplotypes were characterized from these visualizations using amylase segments and in silico map labeling patterns as a guide. The amylase segments were determined as described previously[183,184]. Segments depicted by colored arrows were determined based on the labeling pattern from the Bionano Genomics optical genome mapping data and the sequence similarity between segments using blastn (BLASTN v2.9.0+)[185]. First, using optical genome mapping in silico map labeling patterns of GRCh38p.12 reference assembly, amylase segments were identified using pairwise alignments between smaller duplicons within segmental duplications. Then, to obtain accurate coordinates of segments, the GRCh38p.12 fasta sequence of the locus was aligned against itself using blastn (BLASTN 2.9.0+). The alignments with less than 99% sequence similarity and 60% coverage were filtered out and coordinates of all reference segments were finalized. We identified one copy of AMY2B (purple; 45.7 kbp), two copies of Intergenic1 (green; 5.3 kbp), one copy of AMY2A.1 (red; 12.4 kbp), two copies of AMY2A.2 (orange; 14.1 kbp), three copies of Intergenic2 (maroon; 4.3 kbp), and three copies of AMY1 (pink; 26.3 kbp) segments within the GRCh38p.12 amylase locus. Next, the GRCh38p.12 segment sequences were used as a guide to detect amylase segments from each sample included in this study using blastn. Same sequence similarity (> 99%) and coverage thresholds (> 60%) were used to obtain the amylase segment coordinates from each sample. Finally, all amylase haplotypes were visualized using pygenomeviz (https://github.com/moshi4/pyGenomeViz) to confirm haplotype structures and haplotypes were clustered using pgr-tk[156] (v0.5.1).

#### Results

There are ~60 genomic disorders associated with neurodevelopmental delay, autism, epilepsy and intellectual disability where recurrent CNVs are associated with large complex regions of segmental duplications. The organization and orientation of the SDs is a critical risk factor in predisposing to NAHR that predispose and/or protect from CNV risk[13,186–188]. These SD rich regions have also been a hotspot for nonallelic gene conversion that plays an important role in evolution of many multi-copy genes[189]. Here, we focus on a biomedically relevant and structurally complex region containing *SMN1* and *SMN2* gene copies. *SMN1* is the primary gene associated with spinal muscular atrophy (SMA) and its duplicated copy, SMN2, which modifies SMA severity and is a target of one of the most successful ASO-mediated gene therapies. The genes are embedded in a very large SD region (~1.5 Mbp) that has been almost impossible to fully sequence and assemble over the last two decades[4,28,38,190] (**Supplementary Fig. 46**). In addition to recently released HPRC assemblies, we were able to fully assemble the *SMN1/2* region in approximately two-thirds of the haplotypes in this study, providing us a unique opportunity to investigate the structural complexity of this region (**Fig. 5b**). We characterized this combined set of 101 complete human assemblies of this region that allows us to map copy number (full or partial gene alignments) of *SMN1/2* genes and detailed structure of this region. Focusing on multi-copy genes *SMN1/2*, *SERF1A/B* and *GTF2H2/C* we find that about half (n=48) of all haplotypes carry exactly two copies (aligned to two distinct regions in a single haplotype) of each of these genes. On the other hand *NAIP* gene is present mostly in a single copy (gene alignment per haplotype). (**Fig. 5c**). We highlight 11 human haplotypes including orangutan that carry distinct gene copy numbers and are organized into structurally diverse duplicon structures composed of multiple palindromic regions (**Fig. 5d**). We further defined functional *SMN1* copy and duplicated *SMN2* copy in each haplotype using orangutan ancestral *SMN1* copy as an outgroup (**Supplementary Fig. 47**). For samples where both haplotypes are fully assembled (n=31) we report 100% concordance with the short-read based genotyping methods (**Supplementary Fig. 48**). We have assigned 98 haplotypes to carry ancestral SMN1 copy and 3 haplotypes that do not (**Fig. 5e, Supplementary Fig. 49**). We speculated that such haplotypes are likely to occur via gene conversion. Evaluation of an extent of directly oriented and highly identical repeats we define 2 haplotypes that are likely predisposed to CNVs via NAHR.

The amylase locus in the human genome spans approximately 212.5 kilobase pairs (kbp) and is located on chromosome 1 (GRCh38; chr1:103,554,220–103,766,732). This locus contains the *AMY2B*, *AMY2A*, *AMY1A*, *AMY1B*, and *AMY1C* genes (**Supplementary Fig. 50**). We constructed haplotype-resolved diploid assemblies for 65 individuals, resulting in a total of 130 sampled alleles (haploid sample size; **Supplementary Table 57**). By analyzing copy number and orientation of the amylase segments, we identified 35 distinct amylase haplotypes (**Supplementary Fig. 50**, **Supplementary Table 57**). These haplotypes were supported by both Verkko and optical genome mapping de novo assemblies. To classify these haplotypes, we adhered to the established nomenclature[184]. The notation HXAYBZ represents the copy number of *AMY1*, *AMY2A*, and *AMY2B* genes, respectively, with superscript “a" indicating ancestral and superscript “r” indicating reference haplotypes (**Supplementary Fig. 50**). Notably, this study described the largest number of base-pair resolution amylase haplotypes described to date. The length of these amylase haplotypes varies, ranging from 111 kbp ($H1^{a}$.1 and $H1^{a}$.2) to 582 kbp (H11.1) (**Supplementary Fig. 50**), capturing those that are structurally identical to the GRCh38 ($H3^{r}$.1) and the T2T-chm13

(H7.3) assemblies. Among the identified haplotypes, four common ones, $H1^{a}$.1 (n = 14), $H3^{r}$.1 (n = 13), $H3^{r}$.2 (n = 19), and $H3^{r}$.4 (n = 22), constitute approximately 57% of all amylase haplotypes. Notably, 23 out of 35 distinct amylase haplotypes are singletons. Furthermore, this study provides base-pair level resolution for nine haplotypes that were previously supported only by optical genome mapping data. Finally, we introduce a novel haplotype, H11. 1, which also has base-pair level resolution.

### Centromeres

Contributing authors: Evan E. Eichler, Miriam K. Konkel, Mark Loftus, Glennis A. Logsdon, Keisuke K. Oshima

#### Methods

##### Centromere identification and annotation

To identify the centromeric regions within each Verkko and hifiasm genome assembly, we first aligned the whole-genome assemblies to the T2T-CHM13 (v2.0) reference genome[23] using minimap2[24] (v2.24) with the following parameters: -ax asm20 --secondary=no -s 25000 -K 15G --eqx --cs. We filtered the alignments to only those contigs that traversed each human centromere, from the p- to the q-arm, using BEDTools intersect (v2.29.0)[54]. Then, we ran dna-brnn (v0.1; https://pubmed.ncbi.nlm.nih.gov/30989183/) on each centromeric contig to identify regions containing α-satellite sequences, as indicated by a “2”. Once we had identified the regions containing α-satellite sequences, we ran RepeatMasker[56] (v4.1.0) to identify all repeat elements and their organization within the centromeric region, and we also ran Hum-AS-HMMER (https://github.com/fedorrik/HumAS-HMMER_for_AnVIL) to identify α-satellite higher-order repeat (HOR) sequence composition and their organization. We used the resulting RepeatMasker and HumAS-AMMER stv_row.bed file to visualize the organization of the α-satellite HOR arrays with R[191] (v1.1.383) and the ggplot2 package[192].

##### Validation of centromeric regions

We validated the construction of each centromeric region by first aligning native PacBio HiFi and ONT data from the same genome to each relevant whole-genome assembly using pbmm2 (v1.1.0; https://github.com/PacificBiosciences/pbmm2; for PacBio HiFi data) or minimap2[24] (v2.28; for ONT data). We, then, assessed the assemblies for uniform read depth across the centromeric regions via IGV[95] and NucFreq[27]. Centromeres that were found to have a collapse in sequence, false duplication of sequence, and/or misjoin were flagged and removed from our analysis.

##### Estimation of α-satellite HOR array length

To estimate the length of the α-satellite HOR arrays for each human centromere, we first ran Hum-AS-HMMER (https://github.com/fedorrik/HumAS-HMMER_for_AnVIL) on the centromeric regions using the hmmer-run.sh script and the AS-HORs-hmmer3.0-170921.hmmHidden Markov Model. Then, we used the stv_row.bed file to calculate the length of the α-satellite HOR arrays by taking the minimum and maximum coordinate of the “live” α-satellite HOR arrays, marked by an “L”, and plotting their lengths with Graphpad Prism (v9).

#### Pairwise sequence identity heatmaps

To generate pairwise sequence identity heatmaps of each centromeric region, we ran StainedGlass[193] (v6.7.0) with the following parameters: window=5000, mm_f=30000, and mm_s=1000. We normalized the color scale across the StainedGlass plots by binning the % sequence identities equally and recoloring the data points according to the binning.

#### CpG methylation analysis

To determine the CpG methylation status of each centromere, we aligned ONT reads >30 kbp in length from the same source genome to the relevant whole-genome assembly via minimap2[24] (v2.28) and then assessed the CpG methylation status of the centromeric regions with Epi2me modbam2bed (https://github.com/epi2me-labs/modbam2bed; v0.10.0) and the following parameters: -e -m 5mC --cpg. We converted the resulting BED file to a bigWig using the bedGraphToBigWig tool (https://www.encodeproject.org/software/bedgraphtobigwig/) and then visualized the file in IGV. To determine the length of the hypomethylated region (termed “centromere dip region”, or CDR[194,195] in each centromere, we used CDR-Finder (https://github.com/EichlerLab/CDR-Finder_smk). This tool first bins the assembly into 5 kbp windows, computes the median CpG methylation frequency within windows containing α-satellite (as determined by RepeatMasker[56] (v4.1.0), selects bins that have a lower CpG methylation frequency than the median frequency in the region, merges consecutive bins into a larger bin, filters for merged bins that are >50 kbp, and reports the location of these bins.

### Supplementary Figures

#### Fig. 1

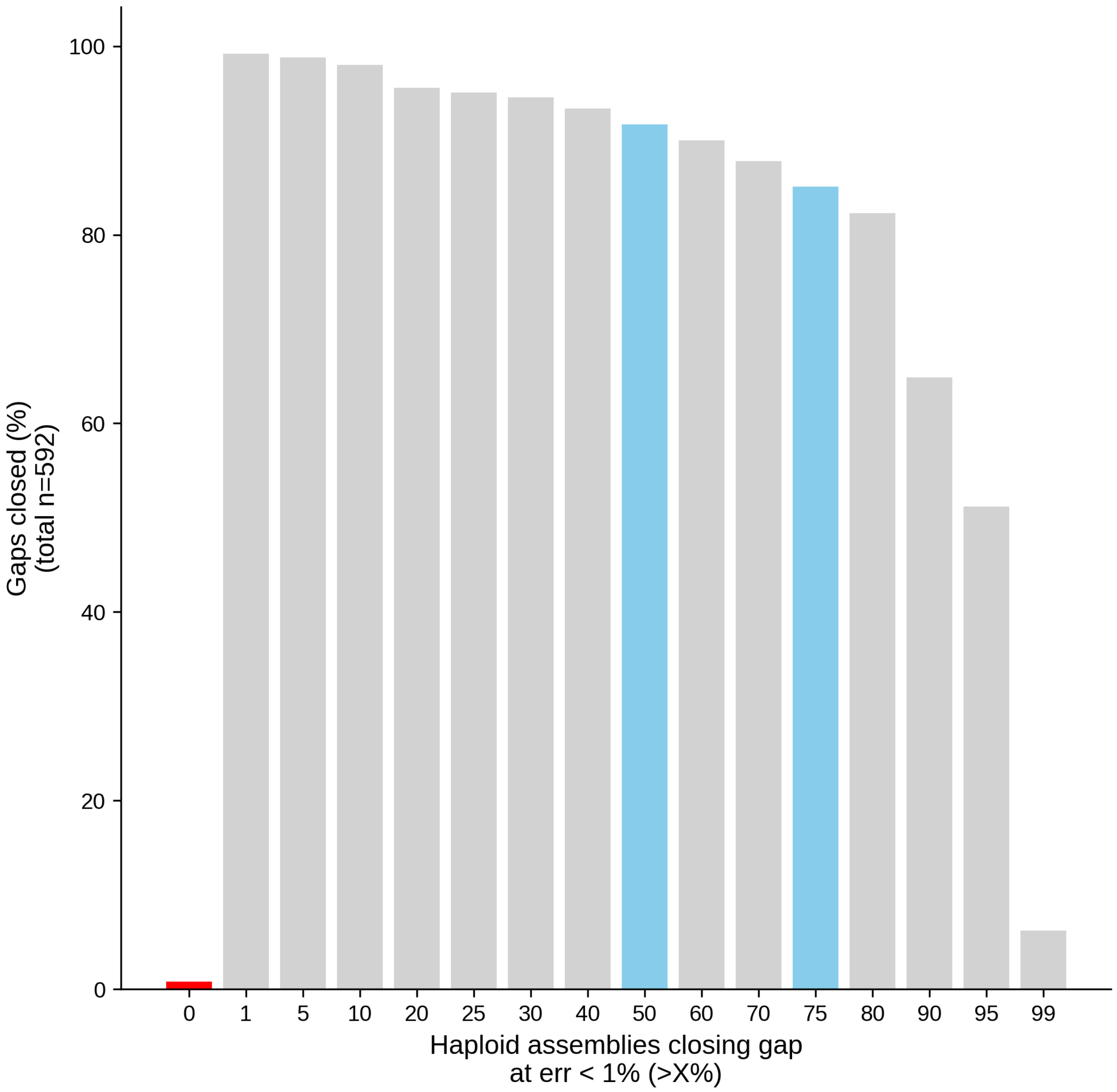


**Supplementary Figure 1. Closed HPRC gaps.** Bar chart depicting the percentage of closed HPRC gaps (vertical axis) [38] at increasingly stringent thresholds for the percentage of haploid assemblies closing the gap (horizontal axis). Highlights are shown for gaps that are never closed (red bar, n=5, ~0.8%), for gaps that are closed in more than 50% of all haploid assemblies (left blue bar, n=543, ~91.7%) and for gaps that are closed in at least 75% of all haploid assemblies (right blue bar, n=504, ~85.1%). For this summary, the error rate of the contigs closing the gap was set to less than 1% (“err < 1%”).

Fig. 2

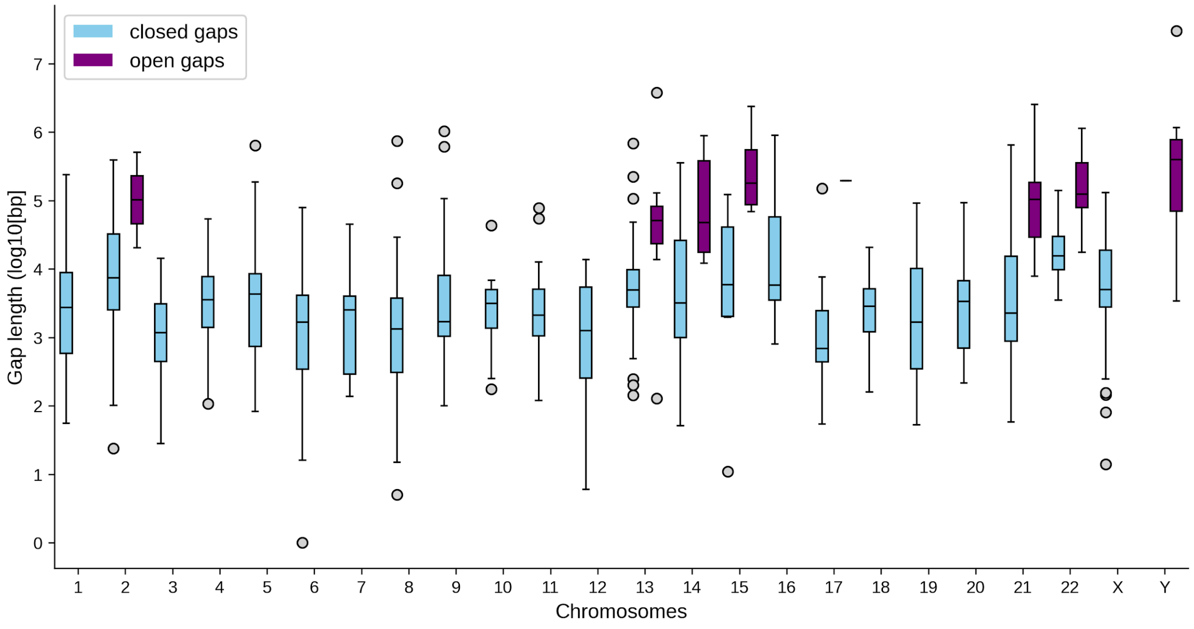


**Supplementary Figure 2. Size distribution of HPRC gaps.** Boxplots of gap lengths (vertical axis, log10) stratified by chromosome (horizontal axis) and status [closed (blue) and, if applicable, open (purple)] of previously reported HPRC gaps[38]**.** In this summary, gaps were counted as "closed" at an error rate of <1% (**Methods**) if they were labeled as closed in more than 50% of all haploid assemblies and "open" otherwise. Boxplot whiskers depict 1.5*(interquartile range); outliers beyond that range appear as gray points; the horizontal line indicates the median.

Fig. 3

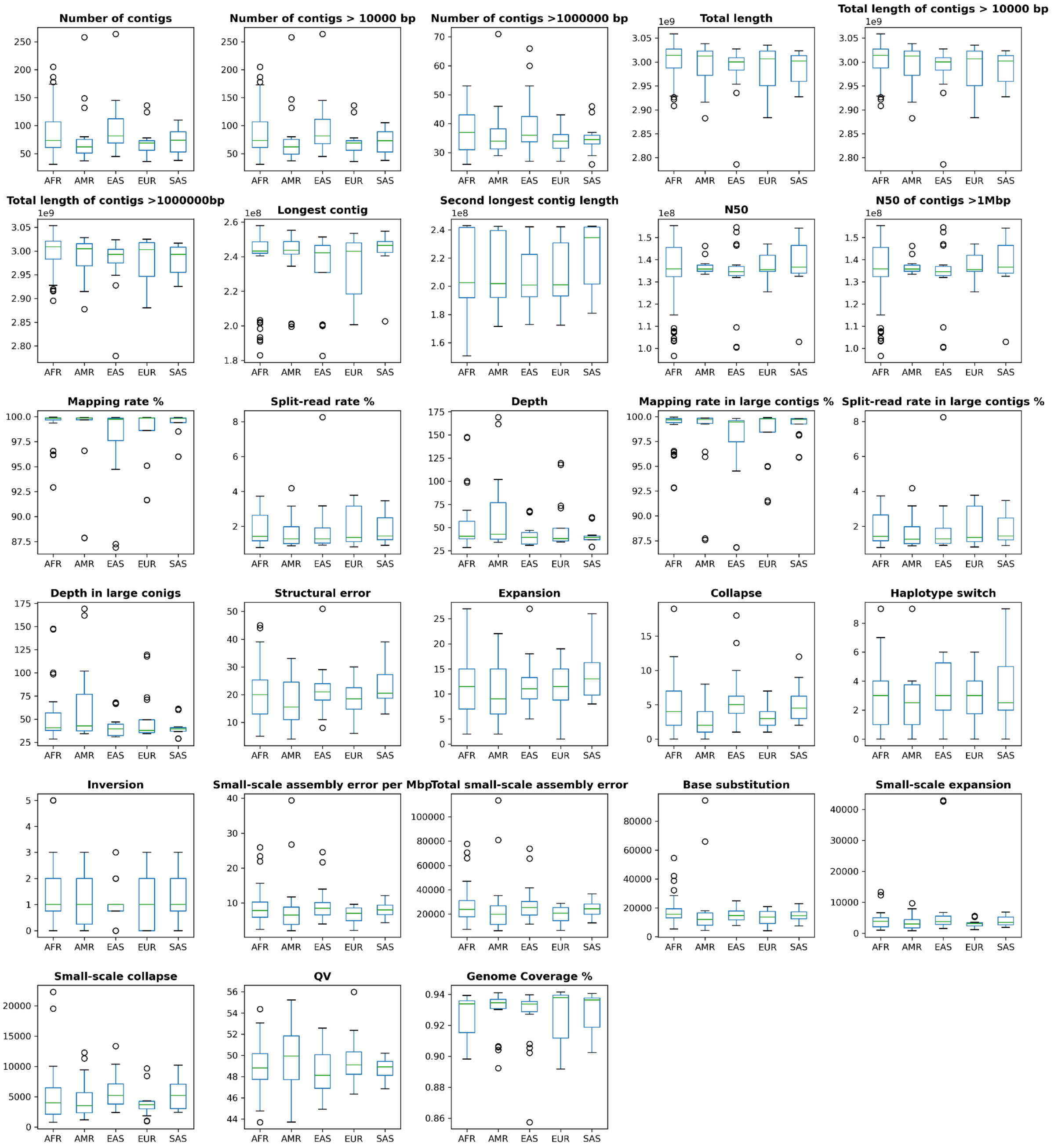


**Supplementary Figure 3. Verkko assemblies quality evaluation with HiFi raw reads.** Boxplot representation of each evaluated parameters for 65 samples with two haplotypes from Verkko assembler. Evaluation was performed using Inspector v1.2[37] with HiFi raw reads. Results were grouped in superpopulation. The boxplot whiskers depict 1.5* interquartile reange, outliers beyond that range appear as grey circles; the horizontal line indicates the median.

Fig. 4

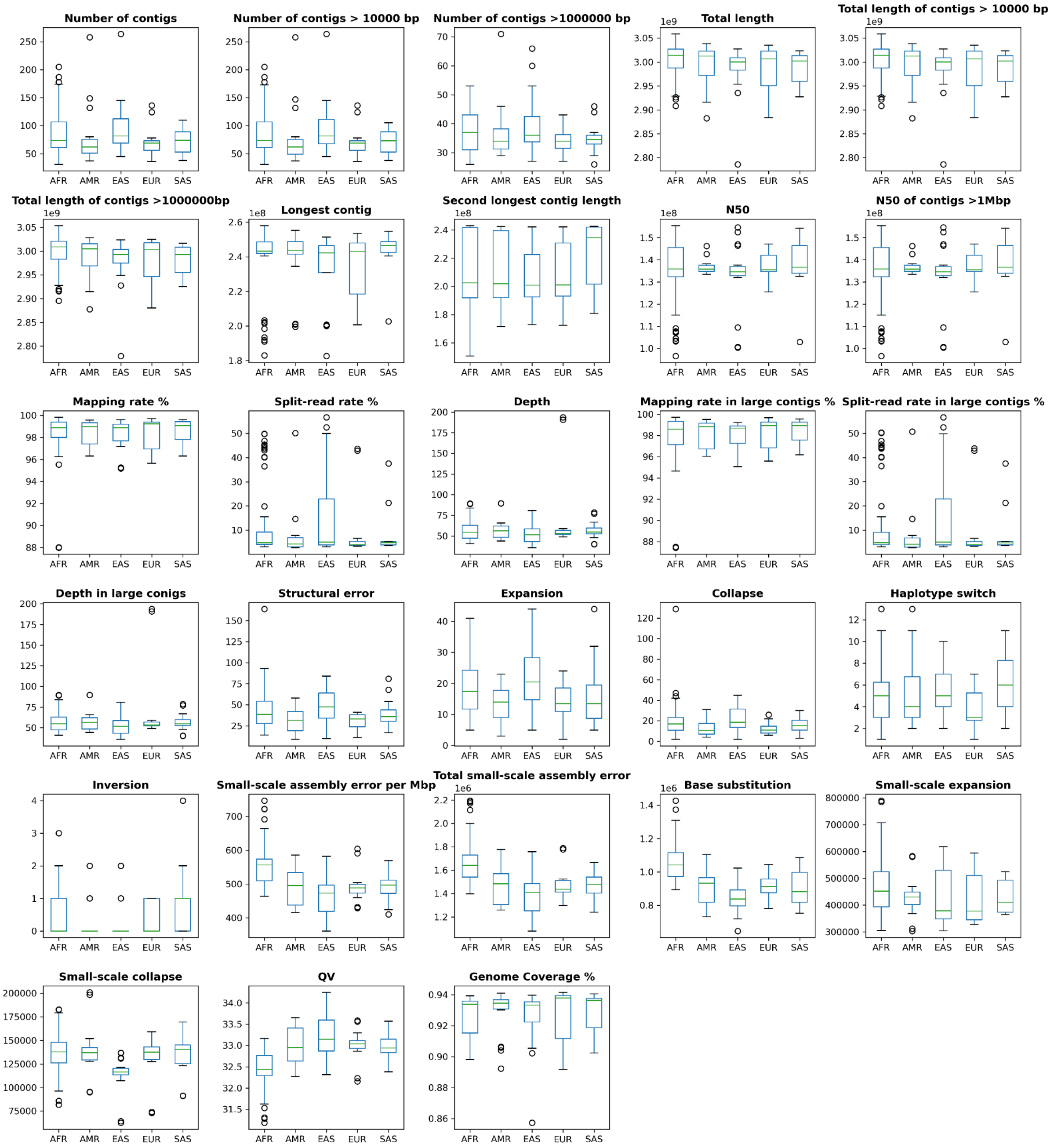


**Supplementary Figure 4. Verkko assemblies quality evaluation with ONT raw reads.** Boxplot representation of each evaluated parameters for 65 samples with two haplotypes from Verkko assembler. Evaluation was performed using Inspector v1.2[37] with ONT raw reads. Results were grouped in superpopulation. The boxplot whiskers depict 1.5* interquartile reange, outliers beyond that range appear as grey circles; the horizontal line indicates the median.

Fig. 5

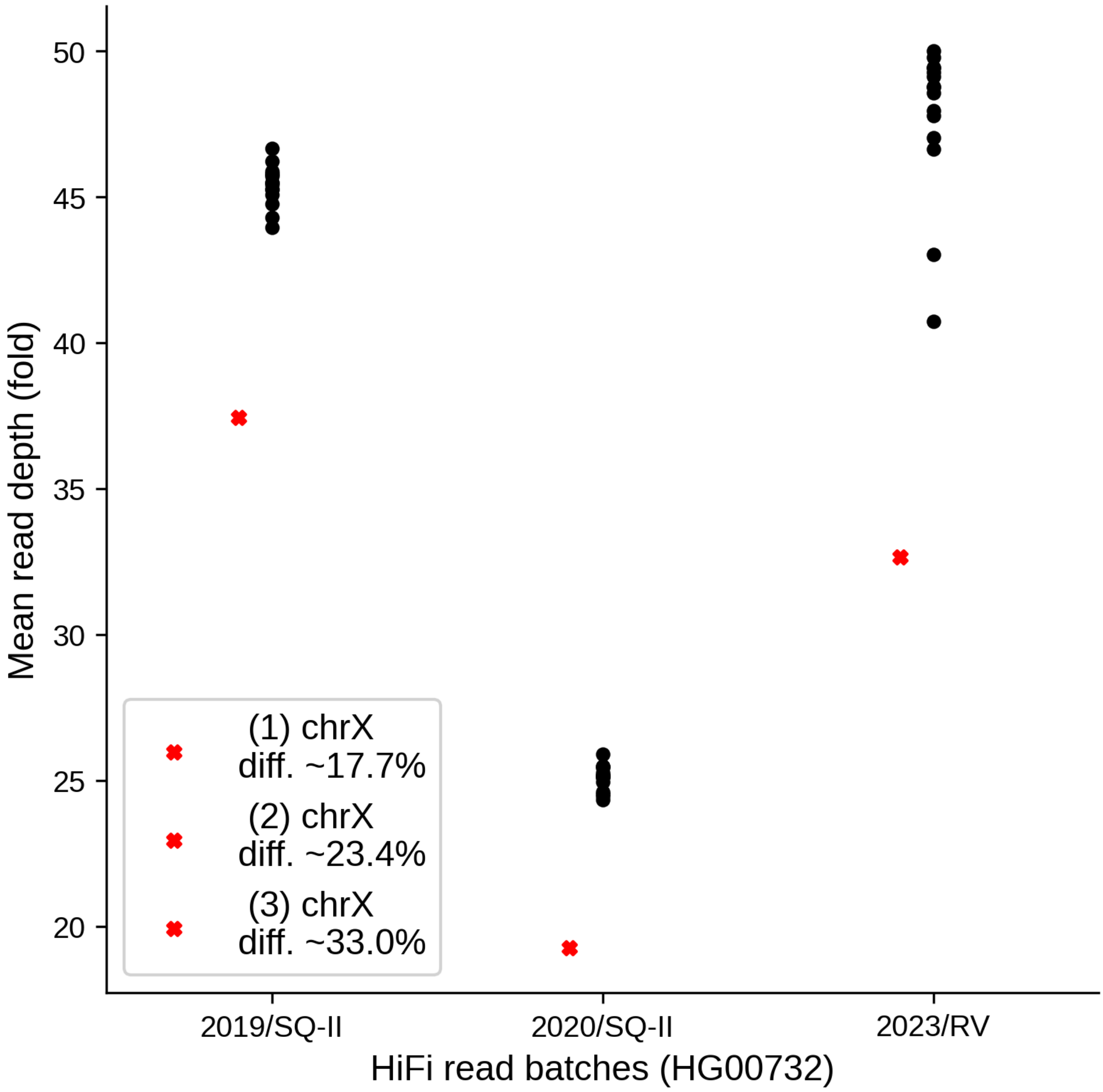


**Supplementary Figure 5. HiFi read coverage dropout over Chromosome X in sample HG00732.** Summary of mean HiFi read fold coverage stratified by sequencing batch/year (horizontal axis) over main reference chromosomes (acrocentric chromosomes and chromosome Y excluded). The mean read depth for chromosome X is highlighted as single red “x” per batch/year. The relative difference of the X-chromosomal read depth compared to all other chromosomes is stated in the legend. SQ-II: PacBio Sequel-II platform; RV: PacBio Revio platform.

Fig. 6

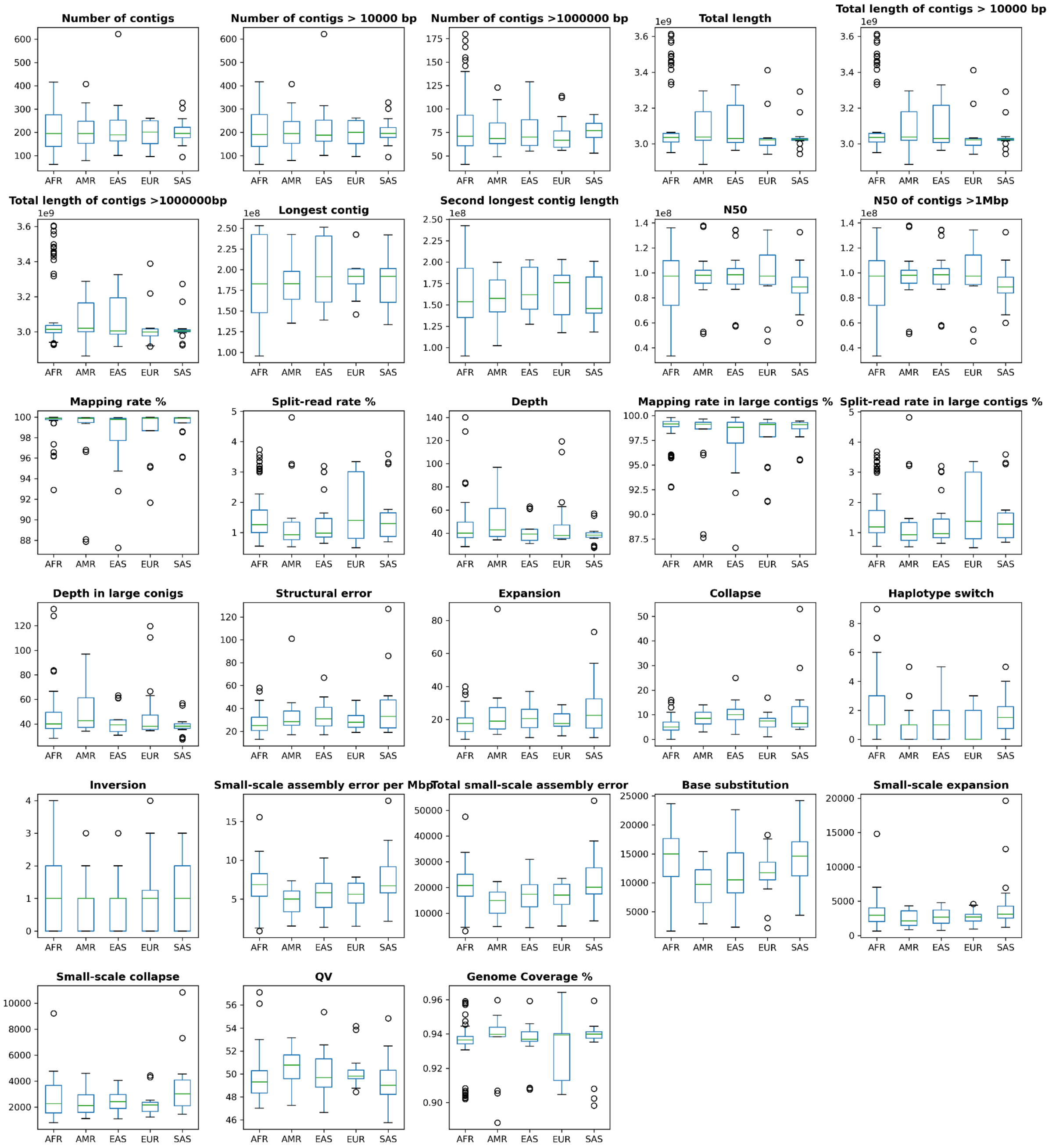


**Supplementary Figure 6. Hifiasm assemblies quality evaluation with HiFi raw reads.** Boxplot representation of each evaluated parameters for 65 samples with two haplotypes from Hifiasm assembler. Evaluation was performed using Inspector v1.2 [37] with HiFi raw reads. Results were grouped in superpopulation. The boxplot whiskers depict 1.5* interquartile reange, outliers beyond that range appear as grey circles; the horizontal line indicates the median.

Fig. 7

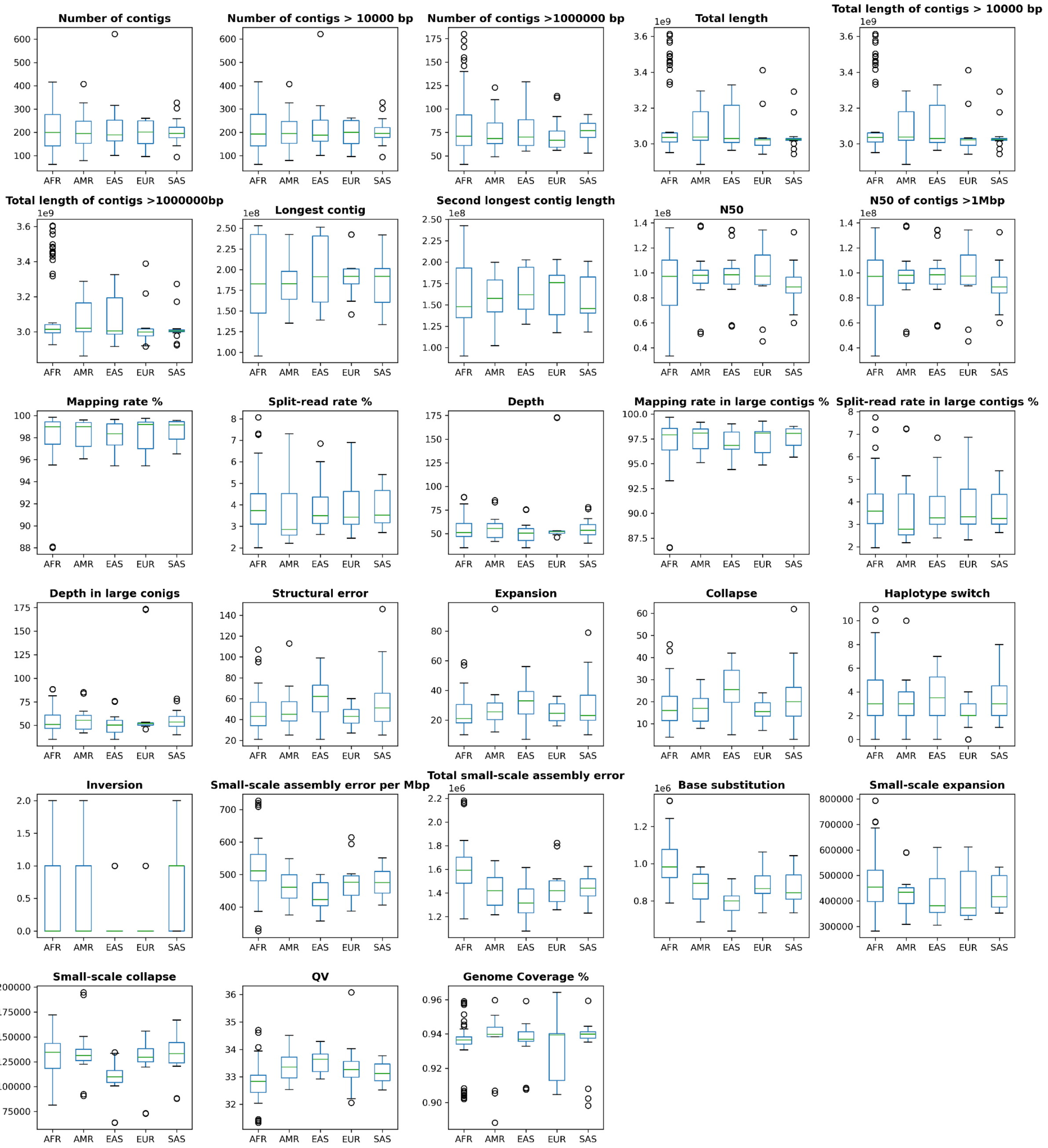


**Supplementary Figure 7. Hifiasm assemblies quality evaluation with ONT raw reads.** Boxplot representation of each evaluated parameters for 65 samples with two haplotypes from Hifiasm assembler. Evaluation was performed using Inspector v1.2 [37] with ONT raw reads. The boxplot whiskers depict 1.5* interquartile reange, outliers beyond that range appear as grey circles; the horizontal line indicates the median.

Fig. 8

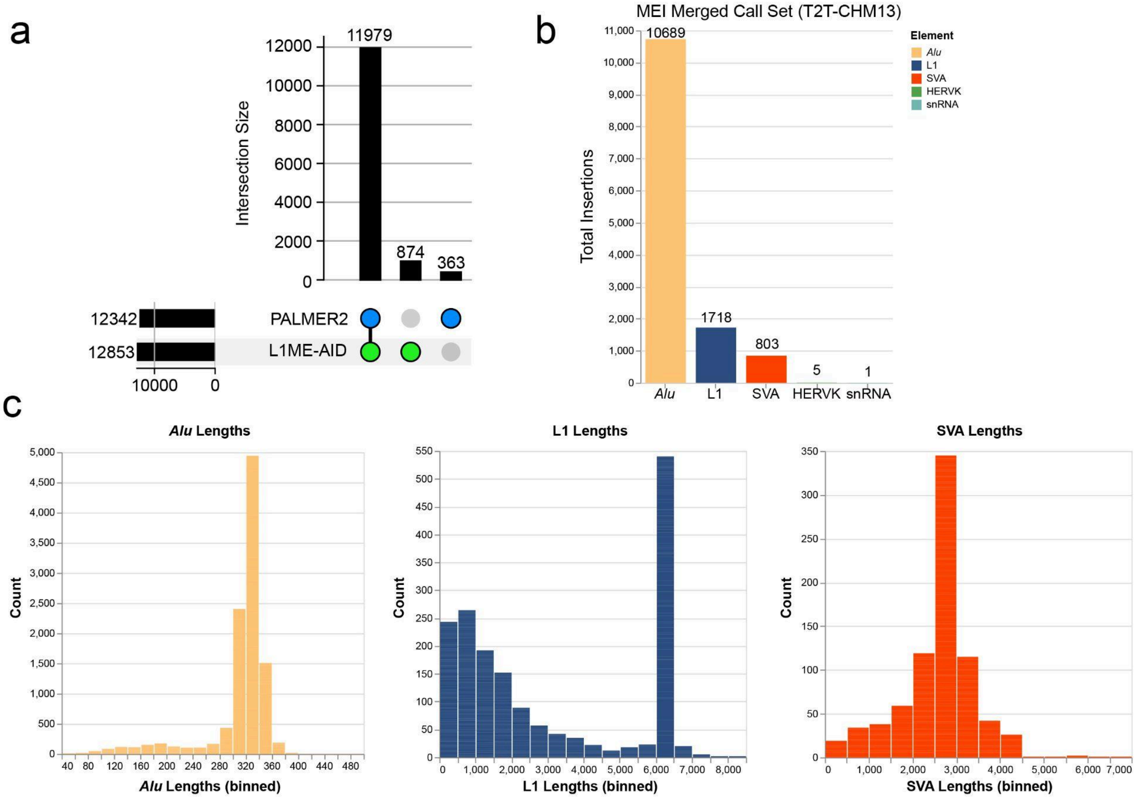


**Supplementary Figure 8. T2T-CHM13 MEI Merged Callset.** a) An upset plot showing the intersection of mobile element insertion (MEI) calls across the two individual MEI callsets built against the T2T-CHM13 reference genome. Most (90.6%) of the 13,216 MEIs were identified by both callers (PALMER2 and L1ME-AID). b) A bar graph displaying the total *Alu*, L1, SVA, HERV-K, and snRNA insertion calls within the merged callset. c) Three histograms displaying the length distribution of *Alu* elements (yellow), L1s (blue), and SVAs (red) within the merged callset.

Fig. 9

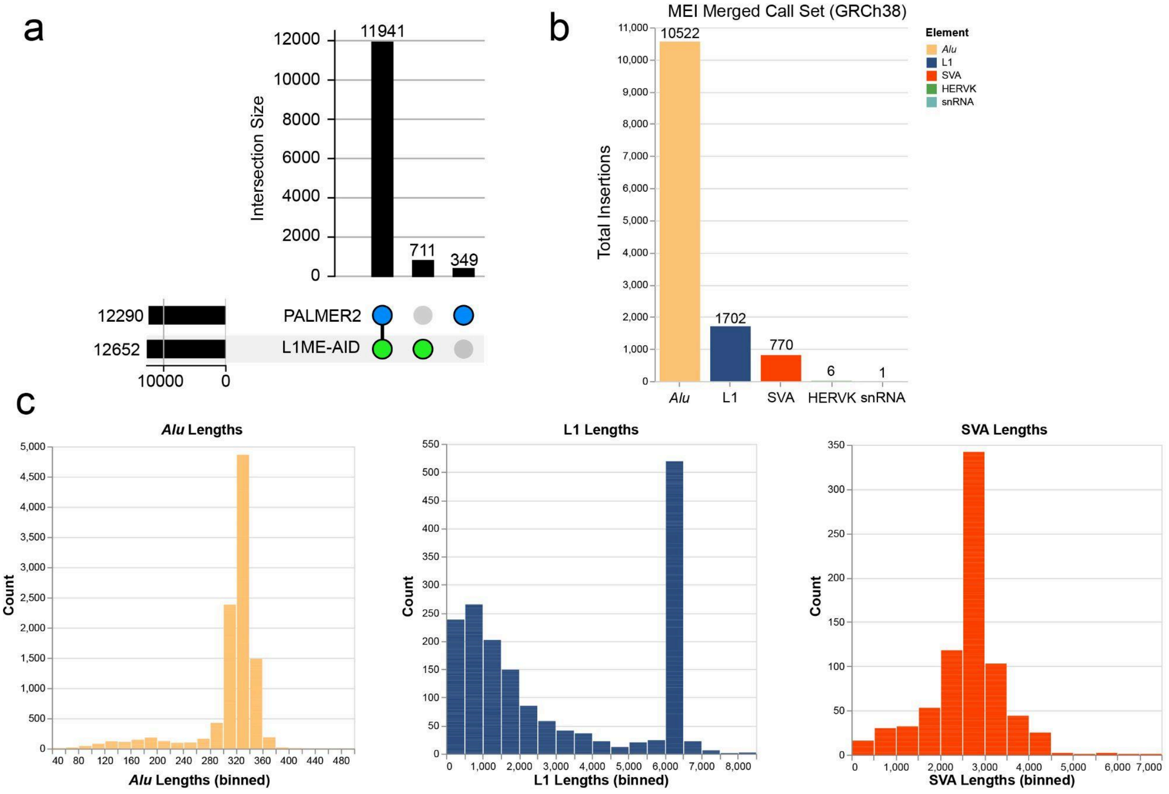


**Supplementary Figure 9. GRCh38 MEI Merged Callset. a)** An upset plot showing the intersection of mobile element insertion (MEI) calls across the two individual MEI callsets built against the GRCh38 reference genome. Most (91.8%) of the 13,001 MEIs were identified by both callers (PALMER2 and L1ME-AID). **b)** A bar graph displaying the total *Alu*, L1, SVA, HERV-K, and snRNA insertion calls within the merged callset. **c)** Three histograms displaying the length distribution of *Alu* elements (yellow), L1s (blue), and SVAs (red) within the merged callset.

Fig. 10

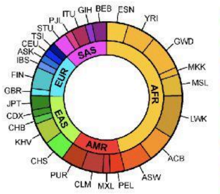


**Total Unique MEIs (GRCh38)**

| Samples (H1 + H2 Unique MEIs) | Sample INS | | | Reference INS | | |
|---|---|---|---|---|---|---|
| | *Alu* | L1 | SVA | *Alu* | L1 | SVA |
| HG02587 | 1622 | 224 | 96 | 1309 | 164 | 70 |
| HG02666 | 1600 | 211 | 85 | 1304 | 167 | 74 |
| HG02769 | 1607 | 222 | 115 | 1286 | 158 | 74 |
| HG02818 | 1587 | 218 | 100 | 1314 | 156 | 75 |
| HG02953 | 1616 | 227 | 105 | 1298 | 170 | 79 |
| HG03065 | 1578 | 222 | 106 | 1290 | 178 | 70 |
| HG03248 | 1606 | 231 | 84 | 1280 | 160 | 74 |
| HG03371 | 1621 | 225 | 94 | 1288 | 171 | 71 |
| HG03452 | 1616 | 229 | 113 | 1288 | 170 | 78 |
| HG03456 | 1614 | 222 | 97 | 1287 | 171 | 78 |
| HG03520 | 1638 | 215 | 103 | 1301 | 169 | 75 |
| NA19036 | 1650 | 208 | 102 | 1297 | 162 | 76 |
| NA19129 | 1647 | 216 | 113 | 1263 | 161 | 78 |
| NA19238 | 1629 | 226 | 104 | 1268 | 166 | 77 |
| NA19239 | 1593 | 233 | 97 | 1279 | 163 | 80 |
| NA19240 | 1683 | 226 | 105 | 1287 | 172 | 78 |
| NA19317 | 1598 | 230 | 94 | 1282 | 156 | 73 |
| NA19331 | 1640 | 228 | 112 | 1292 | 160 | 77 |
| NA19347 | 1580 | 211 | 100 | 1278 | 167 | 75 |
| NA19384 | 1619 | 205 | 105 | 1265 | 173 | 66 |
| NA19434 | 1586 | 208 | 105 | 1294 | 161 | 75 |
| NA21487 | 1558 | 229 | 105 | 1243 | 158 | 70 |
| HG01890 | 1596 | 213 | 98 | 1314 | 156 | 70 |
| HG02011 | 1562 | 203 | 97 | 1298 | 167 | 73 |
| HG02282 | 1538 | 234 | 93 | 1289 | 164 | 80 |
| HG02554 | 1610 | 207 | 100 | 1286 | 161 | 73 |
| NA19705 | 1519 | 244 | 94 | 1291 | 169 | 77 |
| NA19836 | 1556 | 213 | 88 | 1280 | 165 | 75 |
| NA19983 | 1558 | 228 | 88 | 1276 | 169 | 74 |
| NA20355 | 1555 | 212 | 107 | 1273 | 173 | 76 |
| HG00731 | 1216 | 200 | 80 | 1110 | 158 | 68 |
| HG00732 | 1255 | 196 | 84 | 1083 | 144 | 62 |
| HG00733 | 1273 | 179 | 78 | 1119 | 153 | 63 |
| HG01114 | 1275 | 188 | 74 | 1074 | 150 | 65 |
| HG01352 | 1210 | 188 | 74 | 1104 | 148 | 66 |
| HG01457 | 1272 | 194 | 76 | 1090 | 146 | 69 |
| HG01573 | 1207 | 187 | 78 | 1083 | 142 | 67 |
| HG02106 | 1168 | 178 | 72 | 1048 | 146 | 59 |
| NA19650 | 1266 | 186 | 80 | 1106 | 144 | 65 |
| HG00512 | 1225 | 205 | 83 | 1087 | 150 | 62 |
| HG00513 | 1220 | 199 | 79 | 1099 | 150 | 63 |
| HG00514 | 1210 | 202 | 82 | 1095 | 151 | 61 |
| HG00864 | 1228 | 177 | 90 | 1112 | 151 | 72 |
| HG01596 | 1211 | 189 | 77 | 1085 | 147 | 59 |
| HG02018 | 1279 | 194 | 78 | 1124 | 157 | 66 |
| HG02059 | 1214 | 191 | 86 | 1077 | 147 | 63 |
| NA18534 | 1224 | 203 | 90 | 1092 | 141 | 64 |
| NA18939 | 1234 | 186 | 85 | 1100 | 144 | 61 |
| NA18989 | 1231 | 184 | 73 | 1098 | 141 | 60 |
| HG00096 | 1242 | 167 | 64 | 1094 | 143 | 68 |
| HG00171 | 1238 | 186 | 73 | 1074 | 144 | 66 |
| HG00268 | 1217 | 179 | 69 | 1077 | 147 | 60 |
| HG00358 | 1242 | 194 | 71 | 1058 | 148 | 64 |
| HG01505 | 1274 | 178 | 79 | 1082 | 146 | 61 |
| NA12329 | 1229 | 174 | 68 | 1080 | 155 | 63 |
| NA20509 | 1250 | 187 | 75 | 1091 | 138 | 66 |
| NA24385 | 1237 | 166 | 83 | 1085 | 138 | 65 |
| HG02492 | 1228 | 178 | 89 | 1104 | 148 | 72 |
| HG03009 | 1288 | 184 | 80 | 1111 | 141 | 66 |
| HG03683 | 1288 | 194 | 78 | 1118 | 148 | 65 |
| HG03732 | 1246 | 220 | 86 | 1112 | 154 | 61 |
| HG03807 | 1304 | 195 | 80 | 1117 | 155 | 64 |
| HG04036 | 1317 | 187 | 84 | 1130 | 139 | 74 |
| HG04217 | 1298 | 197 | 76 | 1096 | 145 | 64 |
| NA20847 | 1257 | 181 | 85 | 1102 | 155 | 66 |

**Total Unique MEIs (T2T-CHM13)**

| Samples (H1 + H2 Unique MEIs) | Sample INS | | | Reference INS | | |
|---|---|---|---|---|---|---|
| | *Alu* | L1 | SVA | *Alu* | L1 | SVA |
| HG02587 | 1671 | 224 | 93 | 1273 | 164 | 53 |
| HG02666 | 1624 | 212 | 91 | 1274 | 161 | 58 |
| HG02769 | 1639 | 229 | 114 | 1271 | 164 | 61 |
| HG02818 | 1637 | 224 | 98 | 1297 | 163 | 61 |
| HG02953 | 1653 | 225 | 100 | 1300 | 170 | 60 |
| HG03065 | 1654 | 226 | 110 | 1261 | 170 | 58 |
| HG03248 | 1683 | 240 | 87 | 1246 | 168 | 60 |
| HG03371 | 1648 | 236 | 95 | 1267 | 164 | 54 |
| HG03452 | 1643 | 239 | 113 | 1262 | 170 | 67 |
| HG03456 | 1668 | 231 | 103 | 1282 | 174 | 63 |
| HG03520 | 1709 | 223 | 106 | 1275 | 168 | 57 |
| NA19036 | 1684 | 215 | 101 | 1250 | 161 | 57 |
| NA19129 | 1706 | 211 | 117 | 1228 | 153 | 63 |
| NA19238 | 1677 | 231 | 112 | 1251 | 171 | 60 |
| NA19239 | 1645 | 237 | 86 | 1252 | 162 | 59 |
| NA19240 | 1740 | 239 | 105 | 1263 | 170 | 57 |
| NA19317 | 1612 | 238 | 97 | 1268 | 161 | 56 |
| NA19331 | 1669 | 230 | 116 | 1250 | 160 | 66 |
| NA19347 | 1638 | 221 | 100 | 1234 | 164 | 55 |
| NA19384 | 1671 | 202 | 101 | 1258 | 167 | 55 |
| NA19434 | 1636 | 195 | 95 | 1241 | 159 | 60 |
| NA21487 | 1587 | 223 | 99 | 1202 | 158 | 55 |
| HG01890 | 1650 | 220 | 101 | 1266 | 160 | 57 |
| HG02011 | 1576 | 210 | 101 | 1255 | 160 | 62 |
| HG02282 | 1566 | 228 | 94 | 1228 | 163 | 61 |
| HG02554 | 1616 | 211 | 111 | 1255 | 163 | 59 |
| NA19705 | 1590 | 238 | 92 | 1254 | 170 | 59 |
| NA19836 | 1580 | 214 | 88 | 1249 | 159 | 54 |
| NA19983 | 1600 | 235 | 90 | 1242 | 165 | 58 |
| NA20355 | 1592 | 210 | 115 | 1240 | 166 | 64 |
| HG00731 | 1124 | 168 | 76 | 996 | 132 | 51 |
| HG00732 | 1162 | 187 | 77 | 1001 | 133 | 46 |
| HG00733 | 1171 | 157 | 67 | 1013 | 137 | 50 |
| HG01114 | 1172 | 167 | 68 | 982 | 135 | 46 |
| HG01352 | 1160 | 168 | 74 | 1032 | 139 | 50 |
| HG01457 | 1197 | 168 | 72 | 1010 | 137 | 53 |
| HG01573 | 1140 | 155 | 72 | 974 | 129 | 46 |
| HG02106 | 1103 | 159 | 66 | 968 | 134 | 47 |
| NA19650 | 1204 | 177 | 78 | 1024 | 139 | 45 |
| HG00512 | 1171 | 181 | 84 | 1022 | 146 | 51 |
| HG00513 | 1159 | 179 | 69 | 1002 | 141 | 43 |
| HG00514 | 1171 | 186 | 81 | 994 | 146 | 42 |
| HG00864 | 1162 | 162 | 85 | 1039 | 137 | 56 |
| HG01596 | 1174 | 180 | 75 | 1029 | 149 | 44 |
| HG02018 | 1206 | 183 | 77 | 1034 | 149 | 51 |
| HG02059 | 1154 | 183 | 85 | 1012 | 139 | 48 |
| NA18534 | 1179 | 190 | 79 | 1010 | 146 | 47 |
| NA18939 | 1178 | 174 | 84 | 1030 | 139 | 50 |
| NA18989 | 1176 | 174 | 76 | 1006 | 139 | 46 |
| HG00096 | 1159 | 156 | 57 | 986 | 131 | 48 |
| HG00171 | 1158 | 175 | 66 | 985 | 136 | 50 |
| HG00268 | 1126 | 157 | 64 | 980 | 138 | 41 |
| HG00358 | 1177 | 179 | 68 | 978 | 140 | 41 |
| HG01505 | 1199 | 159 | 70 | 984 | 130 | 48 |
| NA12329 | 1161 | 157 | 62 | 990 | 137 | 52 |
| NA20509 | 1184 | 176 | 68 | 1017 | 130 | 44 |
| NA24385 | 1171 | 162 | 74 | 1010 | 136 | 49 |
| HG02492 | 1168 | 182 | 79 | 1004 | 137 | 56 |
| HG03009 | 1213 | 179 | 74 | 998 | 136 | 46 |
| HG03683 | 1191 | 185 | 73 | 1029 | 142 | 49 |
| HG03732 | 1218 | 209 | 76 | 1037 | 154 | 44 |
| HG03807 | 1241 | 183 | 79 | 1028 | 143 | 50 |
| HG04036 | 1219 | 176 | 80 | 1016 | 136 | 56 |
| HG04217 | 1219 | 176 | 64 | 1017 | 141 | 45 |
| NA20847 | 1162 | 176 | 78 | 1012 | 154 | 45 |

**Supplementary Figure 10. Sample Mobile Element Insertion Counts.** Two heatmaps (GRCh38 (left) and T2T-CHM13 (right)) showing the total unique sample-specific and reference-specific mobile element insertions per sample. Both heatmaps show samples from African descent containing a greater total number of sample-specific MEIs.

Fig. 11

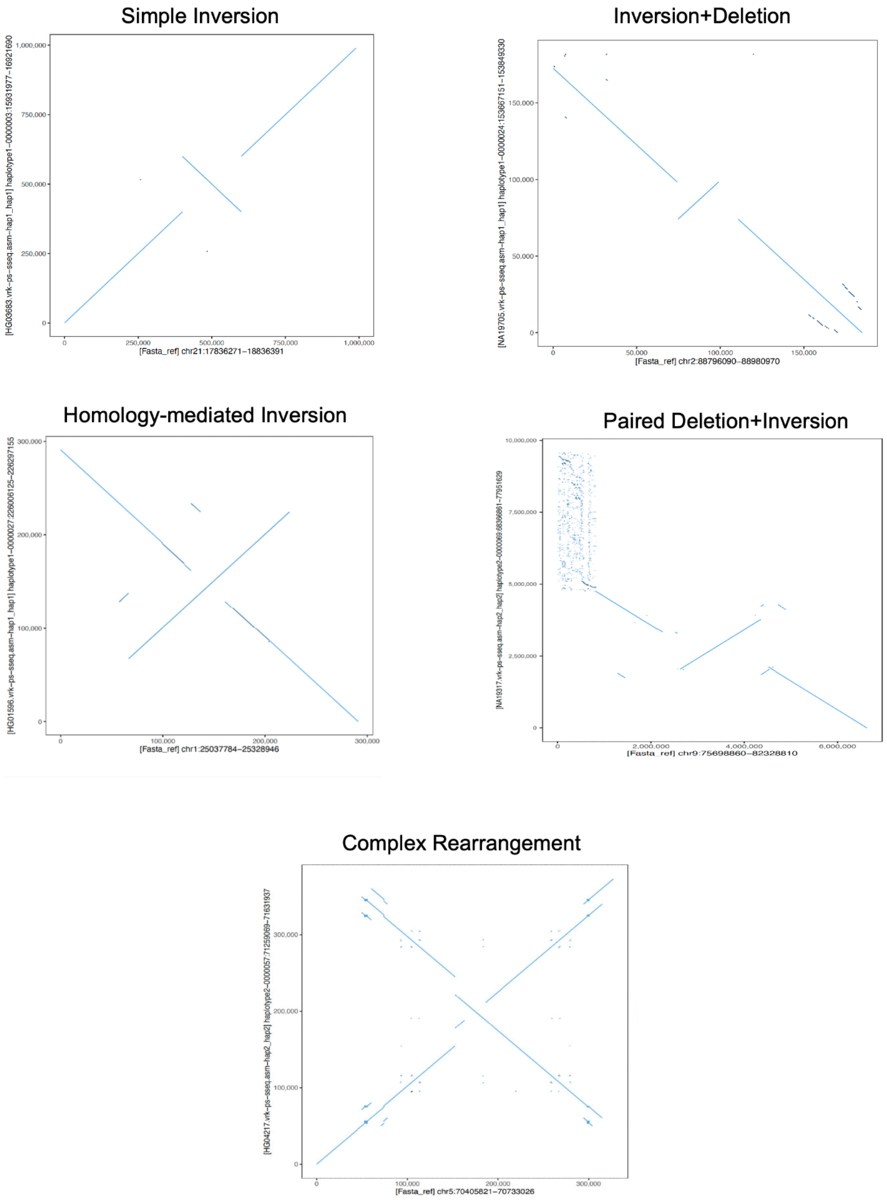


**Supplementary Figure 11. Inversion Dotplot Analysis.** Representative dot plots illustrating the categorization of inversions in different inversion classes in our dataset. The y-axis represents the specified genomic location in one of the inversion carrier samples, while the x-axis represents the corresponding region in the CHM13 reference genome. Each candidate inversion locus was manually examined through dot plot analysis, facilitating the validation of candidate inversion locations and their classification into one of the five distinct categories showcased in the graph.

Fig. 12

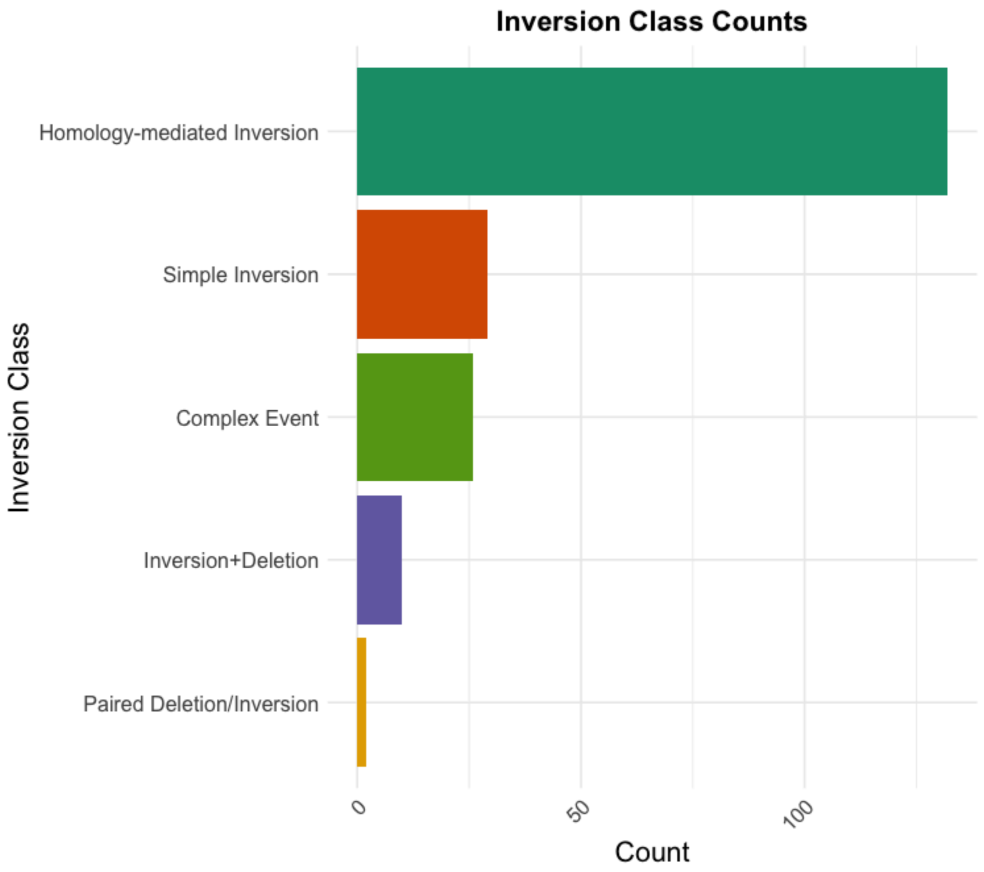


**Supplementary Figure 12. Inversion Classes.** The bar plot represents the count of instances for each inversion class identified in the manual dotplot analysis.

Fig. 13

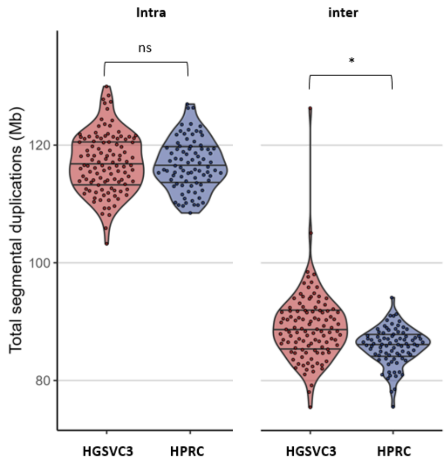


**Supplementary Figure 13. Comparison of segmental duplication content to the HPRC draft genomes.** Total SD content is compared across this study (HGSVC3) and the HPRC draft genomes[28] (HPRC). Three horizontal lines in the violin indicate the first quartile, median and the third quartile. Significance shown on the top indicates two-tailed Wilcox ranked sum test.

Fig. 14

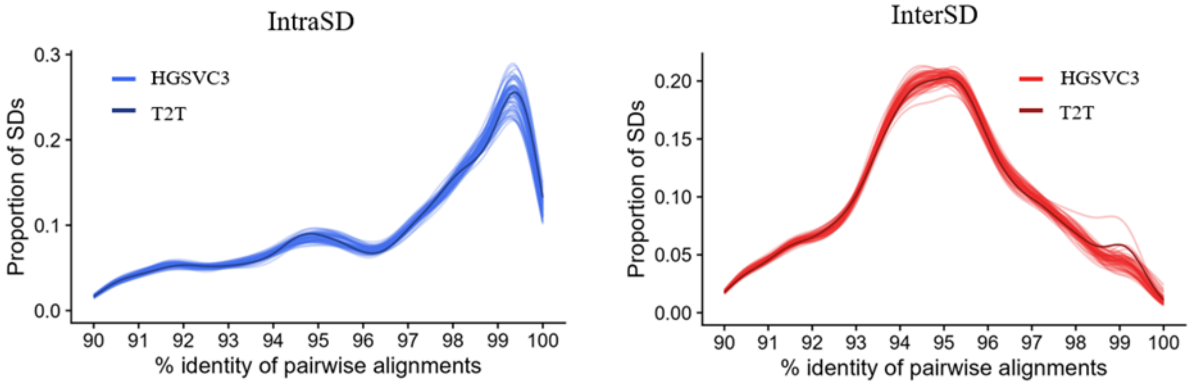


**Supplementary Figure 14. Inter- and intra-SD content.** Distribution pairwise identity among inter/intra-chromosomal segmental duplication pairs in the HGSVC genomes (n=126 haplotypes).

Fig. 15

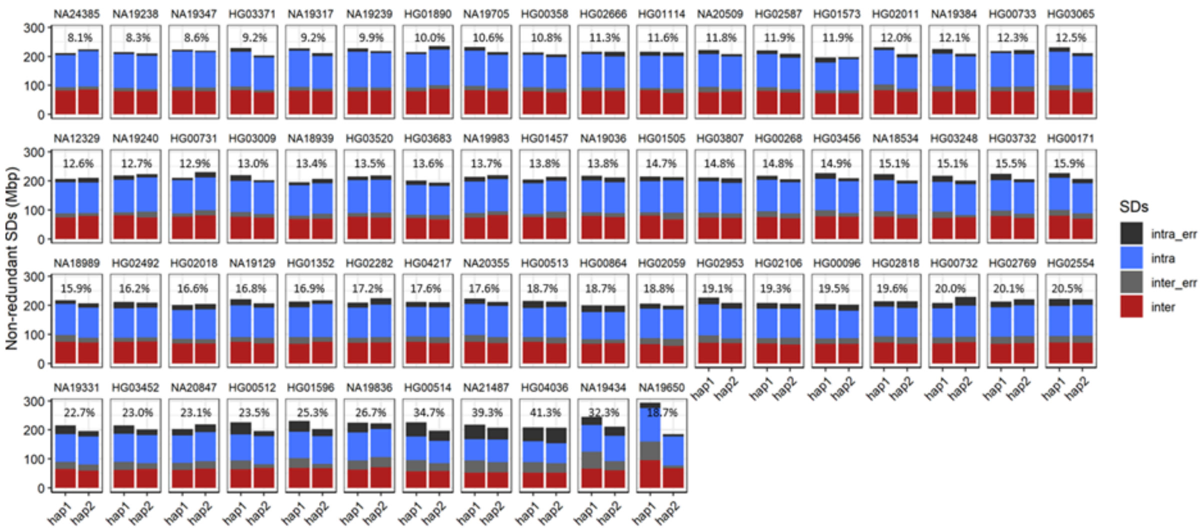


**Supplementary Figure 15. Non-redundant SD content.** Nonredundant bases of inter/intra-chromosomal segmental duplications and those that overlap with flagged regions via Flagger/NucFreq.

Fig. 16

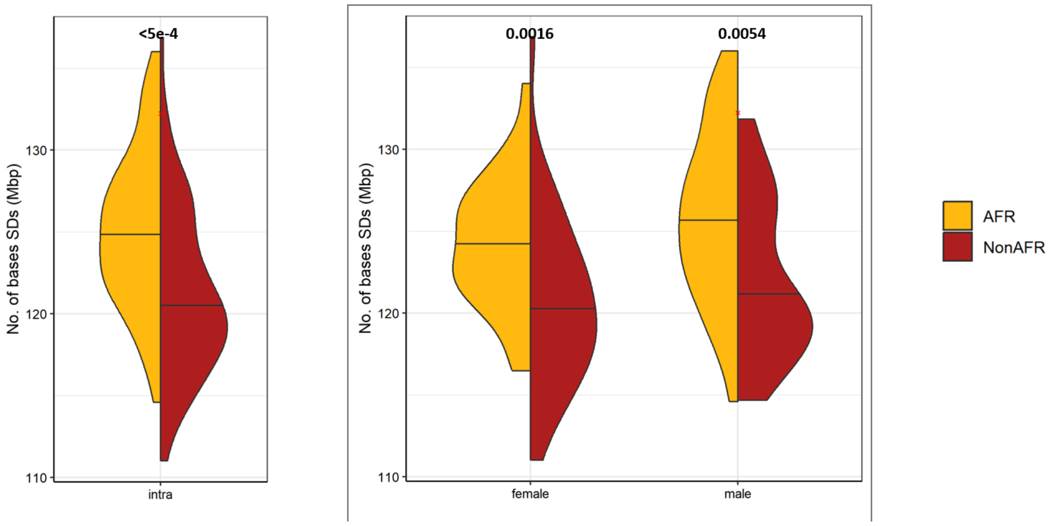


**Supplementary Figure 16. African vs. non-African SD content.**

Fig. 17

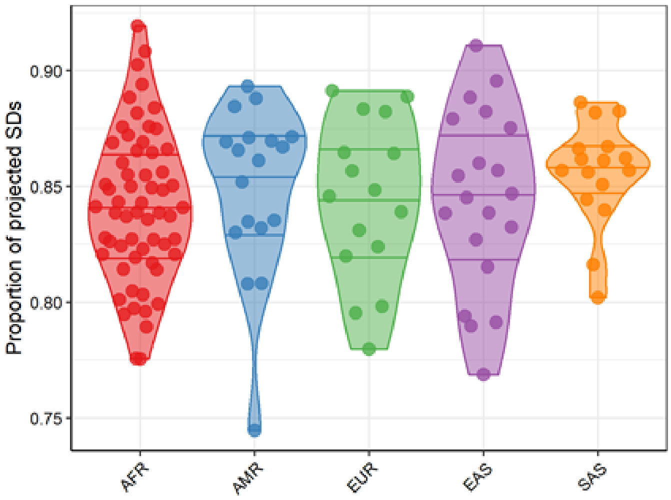


**Supplementary Figure 17. Proportion of segmental duplications successfully projected onto T2T space.**

FIg. 18

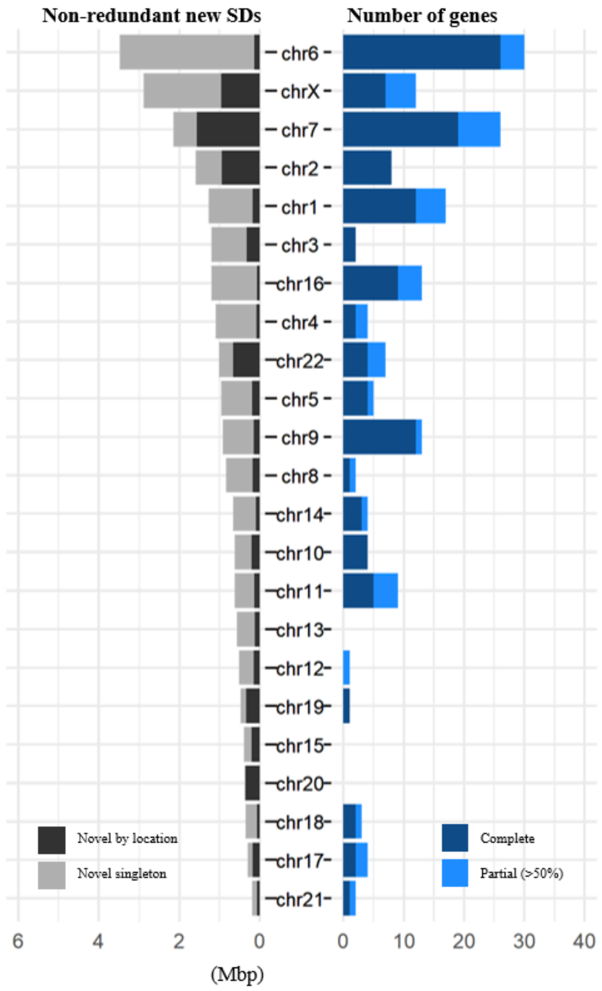


**Supplementary Figure 18. New SD content.** New segmental duplication content of HGSVC3 genomes compared to the segmental duplications of 170 haplotypes [71]. In the respective plots, the left shows SDs with content changes, expansion, and whether they are new (based on location/synteny), while flagging those that are singleton. On the right quantifies the number of complete (100% coverage), and partial overlaps (>50% coverage) with protein-coding genes.

Fig. 19

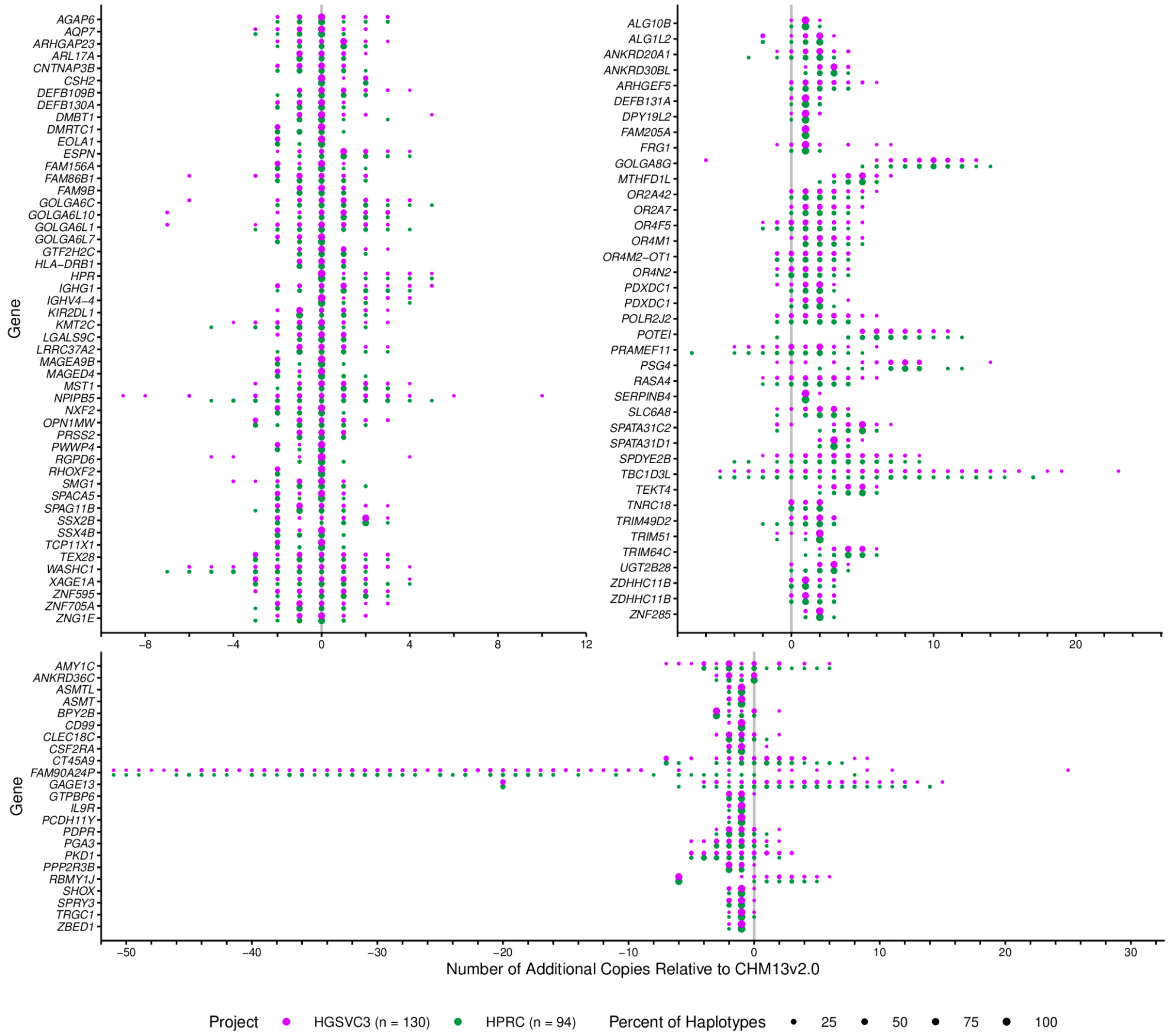


**Supplemental Figure 19. Top 50 CNV genes.** A comparison of the top-50 CNV genes in the HGSVC3 (violet) and HPRC (green) assemblies. (top-left) CNV genes with equal average copy number in HGSVC3 as CHM13. (top-right) CNV genes with greater average copy number in HGSVC3 versus CHM31. (bottom) Genes with lower average copy number than CHM13.

Fig. 20

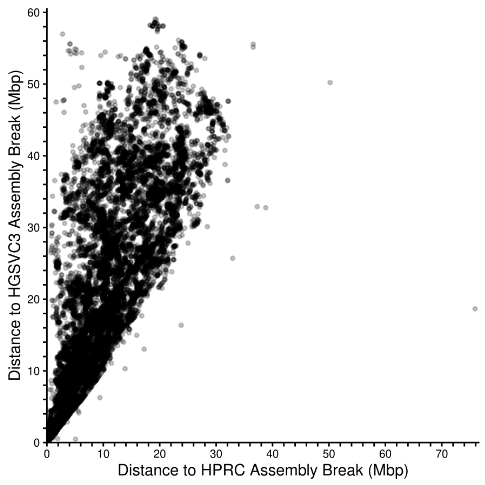


**Supplemental Figure 20. CNV gene contiguity.** Contiguity analysis of CNV genes in HGSVC3 and HPRC assemblies. Each point reflects a gene that is duplicated in both assemblies. The x and y coordinates reflect the closest distance to an assembly break defined by the end of a contig or an unassigned base (“N”). Points above the diagonal reflect greater contiguity in HGSVC3 assemblies.

Fig. 21

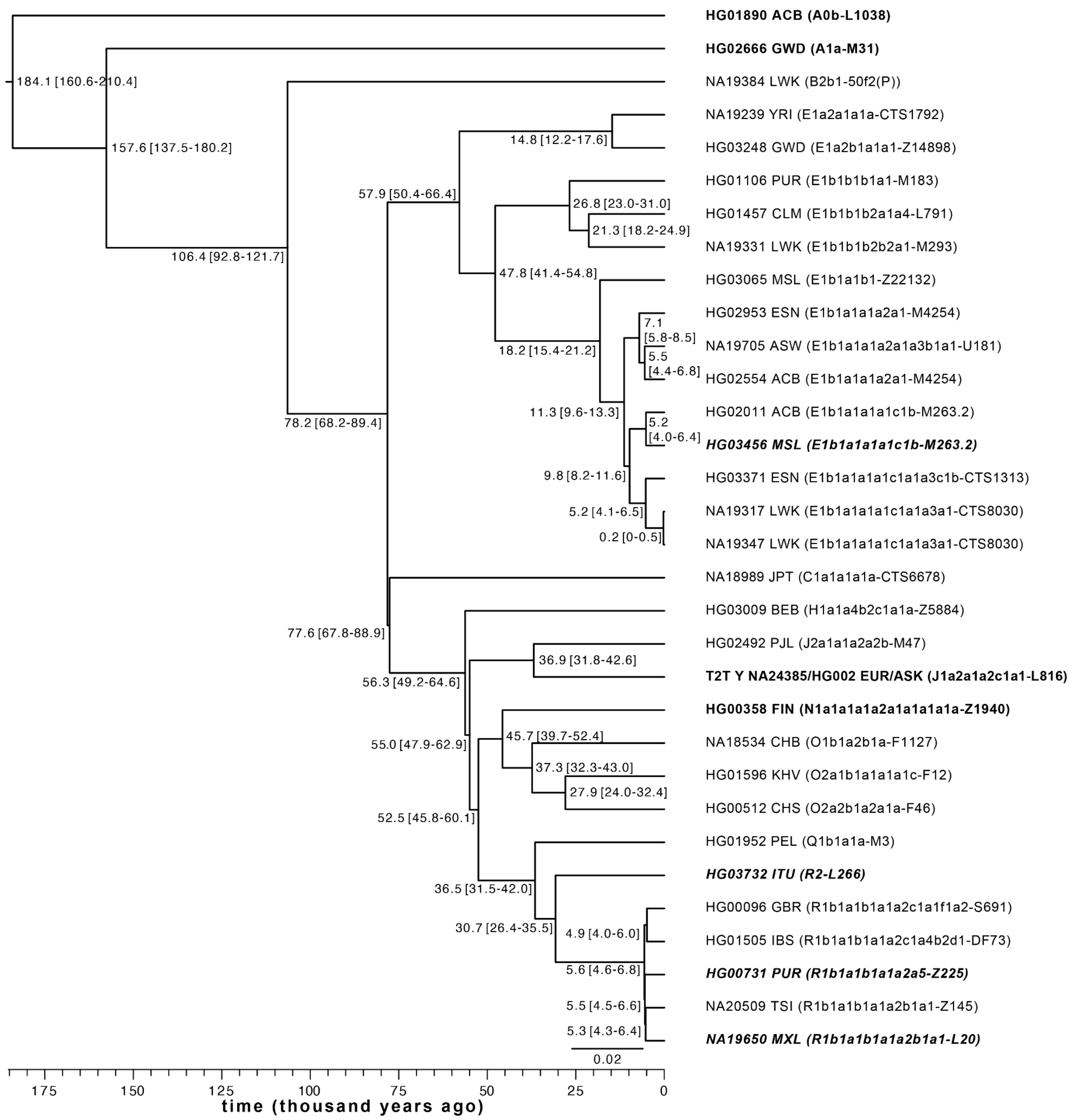


**Supplementary Figure 21. Phylogenetic relationships of 32 Y chromosomes.** Split times as estimated according to the BEAST analysis are shown with 95% HPD interval in brackets. Sample ID is followed by population designation, full Y haplogroup label according to ISOGG v15.73 and terminal marker ID. Population abbreviations: ACB - African Caribbean in Barbados; ASW - African Ancestry in SW USA; BEB - Bengali in Bangladesh; CHB - Han Chinese in Beijing, China; CHS - Han Chinese South; CLM - Colombian in Medellín, Colombia; ESN - Esan in Nigeria; FIN - Finnish in Finland; GBR - British From England and Scotland; GWD - Gambian in Western Division – Mandinka; IBS - Iberian Populations in Spain; ITU - Indian Telugu in the U.K.; JPT - Japanese in Tokyo, Japan; KHV - Kinh in Ho Chi Minh City, Vietnam; LWK - Luhya in Webuye, Kenya; MSL - Mende in Sierra Leone; MXL - Mexican Ancestry in Los Angeles CA USA; PEL - Peruvian in Lima Peru; PJL - Punjabi in Lahore, Pakistan; PUR - Puerto Rican in Puerto Rico; TSI - Toscani in Italia; YRI - Yoruba in Ibadan, Nigeria. Four novel contiguously assembled Y chromosomes (note - HG03456 has an assembly break in pseudoautosomal region 1) are shown in bold italics and previously reported contiguous Y chromosomes[8,39] in bold.

Fig. 22

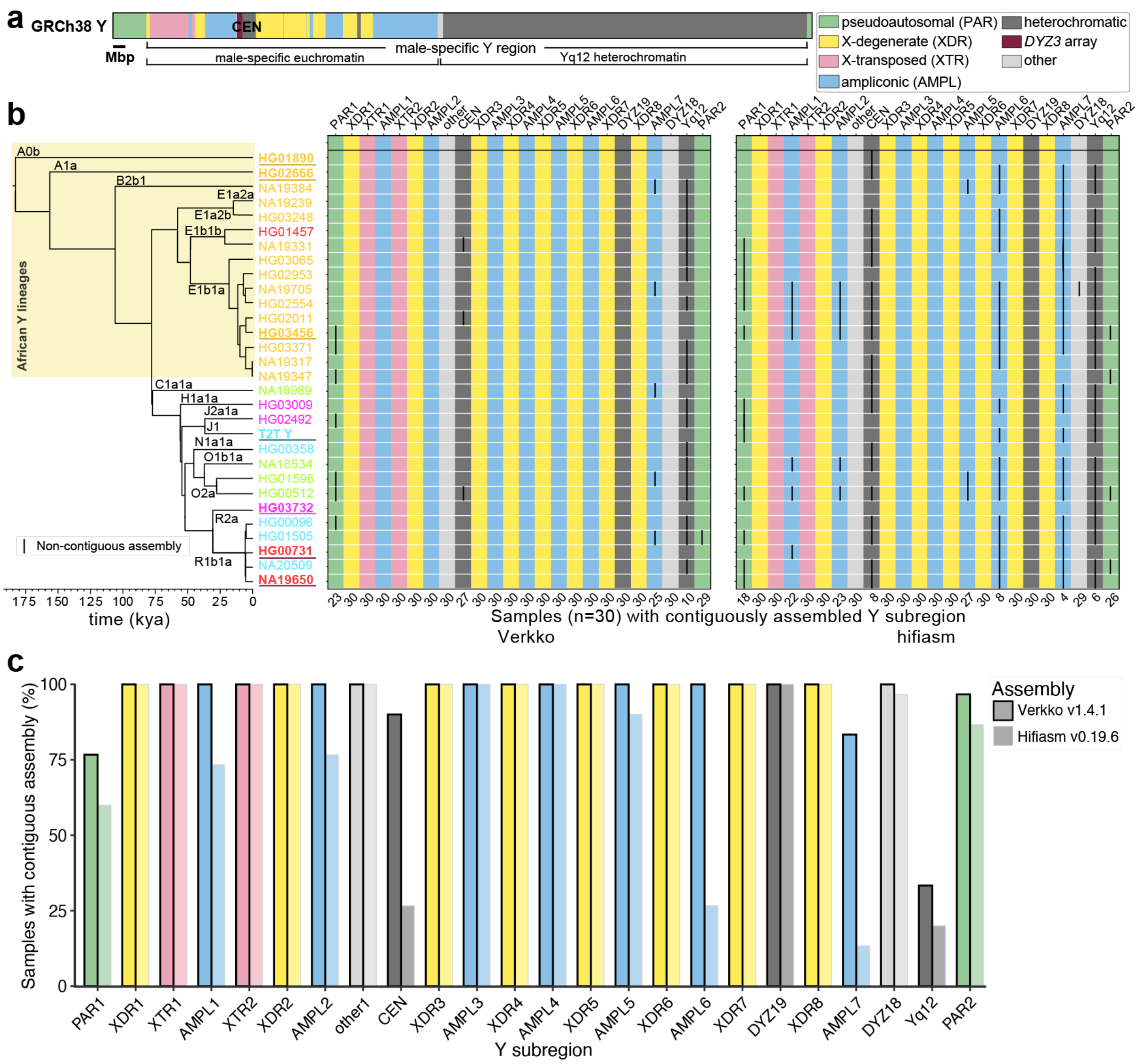


**Supplementary Figure 22. Assembly contiguity of the Y chromosomes**. **a.** Human Y chromosome structure based on the GRCh38 reference sequence. **b.** Dated Y phylogeny (left) and assembly completeness of Y-chromosomal subregions comparing Verkko and hifiasm (right). The color of sample ID corresponds to superpopulation designation (**Fig. 1a**). A Y-chromosomal subregion was considered to be contiguous if it was assembled contiguously from the subclass on the left to the subclass on the right. PAR1 and PAR2 regions were considered contiguously assembled if they contained the telomeric repeat sequence. kya - thousand years ago. **c.** The proportion of contiguously assembled Y subregions across 30 males comparing Verkko and hifiasm. See panel **a** for subregion labels.

Fig. 23

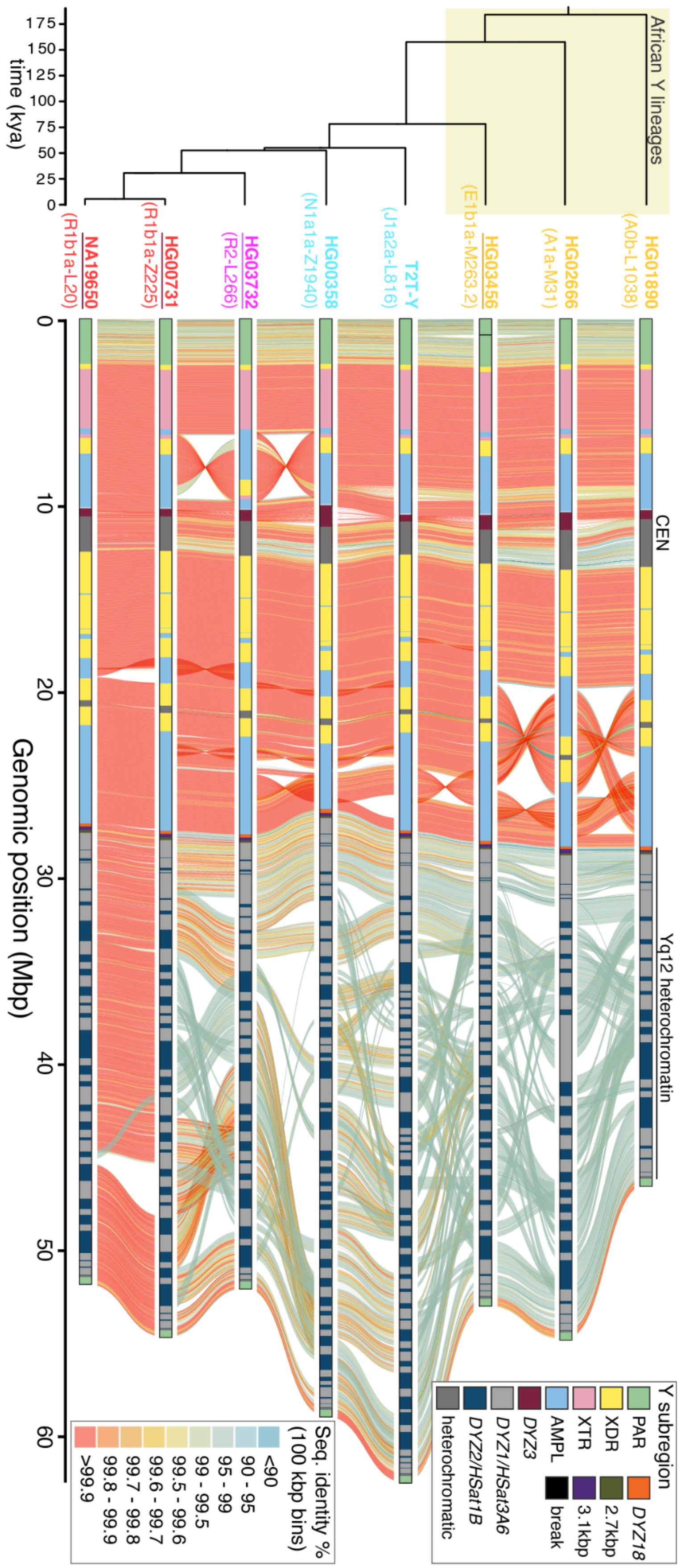

African Y lineages
time (kya)
175
150
125
100
75
50
25
0
NA19650
(R1b1a-L20)
HG00731
(R1b1a-Z225)
HG03732
(R2-L266)
HG00358
(N1a1a-Z1940)
T2T-Y
(J1a2a-L816)
HG03456
(E1b1a-M263.2)
HG02666
(A1a-M31)
HG01890
(A0b-L1038)
CEN
Yq12 heterochromatin
Genomic position (Mbp)
0
10
20
30
40
50
60
Seq. identity %
(100 kbp bins)
<90
90 - 95
95 - 99
99 - 99.5
99.5 - 99.6
99.6 - 99.7
99.7 - 99.8
99.8 - 99.9
>99.9
Y subregion
PAR
XDR
XTR
AMPL
DYZ3
DYZ1/HSat3A6
DYZ2/HSat1B
heterochromatic
DYZ18
2.7kbp
3.1kbp
break

**Supplementary Figure 23. Pairwise comparison of the eight contiguously assembled Y chromosomes.** Four novel contiguous Y assemblies, representing E1b1a, R2a and R1b1a Y lineages (three assembled telomere-to-telomere and one, HG03456, with a break in PAR1 region) are underlined. Six chrY assemblies were generated here using Verkko (v1.4.1) and two (T2T Y and HG00358) were included from [8,39].

Fig. 24

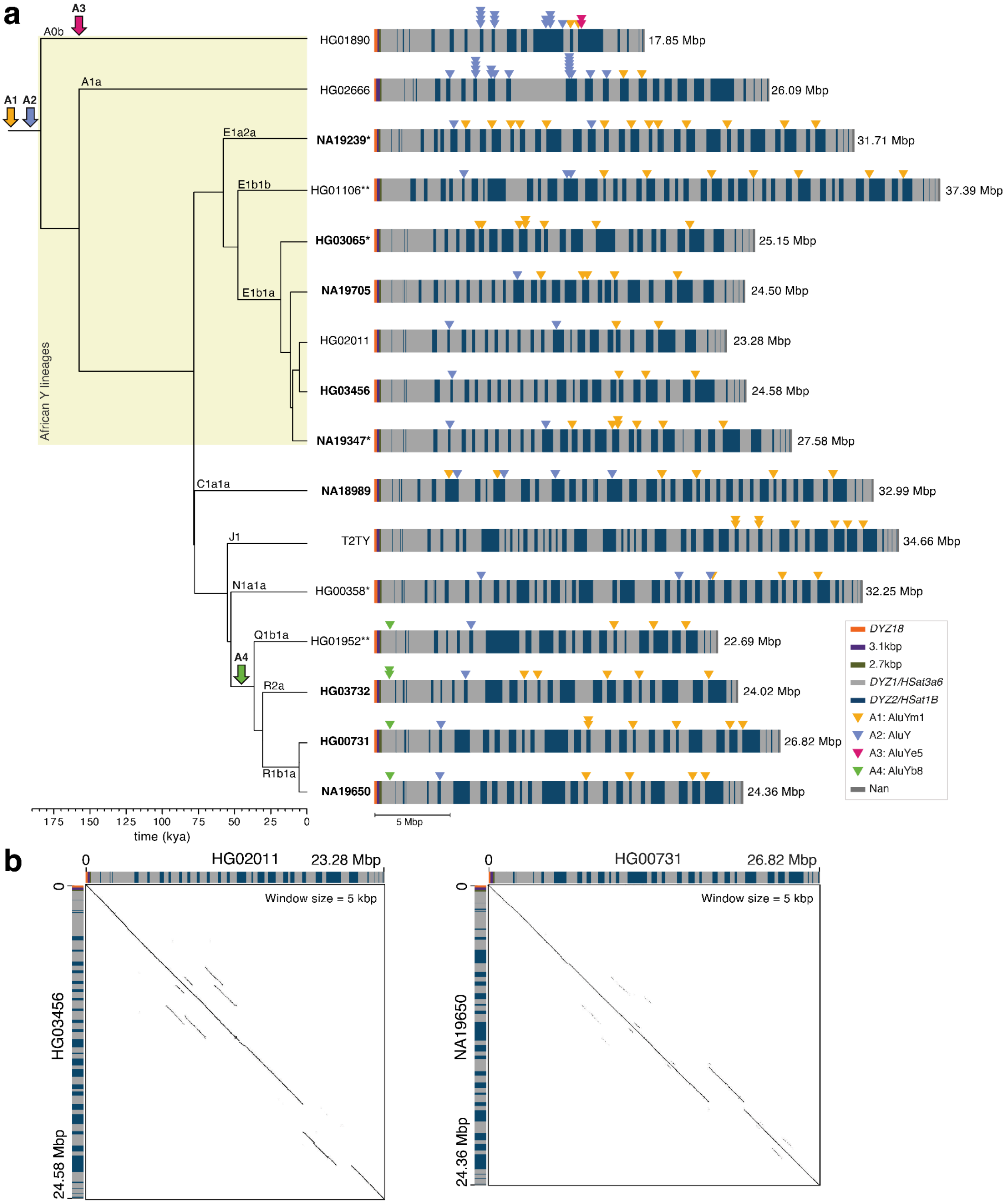


**Supplementary Figure 24. Yq12 heterochromatic region. a)** Repeat composition of the contiguously assembled Yq12 heterochromatic regions across 16 samples. Distribution of the four reported[8] *Alu* insertions across the Yq12 (labeled A1-A4) are shown as filled triangles. Based on their phylogenetic distribution, the insertion of *Alu* elements A1 and A2 occurred prior to the split of studied Y chromosomes, while A3 insertion is present only in the A0b lineage and A4 shared occurred in the common ancestor of Q and R lineages. Samples included were assembled using Verkko (this study, n=9), hifiasm (this study, n=4, indicated by an asterisk). Yq12 sequences for two samples (indicated by double asterisks) were included from Ref. [8] and the T2T Y chromosome sequence from Ref. [39]. Novel, previously unpublished Yq12 sequences (n=9) are shown in bold text. The total length of the Yq12 region is indicated on the right. ka, thousand years ago. **b)** Dotplots of the Yq12 regions of two closely related chrY pairs (haplogroup E1b1a: HG02011 and HG03456 and R1b1a: HG00731 and NA19650) using window size = 5,000 bp and showing evidence of two (left) and one (right) large insertion/deletion events.

Fig. 25

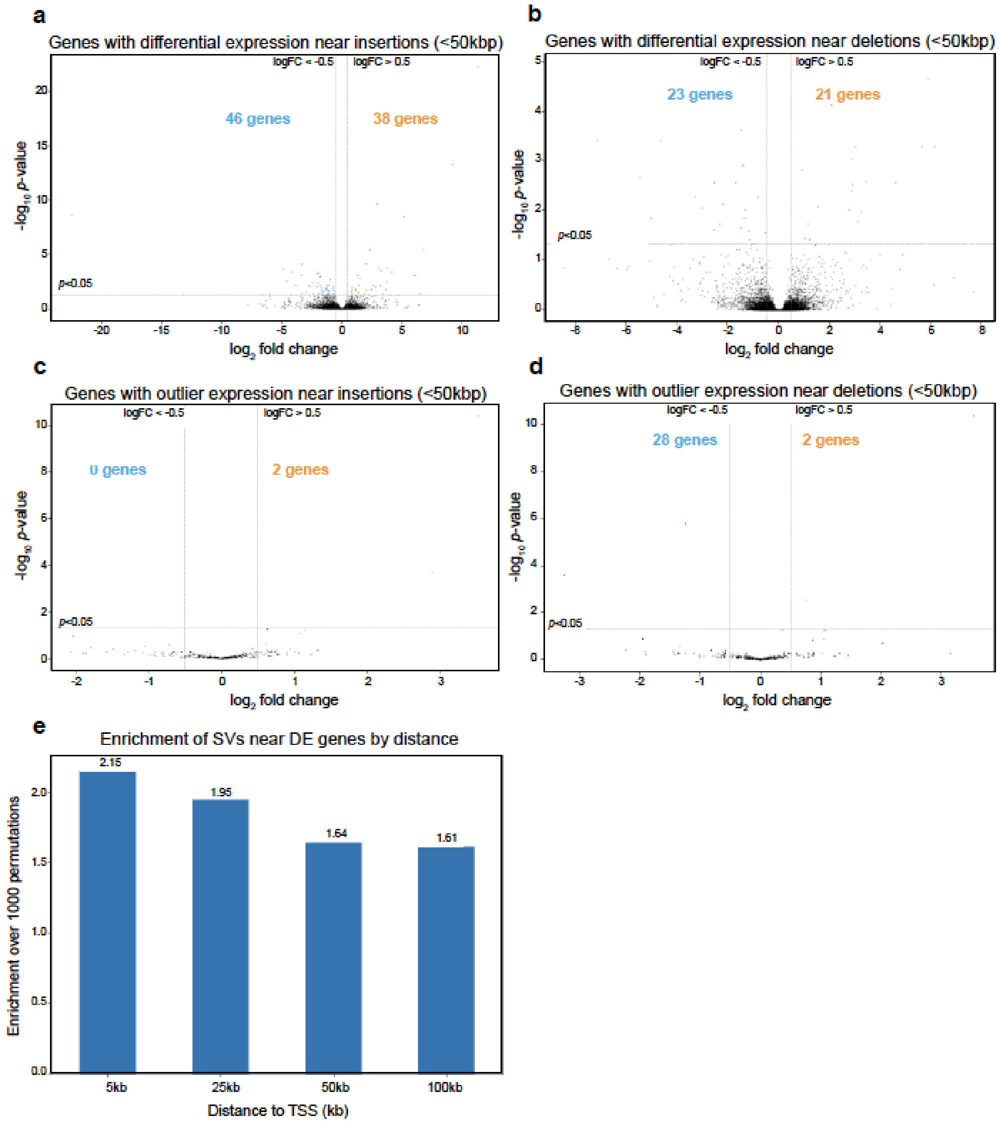


**Supplementary Figure 25. Differential expression analysis. a,b)** Volcano plots of differentially expressed genes with (a) insertions and (b) deletions located <50 kbp from the gene TSS without overlapping the gene. Differential expression was calculated by comparing groups of 2-10 individuals who carried the SV vs. the remainder of individuals without the SV. The x-axis is the $\log_2$-fold-change of read counts, and the y-axis is $-\log_{10}$ p-values (adjusted with Benjamini-Hochberg correction). The significant expression increases and decreases are defined as points with adjusted p-value < 0.05 and $\log_2$-fold-change > 0.5 or < -0.5. The numbers indicate counts of SV-gene pairs considered significant. **c,d)** Volcano plots of outlier expression genes with (c) insertions and (d) deletions located <50 kbp from the gene TSS without overlapping the gene. Outlier expression was calculated by comparing individual samples with or without an SV vs. the remaining 11 samples, with P-values derived from Z-scores (see Methods section “Differential gene expression and outliers” above). **e)** Enrichment of proportions of genes with differential expression occurring near SVs per genomic distance, compared with the proportion of genes near SVs averaged across 1000 permutation tests. The decreasing trend indicates that SVs closer to TSSs are more likely to affect expression.

Fig. 26

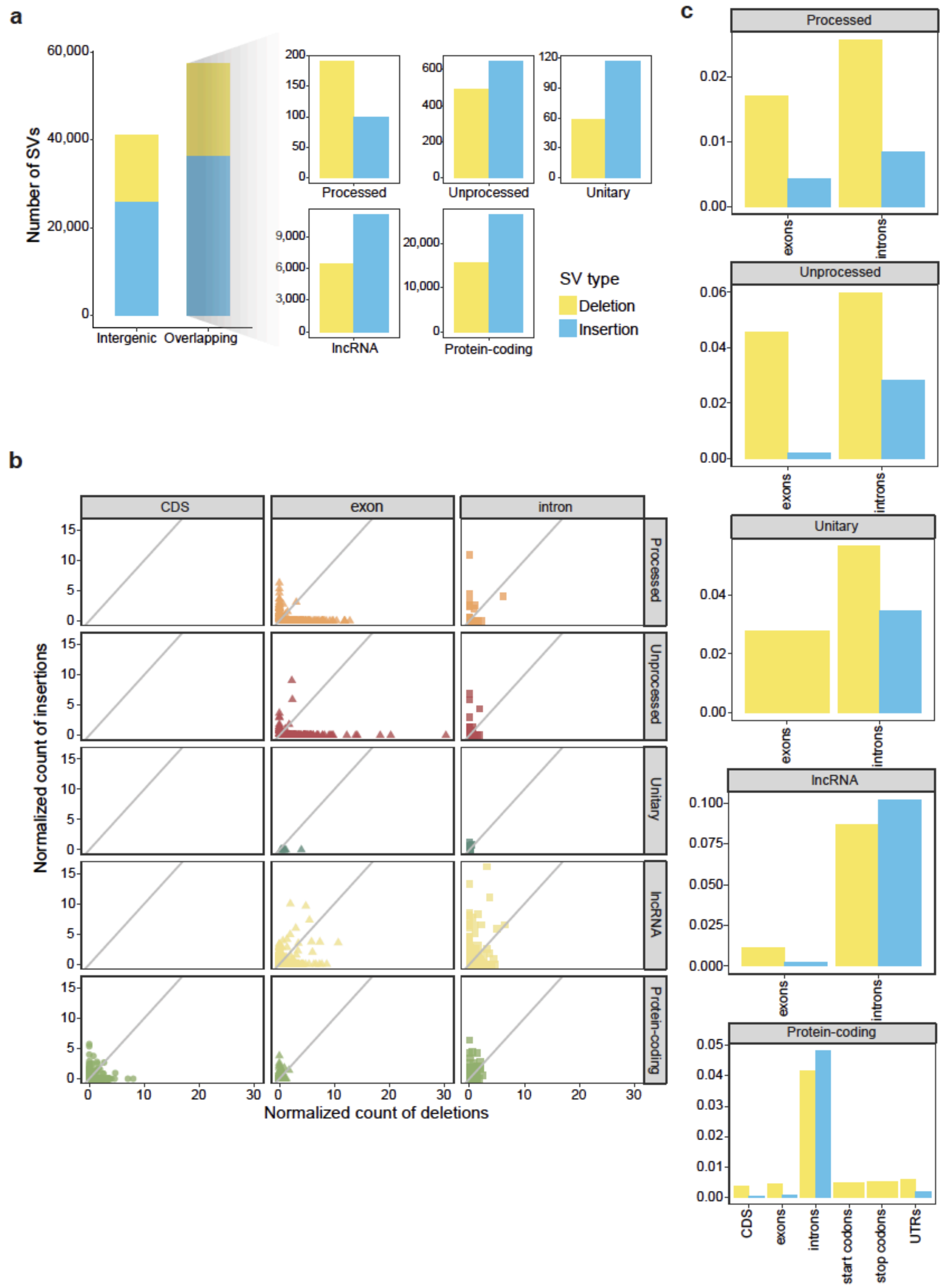


**Supplementary Figure 26. SV intersects with functional elements. a)** Number of intergenic and overlapping SVs. Over 40% of SVs in the human genome were intergenic, indicating that these SVs did not overlap with annotated protein-coding genes, long noncoding RNAs (lncRNAs), or pseudogenes (including processed, unprocessed, and unitary types). For the SVs that did overlap with genes, we categorized the genes into different biotypes based on GENCODE v45 annotations. Insertions outnumbered deletions across all gene biotypes, except for processed pseudogenes. **b)** Normalized counts of deletions and insertions. The X-axis represents the number of deletions, while the Y-axis represents the number of insertions. Number of SV overlaps were counted per gene and per genomic element, and we normalized the counts by dividing them by the length of the genomic elements for a given gene and multiplying by a scaling factor of 1,000. Overall, pseudogenes tended to have more deletions within both exons and introns, while protein-coding genes exhibited a balance between insertions and deletions within coding sequences (CDSs), exons, and introns. **c)** Proportion of different genomic elements affected by SVs. At the transcript level, we calculated the overlapping length of SVs per genomic or transcript element (i.e., intron or exon). The proportion was defined by dividing the overlapping length by the total length in the genome. As expected, a higher proportion was consistently observed for both insertions and deletions within introns compared to exons across different transcript biotypes. Notably, protein-coding transcripts exhibited a significantly reduced proportion for insertions and deletions within CDSs, exons, start codons, stop codons, and untranslated regions (UTRs) compared to introns.

Fig. 27

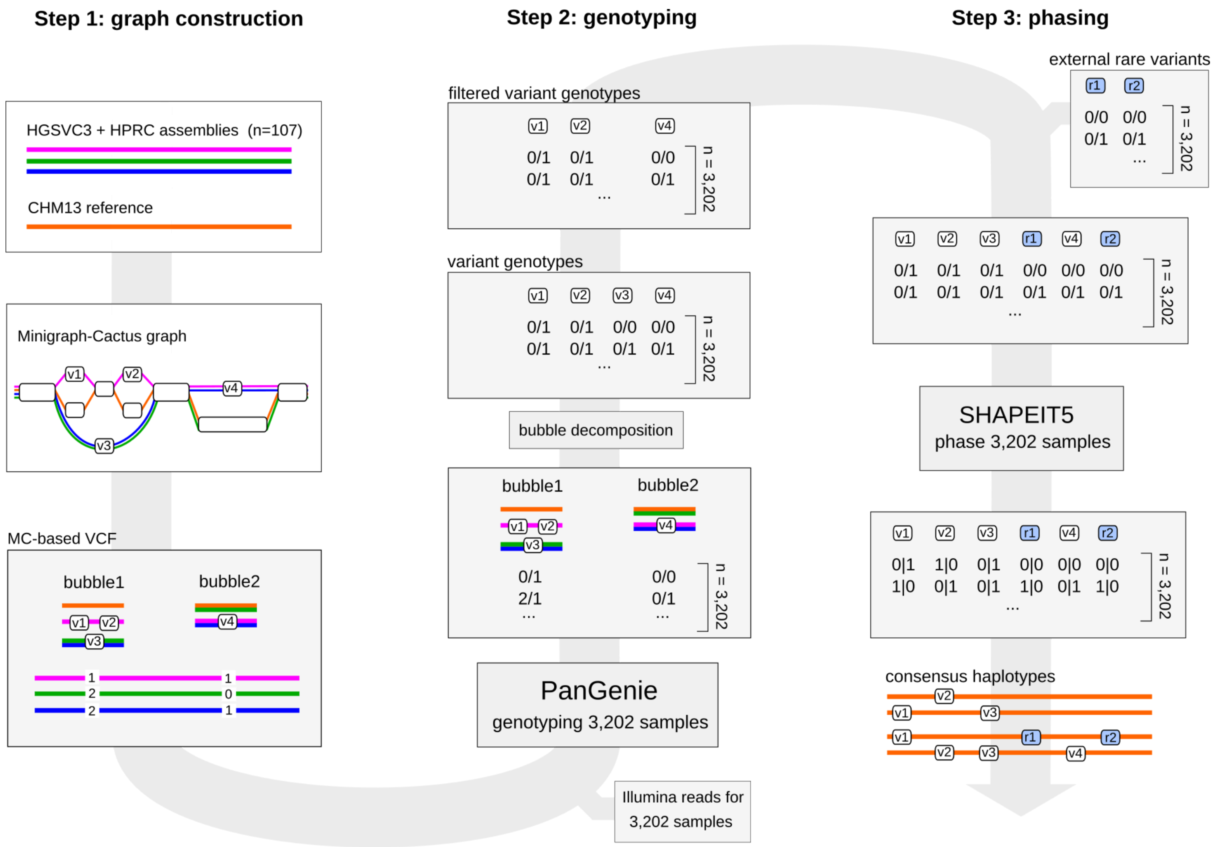


**Supplementary Figure 27. Overview of the graph constructing, genotyping and phasing pipelines. Step 1:** graph construction. A pangenome graph is constructed from haplotype-resolved assemblies of 107 samples (65 HGSVC3, 42 HPRC). A VCF representation of the bubbles with phased genotypes for all assembly samples is generated. **Step 2:** genotyping. The bubbles are genotyped across all 3,202 1kGP samples with PanGenie using the panel VCF generated in step 1 as input, as well as Illumina reads for all samples. Bubbles are decomposed into variant alleles and bubble genotypes are translated to variant genotypes. Finally, genotypes are filtered using a machine-learning approach. Step 3: phasing. The filtered genotypes are combined with additional rare variant calls produced from short reads across all 3,202 samples and the combined set is phased with SHAPEIT5. Based on the resulting phased genotypes, consensus haplotypes are generated for all 3,202 samples by implanting variants located on each haplotype into the reference genome.

Fig. 28

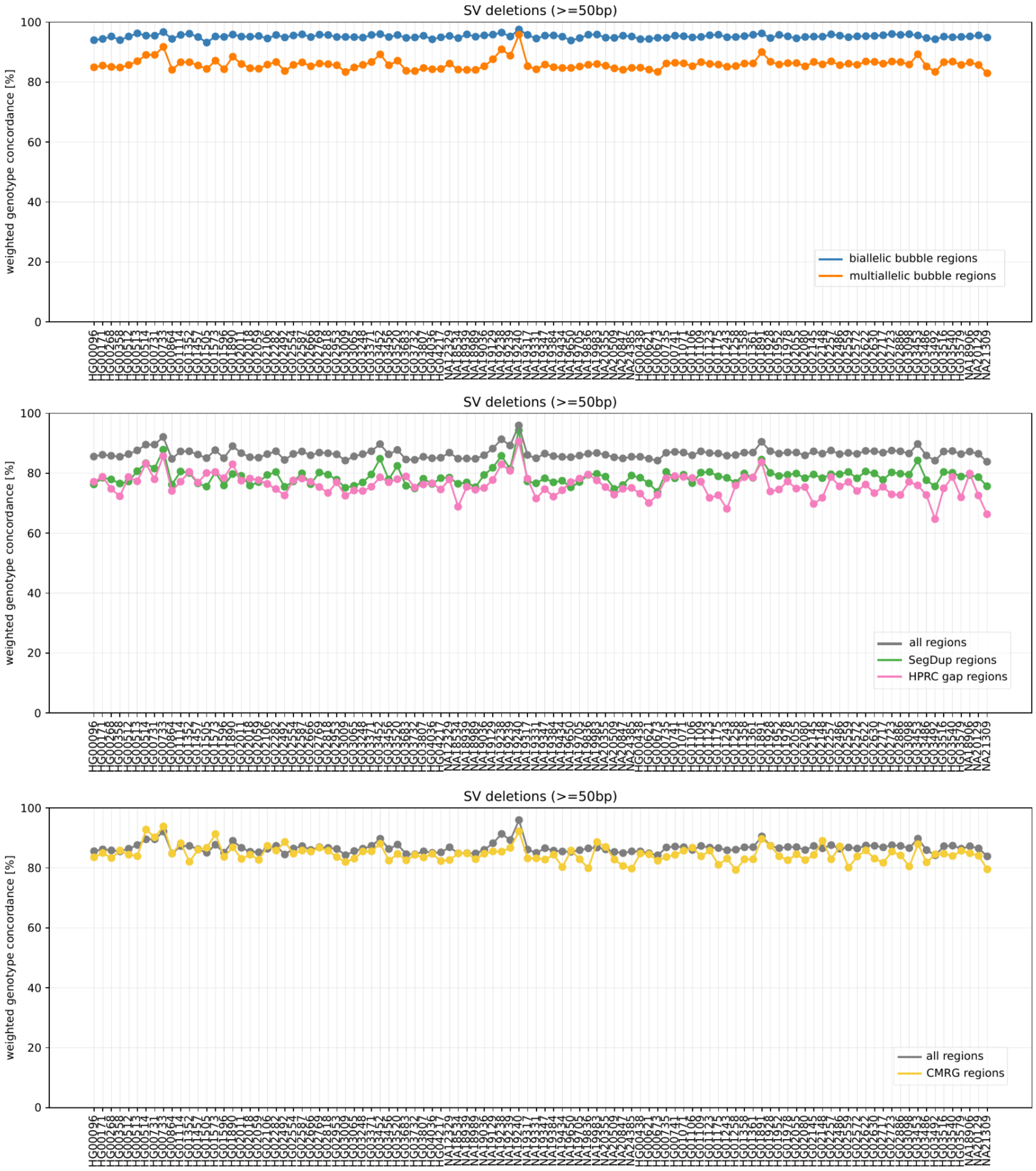


**Supplementary Figure 28. PanGenie Leave-one-out validation for deletions.** We conducted a leave-one-out experiment by repeatedly removing one sample from the MC-based panel VCF and genotyping it with PanGenie. The left out sample was used as a ground truth for evaluation based on the weighted genotype concordance[127]. The plots show concordances of variants falling within certain regions: all biallelic bubble regions of the graph (blue), multiallelic bubble regions (orange), segmental duplications (green), regions of gaps in the HPRC assemblies[28] (pink) and within challenging medically relevant genes[129] (yellow).

### Fig. 29

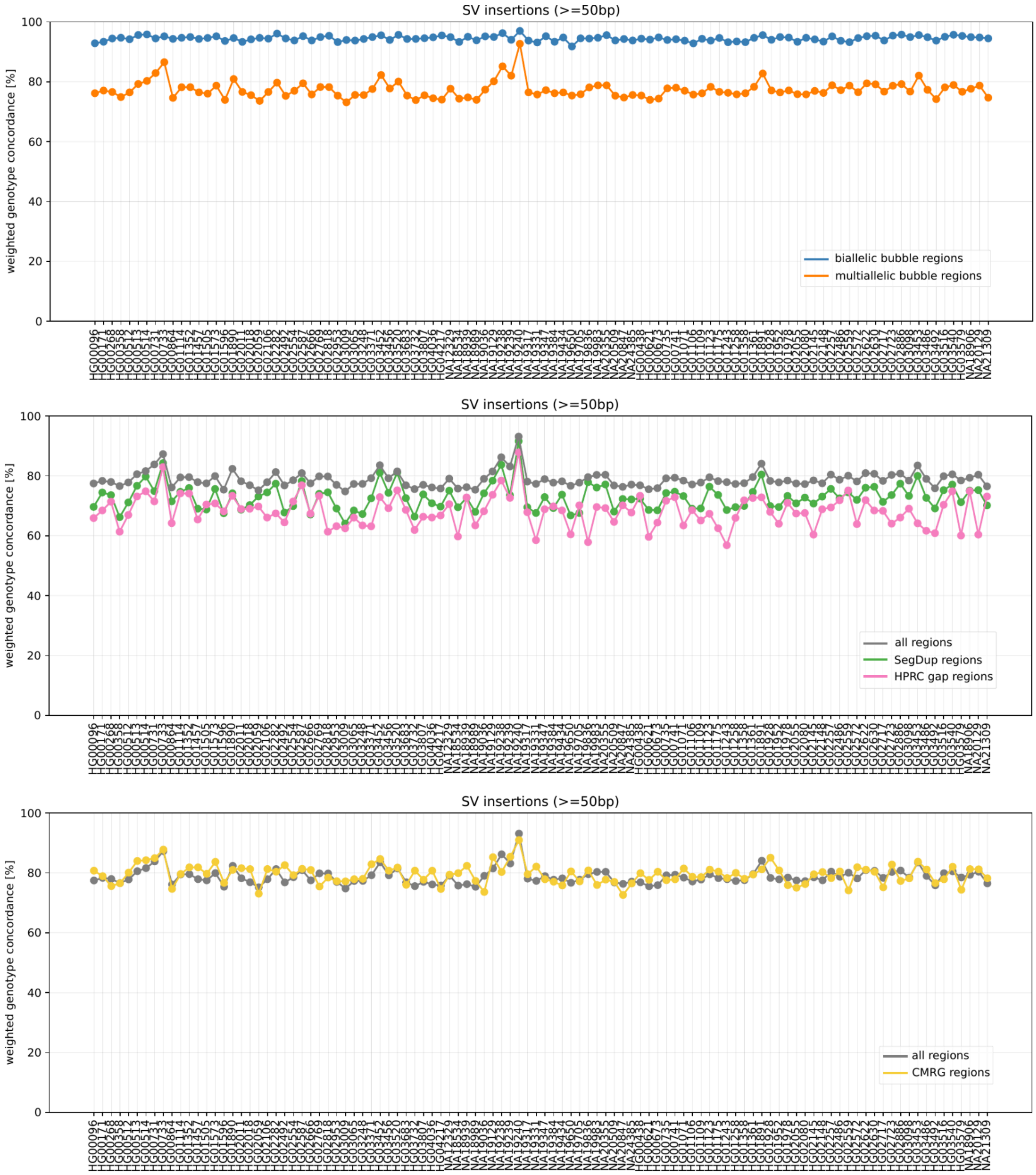


**Supplementary Figure 29. PanGenie Leave-one-out validation for insertions.** We conducted a leave-one-out experiment by repeatedly removing one sample from the MC-based panel VCF and genotyping it with PanGenie. The left out sample was used as a ground truth for evaluation based on the weighted genotype concordance[127]. The plots show concordances of variants falling within certain regions: all biallelic bubble regions of the graph (blue), multiallelic bubble regions (orange), segmental duplications (green), regions of gaps in the HPRC assemblies[28] (pink) and within challenging medically relevant genes[129] (yellow).

### Fig. 30

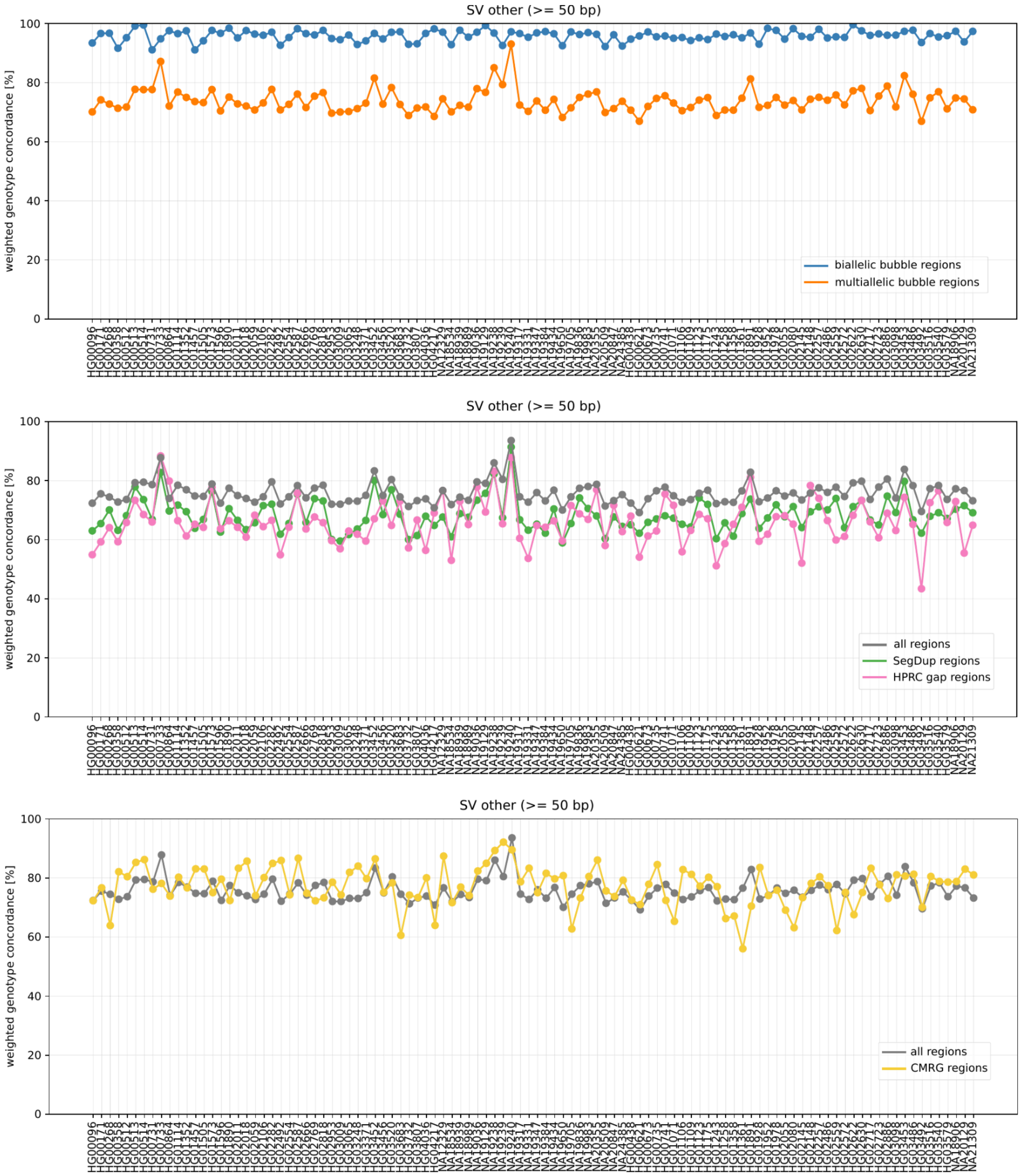


**Supplementary Figure 30. PanGenie Leave-one-out validation for other variants.** We conducted a leave-one-out experiment by repeatedly removing one sample from the MC-based panel VCF and genotyping it with PanGenie. The left out sample was used as a ground truth for evaluation based on the weighted genotype concordance[127]. The plots show concordances of variants falling within certain regions: all biallelic bubble regions of the graph (blue), multiallelic bubble regions (orange), segmental duplications (green), regions of gaps in the HPRC assemblies[28] (pink) and within challenging medically relevant genes[129] (yellow).

#### Fig. 31

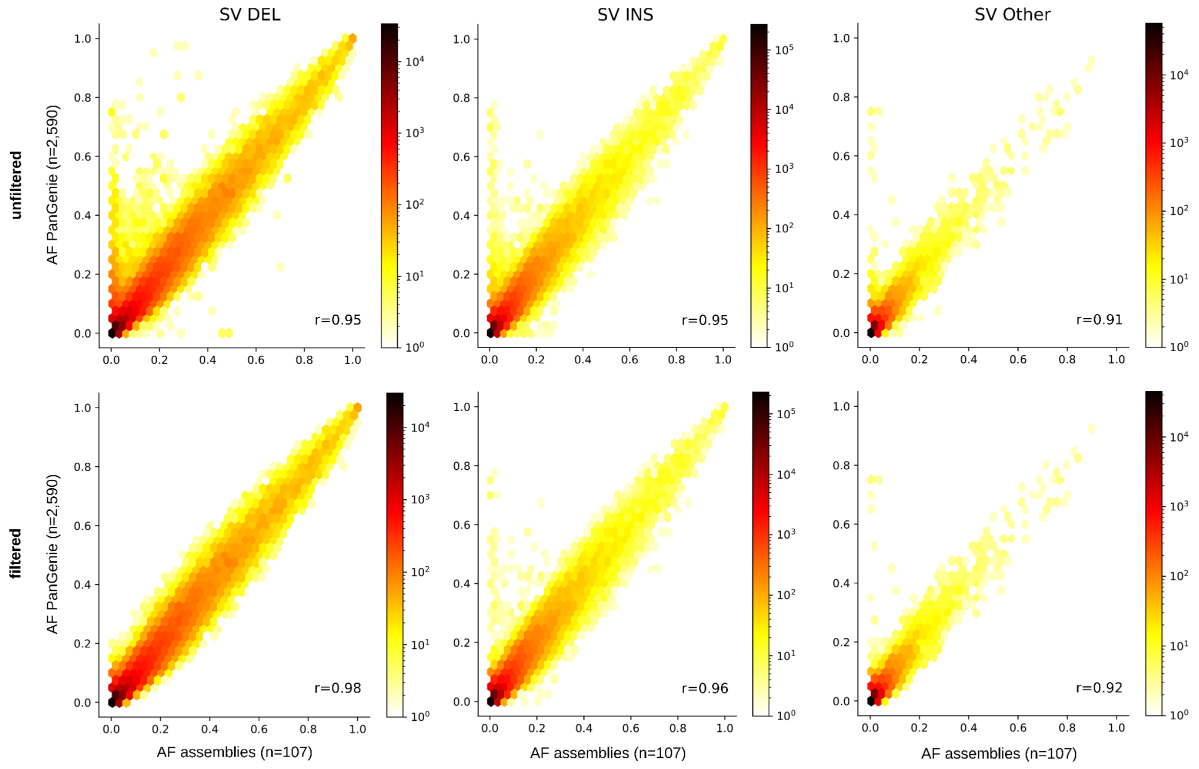


**Supplementary Figure 31. Comparison of allele frequencies by discovery and genotyping.** Shown is a comparison of allele frequencies for SV alleles across the assembly samples in the panel VCF (n=107) and corresponding allele frequencies after genotyping these variants across n=2,590 unrelated 1kGP samples.

Fig. 32

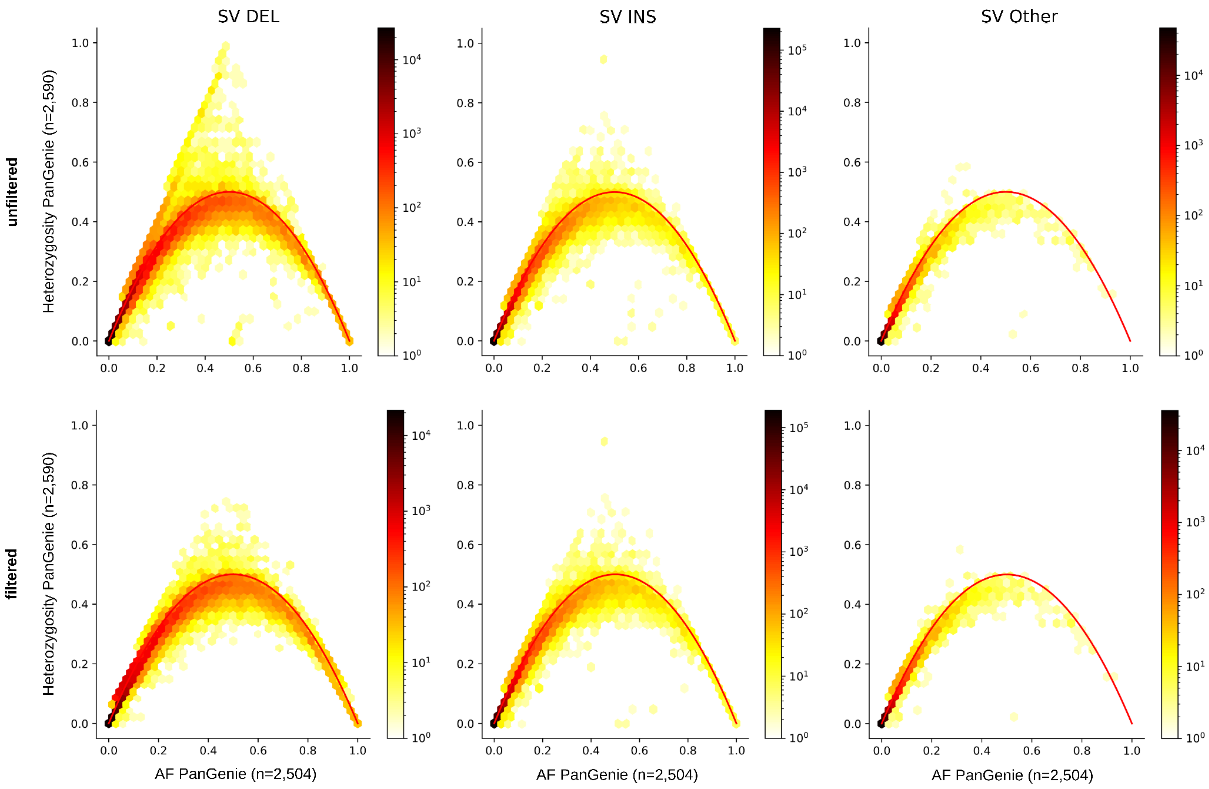


**Supplementary Figure 32. Hardy-Weinberg Equilibrium for PanGenie genotypes.** Shown is a comparison of heterozygosities and allele frequencies after genotyping n=2,590 unrelated 1kGP samples.

Fig. 33

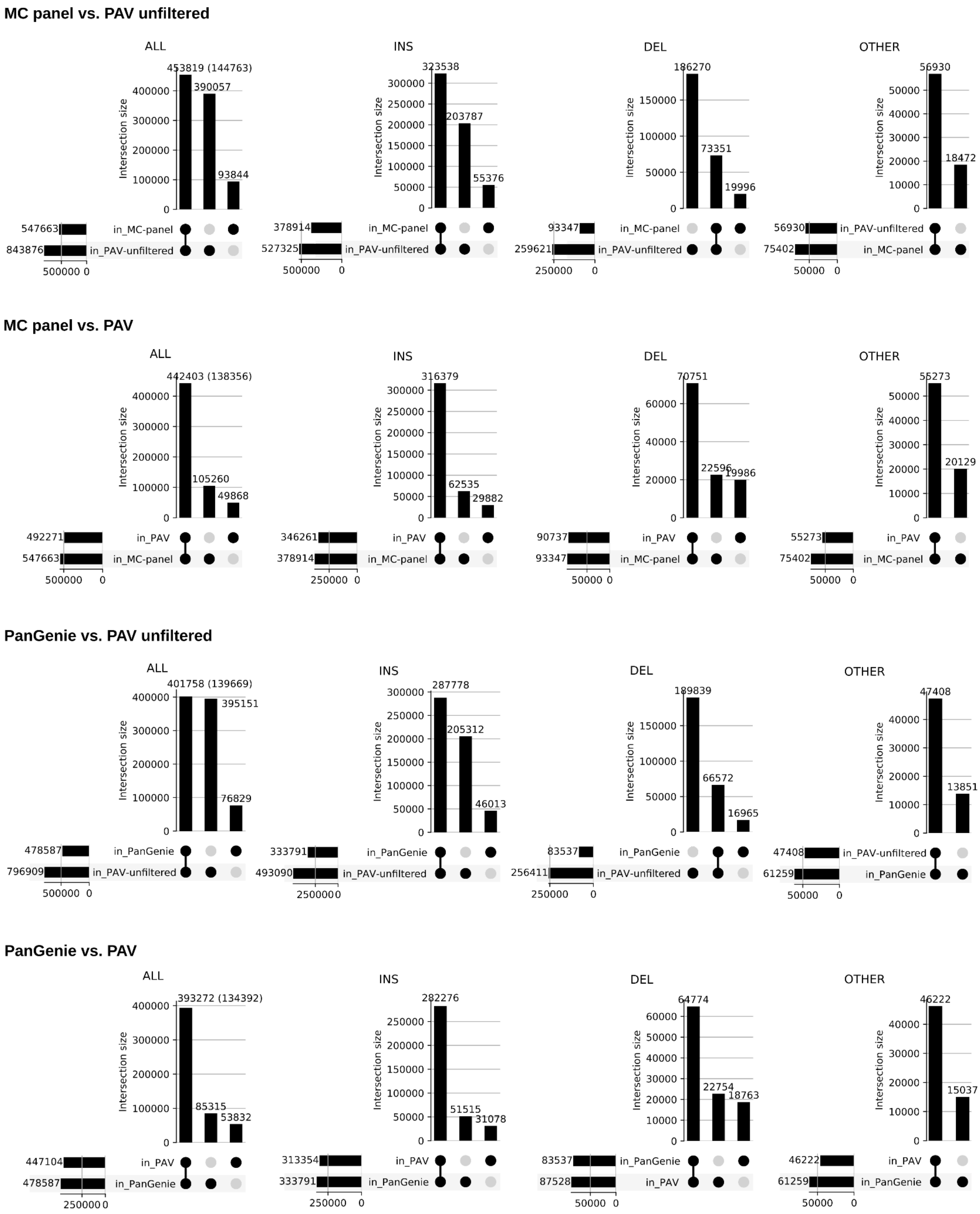


**Supplementary Figure 33. PAV vs graph genotypes.** Comparison of SVs in Minigraph-Cactus panel VCF (“MC-panel”), PanGenie filtered genotypes (“PanGenie”), PAV unfiltered (“PAV-unfiltered”) and PAV (“PAV”) callsets. Variant types and counts in intersections correspond to the MC / PanGenie sets; the numbers in brackets give the number of unique PAV calls matched (only for “ALL”).

Fig. 34

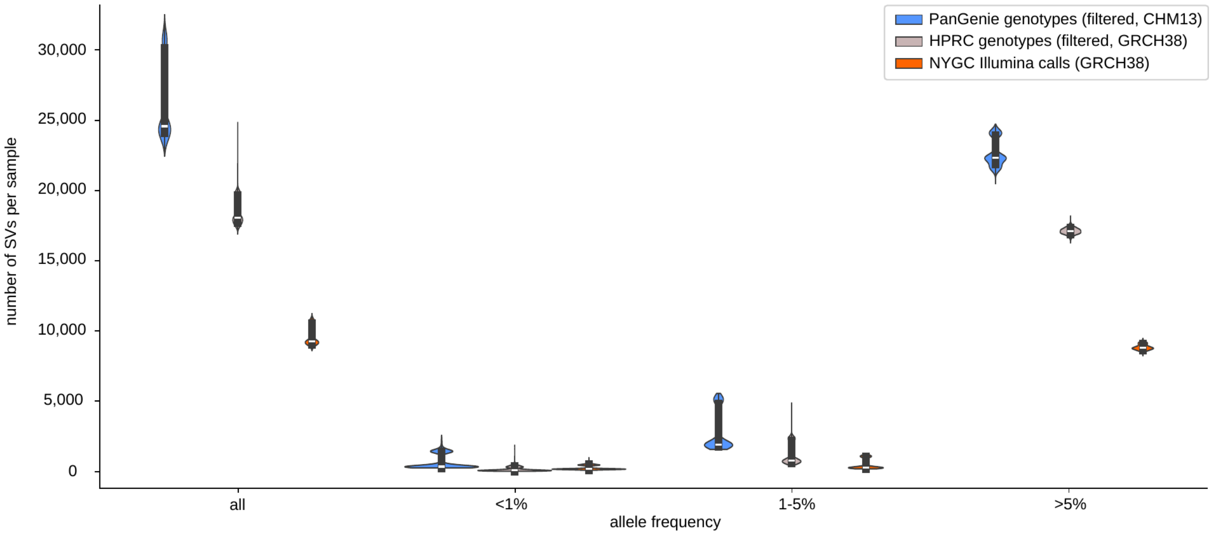


**Supplementary Figure 34. Comparison of PanGenie genotype frequencies.** We show the number of SV alleles present (0/1 or 1/1) in each sample (n=3,202) of each callset. Compared were our filtered genotyped set (blue), the HPRC genotyped set (grey) and the Illumina-based NYGC SV callset (orange). The boxes inside the violins represent the first and third quartiles of the data, white dots represent the medians, and black lines mark minima and maxima of the data.

Fig. 35

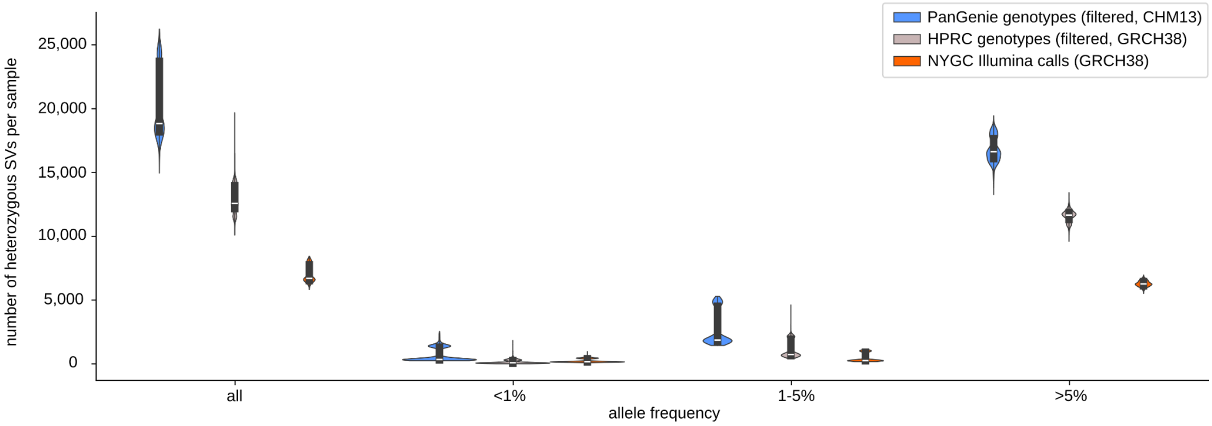


**Supplementary Figure 35**. **Comparison of PanGenie frequencies for heterozygous genotypes.** We show the number of heterozygous SV alleles present (genotype 0/1) in each sample (n=3,202) of each callset. Compared were our filtered genotyped set (blue), the HPRC genotyped set (grey) and the Illumina-based NYGC SV callset (orange). The boxes inside the violins represent the first and third quartiles of the data, white dots represent the medians, and black lines mark minima and maxima of the data.

### Fig. 36

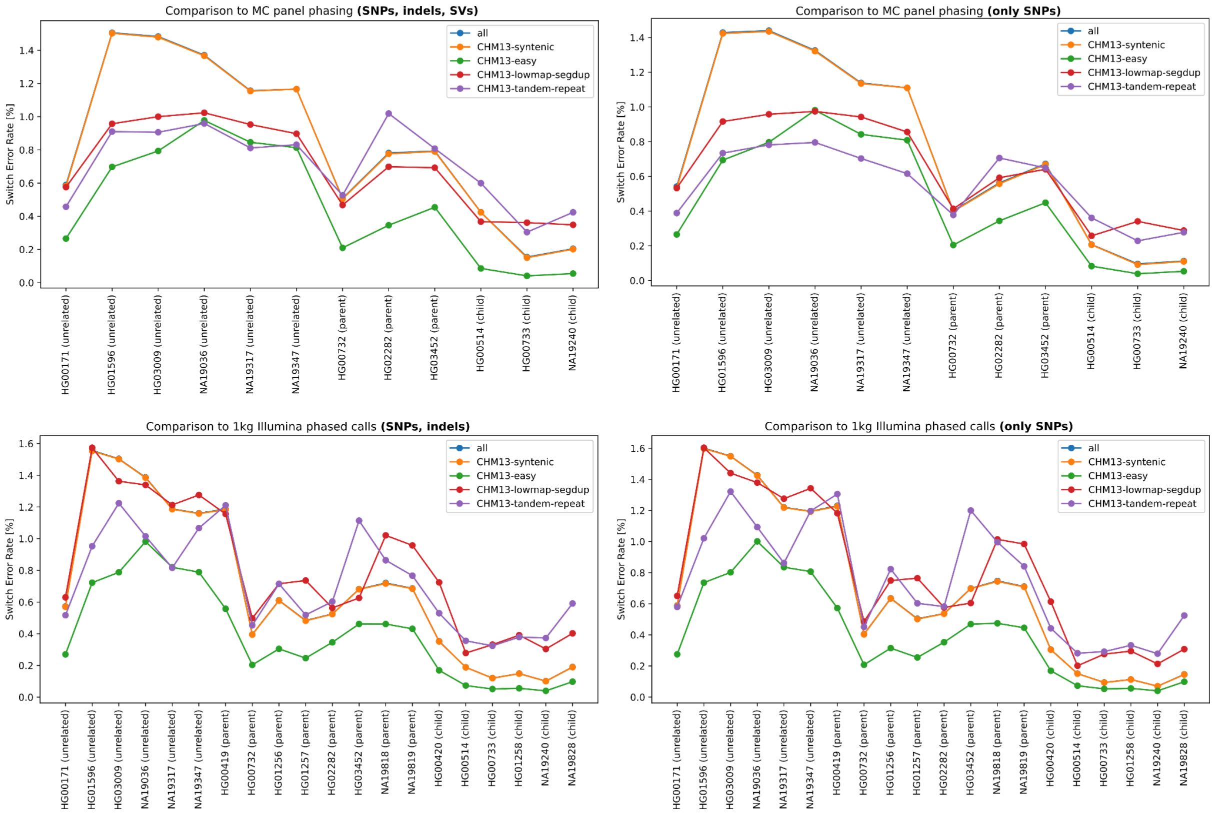


**Supplementary Figure 36. Genotype switch error rate.** Comparison of our phased genotypes to the phasing of the assembly samples in the MC VCF used as input for genotyping (top), as well as to a CHM13-based phased set of Illumina-based SNP and indel calls (bottom). The plots show switch error rates computed for all regions, as well as inside of CHM13-syntenic regions (regions shared between GRCh38 and CHM13), inside of low mappability and segmental duplication regions (“CHM13-lowmap-segdup”), tandem repeats (“CHM13-tandem-repeat”) and “easy” regions (“CHM13-easy”), i.e., regions outside of other difficult regions such as tandem repeats, homopolymers, difficult to map regions, segmental duplications and high/low GC content.

Fig. 37

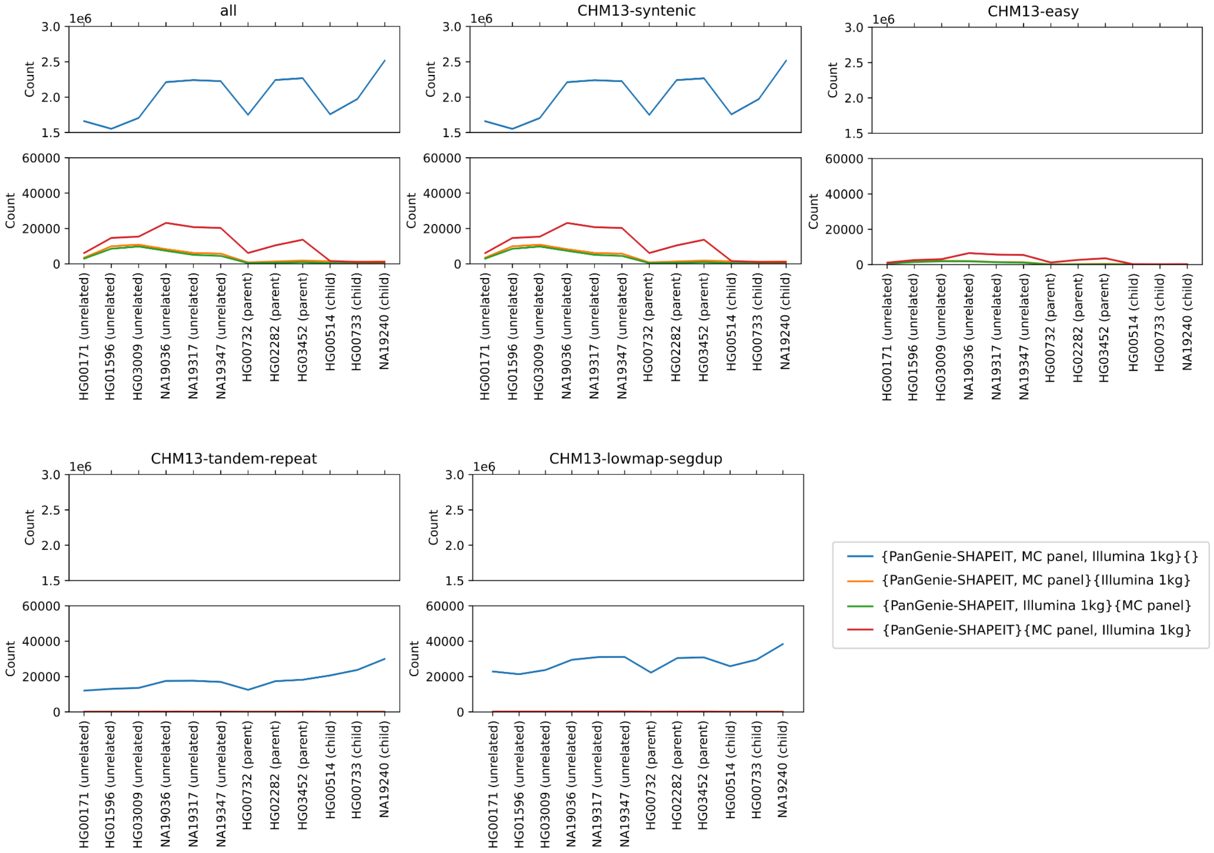


**Supplementary Figure 37. Three-way comparison of phased callsets.** Multiway comparison of three phased sets (all CHM13-based): our phased genotypes ("PanGenie-SHAPEIT"), the 1kGP Illumina-based phased calls and the MC panel VCF (input used for genotyping). The plots report the number of times different sets agreed / disagreed on the phasing of variant sites, i.e., the yellow line shows the number of cases in which our phasing and the MC panel phasing agreed, but the 1kGP phasing was different.

### Fig. 38

a

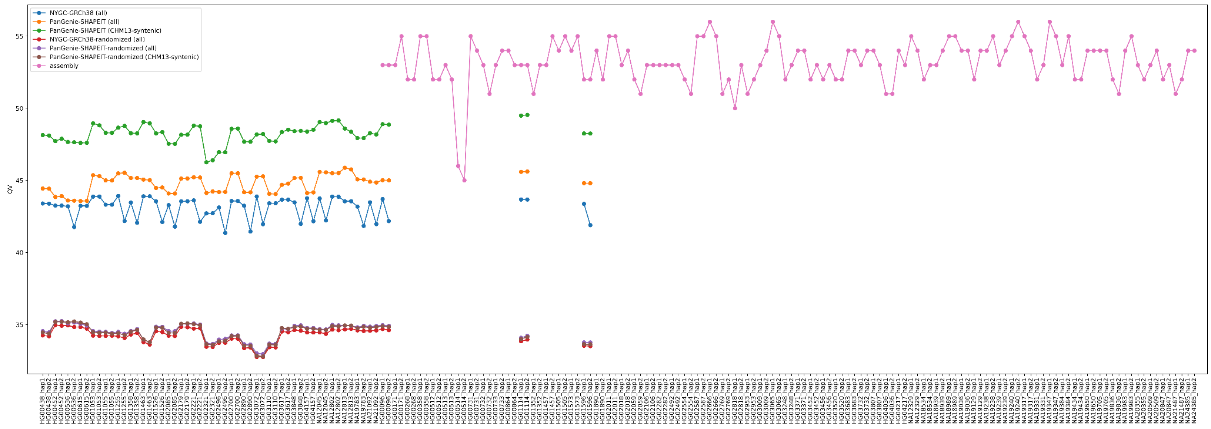


b

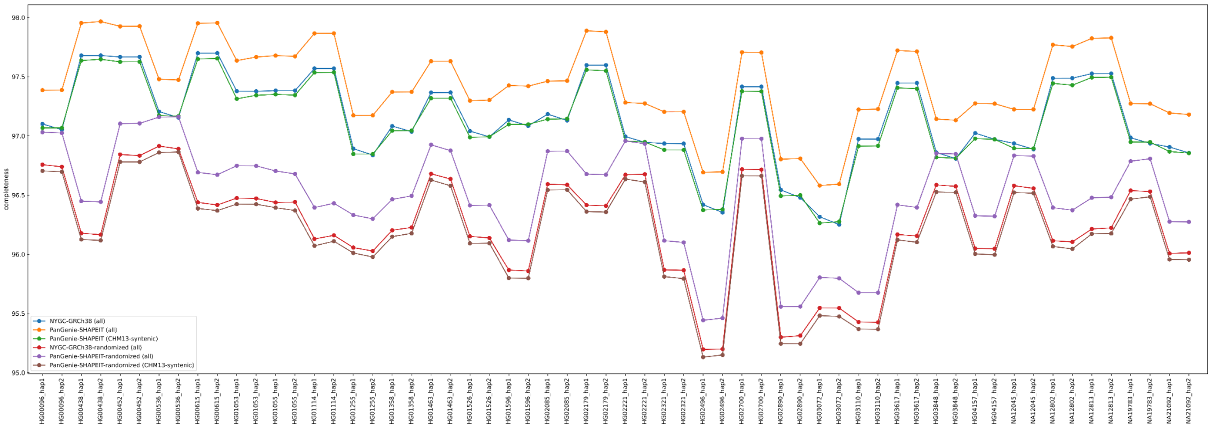


**Supplementary Figure 38. Phasing QV statistics. a)** QV statistics computed for the assemblies (pink), consensus haplotypes produced from the CHM13-based PanGenie-SHAPEIT phased genotypes for all regions (orange), for PanGenie-SHAPEIT within syntenic regions (regions shared between GRCh38 and CHM13) shown in green and consensus haplotypes produced from the GRCh38-based NYGC phased Illumina-calls (blue). In addition, we computed QVs for the two phased sets by switching sample labels ("randomized"), i.e., by using reads and phasings for different samples (in purple, brown, red). **b)** Completeness statistics computed for the consensus haplotypes produced from the CHM13-based PanGenie-SHAPEIT phased genotypes for all regions (yellow), for PanGenie-SHAPEIT within syntenic regions (regions shared between GRCh38 and CHM13) shown in green and consensus haplotypes produced from the GRCh38-based NYGC phased Illumina-calls (blue). In addition, we computed completeness for the two phased sets by switching sample labels ("randomized"), i.e., by using reads and phasings for different samples (in purple, brown, red).

Fig. 39

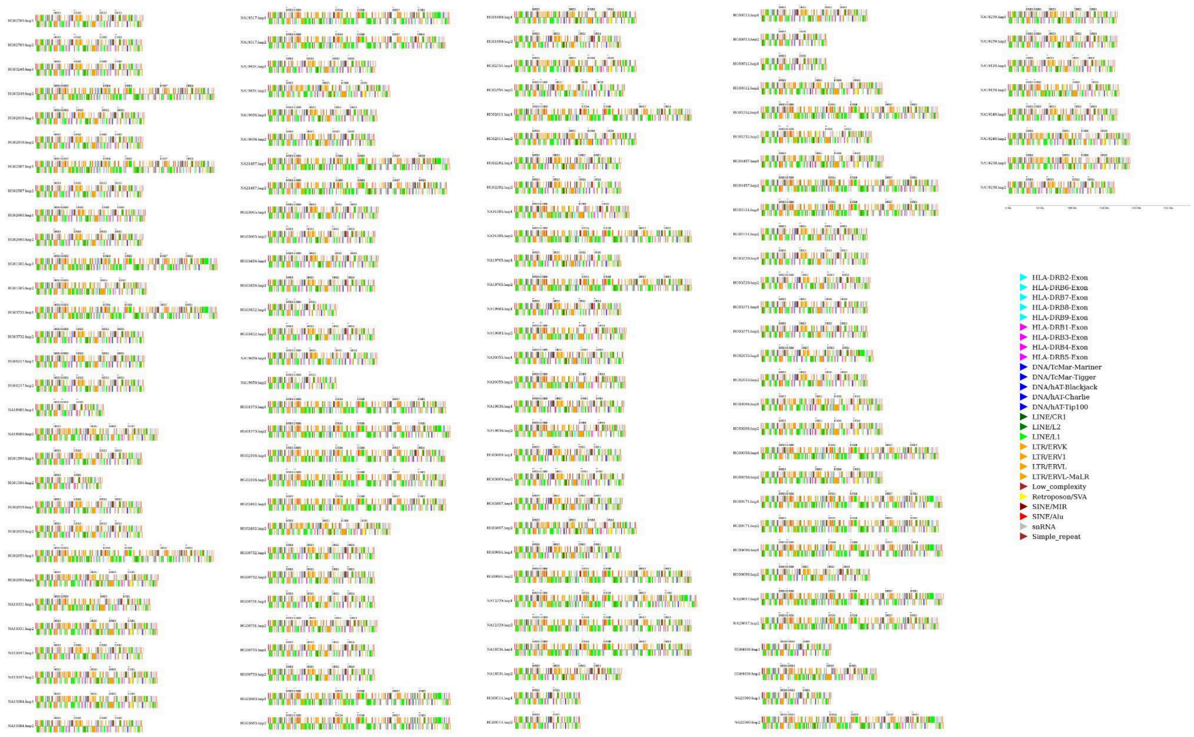


**Supplementary Figure 39. DRB Exon and repeat visualization.** Shows the repetitive element and gene/pseudogene exon distribution within all 128 sample haplotypes analyzed (2/130 sample haplotypes contained a stretch of Ns in the assembly and were not analyzed). Repetitive elements, gene exons, and pseudogene exons are uniquely colored. Lines that are oriented above the midline represent sense orientation, while those below the midline are in the antisense orientation. The first exons of each gene/pseudogene are denoted by an arrow with the name of the gene/pseudogene. The direction of the arrow also denotes the orientation of transcription of the gene compared to the region. Some haplotypes contain additional low divergent copies of the first DRB exon (denoted as a second *DRB9* first exon).

Fig. 40

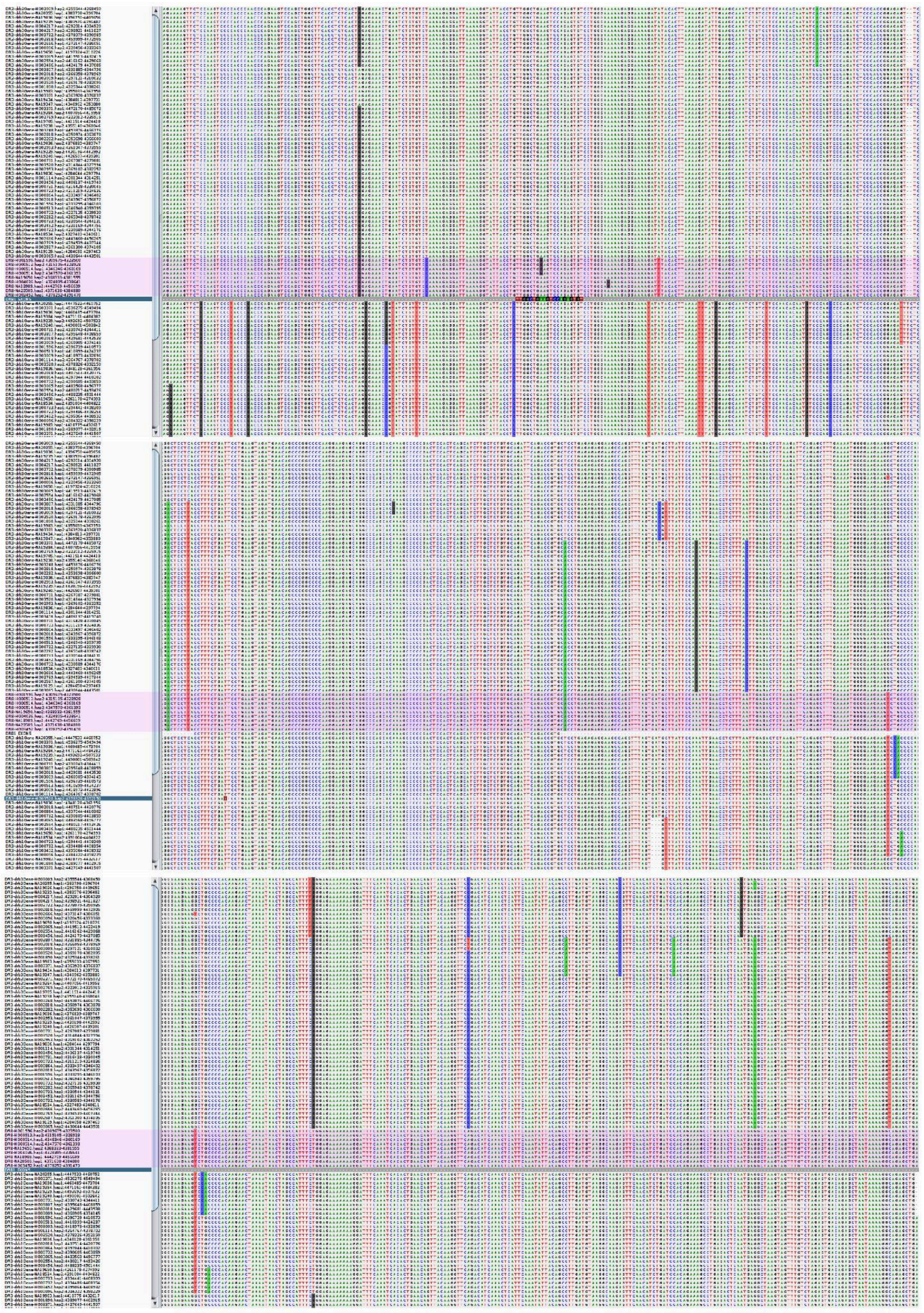


**Supplementary Figure 40. DR8 recombination.** This figure shows the results of a multiple sequence alignment of all sample haplotypes containing DR3/5/6 (*HLA-DRB3* and *HLA-DRB1* genes) and DR8 (*DRB1*). DR8 haplotypes are highlighted in purple and individual exons were placed in the alignment to assist the viewer. The top panel shows the alignment upstream of the identified breakpoint region. In this panel DR8 clearly shares more sequence similarity (more SNVs) with the *DRB3* gene of DR3/5/6 then it does to the *DRB1* gene. The middle panel shows the breakpoint region around exon 4/intron 3. Multiple sample haplotypes contain *DRB3* and *DRB1* genes which share perfect homology while others display shorter stretches of sequence homology. After exon 4 is where the *DRB1* gene in DR8 starts to share SNVs with the *DRB1* gene of DR3/5/6. The bottom panel shows the sequence immediately downstream of the middle panel and how the SNV pattern has switched from that position onward. The putative breakpoint showed a short length (<150bp) of homology between *HLA-DRB1* and *HLA-DRB3* on DR3/5/6 spanning exon 4 and the surrounding downstream sequence (intron 3) in some DR8 haplotypes suggesting that homology-mediated double strand DNA break repair may be a more likely explanation than NAHR. However, due to the putative ages of these haplotypes [152], as well as due to a subset of sample DR8 haplotypes displaying longer stretches (>200bp) of sequence homology, NAHR cannot be entirely discounted.

Fig. 41

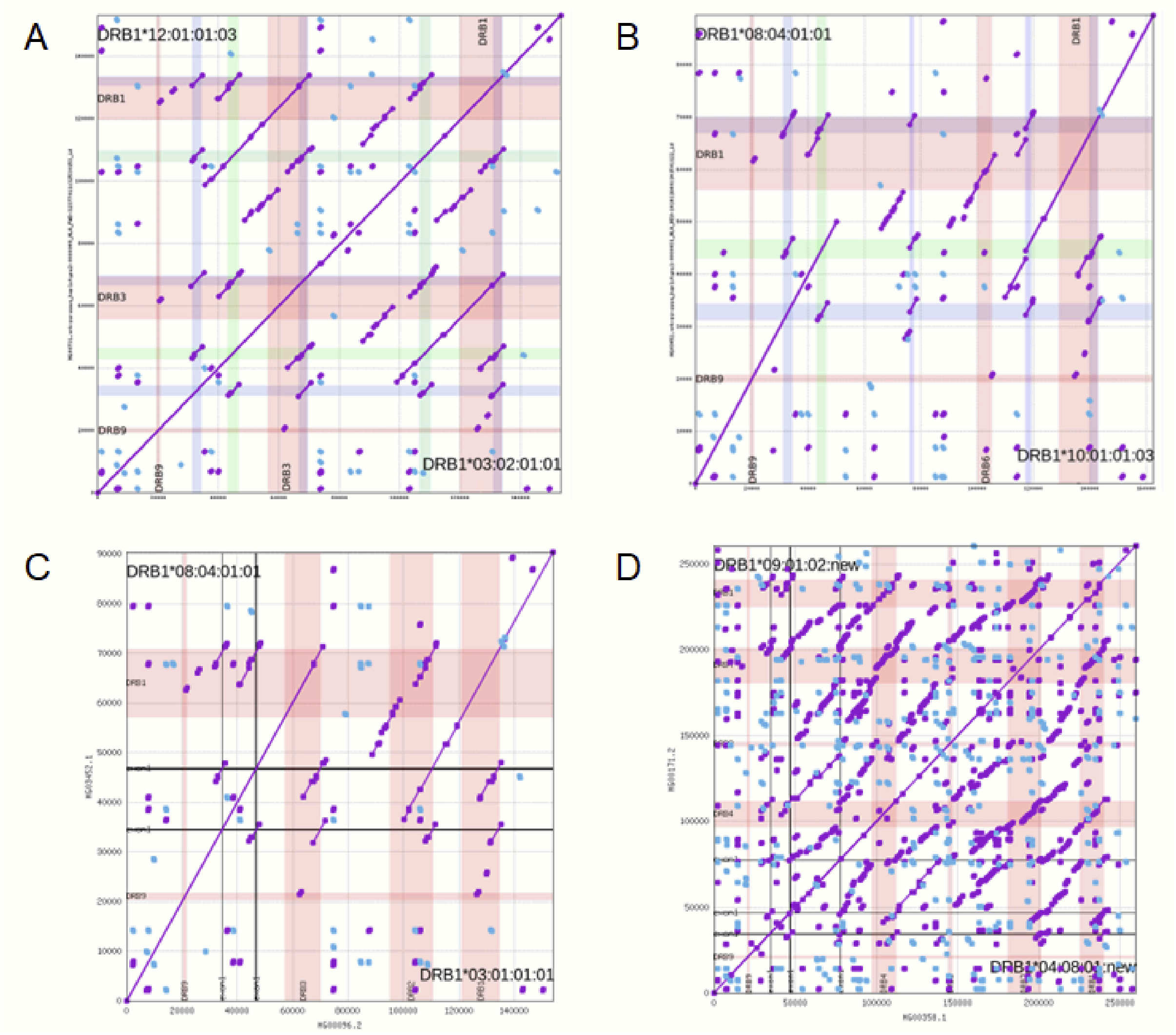


**Supplementary Figure 41. Comparison of DR group haplotypes.** Representative examples of pairwise alignments between DR group haplotypes DR5 and DRB3/6 (panel A); DR8 and DR1 (panel B); DR8 and DR3/6 (panel C); DR9 and DR4/7 (panel D) respectively. Red shading indicates the positions of HLA-DRB genes, blue/green shading matches to the identified HLA-DR small structural variants carrying solitary HLA-DRB exons.

Fig. 42

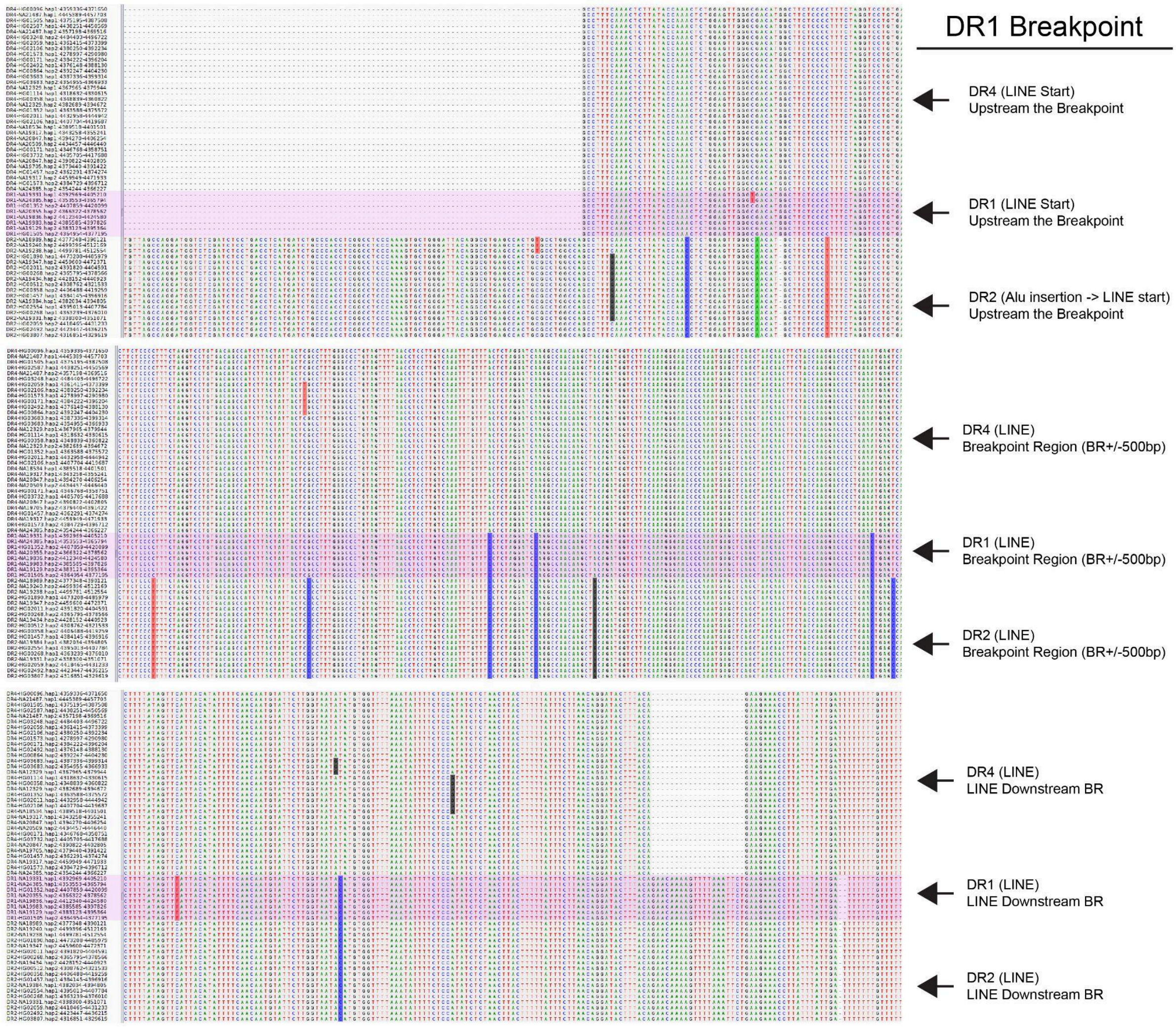


**Supplementary Figure 42. DR1 recombination.** This figure shows the results of a multiple sequence alignment around the identified breakpoint region (+/- 500bp) within all sample haplotypes containing DR4, DR1, and DR2. The top panel starts with an Alu insertion only present in DR2, followed by the LINE/L1 element. Within the LINE/L1 element we can see the shared sequence homology between DR4 and DR1 vs the SNVs within DR2. The middle panel is further within the LINE/L1 element where we start to see a switch in the SNV pattern to where DR1 and DR2 are sharing more SNVs with one another. This marks the +/- 500 base pair region where the breakpoint is believed to have occurred. The bottom panel is downstream this region showing how DR1 and DR2 continue to share SNVs as well as a short expansion/INDELs.

Fig. 43

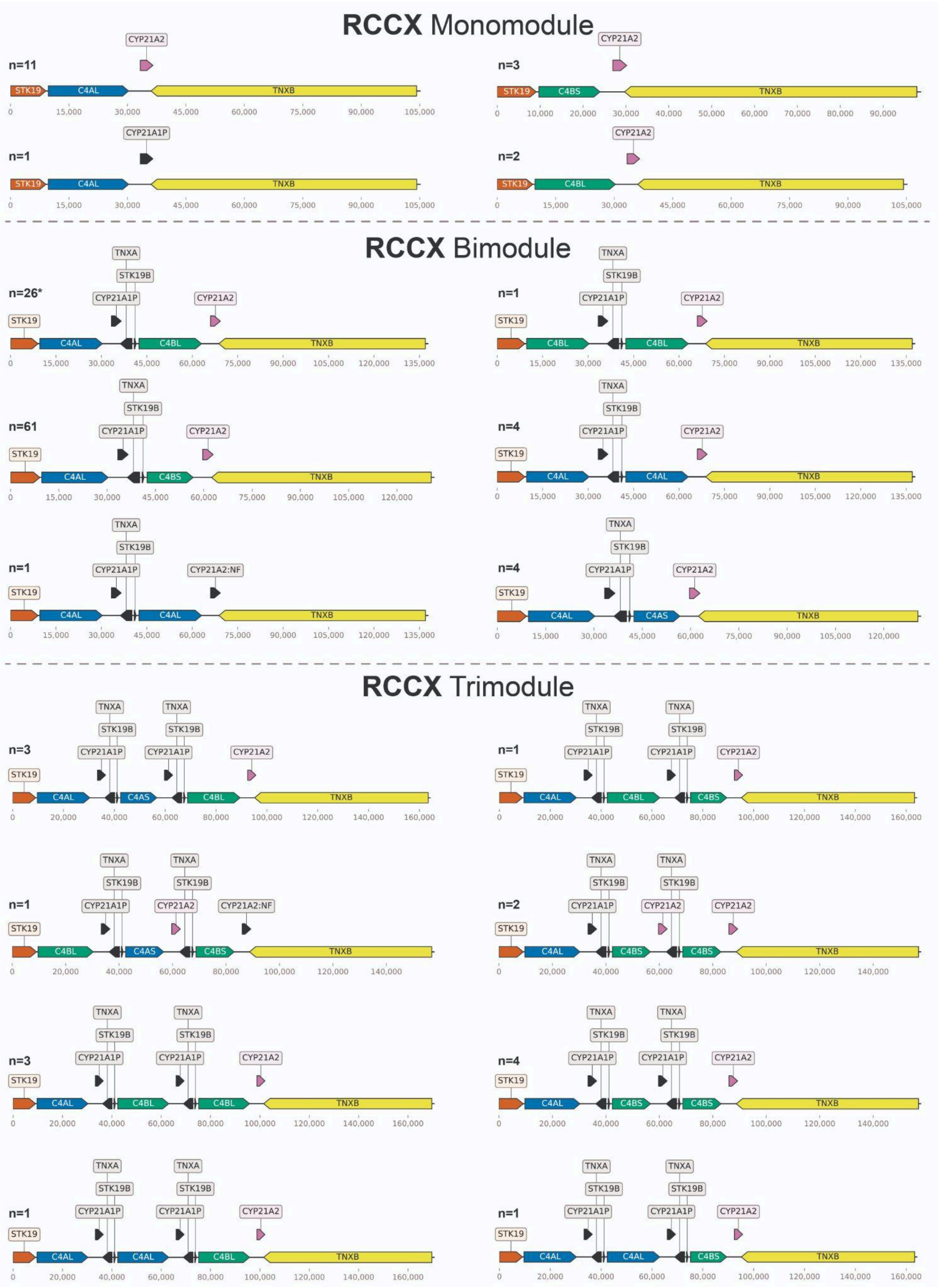
RCCX Monomodule
n=11
CYP21A2
STK19 C4AL TNXB
n=3
CYP21A2
STK19 C4BS TNXB
n=1
CYP21A1P
STK19 C4AL TNXB
n=2
CYP21A2
STK19 C4BL TNXB
RCCX Bimodule
n=26*
TNXA STK19B CYP21A1P CYP21A2
STK19 C4AL C4BL TNXB
n=1
TNXA STK19B CYP21A1P CYP21A2
STK19 C4BL C4BL TNXB
n=61
TNXA STK19B CYP21A1P CYP21A2
STK19 C4AL C4BS TNXB
n=4
TNXA STK19B CYP21A1P CYP21A2
STK19 C4AL C4AL TNXB
n=1
TNXA STK19B CYP21A1P CYP21A2:NF
STK19 C4AL C4AL TNXB
n=4
TNXA STK19B CYP21A1P CYP21A2
STK19 C4AL C4AS TNXB
RCCX Trimodule
n=3
TNXA STK19B CYP21A1P TNXA STK19B CYP21A1P CYP21A2
STK19 C4AL C4AS C4BL TNXB
n=1
TNXA STK19B CYP21A1P TNXA STK19B CYP21A1P CYP21A2
STK19 C4AL C4BL C4BS TNXB
n=1
TNXA STK19B CYP21A1P TNXA STK19B CYP21A2 CYP21A2:NF
STK19 C4BL C4AS C4BS TNXB
n=2
TNXA STK19B CYP21A1P TNXA STK19B CYP21A2 CYP21A2
STK19 C4AL C4BS C4BS TNXB
n=3
TNXA STK19B CYP21A1P TNXA STK19B CYP21A1P CYP21A2
STK19 C4AL C4BL C4BL TNXB
n=4
TNXA STK19B CYP21A1P TNXA STK19B CYP21A1P CYP21A2
STK19 C4AL C4BS C4BS TNXB
n=1
TNXA STK19B CYP21A1P TNXA STK19B CYP21A1P CYP21A2
STK19 C4AL C4AL C4BL TNXB
n=1
TNXA STK19B CYP21A1P TNXA STK19B CYP21A1P CYP21A2
STK19 C4AL C4AL C4BS TNXB

**Supplementary Figure 43. RCCX Architecture.** This schematic displays all unique RCCX mono/bi/tri module architectures found across the 130 sample haplotypes. Protein coding genes are *STK19*, *C4*, *CYP21A2*, and *TNXB*. Pseudogenes are *CYP21A1P*, *TNXA*, and *STK19B*. All C4 copies are functional. *CYP21A2*:NF represents a *CYP21A2* gene copy that is no longer functional due to coding for a premature stop codon. The long (L) or short (S) C4 designation represents if a HERV-K insertion within intron 9 is present. An asterisk (*) denotes the RCCX structure present within the CHM13 reference genome. The most frequently observed representatives of each haplotype configuration carried one functional copy of the *STK19*, *CYP21A2* and *TNXB* genes, as well as *C4AL*, *C4BS* and one copy of the pseudogenes *STK19B*, *CYP21A1P*, and *TNXA* (for n = 61 out of a total of 97 bi-modulesar modules); *C4AL* and no pseudogenes (for n = 11 out of a total of 17 mono-modules); and *C4AL*, 2 copies of *C4BS*, and two copies of the pseudogenes *STK19B*, *CYP21A1P*, and *TNXA* (for n = 4 out of a total of 16 tri-modules).

Fig. 44

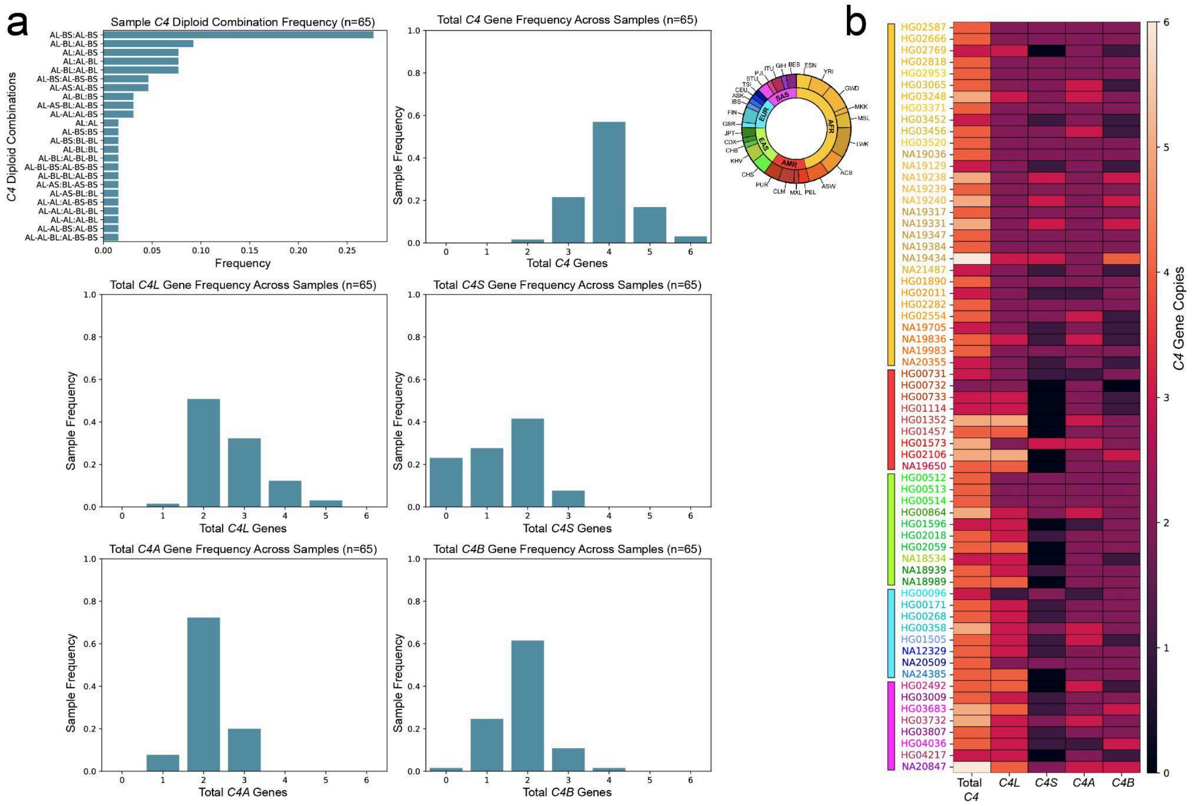


**Supplementary Figure 44. Sample C4 Statistics. a)** Shows six bar graphs displaying the *C4* genome diploid combination sample frequency (i.e., diplotype), total *C4* gene sample frequency, total *C4L* sample frequency, total *C4S* sample frequency, total *C4A* sample frequency and total *C4B* sample frequency. **b)** A heatmap showing the total counts of *C4*, *C4L*, *C4S*, *C4A*, and *C4B* within each sample. The heatmap cells are shaded by the total diploid count (black = zero, white = six copies). Sample names are colored based on their continental group/populations.

Fig. 45

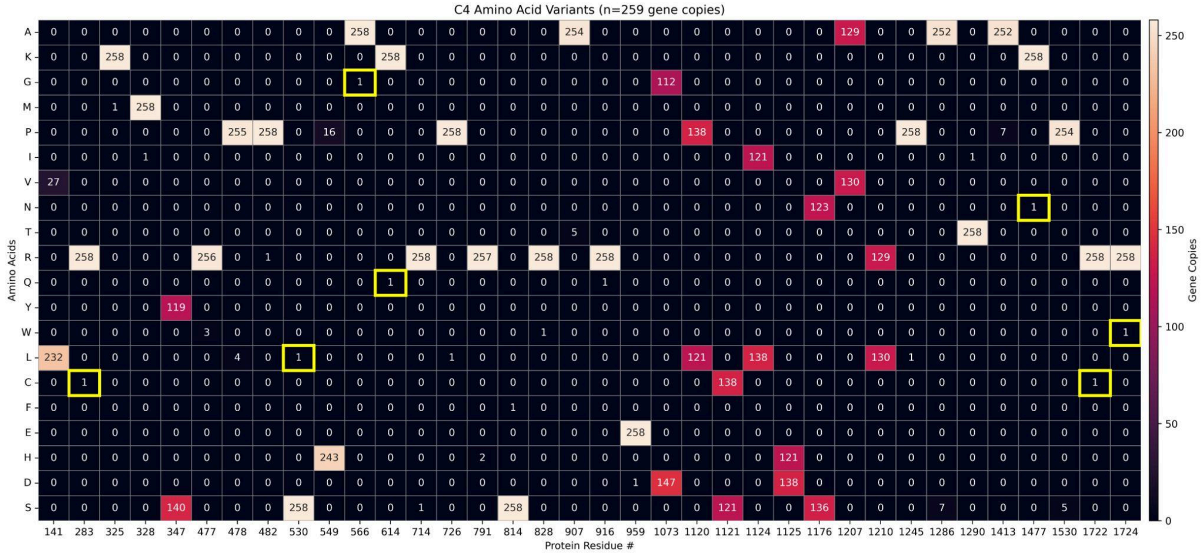


**Supplementary Figure 45. C4 Novel Variants.** Heatmap showing the variable amino acid positions within our 259 *C4* gene copies (across 130 sample haplotypes). The x-axis is the specific amino acid position while the y-axis are amino acids represented by a single letter. The seven novel amino acids found in our dataset are outlined in yellow. Each of these variants were only identified in a single *C4* copy.

Fig. 46

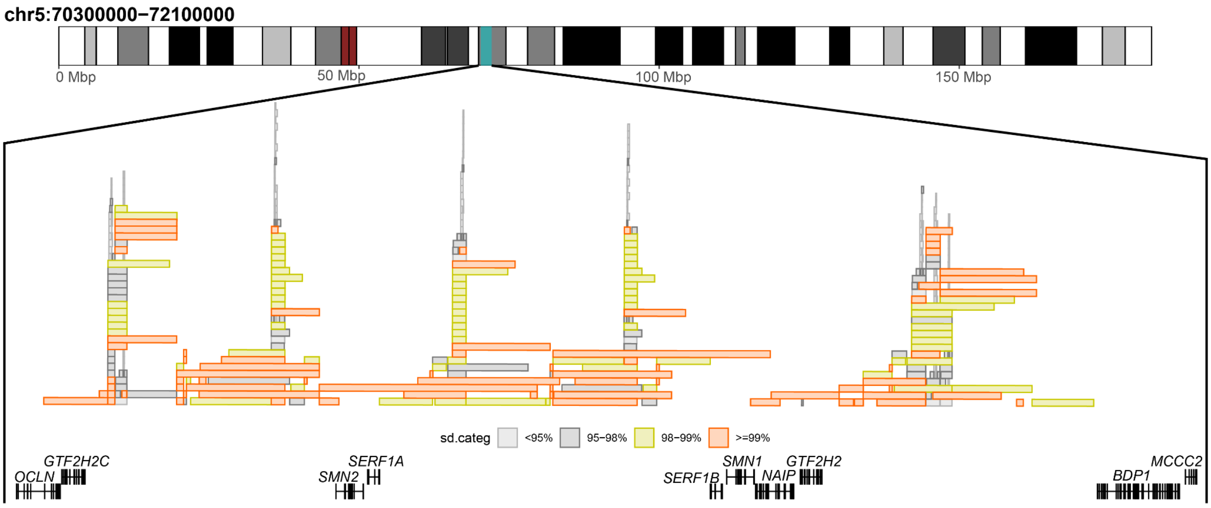


**Supplementary Figure 46. SMN region overview.** An overview of the SMN region that is highlighted on the chromosome 5 ideogram as a blue rectangle. Below we plot the extent of segmental duplications in this region color by the sequence identity (sd.categ). Further below we plot the gene model of protein-coding genes in this region.

Fig. 47

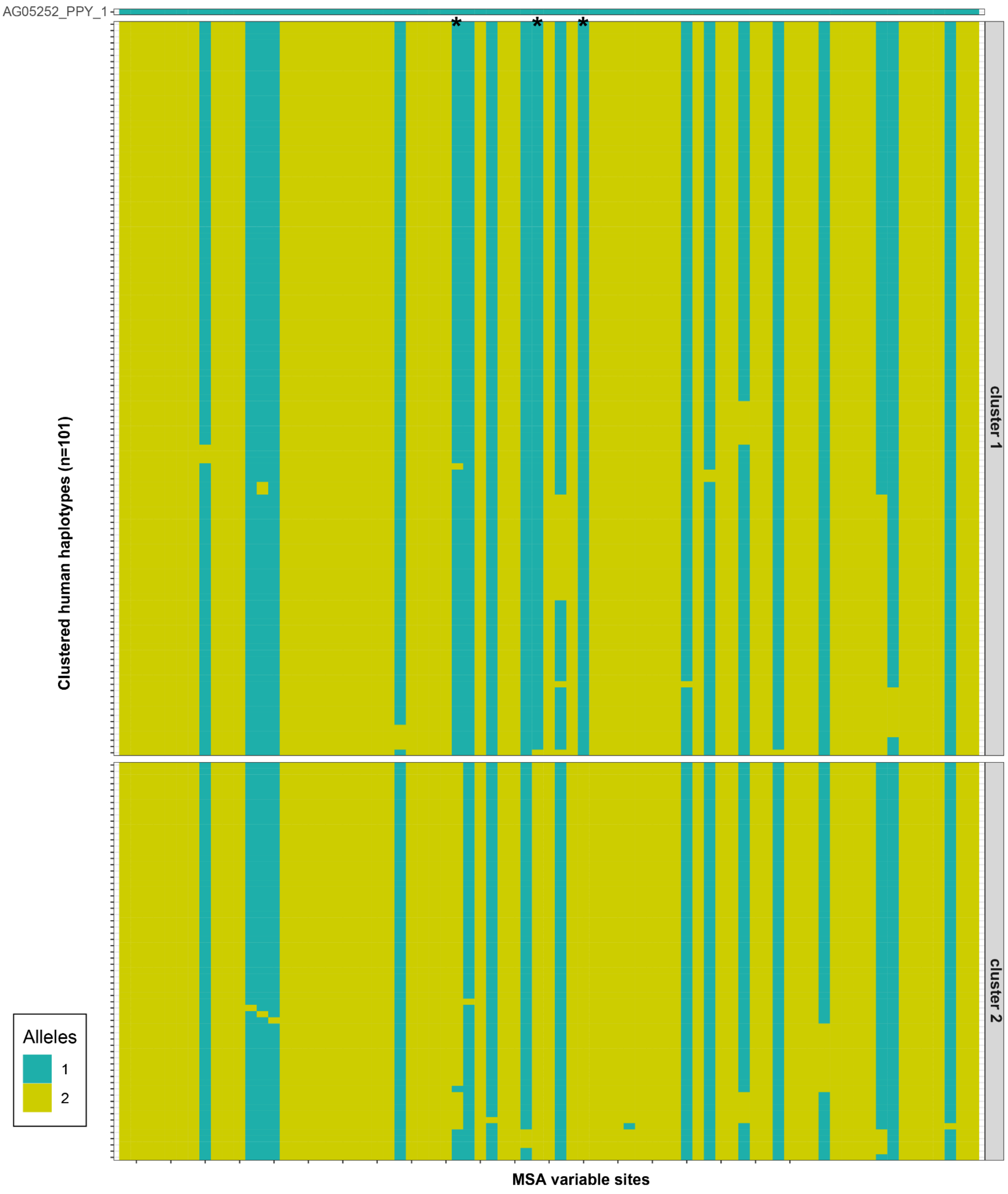


**Supplementary Figure 47. Distinguishing SMN1 and SMN2 gene copies.** A visualization of a multiple sequence alignment between exonic sequences for all SMN gene copies across all 101 human haplotypes. We used orangutan (PPY) as an outgroup to mark ancestral (green) and derived alleles (yellow). We cluster all haplotypes into two clustered mostly distinguished based on 3 ancestral alleles present in cluster 1. We mark cluster 1 as *SMN1* gene copy while cluster 2 is marked as *SMN2* gene copy.

Fig. 48

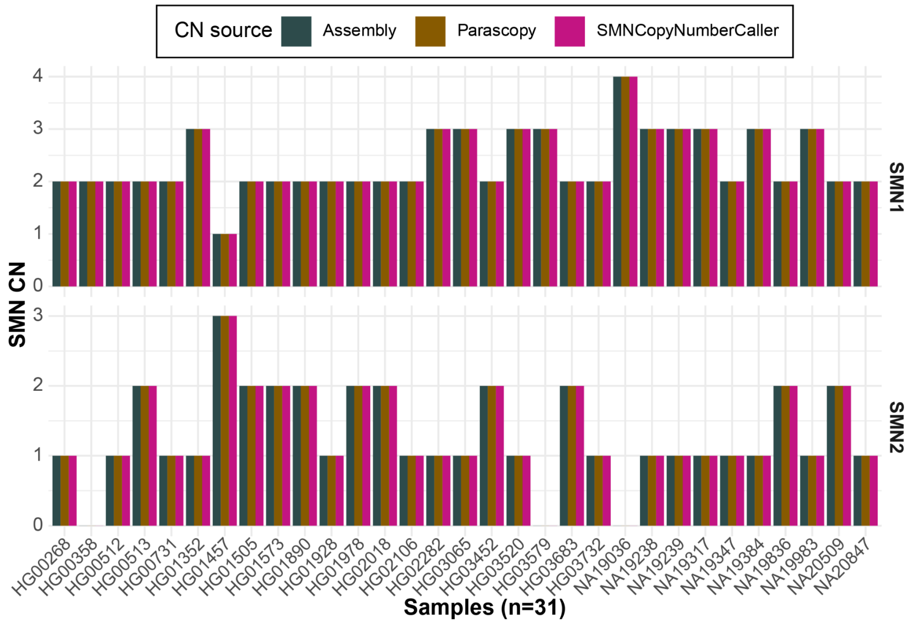


**Supplementary Figure 48: Genotyping of SMN genes.** Comparison of predicted *SMN1* and *SMN2* gene copy numbers defined by assemblies (dark blue) in comparison to short-read based genotyping methods (Parascopy - brown, SMNCopyNumberCaller - pink). We could compare results only for 31 samples which have fully assembled both haplotypes and could be compared to diploid copy number reported by unphased short-reads.

Fig.49

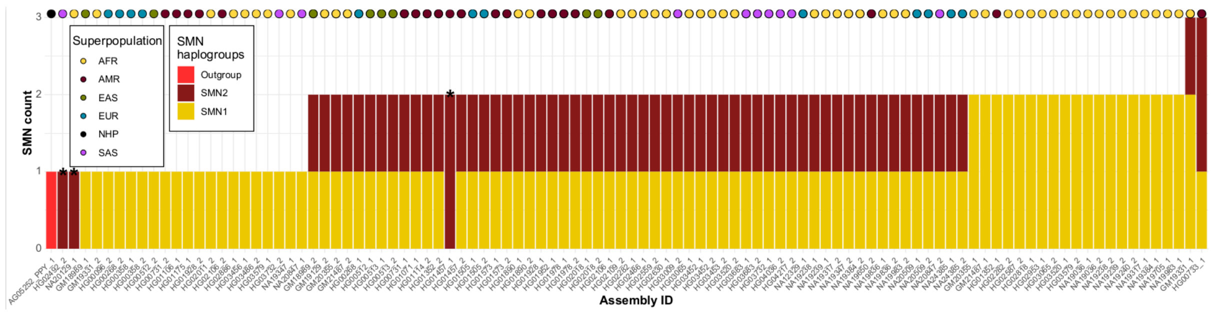


**Supplementary Figure 49. Summary of SMN genes.** Summary of *SMN1* (yellow) and *SMN2* (red) gene copies genotyped in each human haplotype (n=101). Haplotype that carry only *SMN2* gene copy are highlighted by the asterisks. Superpopualtion identity of each haplotype is marked by the colored dot on top of each bar (see legend).

Fig. 50

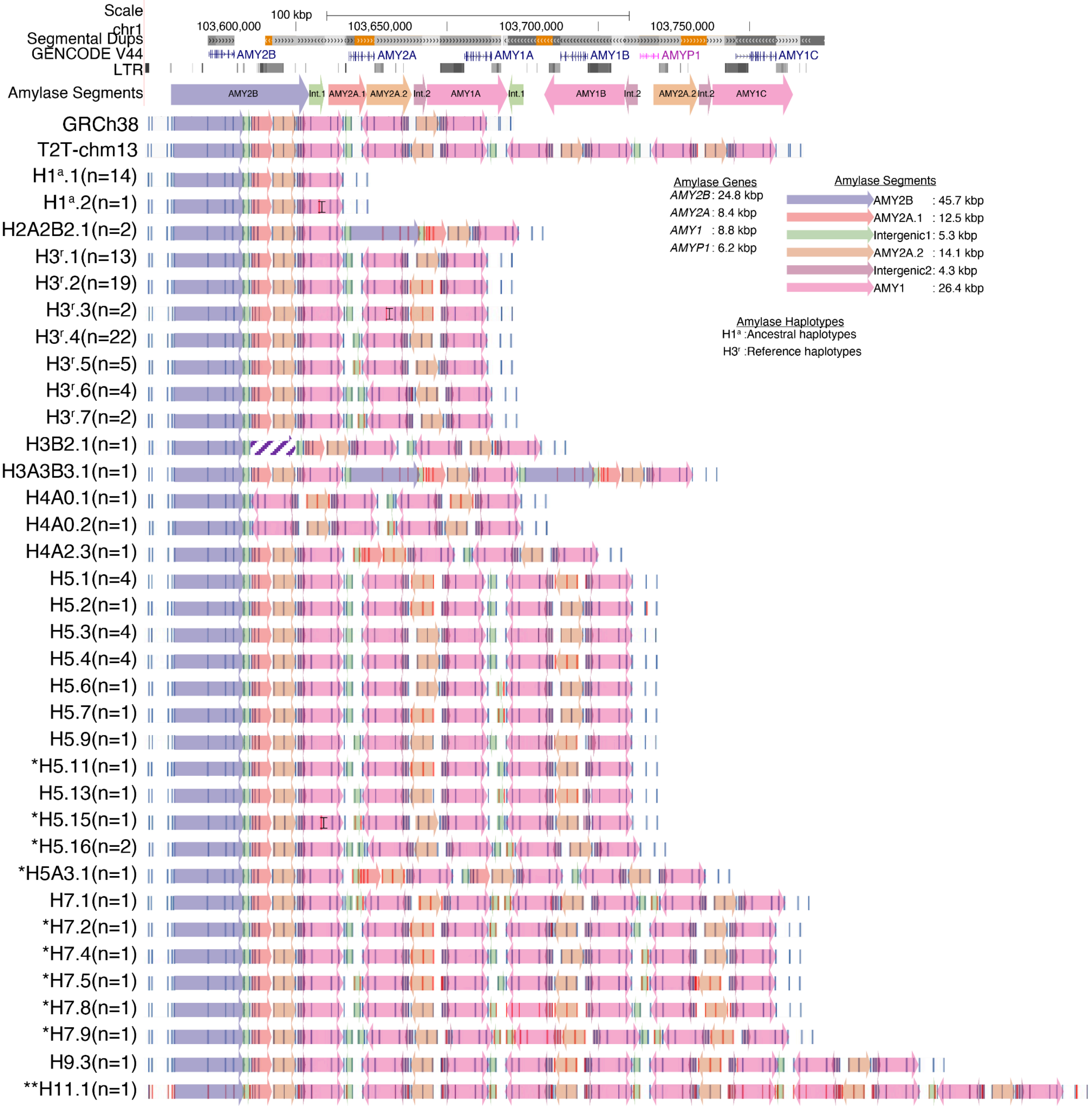


**Supplementary Figure 50. Amylase haplotypes identified in this study.** Segmental duplications (light to dark gray: 90-98% similarity, light to dark orange: > 99% similarity), GENCODE V44 gene annotations, and long terminal repeats (LTRs) are represented as tracks. The lower panel shows amylase segments (colored arrows). The AMY2B segment overlaps the *AMY2B* gene, AMY2A.1 and AMY2A.2 segments overlap the *AMY2A* gene, and the AMY1 segment overlaps the *AMY1* gene. The GRCh38 and T2T-chm13 reference genomes and the unique amylase haplotypes resolved in our dataset (n = 35) are represented with in silico maps with white background displaying label positions in dark blue and amylase segments with color-coded arrows. The vertical black line in the second AMY1 segment of H3ʳ.3 represents the polymorphic label present in three alleles. The diagonal stripes in the second AMY2B segment of H3B2.1 indicate that it is a partial copy of the first AMY2B segment. Haplotype IDs describe: HX: X denotes the number of *AMY1* copies; AX:

X denotes the number of *AMY2A* copies; BX: X denotes the number of *AMY2B* copies. The superscript “a" denotes the ancestral amylase haplotype structure, and the superscript “r” denotes the reference amylase haplotype structure. The number in parentheses indicates the number of alleles. Single asterisk (*) denotes haplotypes that were resolved at the base-pair level for the first time. Double asterisk (**) denotes novel amylase haplotype.

Fig. 51

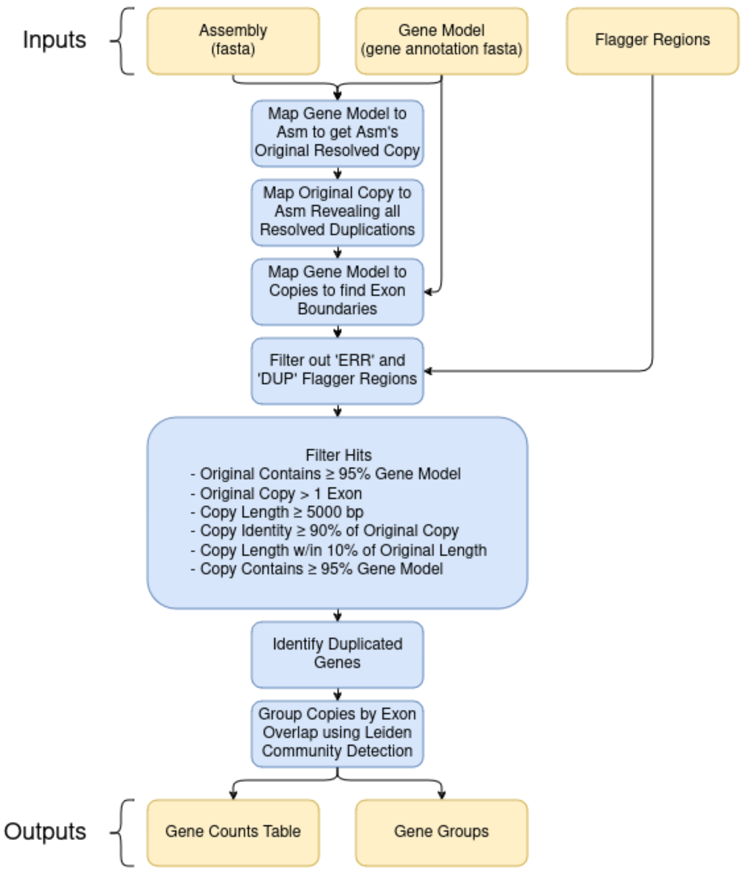


**Supplementary Figure 51. Segmental Duplication Annotation Pipeline flowchart.** Minimap2 was used for all mapping and alignment.

Fig. 52

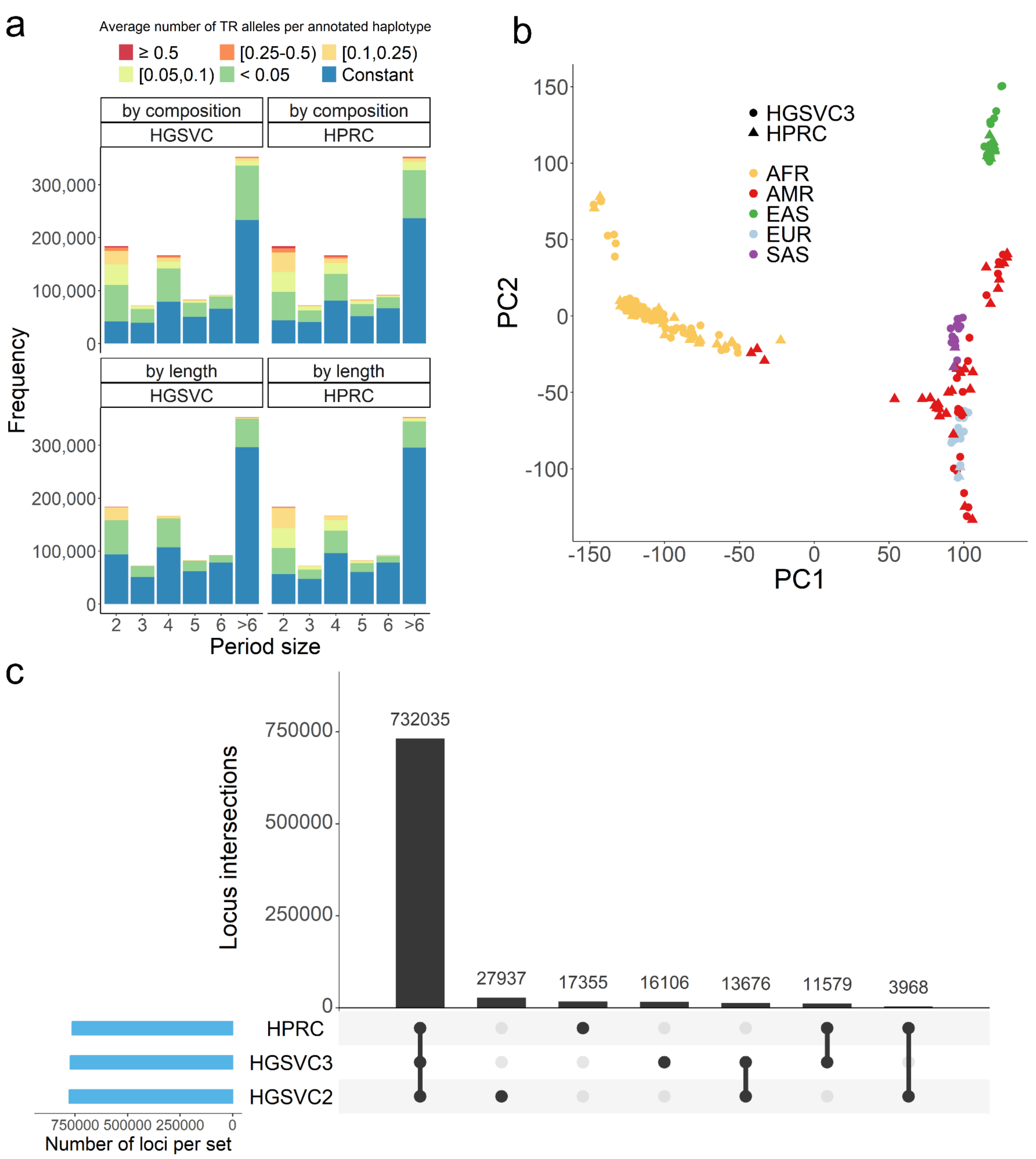


**Supplementary Figure 52. VNTR diversity.** a: variability of TRs in HPRC and HGSVC3 genomes. b: population clustering of HPRC and HGSVC3 genomes by TRs. c: TRs from Tandem Repeat Finder were merged and compared for the three datasets. Centromere regions were masked.

Fig. 53

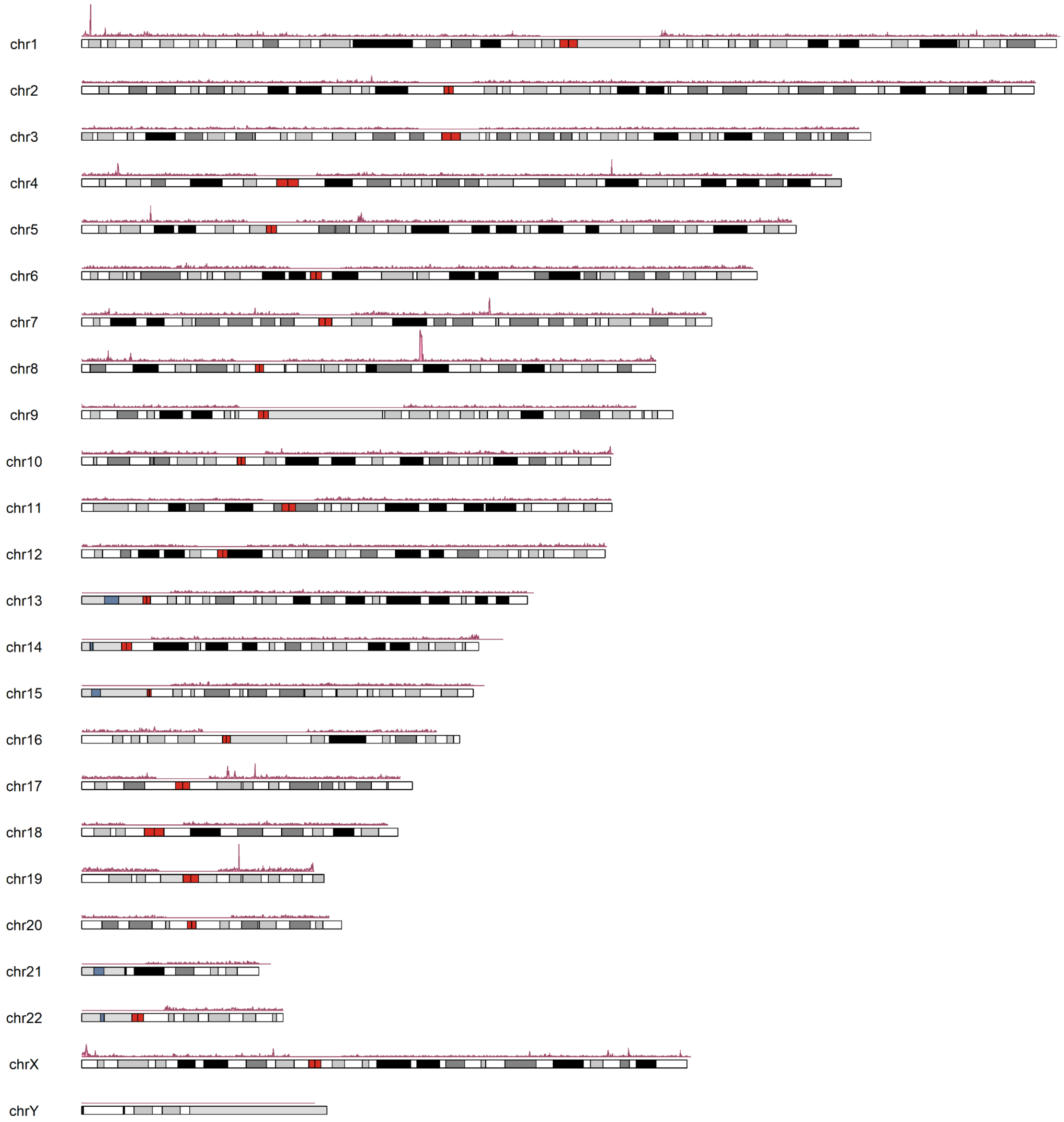


**Supplementary Figure 53. Unique TRs in HGSVC3 compared to HPRC.** Distribution of unique TRs by Tandem Repeat Finder in HGSVC3 genomes compared to HPRC genomes. Ideograms were generated using a bin size of 1,000,000bp and centromere regions were masked.

Fig. 54

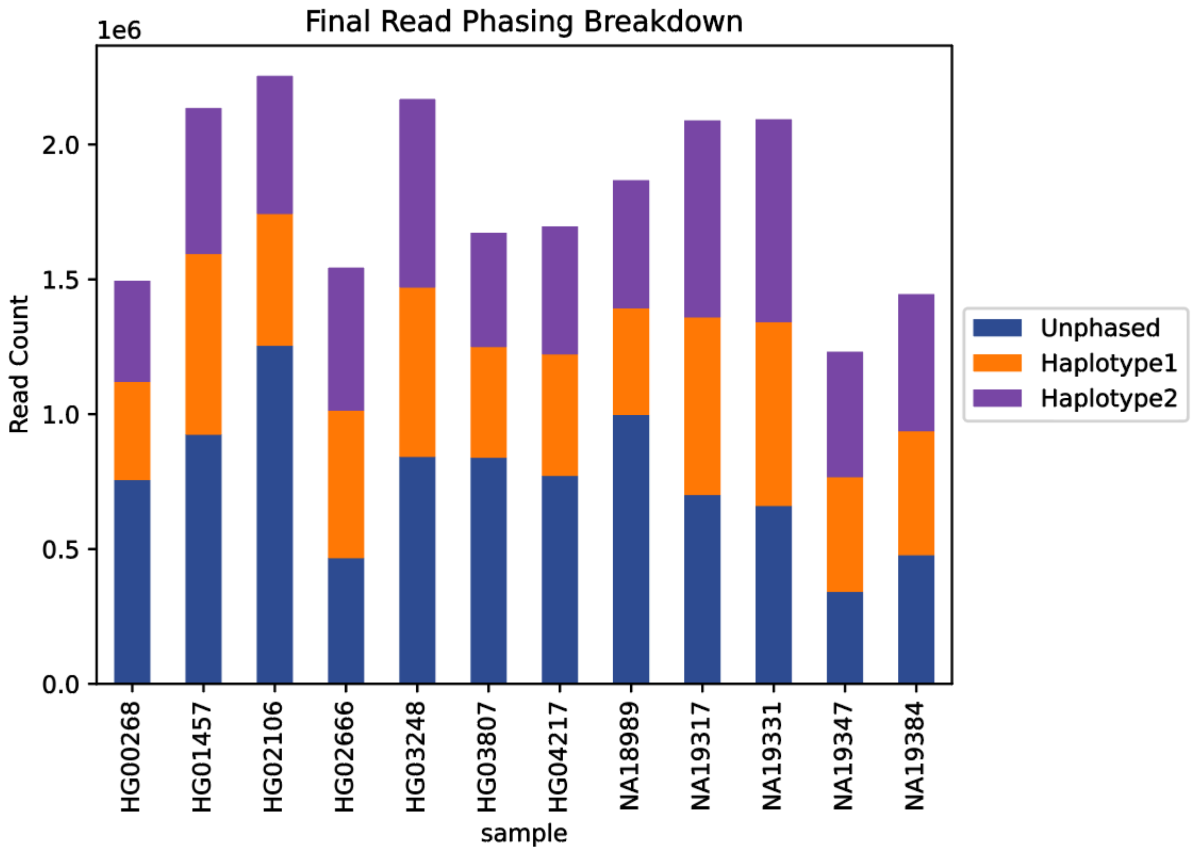


**Supplementary Figure 54. Iso-Seq Read Phasing Breakdown.** The total number of PacBio Iso-Seq read phasing assignments to haplotype 1 (orange), haplotype2 (purple), and unphased (unable to be phased) (navy) per sample using an ensemble alignment-based phasing approach [methods]. Reads designated as unphased aligned to both haplotypes equally well. On average 58.44% of reads were able to be phased to a haplotype of a sample.

#### Fig. 55

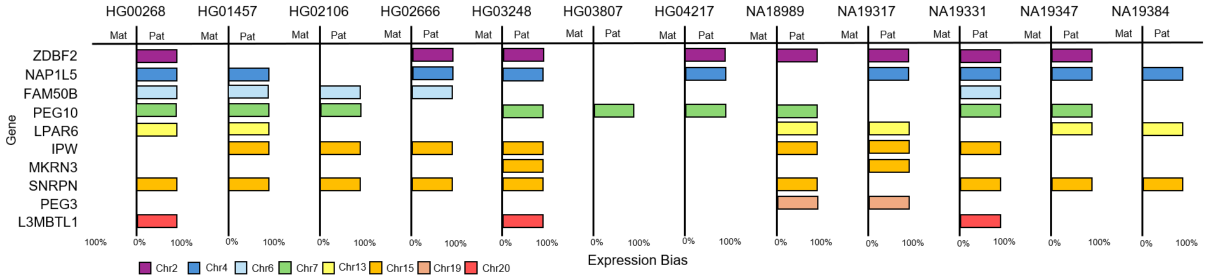


**Supplementary Figure 55. Imprinted Genes According to Iso-Seq.** Depiction of ten genes (y-axis) expressed in EBV-transformed B-lymphocyte cell lines that were identified as imprinted utilizing phased sample-specific assemblies for mapping of Iso-Seq reads. Mono-haplotype expression was declared if sample-specific reads phased to a single haplotype. Bar color represents the chromosome on which the gene resides. The imprinted expression pattern observed in our study is in concordance with the previous literature [89–91].

#### Fig. 56

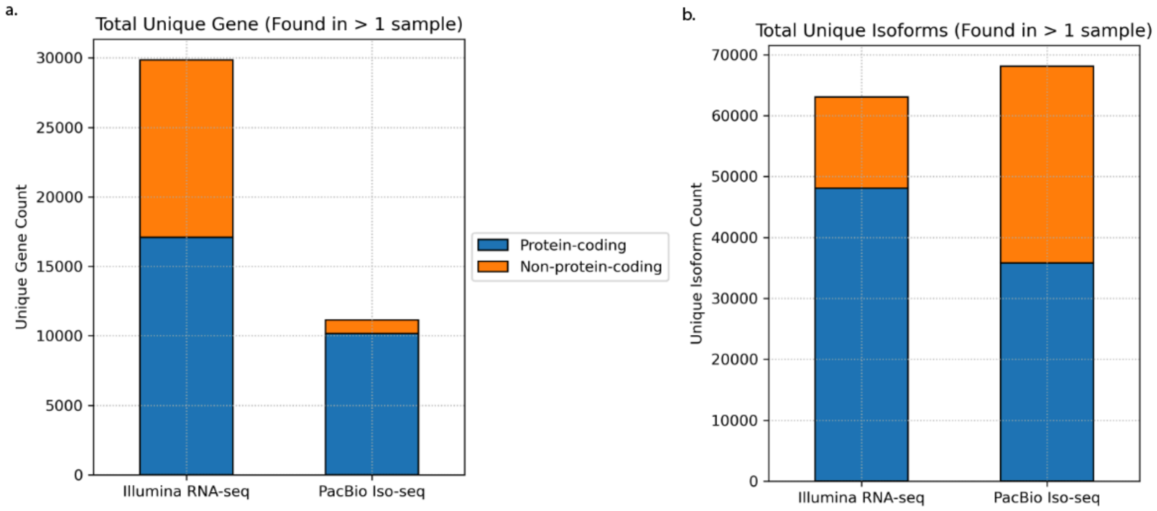


**Supplementary Figure 56. Short- vs Long-Read RNA Sequencing Total Unique Genes and Isoforms. a)** Comparison of the total number of unique genes identified across 12 samples using Illumina RNA-seq vs PacBio Iso-Seq. Only genes expressing isoforms identified in two or more samples were included in this analysis. Hue indicates the proportion of genes that are protein coding (blue) or non-protein coding (orange). Results show that a greater total number of unique genes were identified by the Illumina RNA-seq (29,858 unique genes) versus the PacBio Iso-Seq (11,134 unique genes) across 12 samples. **b)** Comparison of the total number of unique isoforms present in at least two out of 12 samples in Illumina RNA-seq and PacBio Iso-Seq. Hue indicates the proportion of isoforms that are from protein-coding (blue) or non-protein-coding genes (orange). This graph indicates that a greater total number of unique isoforms were identified in the the PacBio Iso-Seq dataset (68,097 unique isoforms) compared to the Illumina RNA-seq callset (63,028 unique isoforms) across the 12 samples.

Fig. 57

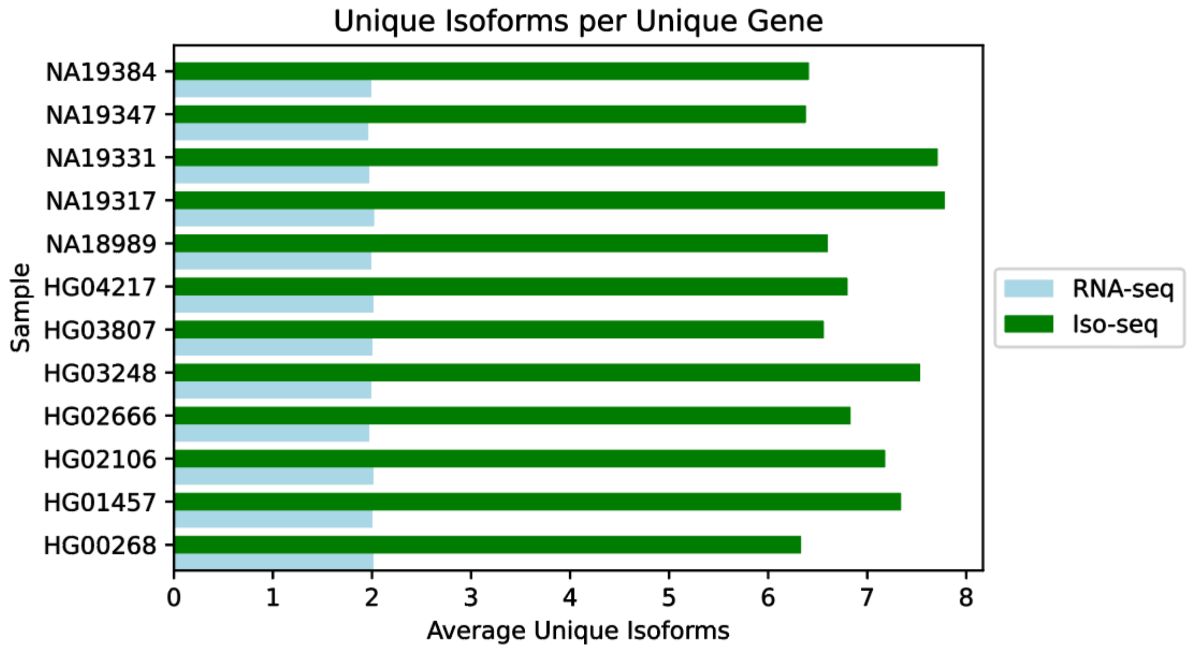


**Supplementary Figure 57. Average Unique Isoforms Per Unique Gene.** Comparison of the average number of unique isoforms identified (present in at least two out of twelve samples) per unique gene represented in Illumina RNA-seq (light blue) and PacBio Iso-Seq (green) data. On average, a greater number of unique isoforms per unique gene were found using the Iso-Seq versus the RNA-seq datasets.

Fig. 58

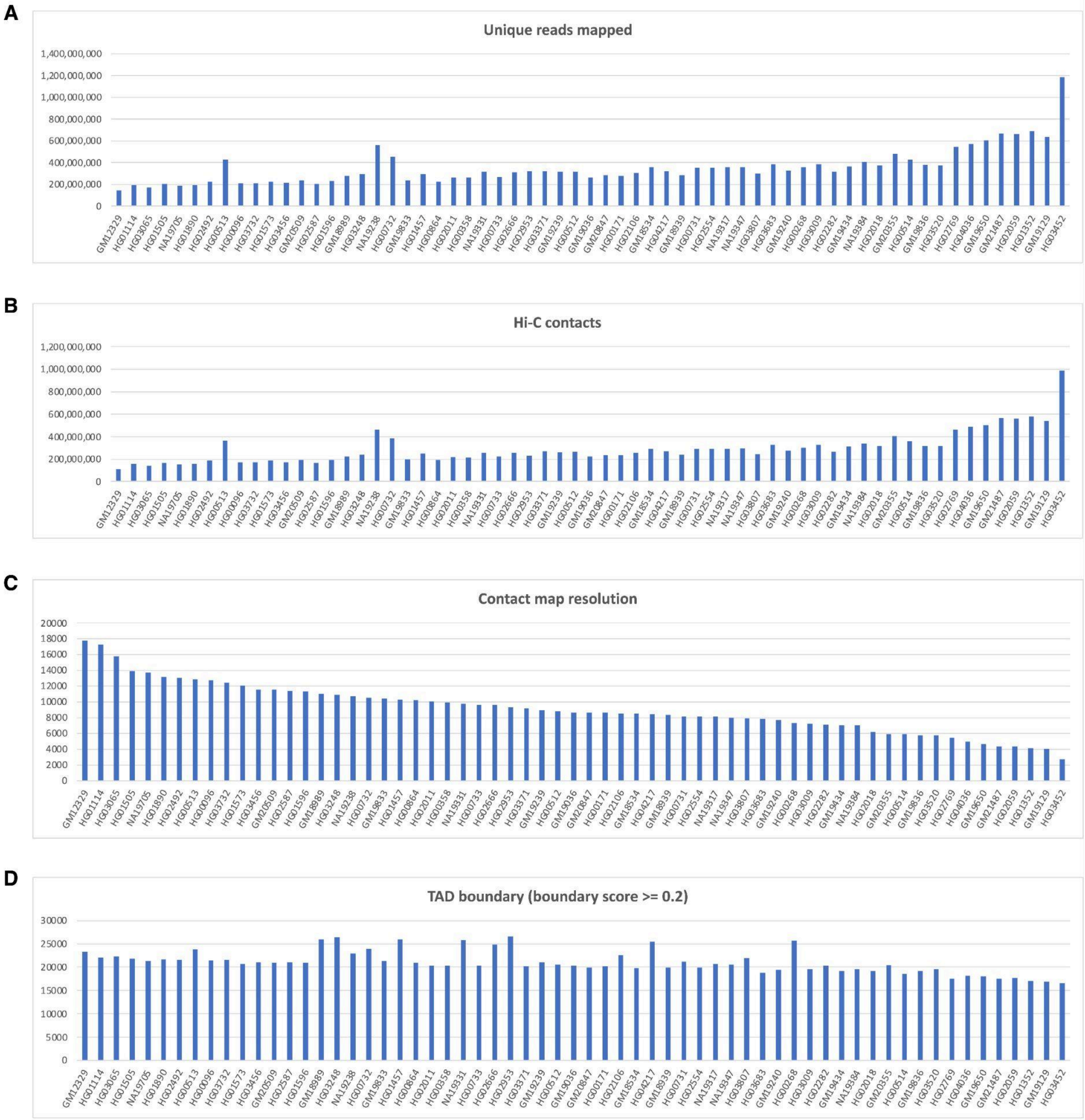


**Supplementary Figure 58. Summary statistics of Hi-C data and chromatin structures**. **A)** Unique reads were mapped to the hg38 reference genome. The x-axis displays sample IDs ordered from lower to higher resolution (calculated based on the GRCh38). The y-axis shows the total number of the mapped reads. **B)** The number of Hi-C contacts for each mapped sample. **C)** The calculated resolution (in base pair) of Hi-C contacts for each sample. **D)** The number of TAD boundaries identified for each sample (filtered by the boundary score as 0.2). The average number of TAD boundaries identified was 21,021 for the hg38 reference genome. No significant correlation was observed between the number of TAD boundaries and contact map resolution, underscoring the robustness of the *IS* algorithm, which is not constrained by the data resolution.

### Fig. 59

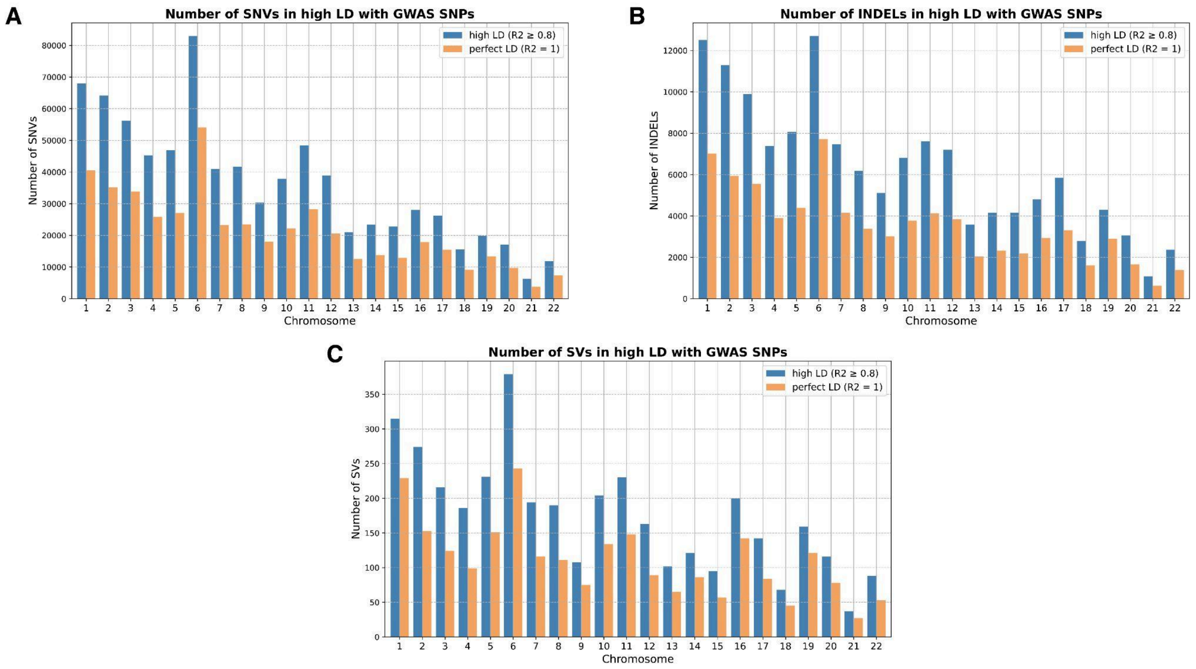


**Supplementary Figure 59. SNVs, INDELs, and SVs that are in high LD with GWAS SNPs. A)** Number of SNVs in high LD with GWAS SNPs. X-axis represents chromosomes and the y-axis indicates the number of variants. Blue bars denote variants with an LD threshold of $R^2 \geq 0.8$. Orange bars denote variants in the perfect LD which has $R^2 = 1$. **B)** Number of INDELs in high LD with GWAS SNPs. **C)** Number of SVs in high LD with GWAS SNPs.

Fig. 60

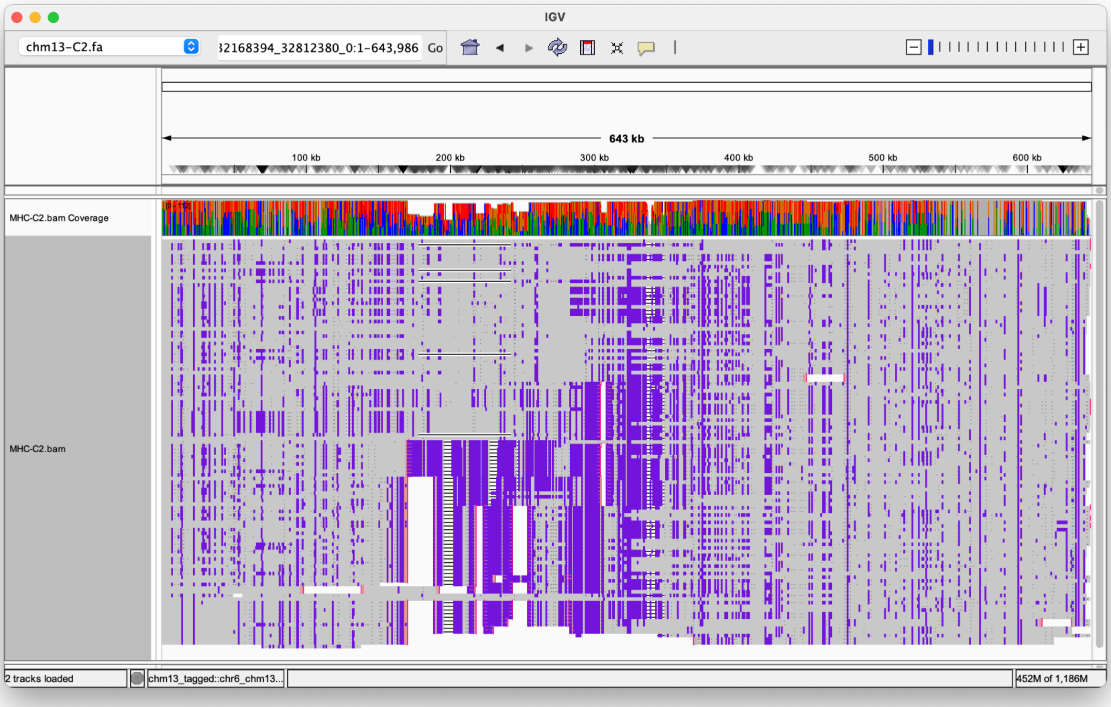


**Supplementary Figure 60. MHC C2 alignment.** The sequence alignment generated by minimap2 of the HGSVC genomes to the CHM-13 reference. The standard alignment process fails to alignment the high diversified regions showing as gaps and high density of variations around the gaps.

Fig. 61

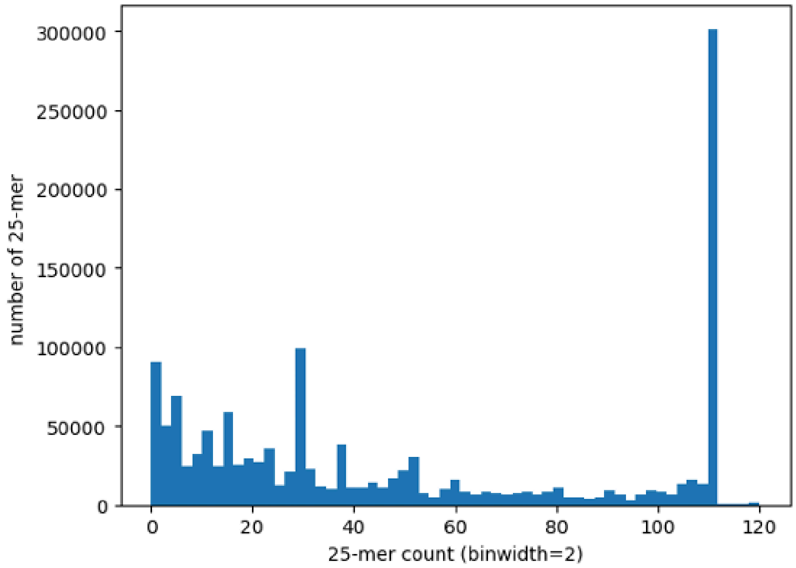


**Supplementary Figure 61. MHC C2 k-mer count histogram.** All 25-mer count through the MHC class II region of the pangenome. While the majority of the 25-mers are from conserved (among different genomes) and uniqued (in single genome) as shown by the peak around 25-mer count = ~111, there are abundance of less conserved *k*-mers ranged 1 to 50.

Fig. 62

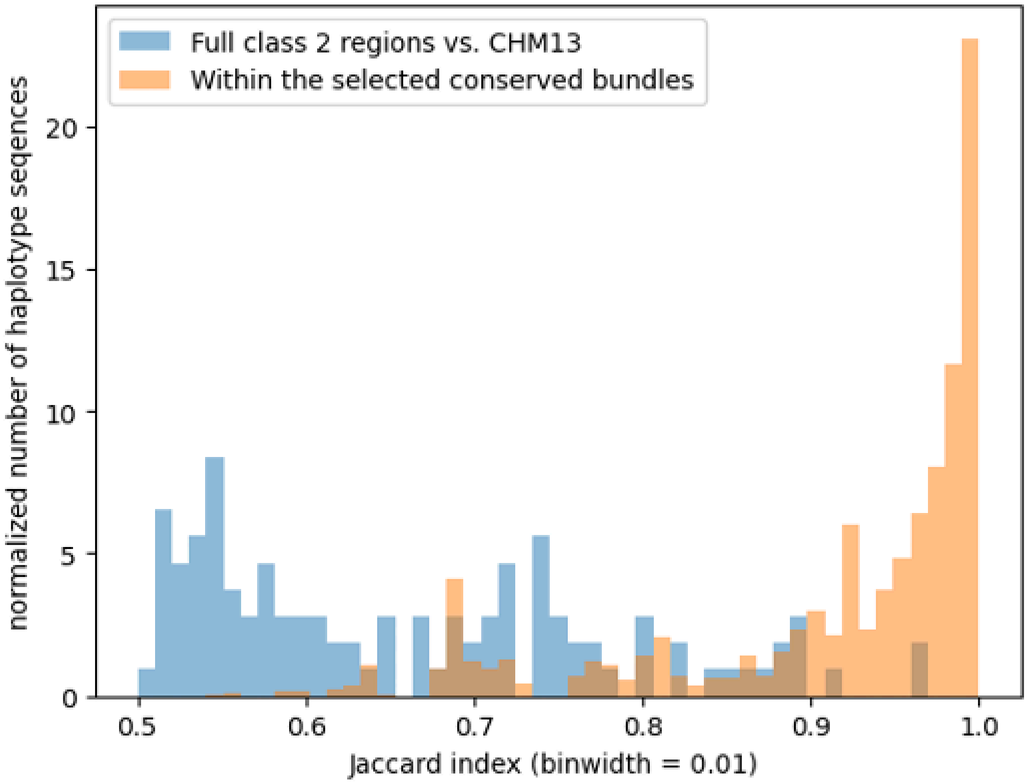


**Supplementary Figure 62. Class II MHC vs CHM13 Jaccard index.** The Jaccard index comparison of the class II regions vs. CHM13 and within the conserved bundles.

Fig. 63

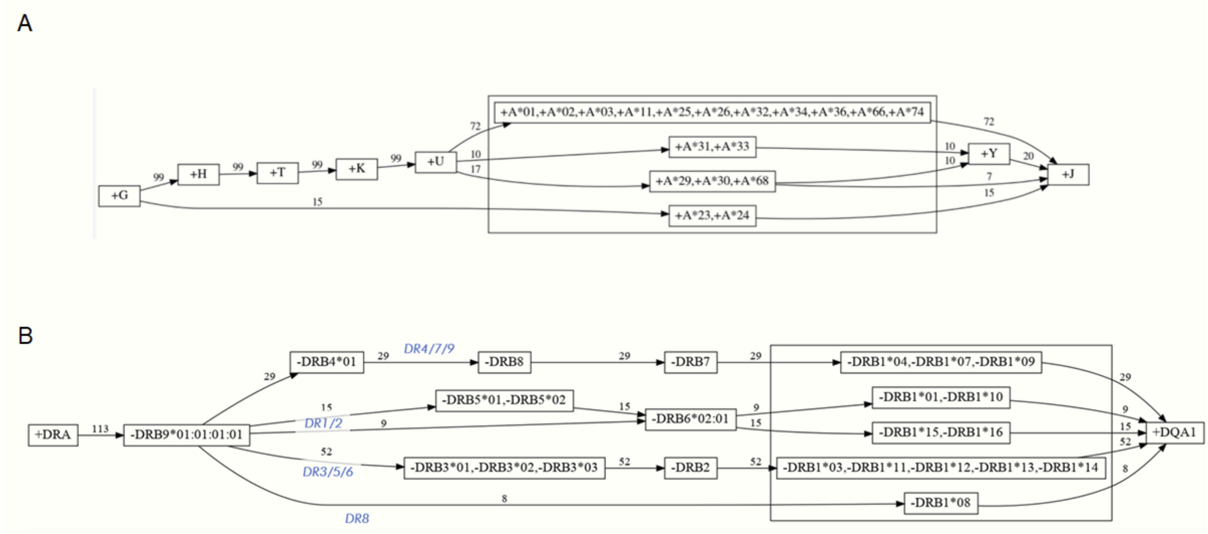


**Supplementary Figure 63. MHC gene structure.** Visualization of the MHC class I (panel A) and MHC class II (panel B) haplotype structure with respect to linkage equilibrium between HLA gene allele classes and presence/absence patterns, based on annotation with Immuannot [145]. Blue labels in panel B indicate which DR group the shown haplotypes belong to.

Fig. 64

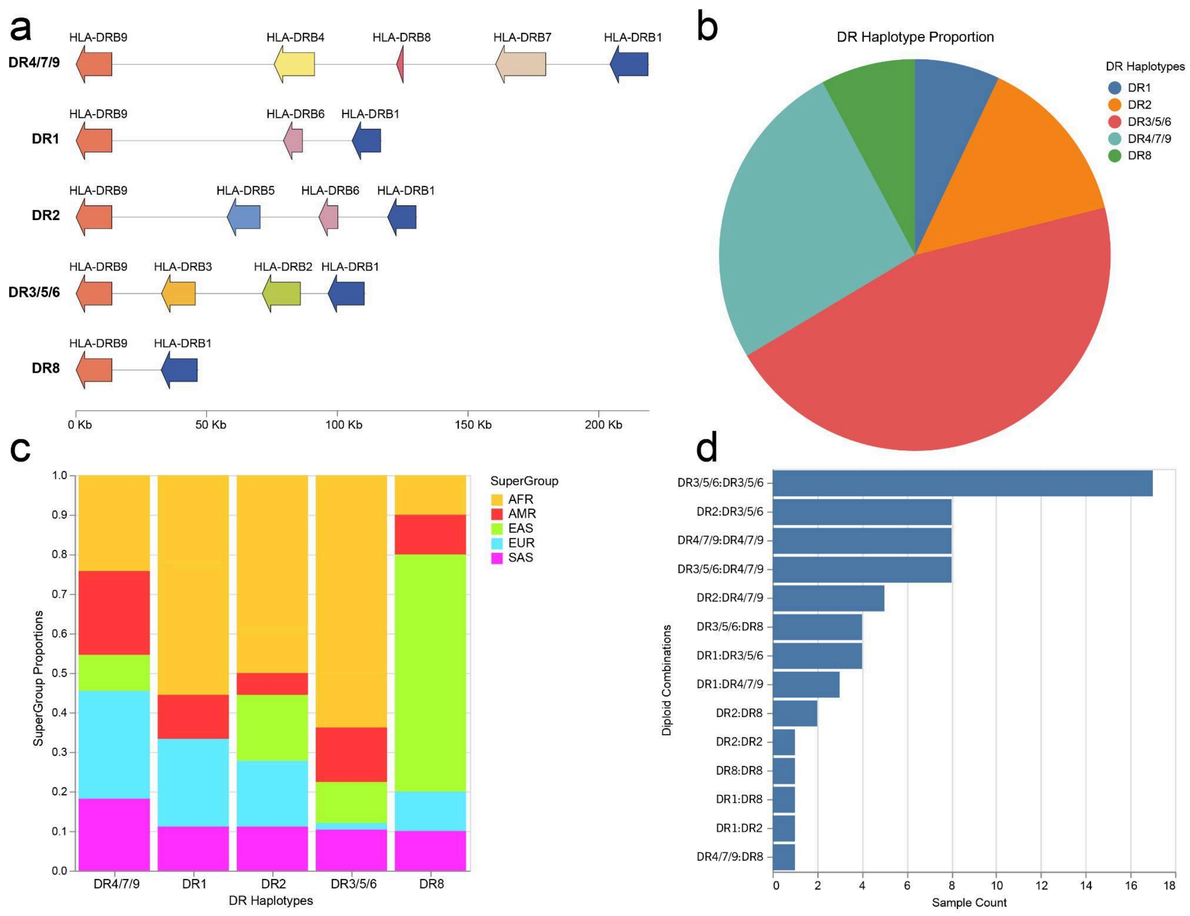


**Supplementary Figure 64. DR haplotype diagram and statistics. a)** A diagram showing the typical length of the five unique DR haplotypes and the genes (*HLA-DRB1*, *-DRB3*, *-DRB4*, and *-DRB5*) as well as the pseudogenes (*-DRB2*, *-DRB6*, *-DRB7*, *-DRB8*, and *-DRB9*) which comprise them. **b)** A pie chart showing the frequency of DR haplogroups within the 128 sample haplotypes (2 haplotypes contained N's in the DRB region and were not analyzed). DR3/5/6 was the most frequently observed DR haplotype **c)** A stacked bar graph showing the proportion of sample supergroups (AFR: African, AMR: American, EAS: East Asian, EUR: European, SAS: South Asian) found containing each DR haplotype. Interestingly, no east asians (EAS) in our sample set contained a DR1 haplotype, though they made up the majority of the DR8 haplotype. **d)** A bar graph showing the total samples (x-axis) containing DR haplotype diploid combinations (y-axis). The diploid combination of two DR3/5/6 haplotypes within a genome was the most frequently observed across our sample set.

Fig. 65

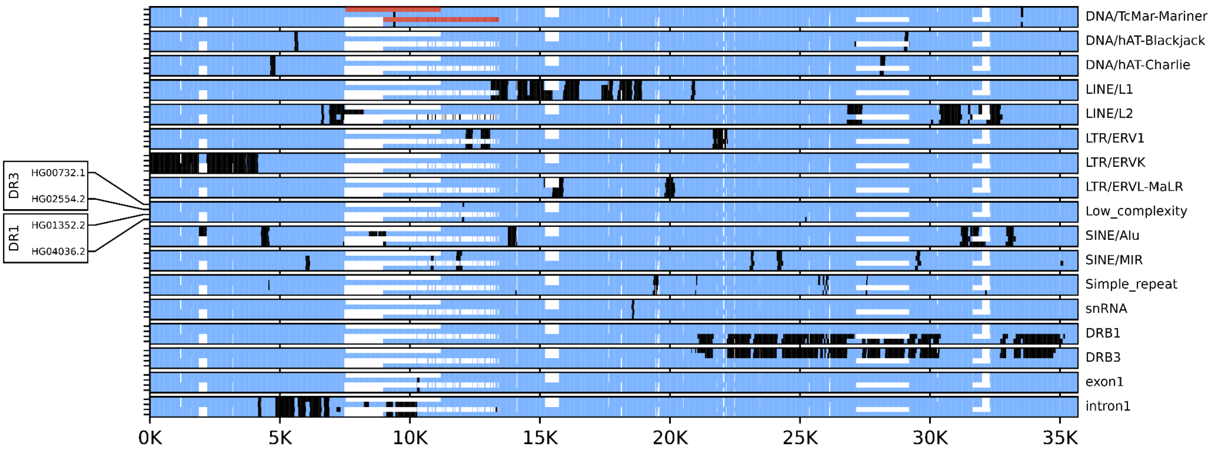


**Supplementary Figure 65. HLA SV insertion visualization.** Visualization of haplotype structures and homology for two different loci in the HLA-DR region, both of which were found to contain small (~ 3.7 kbp and 4.4 kbp) structural variants with solitary intron and exon *HLA-DRB* sequences (see main text). The figure shows that the identified structural variants as well as their surrounding sequence regions from different loci in the HLA-DR region exhibit a high degree of homology, and that the structural variants carry solitary *HLA-DRB* intron and exon sequences as well as repeat elements. Shown is a multiple sequence alignment (MSA) of four sequences; each panel shows the same MSA, annotated with a specific sequence feature (e.g., repeat element positions, homologies to HLA-DRB intron and exon sequences; sequence features are shown in black, white indicates gaps relative to the longest sequence alignment). The included four sequences comprise a) two representative DR3 haplotypes from the region around the identified structural variant in the *HLA-DRB9* region, one carrying the insertion and one not carrying the insertion; b) two representative DR1 haplotypes from the region around the identified structural variant in the region 10 kbp 3′ of *HLA-DRB1,* one carrying the insertion and one not carrying the insertion. The location of the identified structural variants (relative to other sequences from the same locus) is highlighted in red in the top panel.

66. Porubsky D, Harvey WT, Rozanski AN, Ebler J, Höps W, Ashraf H, Hasenfeld P, Human

Pangenome Reference Consortium (HPRC), Human Genome Structural Variation Consortium (HGSVC), Paten B, Sanders AD, Marschall T, Korbel JO, Eichler EE. Inversion polymorphism in a complete human genome assembly. *Genome Biol*. 2023;24:100.

80. Fu Q, Li H, Moorjani P, Jay F, Slepchenko SM, Bondarev AA, Johnson PLF, Aximu-Petri A, Prüfer K, de Filippo C, Meyer M, Zwyns N, Salazar-García DC, Kuzmin YV, Keates SG, Kosintsev PA, Razhev DI, Richards MP, Peristov NV, Lachmann M, Douka K, Higham TFG, Slatkin M, Hublin J-J, Reich D, Kelso J, Viola TB, Pääbo S. Genome

sequence of a 45,000-year-old modern human from western Siberia. *Nature*. 2014;514:445–449.

90. Baran Y, Subramaniam M, Biton A, Tukiainen T, Tsang EK, Rivas MA, Pirinen M, Gutierrez-Arcelus M, Smith KS, Kukurba KR, Zhang R, Eng C, Torgerson DG, Urbanek C, GTEx Consortium, Li JB, Rodriguez-Santana JR, Burchard EG, Seibold MA, MacArthur DG, Montgomery SB, Zaitlen NA, Lappalainen T. The landscape of genomic

imprinting across diverse adult human tissues. *Genome Res*. 2015;25:927–936.

91. Glenn CC, Driscoll DJ, Yang TP, Nicholls RD. Genomic imprinting: potential function and mechanisms revealed by the Prader-Willi and Angelman syndromes. *Mol Hum Reprod*. 1997;3:321–332.

92. Meguro M, Mitsuya K, Nomura N, Kohda M, Kashiwagi A, Nishigaki R, Yoshioka H, Nakao M, Oishi M, Oshimura M. Large-scale evaluation of imprinting status in the Prader-Willi syndrome region: an imprinted direct repeat cluster resembling small nucleolar RNA genes. *Hum Mol Genet*. 2001;10:383–394.

93. Tilgner H, Grubert F, Sharon D, Snyder MP. Defining a personal, allele-specific, and single-molecule long-read transcriptome. *Proc Natl Acad Sci U S A*. 2014;111:9869–9874.

94. O'Leary NA, Wright MW, Brister JR, Ciufo S, Haddad D, McVeigh R, Rajput B, Robbertse B, Smith-White B, Ako-Adjei D, Astashyn A, Badretdin A, Bao Y, Blinkova O, Brover V, Chetvernin V, Choi J, Cox E, Ermolaeva O, Farrell CM, Goldfarb T, Gupta T, Haft D, Hatcher E, Hlavina W, Joardar VS, Kodali VK, Li W, Maglott D, Masterson P, McGarvey KM, Murphy MR, O'Neill K, Pujar S, Rangwala SH, Rausch D, Riddick LD, Schoch C, Shkeda A, Storz SS, Sun H, Thibaud-Nissen F, Tolstoy I, Tully RE, Vatsan AR, Wallin C, Webb D, Wu W, Landrum MJ, Kimchi A, Tatusova T, DiCuccio M, Kitts P, Murphy TD, Pruitt KD. Reference sequence (RefSeq) database at NCBI: current status, taxonomic expansion, and functional annotation. *Nucleic Acids Res*. 2016;44:D733–45.

100. ENCODE Project Consortium, Moore JE, Purcaro MJ, Pratt HE, Epstein CB, Shoresh N, Adrian J, Kawli T, Davis CA, Dobin A, Kaul R, Halow J, Van Nostrand EL, Freese P, Gorkin DU, Shen Y, He Y, Mackiewicz M, Pauli-Behn F, Williams BA, Mortazavi A, Keller CA, Zhang X-O, Elhajjajy SI, Huey J, Dickel DE, Snetkova V, Wei X, Wang X, Rivera-Mulia JC, Rozowsky J, Zhang J, Chhetri SB, Zhang J, Victorsen A, White KP, Visel A, Yeo GW, Burge CB, Lécuyer E, Gilbert DM, Dekker J, Rinn J, Mendenhall EM,

Ecker JR, Kellis M, Klein RJ, Noble WS, Kundaje A, Guigó R, Farnham PJ, Cherry JM, Myers RM, Ren B, Graveley BR, Gerstein MB, Pennacchio LA, Snyder MP, Bernstein BE, Wold B, Hardison RC, Gingeras TR, Stamatoyannopoulos JA, Weng Z. Expanded encyclopaedias of DNA elements in the human and mouse genomes. *Nature*. 2020;583:699–710.

111. Dixon JR, Selvaraj S, Yue F, Kim A, Li Y, Shen Y, Hu M, Liu JS, Ren B. Topological domains in mammalian genomes identified by analysis of chromatin interactions.

*Nature*. 2012;485:376–380.

123. van der Meer D, Kaufmann T, Shadrin AA, Makowski C, Frei O, Roelfs D, Monereo-Sánchez J, Linden DEJ, Rokicki J, Alnæs D, de Leeuw C, Thompson WK, Loughnan R, Fan CC, Westlye LT, Andreassen OA, Dale AM. The genetic architecture

of human cortical folding. *Sci Adv*. 2021;7:eabj9446.

124. Sakaue S, Kanai M, Tanigawa Y, Karjalainen J, Kurki M, Koshiba S, Narita A, Konuma T, Yamamoto K, Akiyama M, Ishigaki K, Suzuki A, Suzuki K, Obara W, Yamaji K, Takahashi K, Asai S, Takahashi Y, Suzuki T, Shinozaki N, Yamaguchi H, Minami S, Murayama S, Yoshimori K, Nagayama S, Obata D, Higashiyama M, Masumoto A, Koretsune Y, FinnGen, Ito K, Terao C, Yamauchi T, Komuro I, Kadowaki T, Tamiya G, Yamamoto M, Nakamura Y, Kubo M, Murakami Y, Yamamoto K, Kamatani Y, Palotie A, Rivas MA, Daly MJ, Matsuda K, Okada Y. A cross-population atlas of genetic associations for 220 human phenotypes. *Nat Genet*. 2021;53:1415–1424.

143. Moutsianas L, Jostins L, Beecham AH, Dilthey AT, Xifara DK, Ban M, Shah TS, Patsopoulos NA, Alfredsson L, Anderson CA, Attfield KE, Baranzini SE, Barrett J, Binder TMC, Booth D, Buck D, Celius EG, Cotsapas C, D'Alfonso S, Dendrou CA, Donnelly P, Dubois B, Fontaine B, Fugger L, Goris A, Gourraud P-A, Graetz C, Hemmer B, Hillert J, International IBD Genetics Consortium (IIBDGC), Kockum I, Leslie S, Lill CM, Martinelli-Boneschi F, Oksenberg JR, Olsson T, Oturai A, Saarela J, Søndergaard HB, Spurkland A, Taylor B, Winkelmann J, Zipp F, Haines JL, Pericak-Vance MA, Spencer CCA, Stewart G, Hafler DA, Ivinson AJ, Harbo HF, Hauser SL, De Jager PL, Compston A, McCauley JL, Sawcer S, McVean G. Class II HLA interactions modulate genetic risk for multiple sclerosis. *Nat Genet*. 2015;47:1107–1113.

144. Dilthey AT. State-of-the-art genome inference in the human MHC. *Int J Biochem Cell Biol*. 2021;131:105882.

145. Zhou Y, Song L, Li H. Full resolution HLA and KIR gene annotations for human genome assemblies. *Genome Res* [Internet]. 2024;Available from: http://dx.doi.org/10.1101/gr.278985.124

146. Barker DJ, Maccari G, Georgiou X, Cooper MA, Flicek P, Robinson J, Marsh SGE. The IPD-IMGT/HLA Database. *Nucleic Acids Res*. 2023;51:D1053–D1060.

147. Mentzer AJ, Dilthey AT, Pollard M, Gurdasani D, Karakoc E, Carstensen T, Muhwezi A, Cutland C, Diarra A, da Silva Antunes R, Paul S, Smits G, Wareing S, Kim H, Pomilla C, Chong AY, Brandt DYC, Nielsen R, Neaves S, Timpson N, Crinklaw A, Lindestam Arlehamn CS, Rautanen A, Kizito D, Parks T, Auckland K, Elliott KE, Mills T, Ewer K, Edwards N, Fatumo S, Webb E, Peacock S, Jeffery K, van der Klis FRM, Kaleebu P, Vijayanand P, Peters B, Sette A, Cereb N, Sirima S, Madhi SA, Elliott AM, McVean G, Hill AVS, Sandhu MS. High-resolution African HLA resource uncovers HLA-DRB1 expression effects underlying vaccine response. *Nat Med*. 2024;30:1384–1394.

158. Yang Y, Chung EK, Wu YL, Savelli SL, Nagaraja HN, Zhou B, Hebert M, Jones KN, Shu Y, Kitzmiller K, Blanchong CA, McBride KL, Higgins GC, Rennebohm RM, Rice RR,

Hackshaw KV, Roubey RAS, Grossman JM, Tsao BP, Birmingham DJ, Rovin BH, Hebert LA, Yu CY. Gene copy-number variation and associated polymorphisms of complement component C4 in human systemic lupus erythematosus (SLE): low copy number is a risk factor for and high copy number is a protective factor against SLE susceptibility in European Americans. *Am J Hum Genet*. 2007;80:1037–1054.

170. Mc Cartney AM, Shafin K, Alonge M, Bzikadze AV, Formenti G, Fungtammasan A, Howe K, Jain C, Koren S, Logsdon GA, Miga KH, Mikheenko A, Paten B, Shumate A,

Soto DC, Sović I, Wood JMD, Zook JM, Phillippy AM, Rhie A. Chasing perfection: validation and polishing strategies for telomere-to-telomere genome assemblies. *Nat Methods*. 2022;19:687–695.

184. Usher CL, Handsaker RE, Esko T, Tuke MA, Weedon MN, Hastie AR, Cao H, Moon JE, Kashin S, Fuchsberger C, Metspalu A, Pato CN, Pato MT, McCarthy MI, Boehnke M, Altshuler DM, Frayling TM, Hirschhorn JN, McCarroll SA. Structural forms of the

human amylase locus and their relationships to SNPs, haplotypes and obesity. *Nat Genet*. 2015;47:921–925.

190. Schmutz J, Martin J, Terry A, Couronne O, Grimwood J, Lowry S, Gordon LA, Scott D, Xie G, Huang W, Hellsten U, Tran-Gyamfi M, She X, Prabhakar S, Aerts A, Altherr M, Bajorek E, Black S, Branscomb E, Caoile C, Challacombe JF, Chan YM, Denys M, Detter JC, Escobar J, Flowers D, Fotopulos D, Glavina T, Gomez M, Gonzales E, Goodstein D, Grigoriev I, Groza M, Hammon N, Hawkins T, Haydu L, Israni S, Jett J, Kadner K, Kimball H, Kobayashi A, Lopez F, Lou Y, Martinez D, Medina C, Morgan J, Nandkeshwar R, Noonan JP, Pitluck S, Pollard M, Predki P, Priest J, Ramirez L, Retterer J, Rodriguez A, Rogers S, Salamov A, Salazar A, Thayer N, Tice H, Tsai M, Ustaszewska A, Vo N, Wheeler J, Wu K, Yang J, Dickson M, Cheng J-F, Eichler EE, Olsen A, Pennacchio LA, Rokhsar DS, Richardson P, Lucas SM, Myers RM, Rubin EM. The DNA sequence and comparative analysis of human chromosome 5. *Nature*. 2004;431:268–274.

191. R Core Team. R: A language and environment for statistical computing [Internet]. Vienna, Austria: R Foundation for Statistical Computing; 2020. Available from: https://www.R-project.org/

192. Wickham H. Ggplot2: elegant graphics for data analysis. New York: Springer; 2009.

193. Vollger MR, Kerpedjiev P, Phillippy AM, Eichler EE. StainedGlass: interactive visualization of massive tandem repeat structures with identity heatmaps. *Bioinformatics*. 2022;38:2049–2051.

194. Altemose N, Logsdon GA, Bzikadze AV, Sidhwani P, Langley SA, Caldas GV, Hoyt SJ, Uralsky L, Ryabov FD, Shew CJ, Sauria MEG, Borchers M, Gershman A, Mikheenko A, Shepelev VA, Dvorkina T, Kunyavskaya O, Vollger MR, Rhie A, McCartney AM, Asri M, Lorig-Roach R, Shafin K, Lucas JK, Aganezov S, Olson D, de Lima LG, Potapova T, Hartley GA, Haukness M, Kerpedjiev P, Gusev F, Tigyi K, Brooks S, Young A, Nurk S, Koren S, Salama SR, Paten B, Rogaev EI, Streets A, Karpen GH, Dernburg AF, Sullivan BA, Straight AF, Wheeler TJ, Gerton JL, Eichler EE, Phillippy AM, Timp W, Dennis MY, O'Neill RJ, Zook JM, Schatz MC, Pevzner PA, Diekhans M, Langley CH, Alexandrov IA, Miga KH. Complete genomic and epigenetic maps of

human centromeres. *Science*. 2022;376:eabl4178.

195. Gershman A, Sauria MEG, Guitart X, Vollger MR, Hook PW, Hoyt SJ, Jain M, Shumate A, Razaghi R, Koren S, Altemose N, Caldas GV, Logsdon GA, Rhie A, Eichler EE, Schatz MC, O'Neill RJ, Phillippy AM, Miga KH, Timp W. Epigenetic patterns in a complete human genome. *Science*. 2022;376:eabj5089.
